## Supplementary Figures 1-13 for "On the limits of fitting complex models of population history to genetic data"

Figure S1

**Figure S1.** The admixture graphs compared in **Figure 1a**.

**a**, graph 1 from **Figure 3a**, LL = 4.9, WR = 2.0 SE.

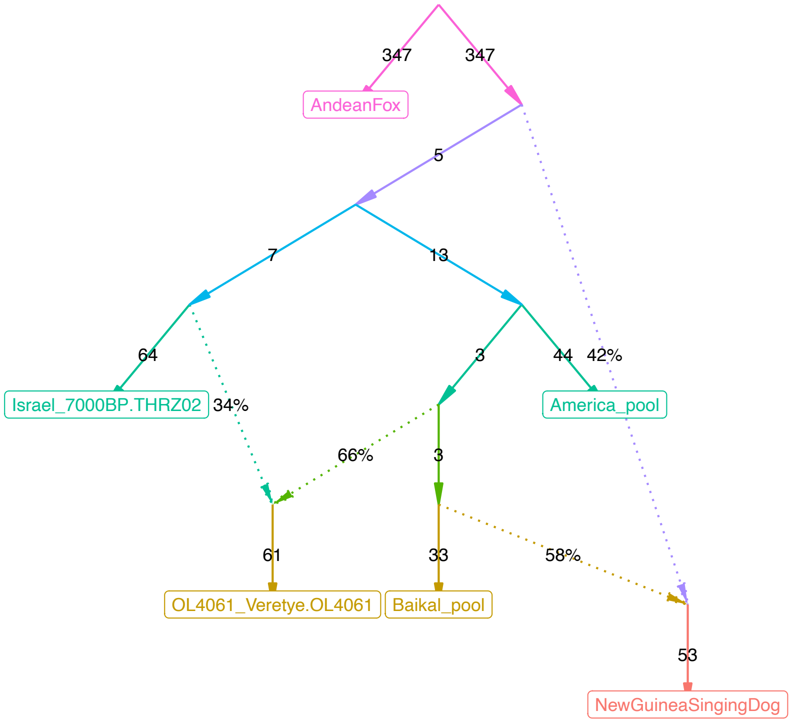

**b**, graph 2 from **Figure 3a**, LL = 25.7, WR = 5.0 SE.

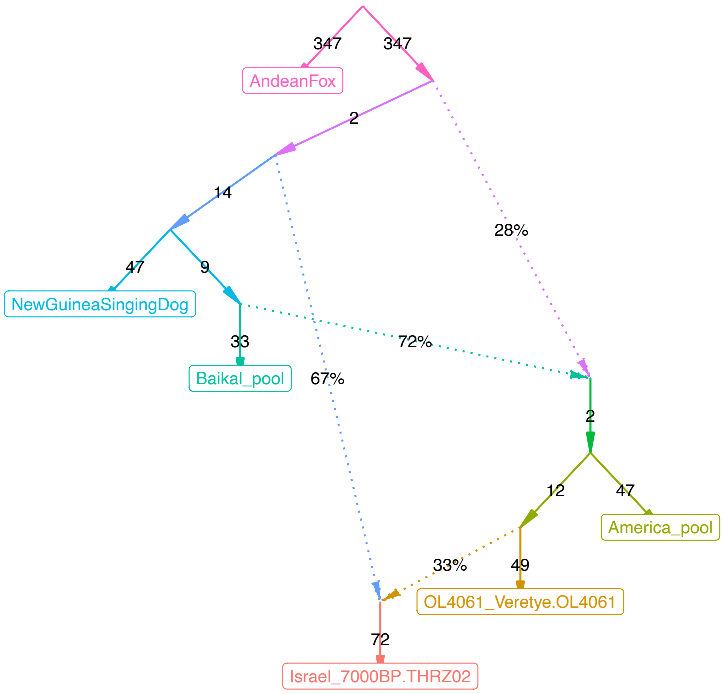

Figure S2

**Figure S2.** The published (Bergström et al. 2020) and alternative admixture graphs for dogs found with *findGraphs*.

**a**, published model

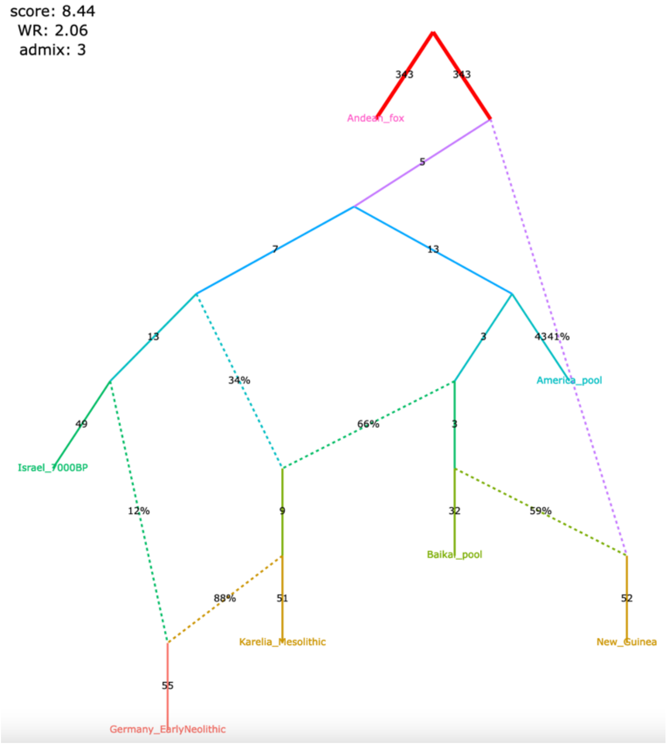

Figure S2

**c**, alternative models fitting nominally better than the published one, sorted by the fit score

the best model corresponding perfectly to the human history

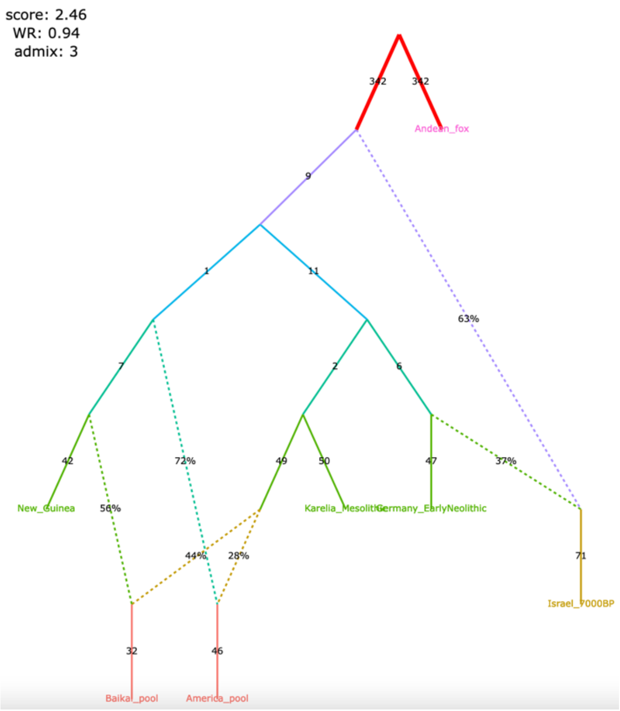

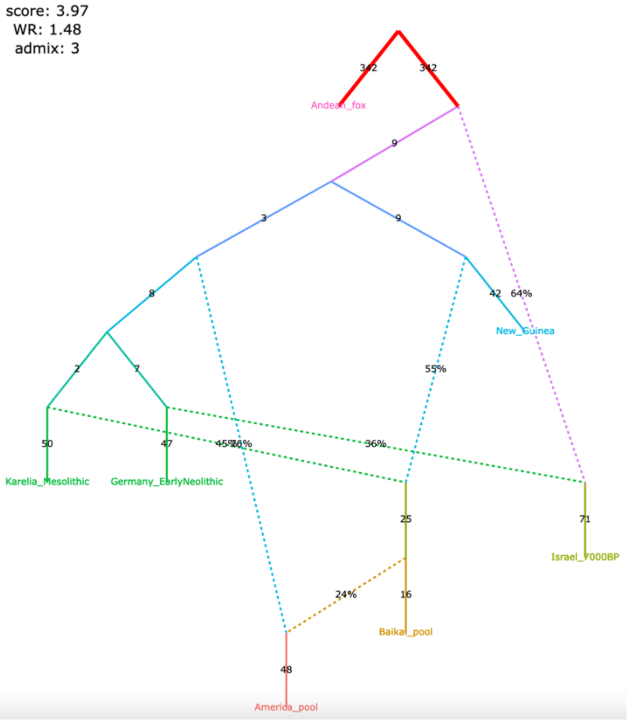

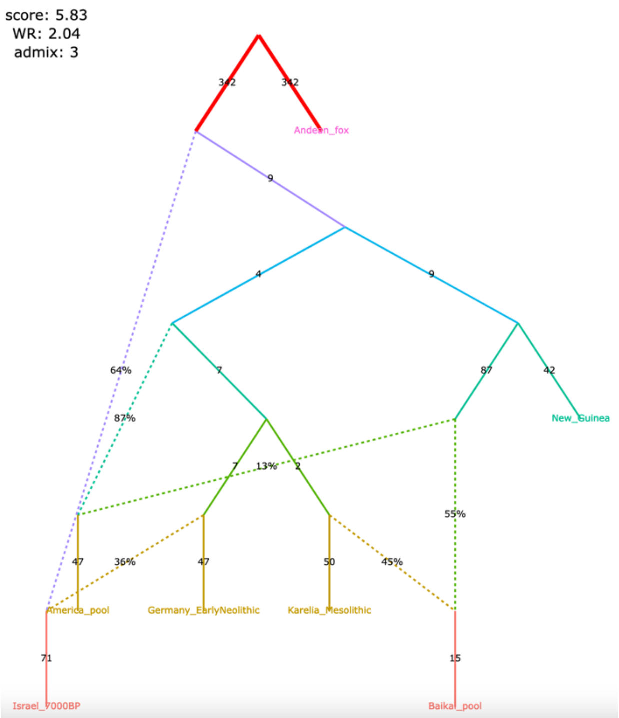

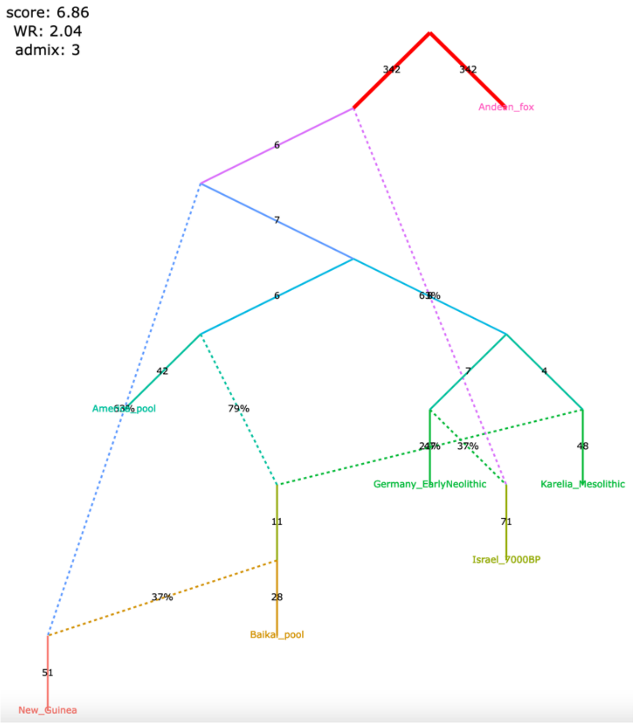

Figure S2

**d**, alternative models fitting nominally or significantly (the graph framed in blue) better than the published one, sorted by the fit score

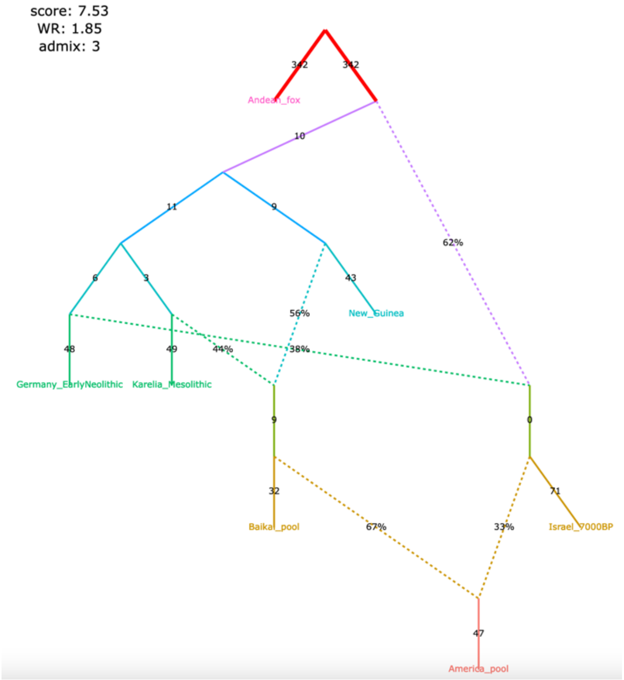

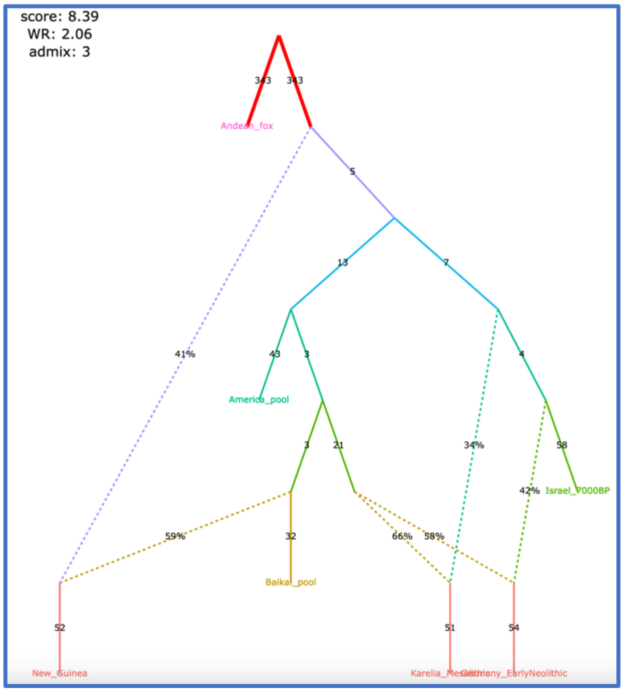

Figure S3

**Figure S3.** Alternative admixture graphs for humans found with *findGraphs* for the dataset from Bergström *et al*. (2020).

**a**, best-fitting models for humans sorted by the fit score

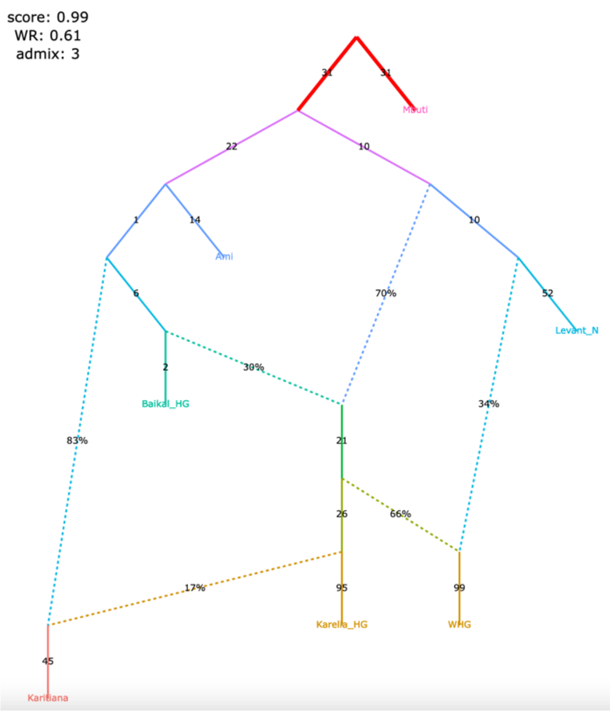

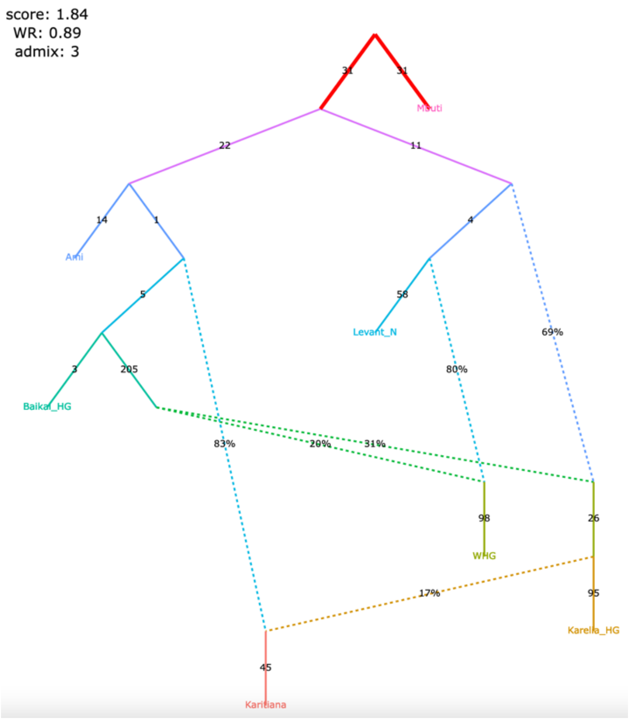

a model ideally corresponding to the best-fitting model for dogs

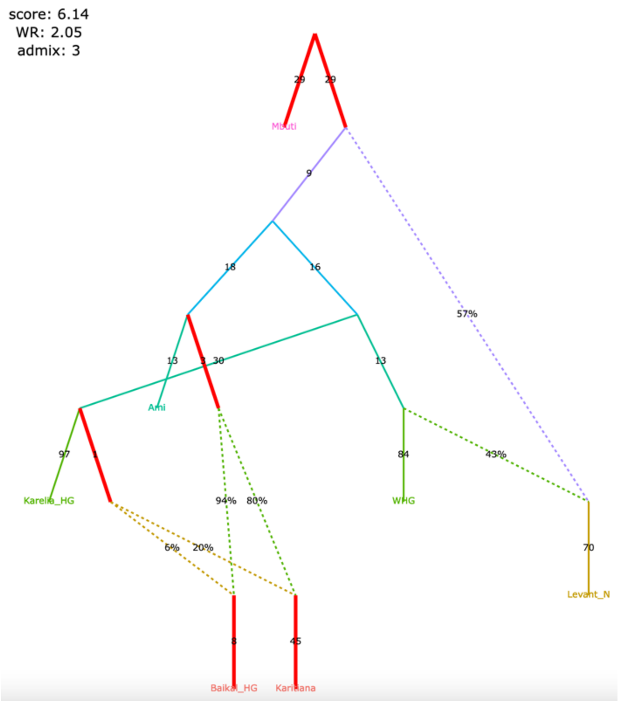

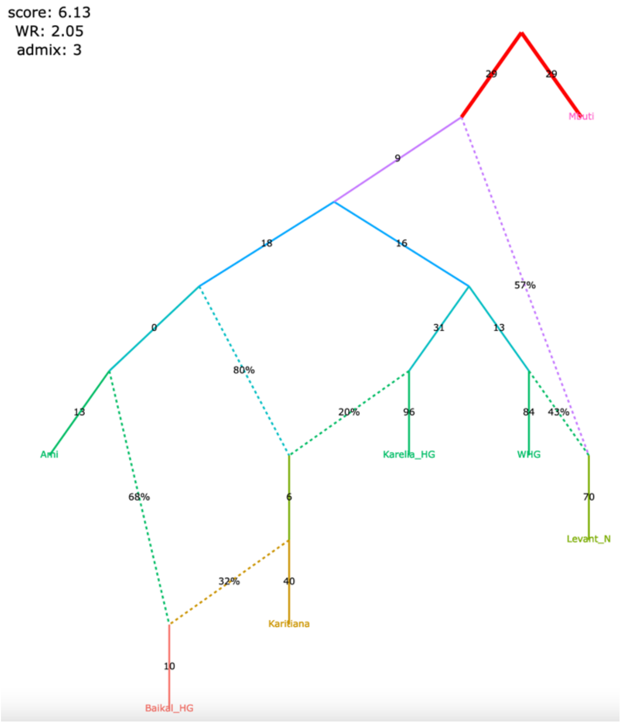

Figure S3

**b**, best-fitting models for humans sorted by the fit score

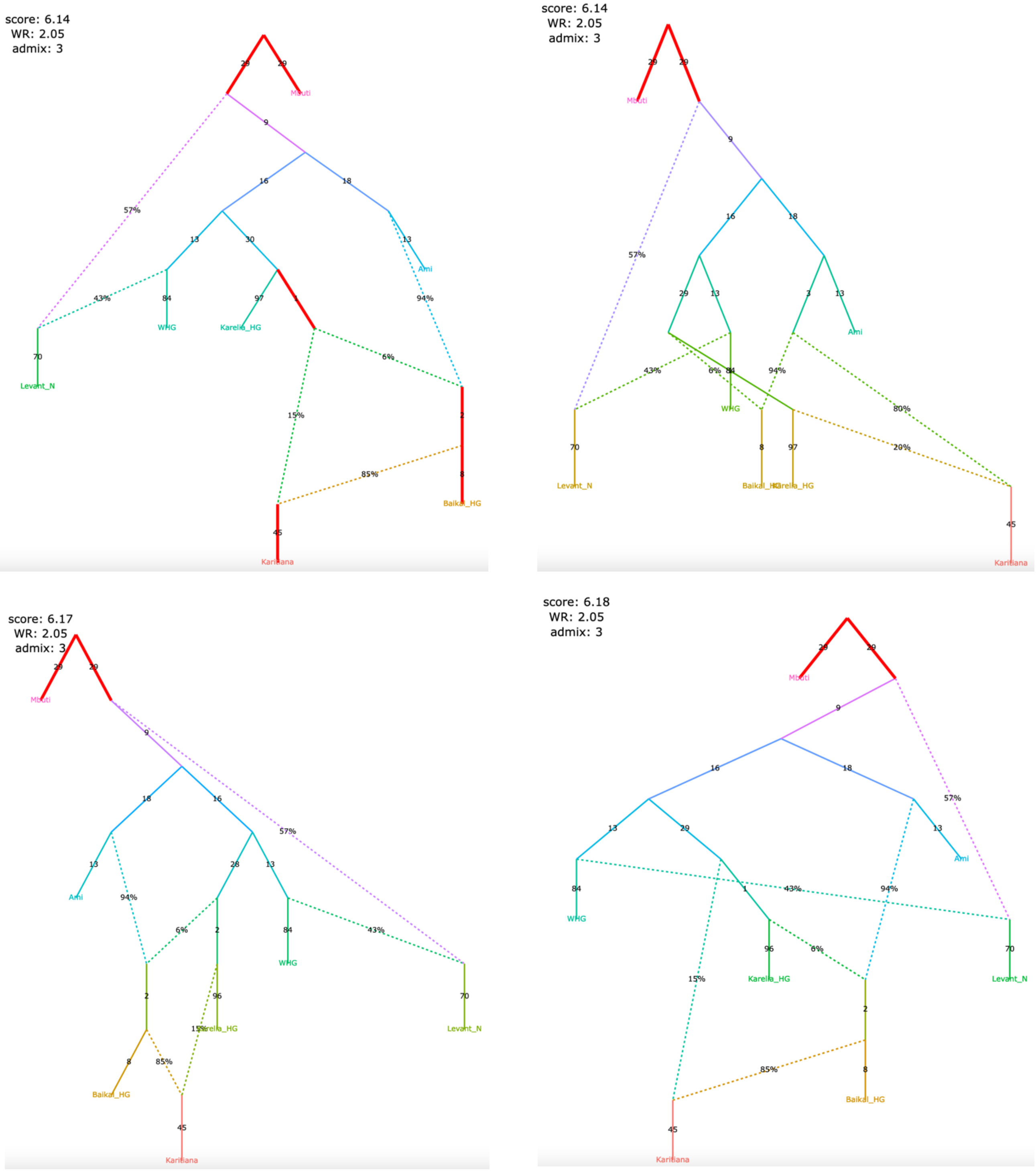

Figure S4

**Figure S4.** Published admixture graph from Lazaridis *et al*. (2014) and alternative graphs found with *findGraphs* (7 populations, 4 admixture events).

**a**, the re-fitted published graph

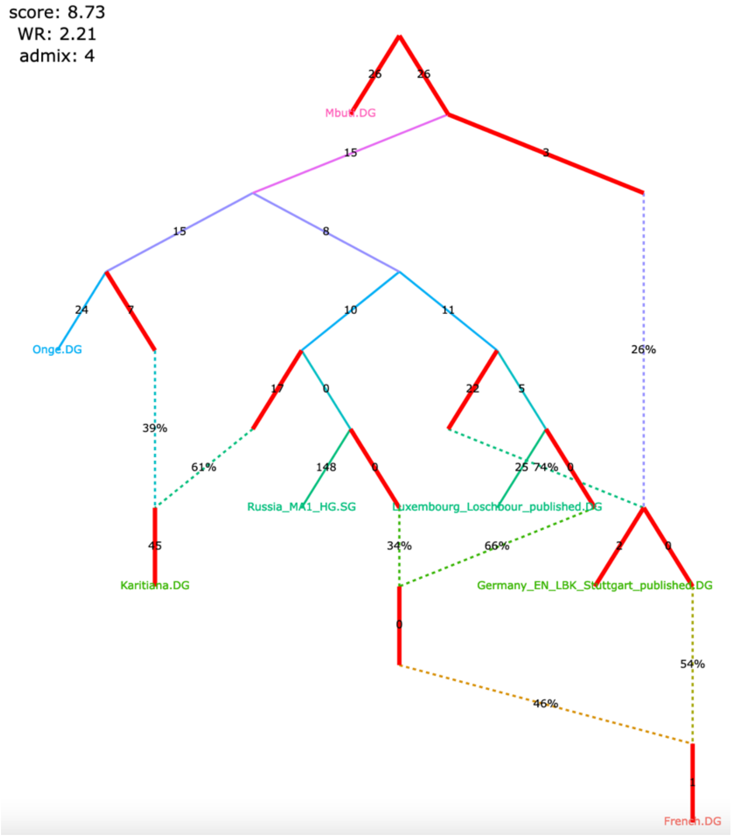

Figure S4

**b**, ten examples of graphs inferred by *findGraphs* (arranged according to LL score) and fitting significantly (the graph framed in blue) or nominally better than the published model.

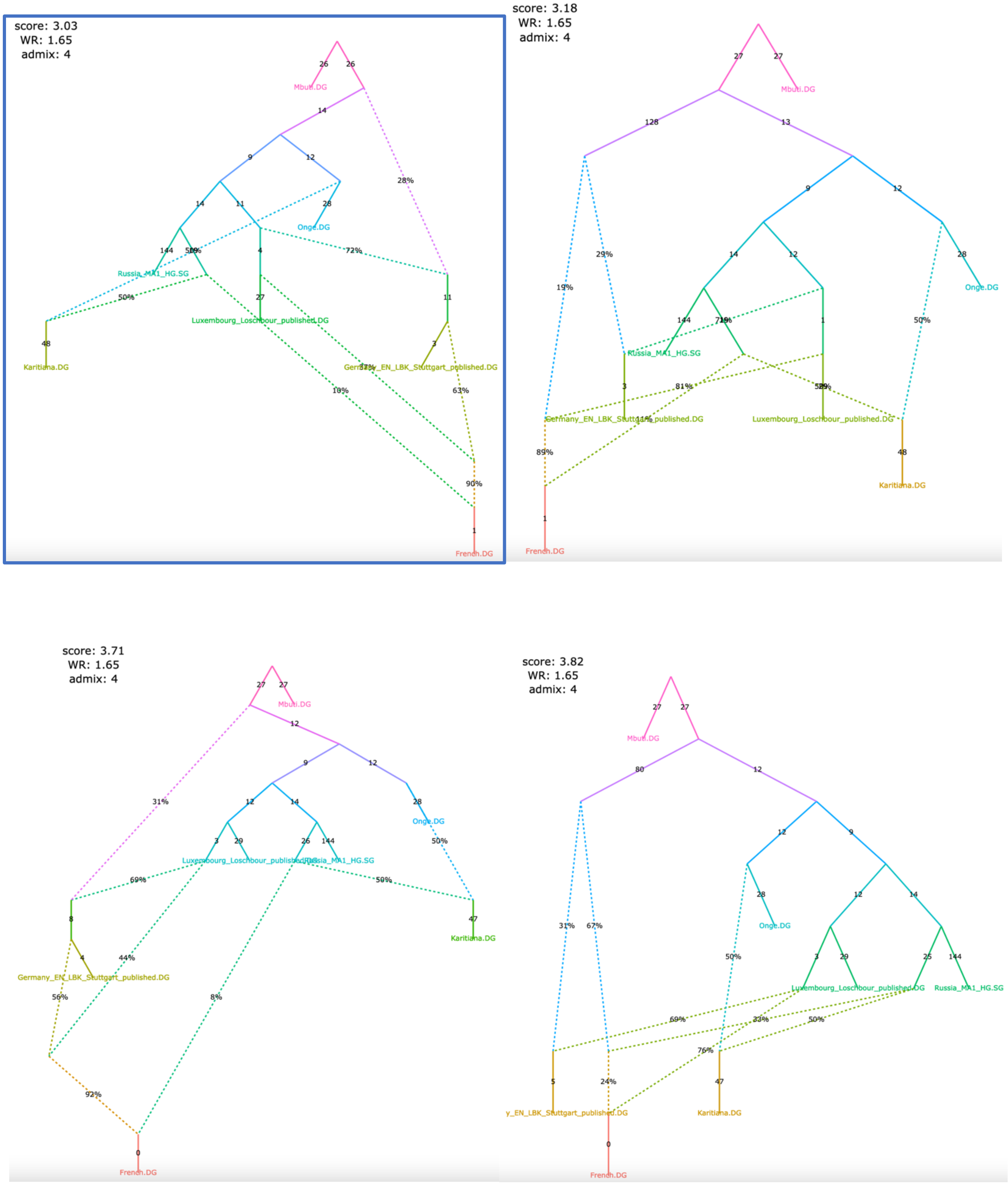

### Figure S4

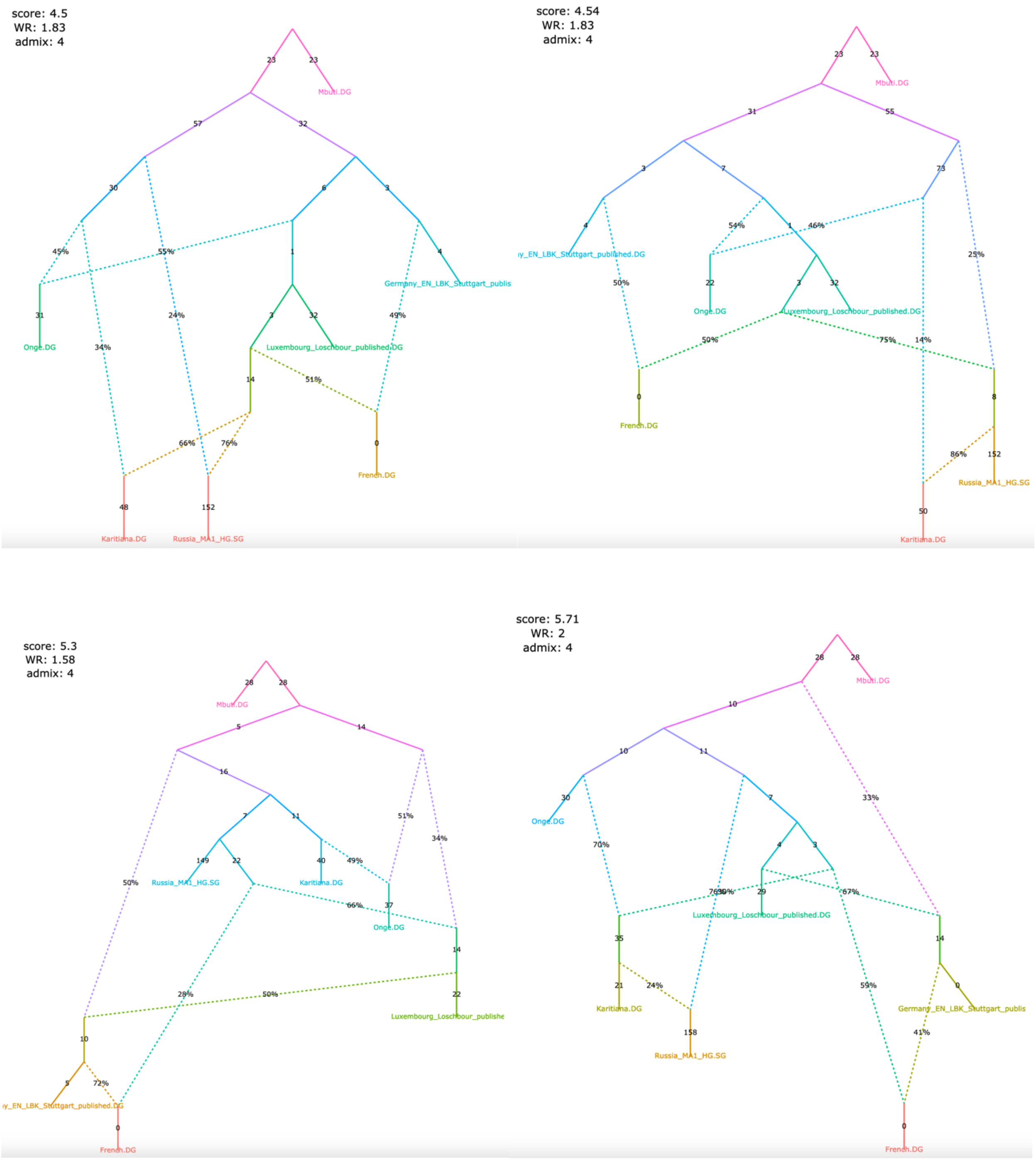

score: 4.5
WR: 1.83
admix: 4
Mbuti.DG
Onge.DG
Germany_EN_LBK_Stuttgart_publis
Luxembourg_Loschbour_published.DG
French.DG
Karitiana.DG
Russia_MA1_HG.SG
score: 4.54
WR: 1.83
admix: 4
Mbuti.DG
Onge.DG
Luxembourg_Loschbour_published.DG
French.DG
Russia_MA1_HG.SG
Karitiana.DG
score: 5.3
WR: 1.58
admix: 4
Mbuti.DG
Russia_MA1_HG.SG
Karitiana.DG
Onge.DG
Luxembourg_Loschbour_publishe
French.DG
score: 5.71
WR: 2
admix: 4
Mbuti.DG
Onge.DG
Luxembourg_Loschbour_published.DG
Karitiana.DG
Russia_MA1_HG.SG
Germany_EN_LBK_Stuttgart_publis
French.DG

Figure S4

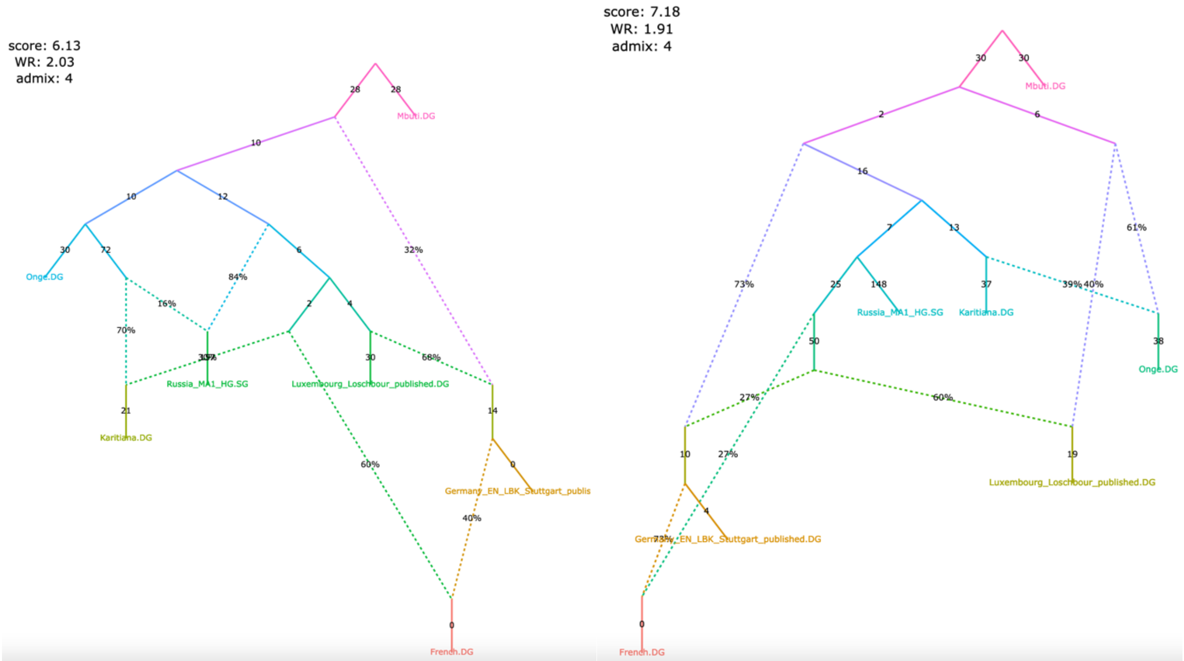

score: 6.13
WR: 2.03
admix: 4
Mbuti.DG
Onge.DG
Russia_MA1_HG.SG
Luxembourg_Loschbour_published.DG
Karitiana.DG
Germany_EN_LBK_Stuttgart_publis
French.DG
score: 7.18
WR: 1.91
admix: 4
Mbuti.DG
Russia_MA1_HG.SG
Karitiana.DG
Onge.DG
Luxembourg_Loschbour_published.DG
Germany_EN_LBK_Stuttgart_published.DG
French.DG

Figure S5

**Figure S5.** Published admixture graph from Shinde *et al*. (2019) and alternative graphs found with *findGraphs* (8 populations, 3 admixture events) relying on the original set of SNPs, original group composition, and original (incorrect) algorithm for calculating *f*-statistics.

**a**, published model; the original set of SNPs and individuals, and the original algorithm for calculating $f_3$-statistics was used (470,389 variable site with no missing data at the group level available)

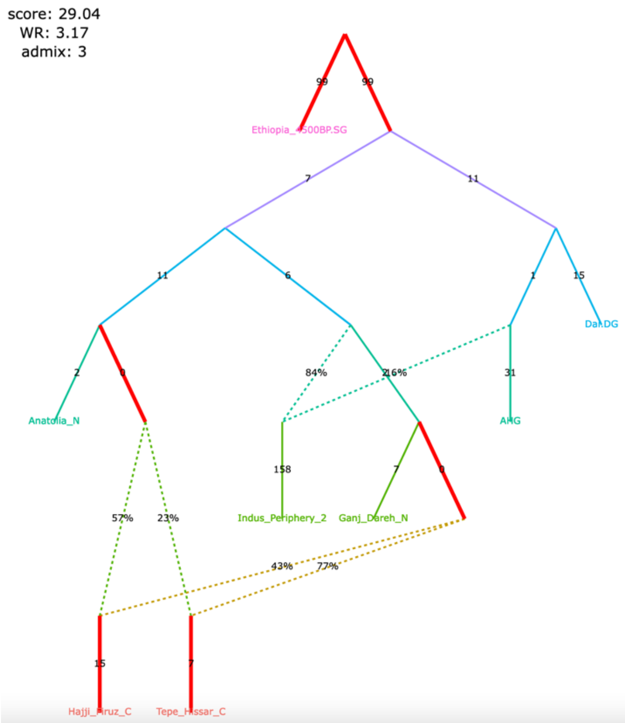

The claim by Shinde et al. 2019 relying on the admixture graph:

Primary ancestry in the Indus Periphery group forms the deepest branch in the Iranian Neolithic clade composed of the Indus Periphery, Ganj Dareh Neolithic, Hajji Firuz Chalcolithic and Tepe Hissar Chalcolithic groups.

Figure S5

**b**, selected alternative models fitting significantly better (graphs framed in blue), nominally better (graphs without frames), or not significantly worse (graphs framed in red) than the published one

Indus Periphery is not the deepest branch in the Iranian Neolithic clade

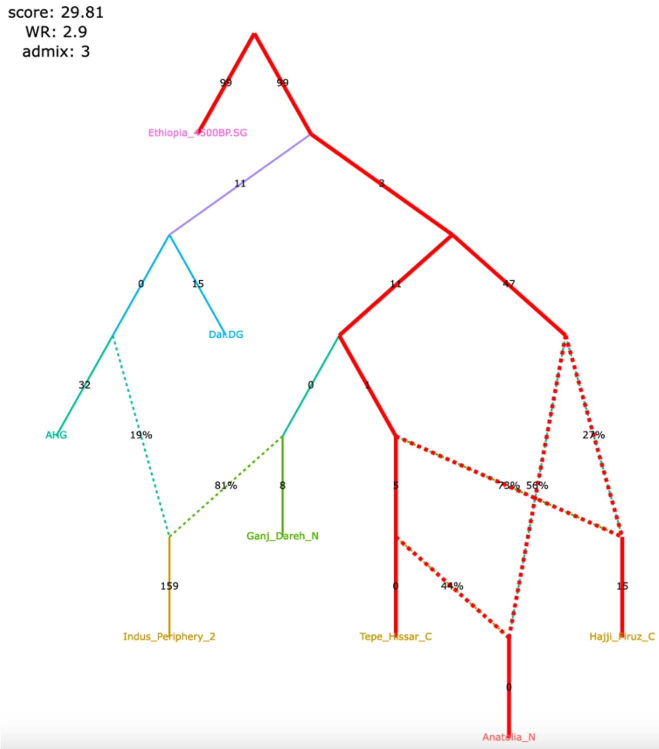

Indus Periphery is not the deepest branch in the Iranian Neolithic clade

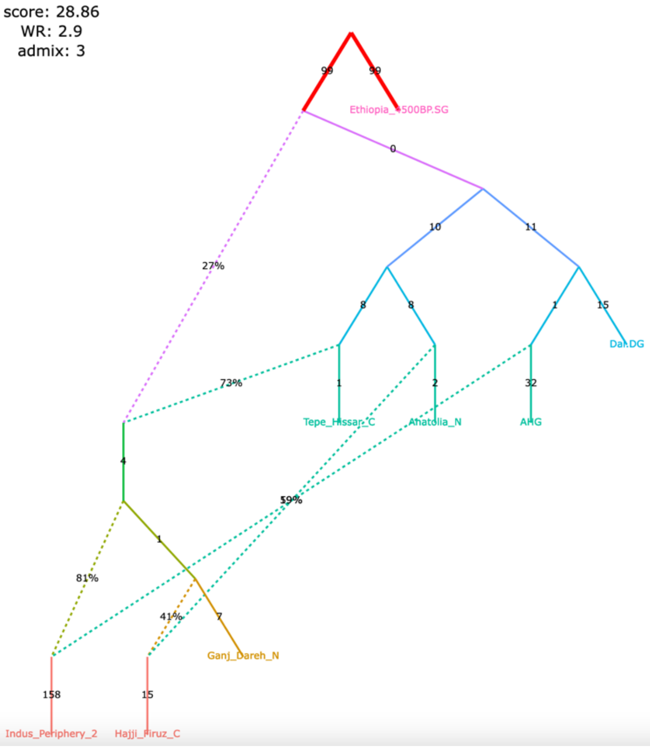

Indus Periphery is not the deepest branch in the Iranian Neolithic clade

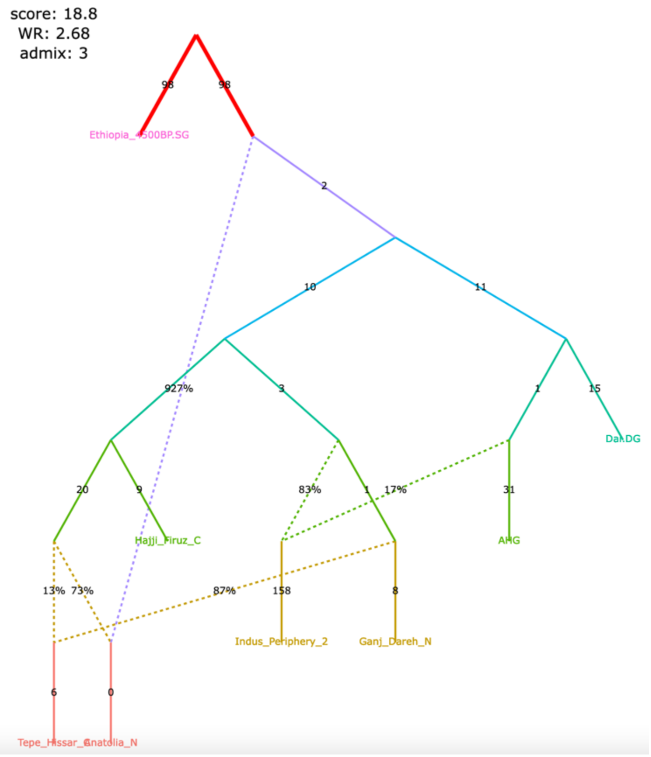

Indus Periphery is not the deepest branch in the Iranian Neolithic clade

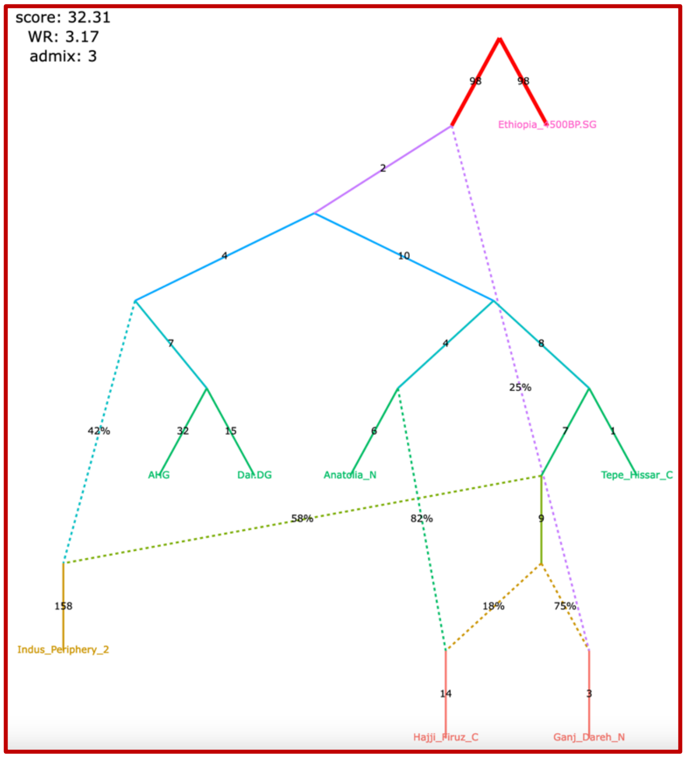

Figure S5

**c**, selected alternative models fitting significantly better (graphs framed in blue), nominally better (graphs without frames), or not significantly worse (graphs framed in red) than the published one

Indus Periphery is not the deepest branch in the Iranian Neolithic clade

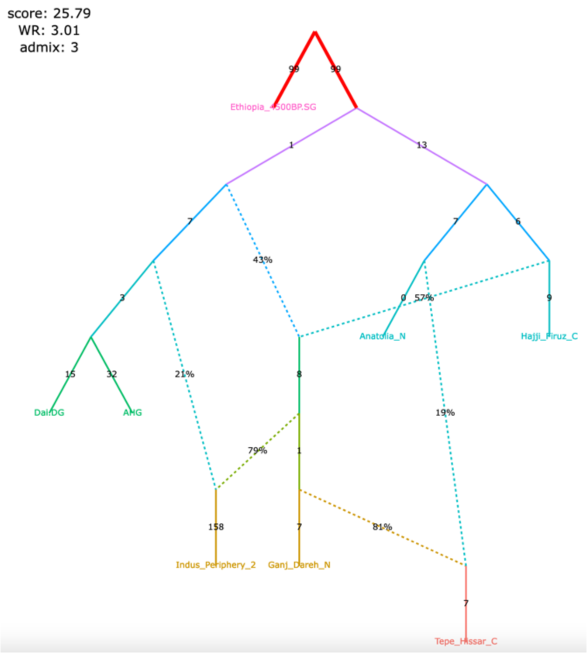

Indus Periphery is not the deepest branch in the Iranian Neolithic clade

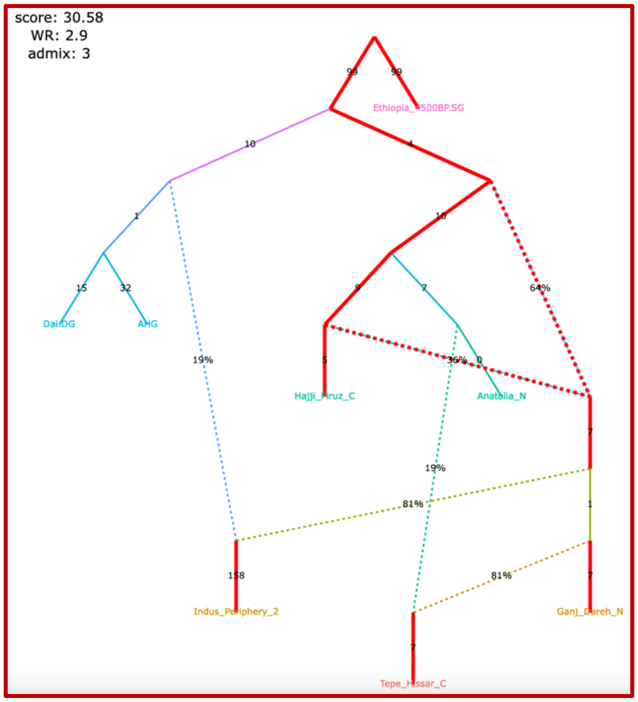

Indus Periphery is not the deepest branch in the Iranian Neolithic clade

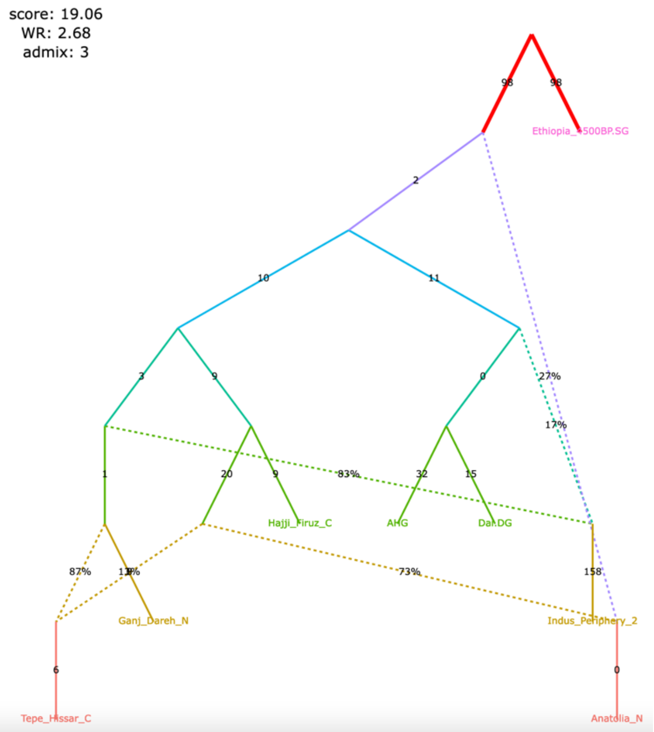

Indus Periphery is not the deepest branch in the Iranian Neolithic clade

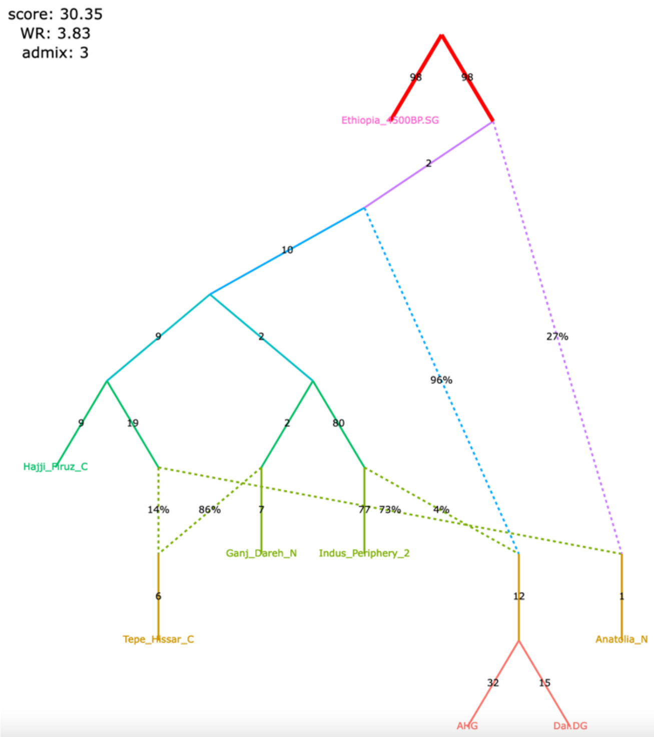

Figure S5

**d**, selected alternative models fitting significantly better (graphs framed in blue), nominally better (graphs without frames), or not significantly worse (graphs framed in red) than the published one

Primary ancestry in Indus Periphery is a basal Asian branch

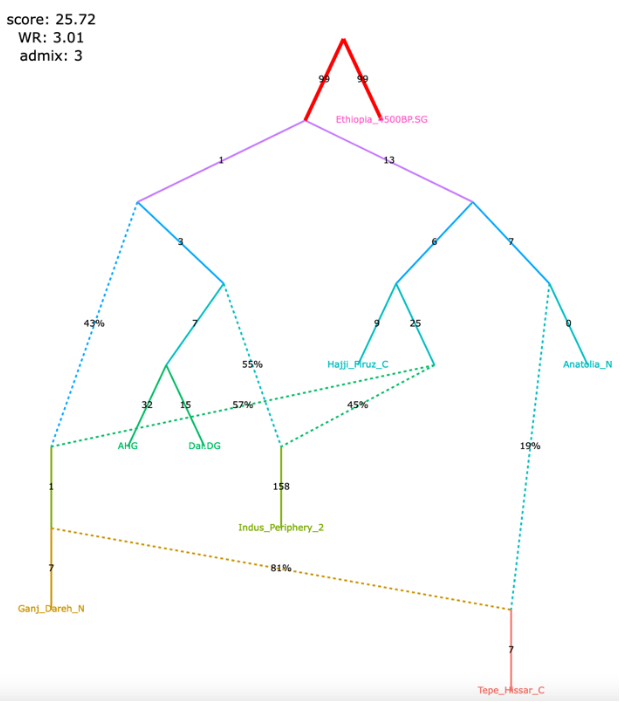

Primary ancestry in Indus Periphery is a basal Asian branch

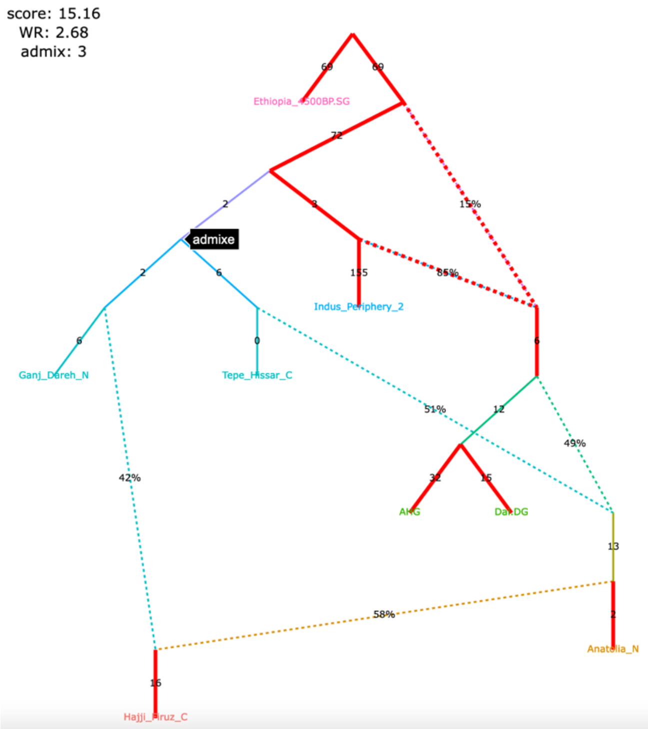

Primary ancestry in Indus Periphery is a basal Asian branch

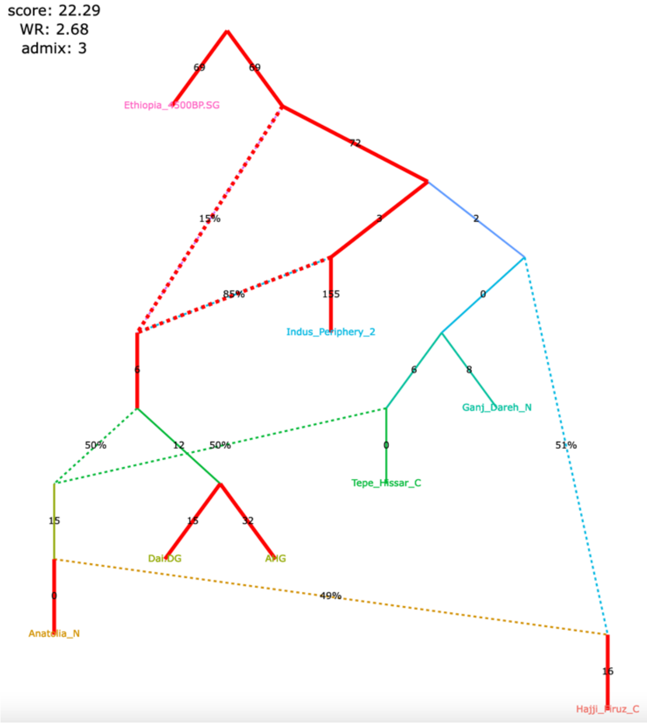

Primary ancestry in Indus Periphery is a basal West Eurasian branch

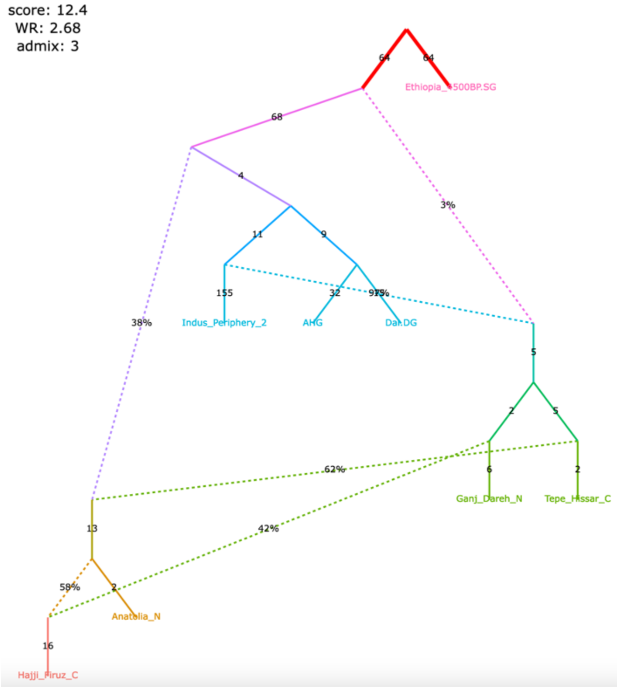

Figure S5

**e**, selected alternative models fitting significantly better (graphs framed in blue), nominally better (graphs without frames), or not significantly worse (graphs framed in red) than the published one

Indus Periphery is the deepest branch in the Iranian Neolithic clade

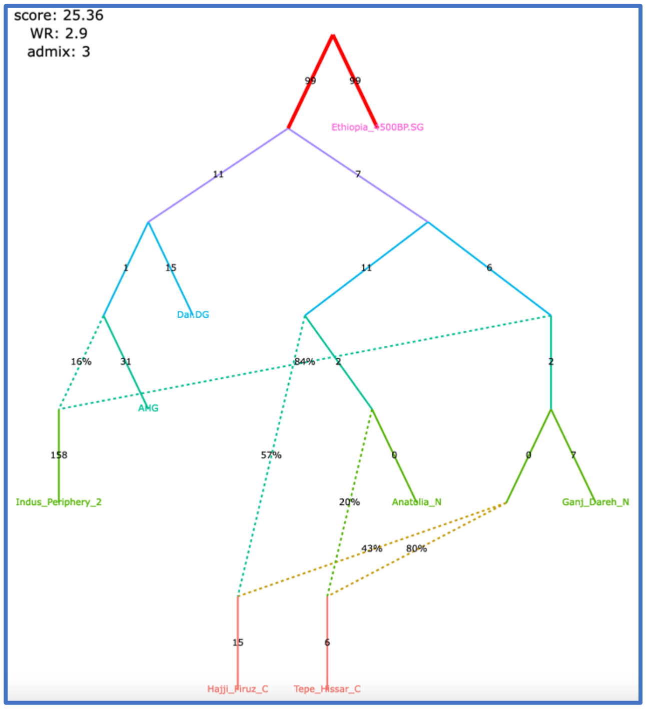

Indus Periphery is the deepest branch in the Iranian Neolithic clade

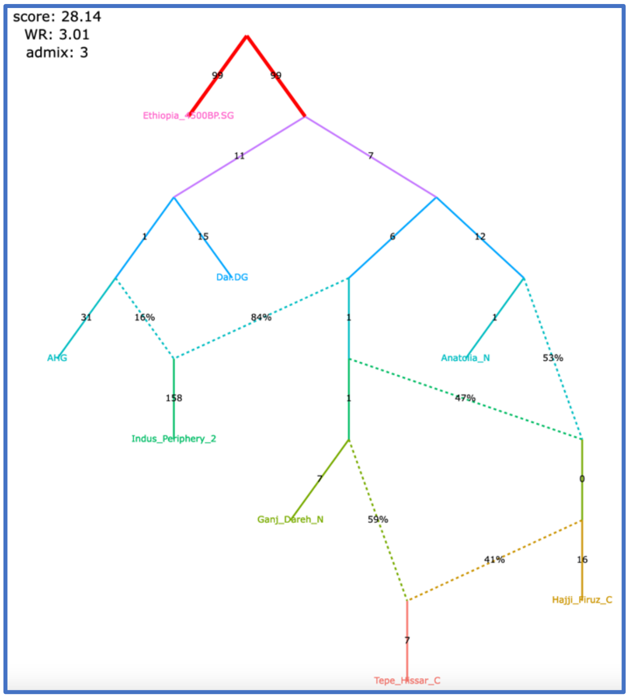

Indus Periphery is the deepest branch in the Iranian Neolithic clade

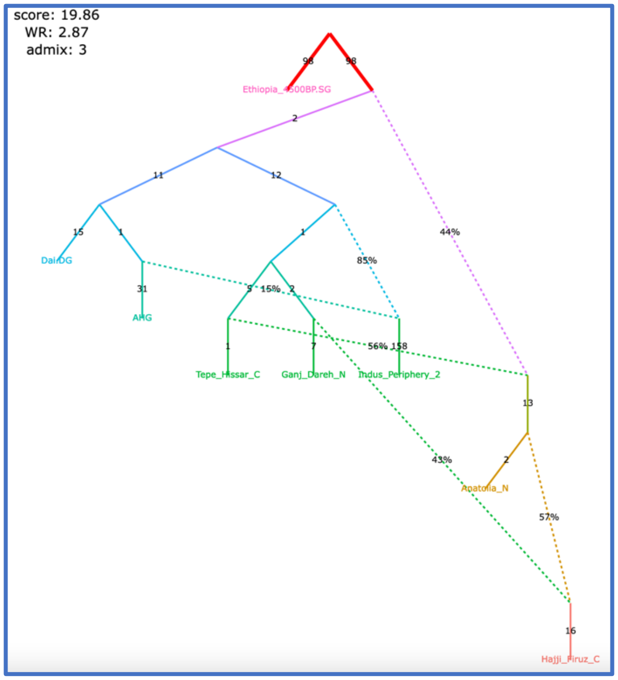

Indus Periphery is the deepest branch in the Iranian Neolithic clade

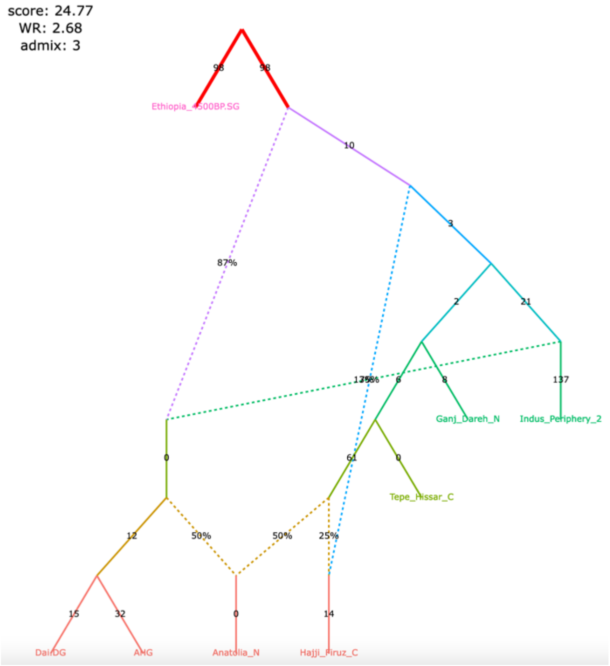

Figure S5

**f**, selected alternative models fitting significantly better (graphs framed in blue), nominally better (graphs without frames), or not significantly worse (graphs framed in red) than the published one

Indus Periphery is the deepest branch in the Iranian Neolithic clade

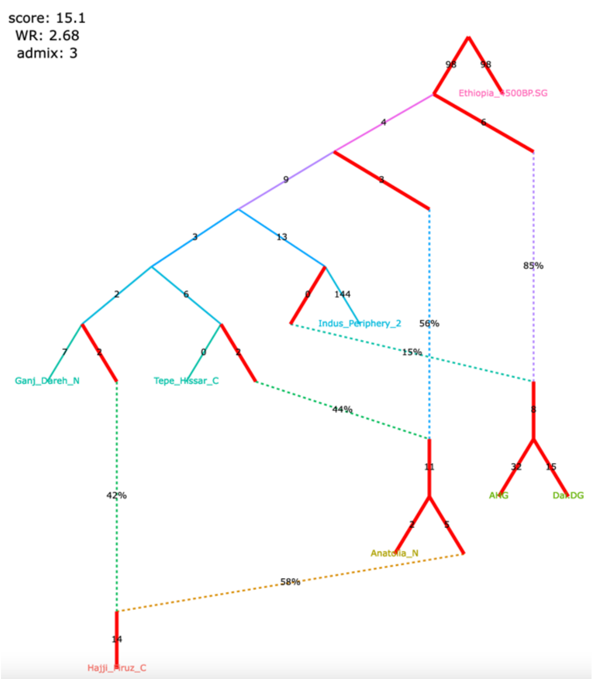

Indus Periphery is the deepest branch in the Iranian Neolithic clade

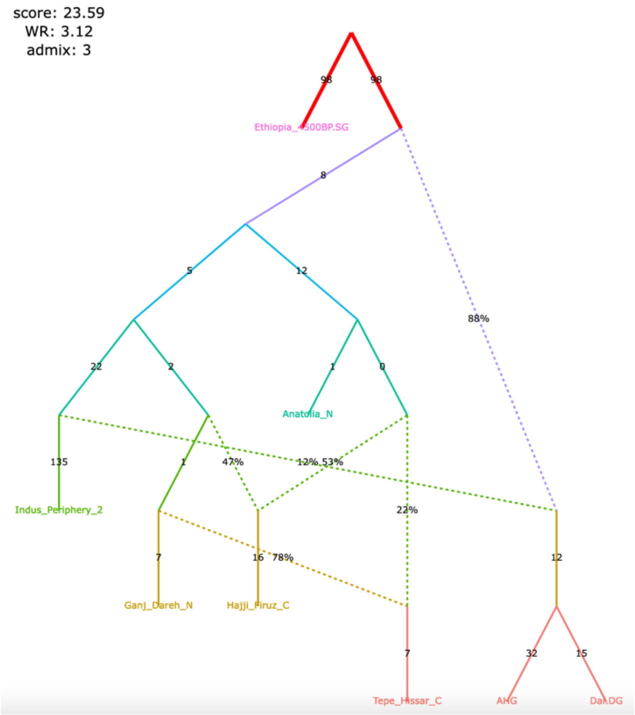

Figure S6

**Figure S6.** Published admixture graph from Shinde *et al*. (2019) and alternative graphs found with *findGraphs* (8 populations, 3 admixture events) for the modified group composition and using the updated algorithm for calculating *f*-statistics. The graphs were also re-fitted on the original set of SNPs/individuals and using the original algorithm for calculating *f*-statistics.

**a**, published model with 3 admixture events

| Relatives (2nd and 3rd degree) and potentially contaminated individuals removed, the Indus Periphery group expanded to 4 ind., the corrected algorithm for calculating $f_3$-statistics (249,009 variable site with no missing data at the group level) | The original set of SNPs and individuals, and the original algorithm for calculating $f_3$-statistics (470,389 variable site with no missing data at the group level) |
| --- | --- |
| 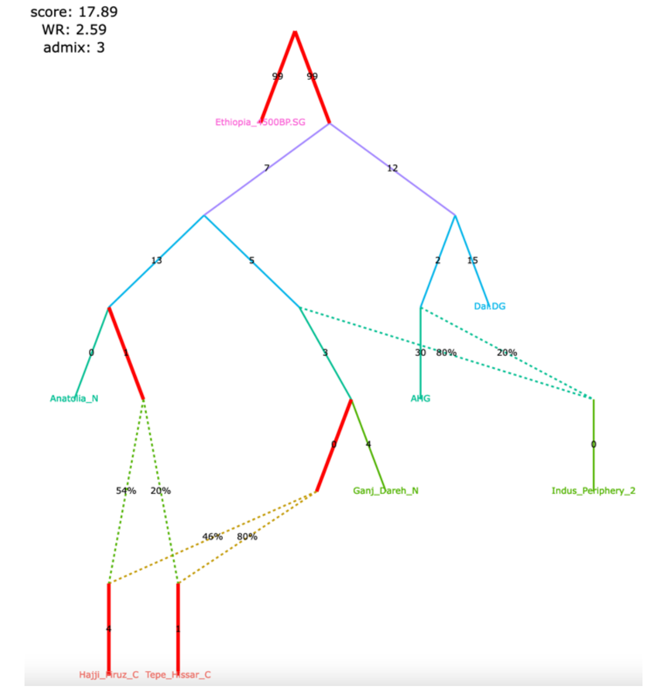 | 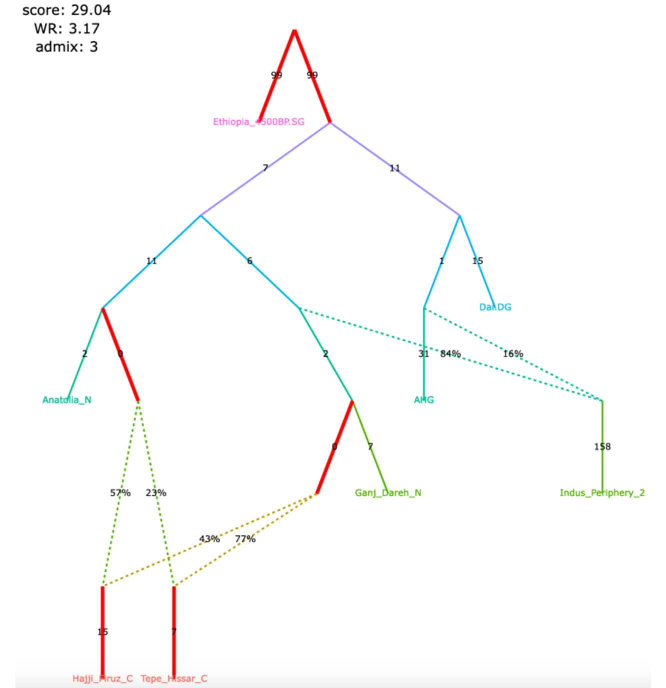 |

A claim by Shinde et al. 2019 relying on the admixture graph:

Primary ancestry in the Indus Periphery group forms the deepest branch in the Iranian Neolithic clade composed of the Indus Periphery, Ganj Dareh Neolithic, Hajji Firuz Chalcolithic and Tepe Hissar Chalcolithic groups.

Figure S6

**b**, alternative models fitting nominally better than the published one and confirming all of its important topological details

| Relatives (2nd and 3rd degree) and potentially contaminated individuals removed, the Indus Periphery group expanded to 4 ind., the corrected algorithm for calculating $f_3$-statistics (249,009 variable site with no missing data at the group level) | The original set of SNPs and individuals, and the original algorithm for calculating $f_3$-statistics (470,389 variable site with no missing data at the group level) |
|---|---|

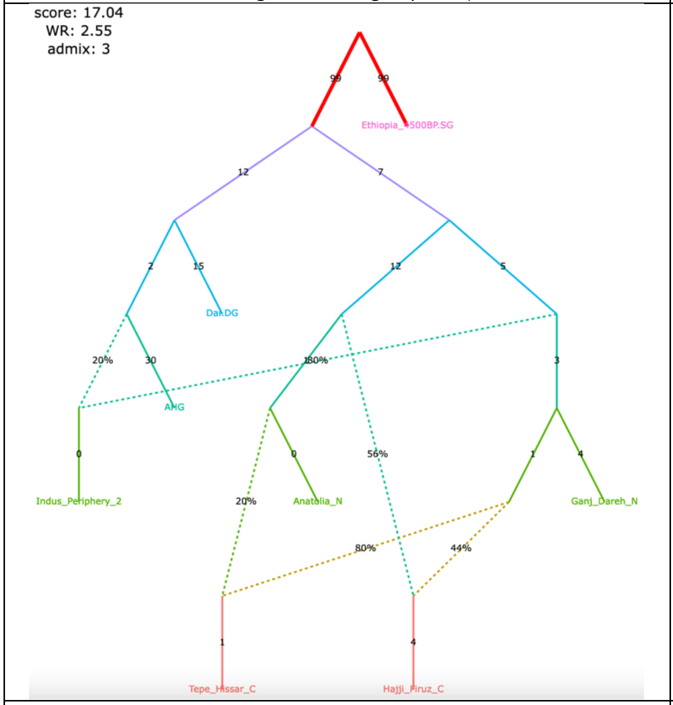

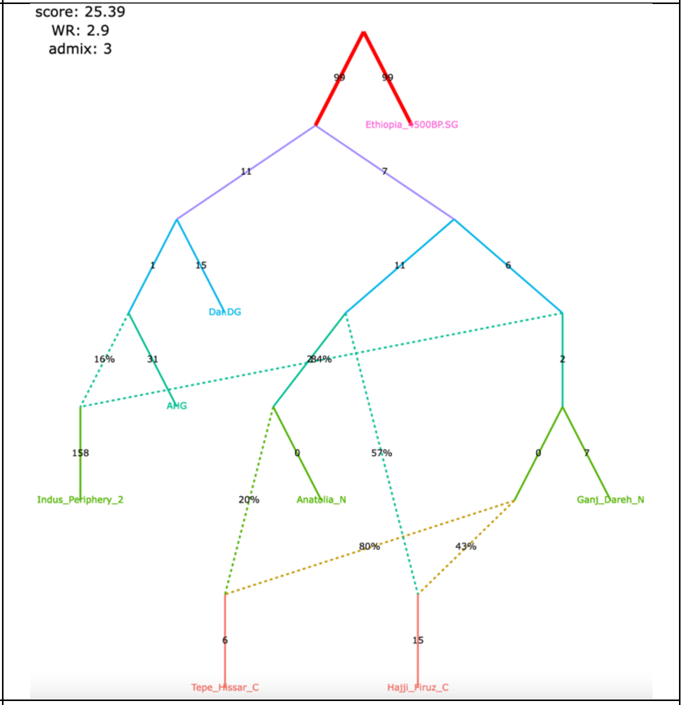

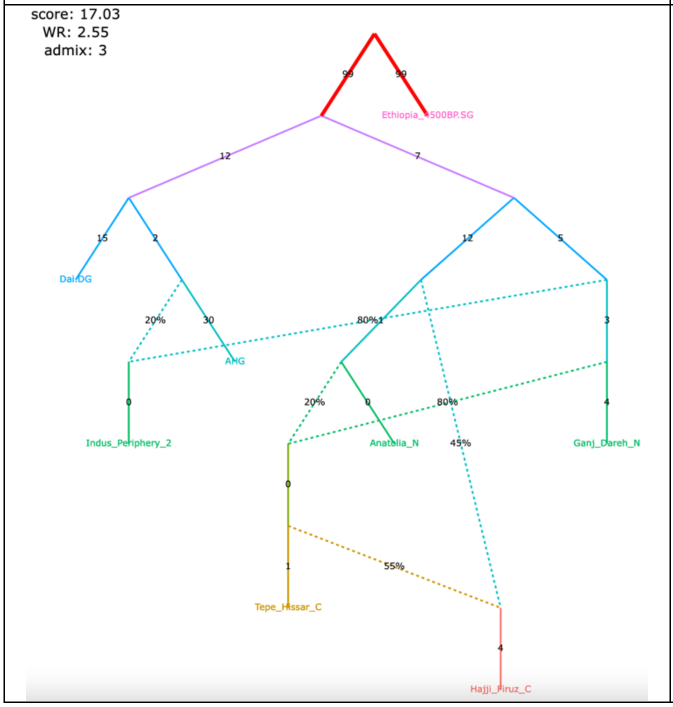

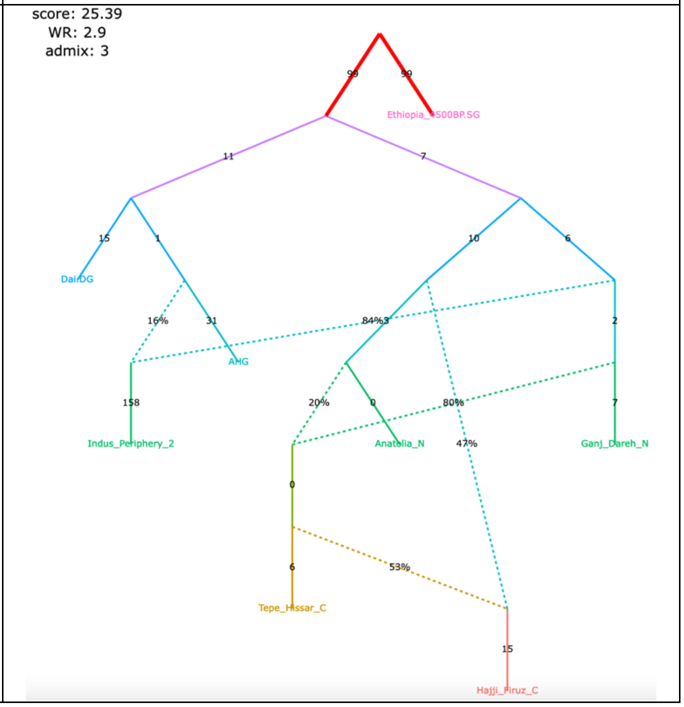

Figure S6

| Relatives (2nd and 3rd degree) and potentially contaminated individuals removed, the Indus Periphery group expanded to 4 ind., the corrected algorithm for calculating $f_3$-statistics (249,009 variable site with no missing data at the group level) | The original set of SNPs and individuals, and the original algorithm for calculating $f_3$-statistics (470,389 variable site with no missing data at the group level) |
|---|---|

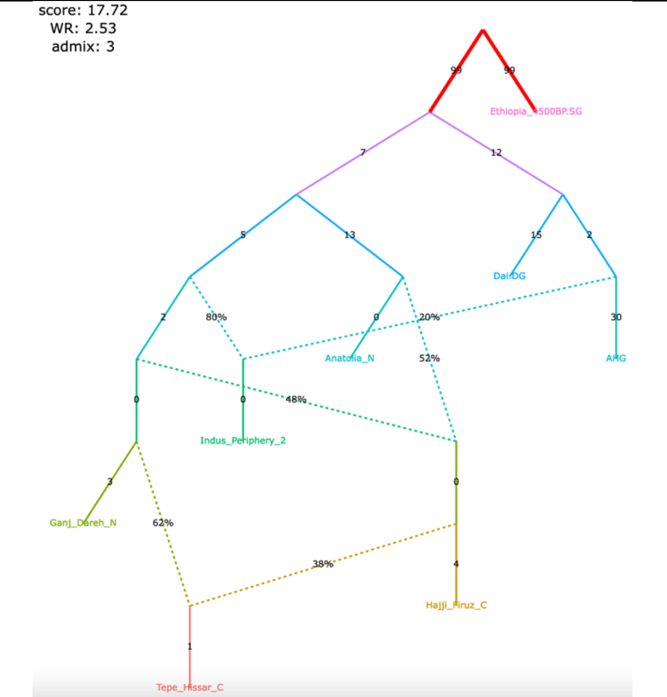

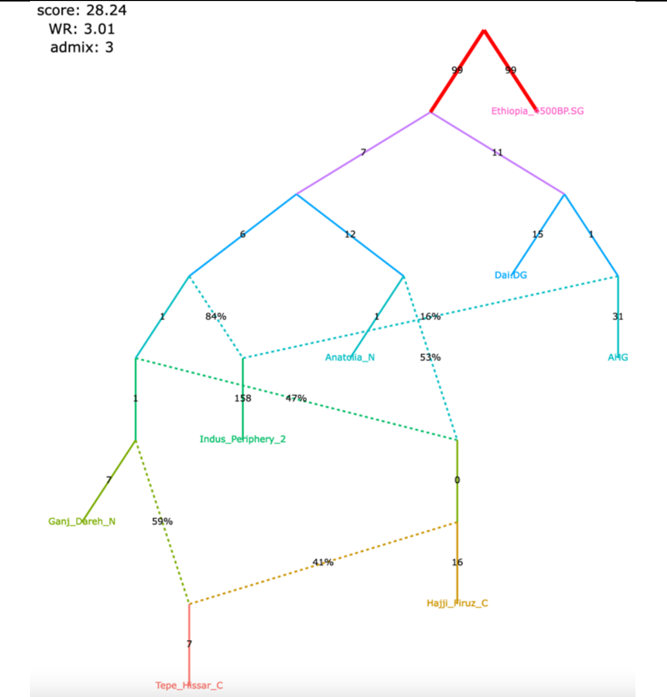

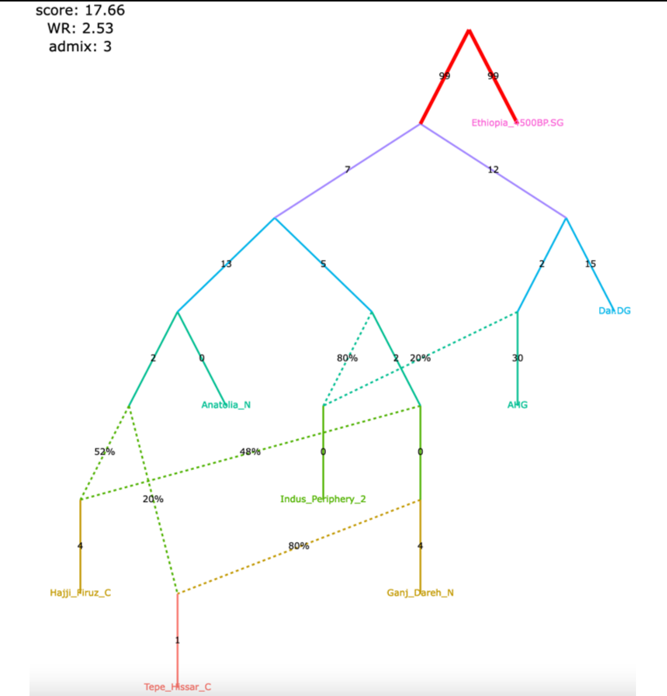

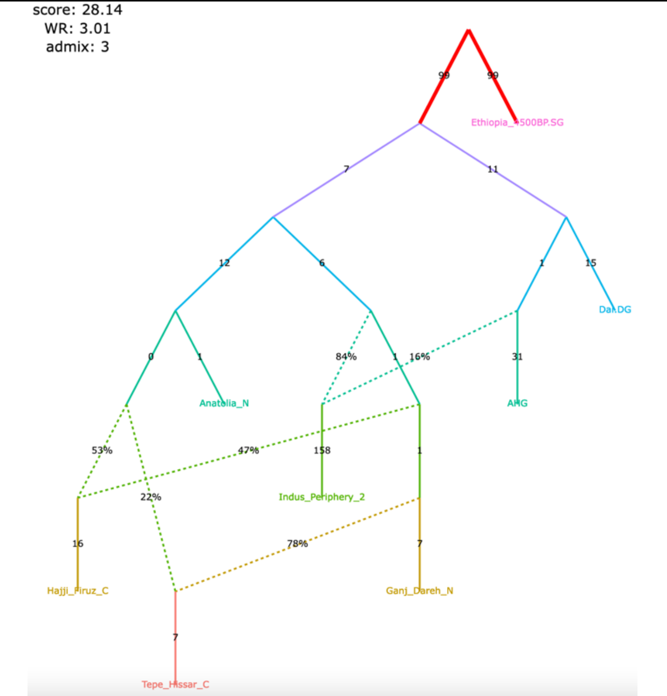

Figure S6

**c**, alternative models fitting not significantly worse than the published one and differing from it in important ways

| Relatives (2nd and 3rd degree) and potentially contaminated individuals removed, the Indus Periphery group expanded to 4 ind., the corrected algorithm for calculating $f_3$-statistics (249,009 variable site with no missing data at the group level) | The original set of SNPs and individuals, and the original algorithm for calculating $f_3$-statistics (470,389 variable site with no missing data at the group level) |
|---|---|
| 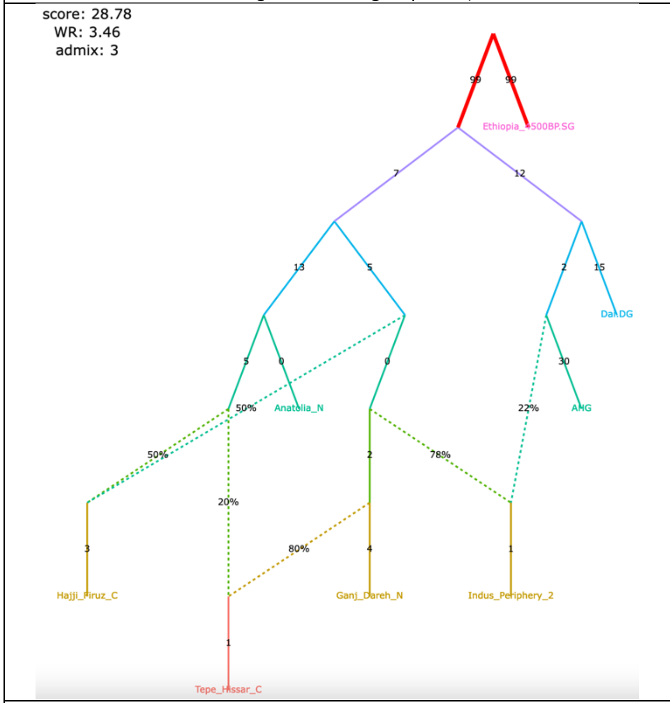 | 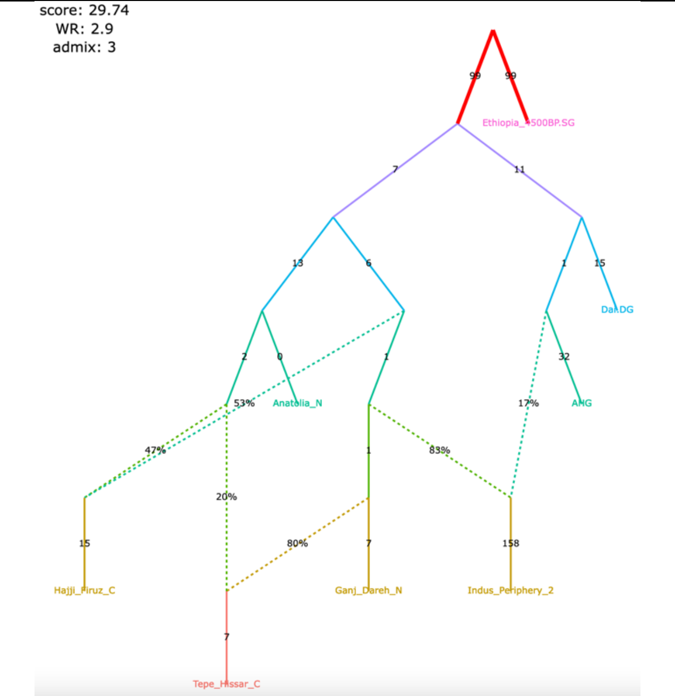 |
| 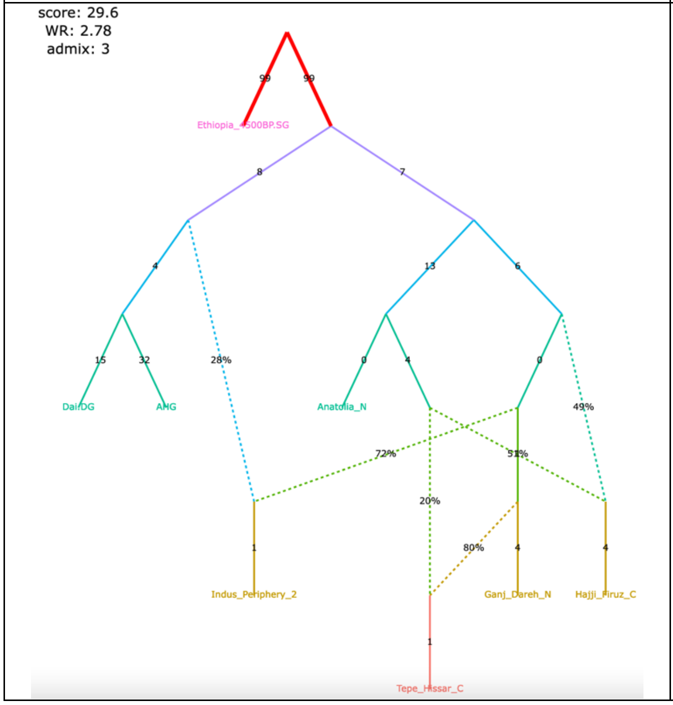 | 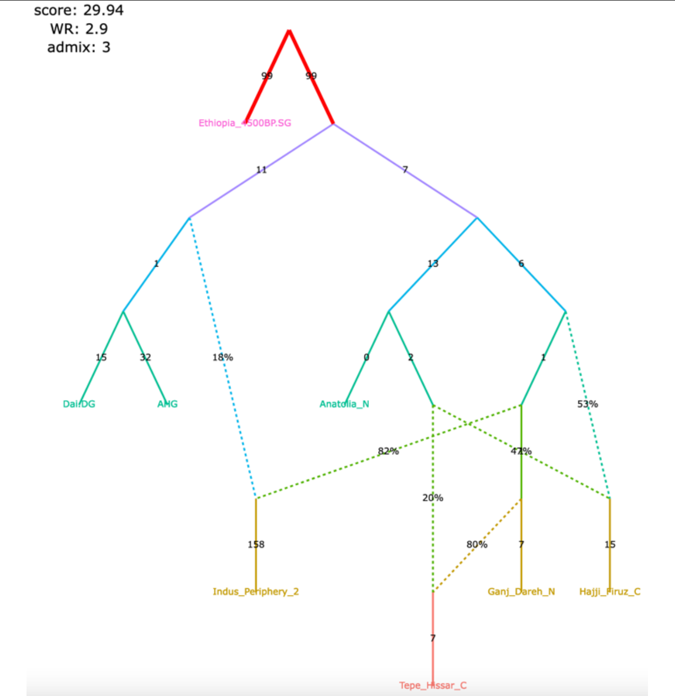 |

Figure S6

| Relatives (2nd and 3rd degree) and potentially contaminated individuals removed, the Indus Periphery group expanded to 4 ind., the corrected algorithm for calculating $f_3$-statistics (249,009 variable site with no missing data at the group level) | The original set of SNPs and individuals, and the original algorithm for calculating $f_3$-statistics (470,389 variable site with no missing data at the group level) |
|---|---|

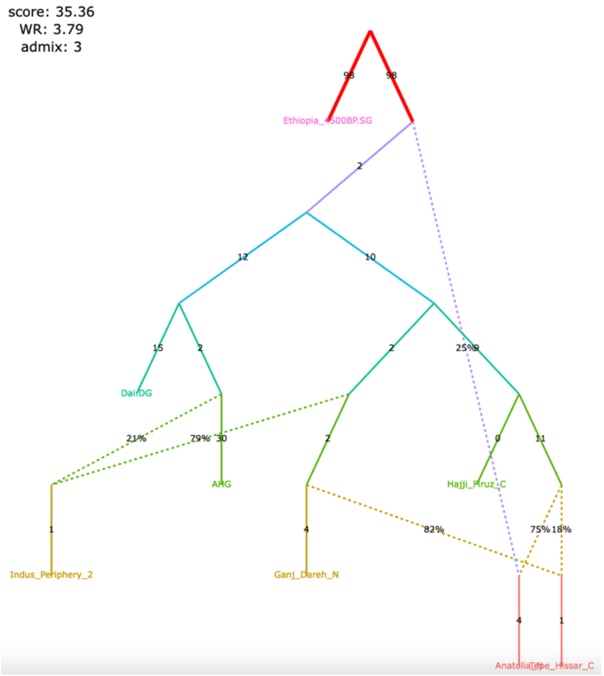

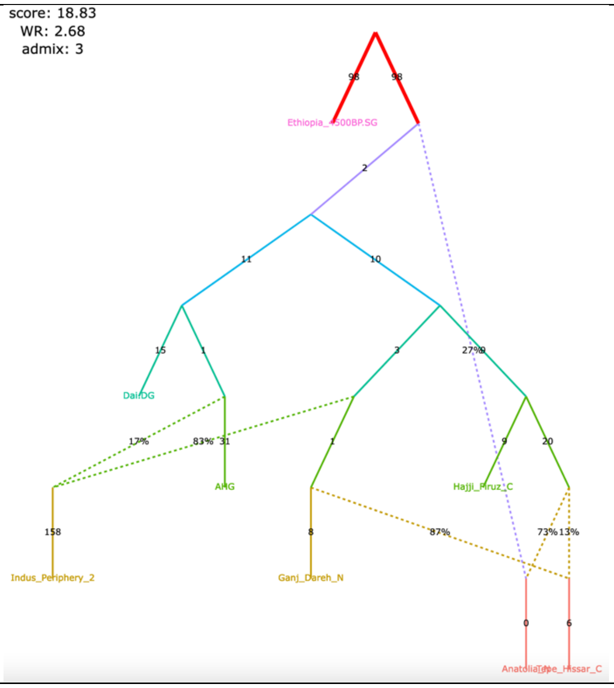

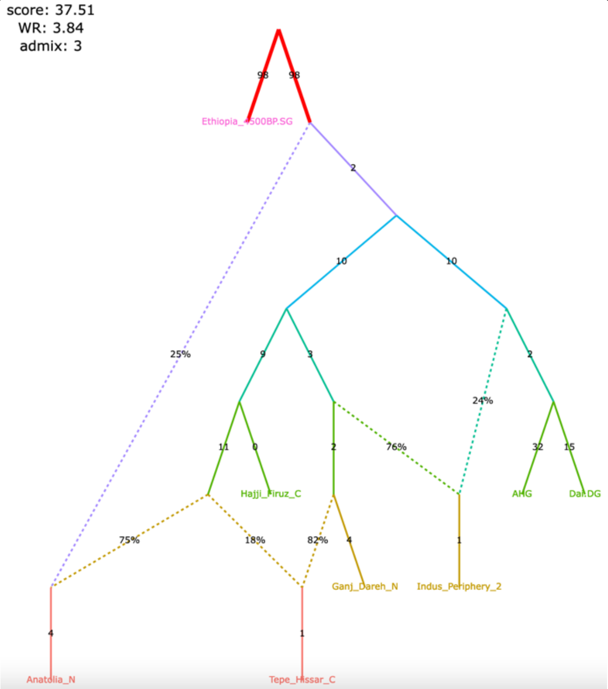

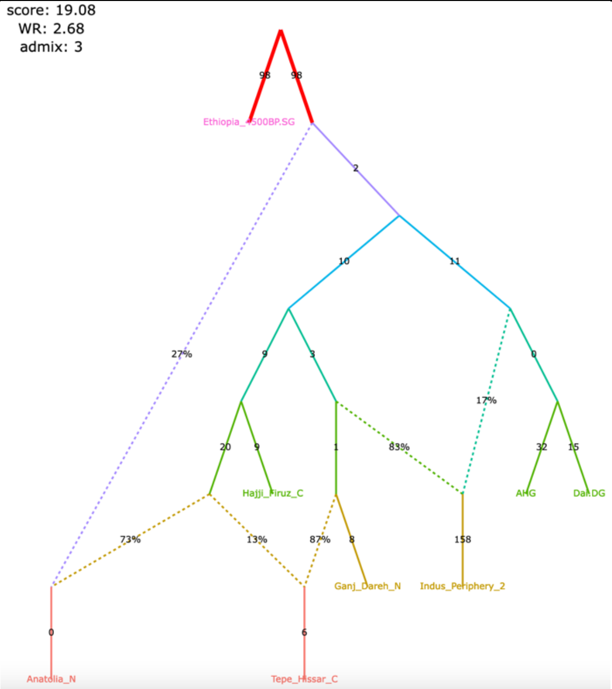

Figure S7

**Figure S7.** Alternative graphs allowing for an additional admixture event found with *findGraphs* for the dataset from Shinde *et al*. (2019): 8 populations, 4 admixture events, the modified group composition and the updated algorithm for calculating *f*-statistics. The graphs were also re-fitted on the original set of SNPs/individuals and using the original algorithm for calculating *f*-statistics.

**a**, highest-ranking model with 4 admixture events; confirms all important features of the published model with 3 admixture events

| Relatives (2nd and 3rd degree) and potentially contaminated individuals removed, the Indus Periphery group expanded to 4 ind., the corrected algorithm for calculating $f_3$-statistics (249,009 variable site with no missing data at the group level) | The original set of SNPs and individuals, and the original algorithm for calculating $f_3$-statistics (470,389 variable site with no missing data at the group level) |
|---|---|
| 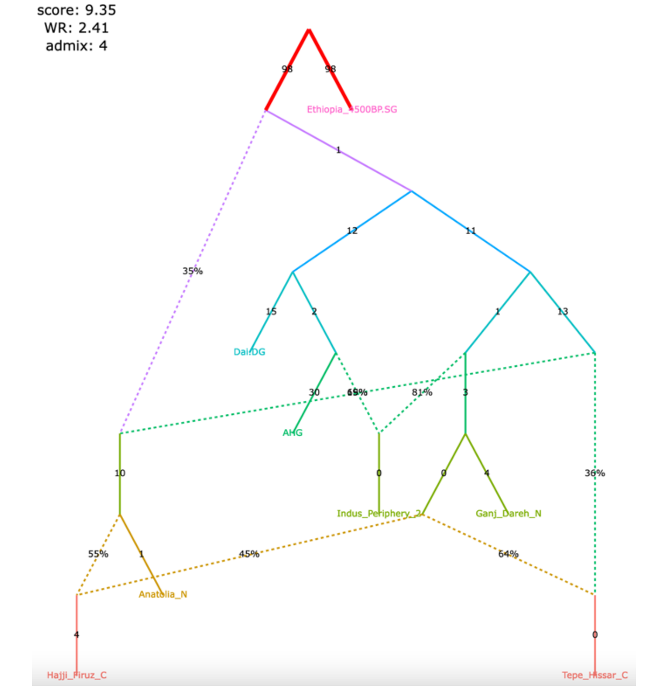  | 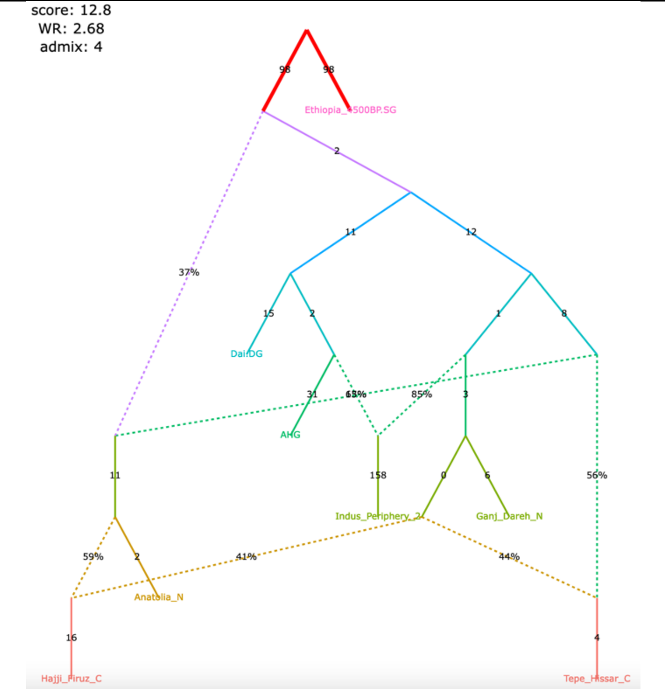  |

A claim by Shinde et al. 2019 relying on the admixture graph:

Primary ancestry in the Indus Periphery group forms the deepest branch in the Iranian Neolithic clade composed of the Indus Periphery, Ganj Dareh Neolithic, Hajji Firuz Chalcolithic and Tepe Hissar Chalcolithic groups.

Figure S7

**b**, alternative models fitting not significantly worse than the highest-ranking one and contradicting the historical interpretation of the admixture graph results by Shinde et al. 2019

| Relatives (2nd and 3rd degree) and potentially contaminated individuals removed, the Indus Periphery group expanded to 4 ind., the corrected algorithm for calculating $f_3$-statistics (249,009 variable site with no missing data at the group level) | The original set of SNPs and individuals, and the original algorithm for calculating $f_3$-statistics (470,389 variable site with no missing data at the group level) |
|---|---|
| 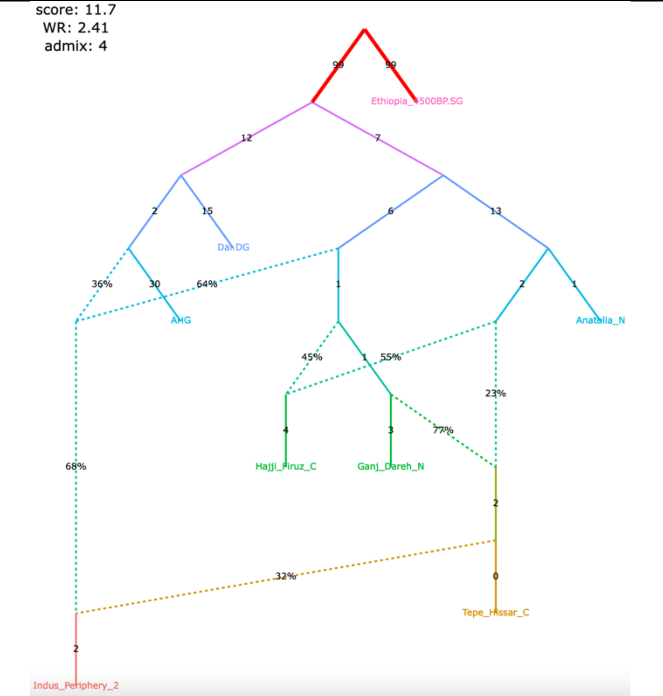 | 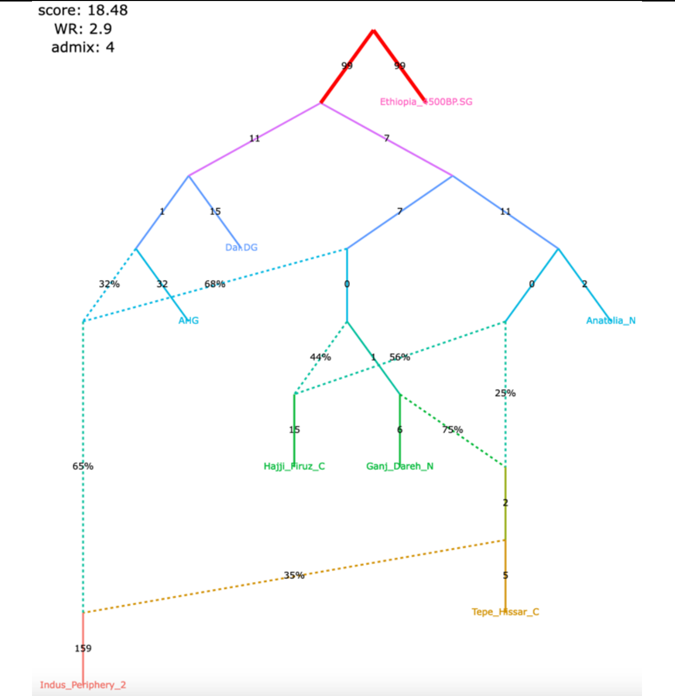 |
| 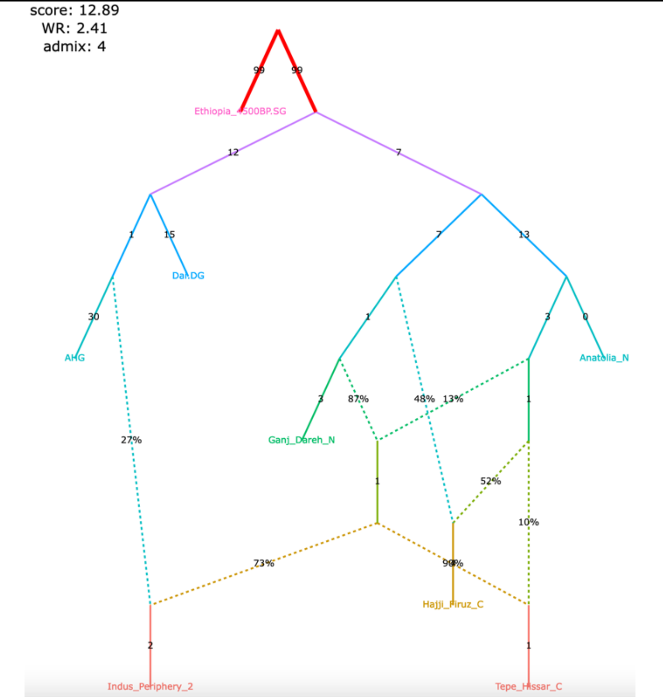 | 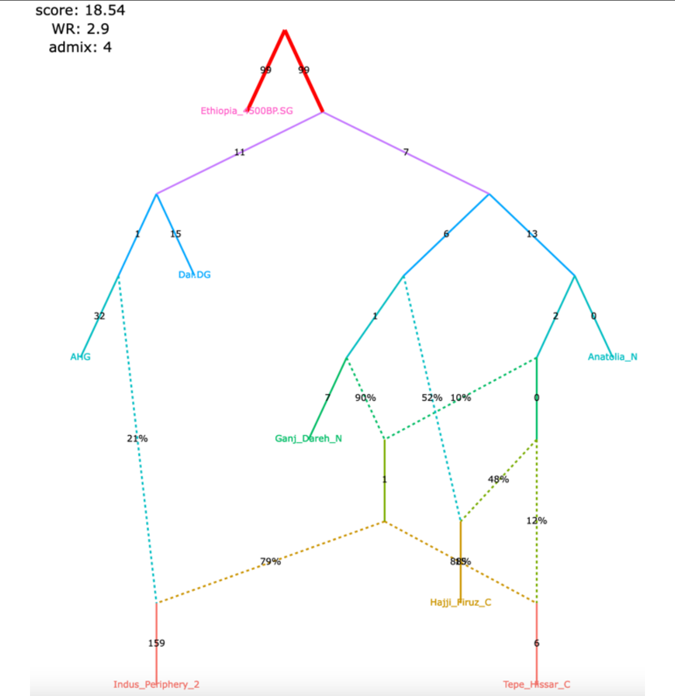 |

### Figure S7

| Relatives (2nd and 3rd degree) and potentially contaminated individuals removed, the Indus Periphery group expanded to 4 ind., the corrected algorithm for calculating $f_3$-statistics (249,009 variable site with no missing data at the group level) | The original set of SNPs and individuals, and the original algorithm for calculating $f_3$-statistics (470,389 variable site with no missing data at the group level) |
| --- | --- |
| 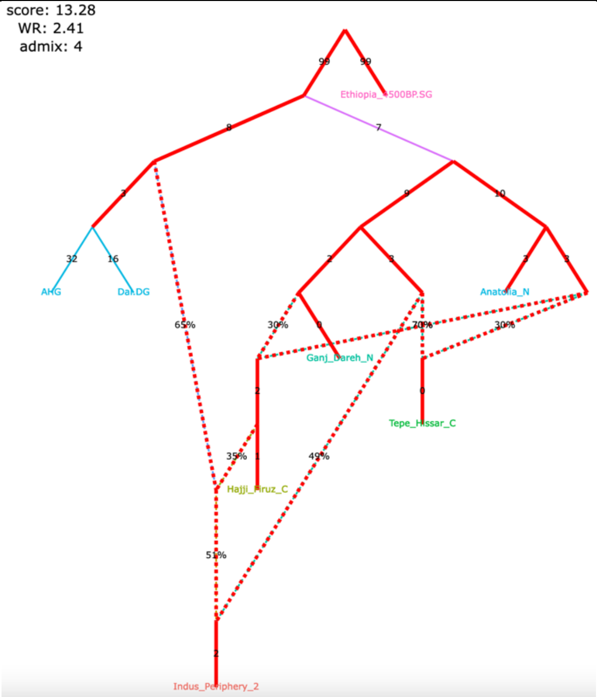 | 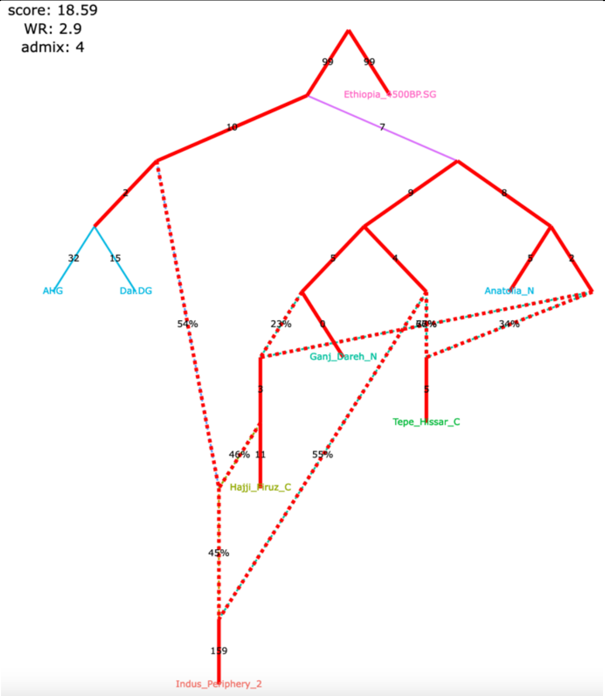 |
| 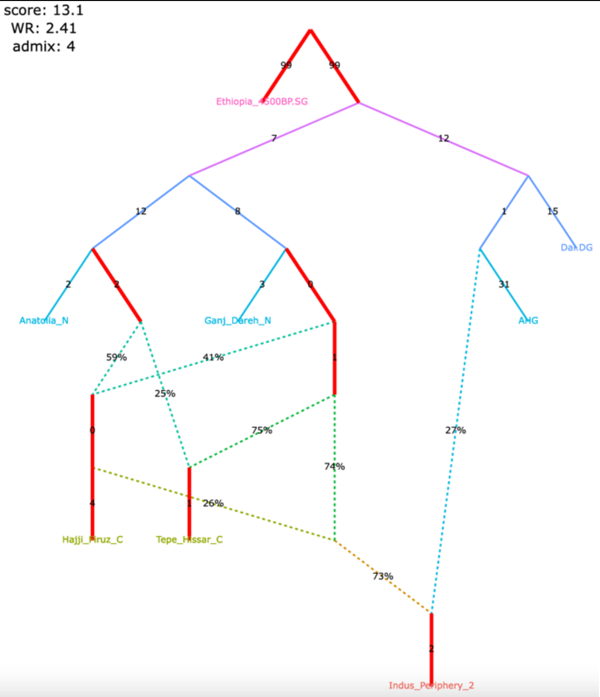 | 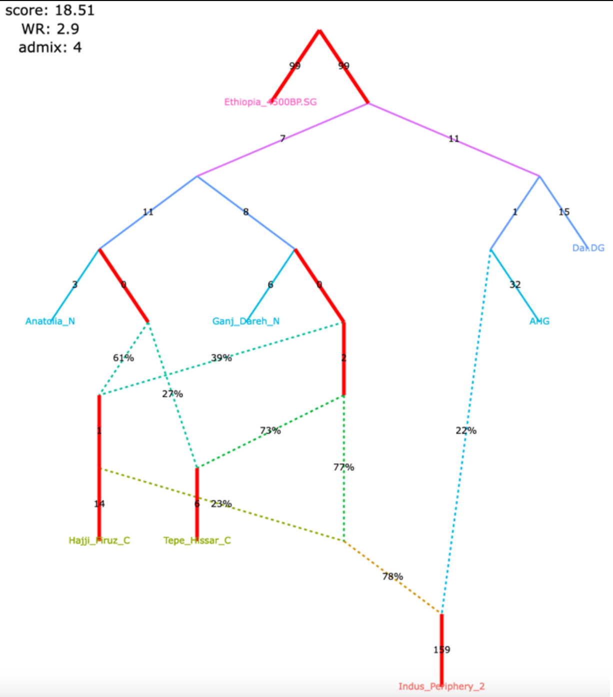 |

Figure S8

**Figure S8.** Published admixture graphs from Librado *et al*. (2019) and alternative graphs found with *findGraphs* (10 populations, 3 to 5 admixture events) for the modified group composition and using the updated algorithm for calculating *f*-statistics. The graphs were also re-fitted on the original set of SNPs/individuals and using the original algorithm for calculating *f*-statistics. Selected alternative graphs found with *findGraphs* when more admixture events were allowed (from 6 to 9) are also shown.

**a**, published model, 3 admixture events

| | |
|---|---|
| **claim 1** yes<br>**claim 2** yes<br>**claim 3** yes<br>**claim 4** yes<br>**claim 5** yes | **claim 1** yes<br>**claim 2** yes<br>**claim 3** yes<br>**claim 4** yes<br>**claim 5** yes |
| The modified set of individuals and the corrected algorithm for calculating $f_3$-statistics (1,767,419 variable site with no missing data at the group level) | The modified set of individuals and the original algorithm for calculating $f_3$-statistics (6,343,116 variable site with no missing data at the group level) |
| 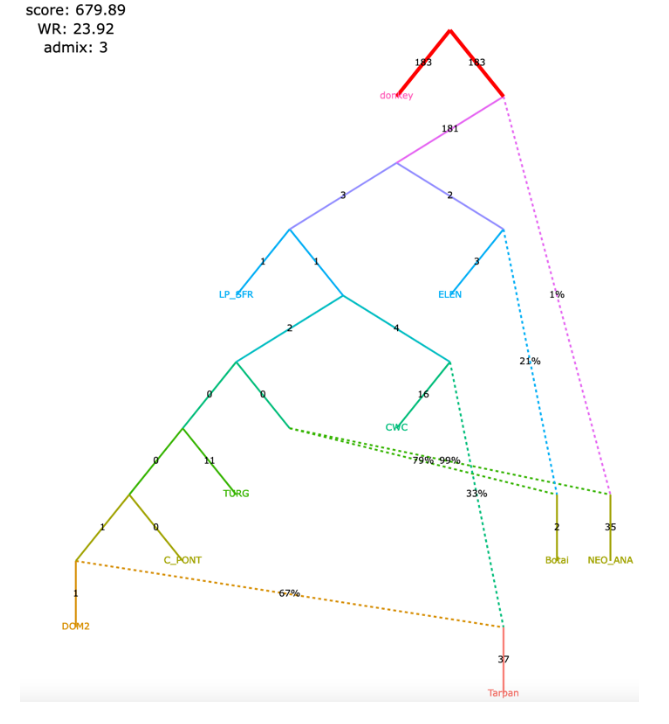 | 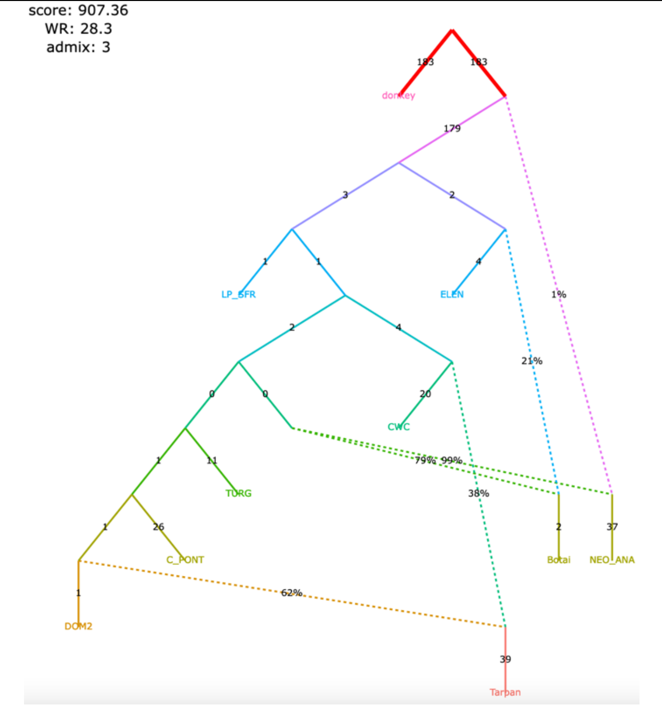 |

Claims by Librado et al. 2021 relying on the admixture graph:

1) NEO-ANA-related admixture is absent in DOM2;
2) DOM2 and C-PONT are sister groups;
3) there is no gene flow connecting the CWC and the cluster associated with Yamnaya horses and horses of the later Sintashta culture whose ancestry is maximized in the Western Steppe (DOM2, C-PONT, TURG);
4) there was a gene flow from a deep-branching ghost group to NEO-ANA;
5) Tarpan is a mixture of a CWC-related and a DOM2-related lineage.

Figure S8

**b**, an alternative model with 3 admixture events fitting significantly better than the published one

<table>
<tr>
<td>

| | |
|---|---|
| **claim 1** | yes |
| **claim 2** | yes |
| **claim 3** | yes |
| **claim 4** | yes |
| **claim 5** | no |

</td>
<td>

| | |
|---|---|
| **claim 1** | yes |
| **claim 2** | yes |
| **claim 3** | yes |
| **claim 4** | yes |
| **claim 5** | no |

</td>
</tr>
<tr>
<td>The modified set of individuals and the corrected algorithm for calculating f3-statistics (1,767,419 variable site with no missing data at the group level)</td>
<td>The modified set of individuals and the original algorithm for calculating f3-statistics (6,343,116 variable site with no missing data at the group level)</td>
</tr>
<tr>
<td>

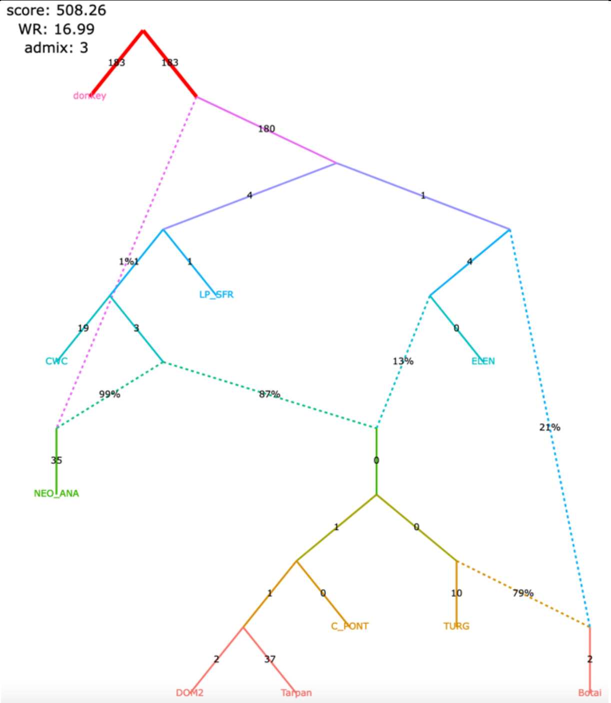

</td>
<td>

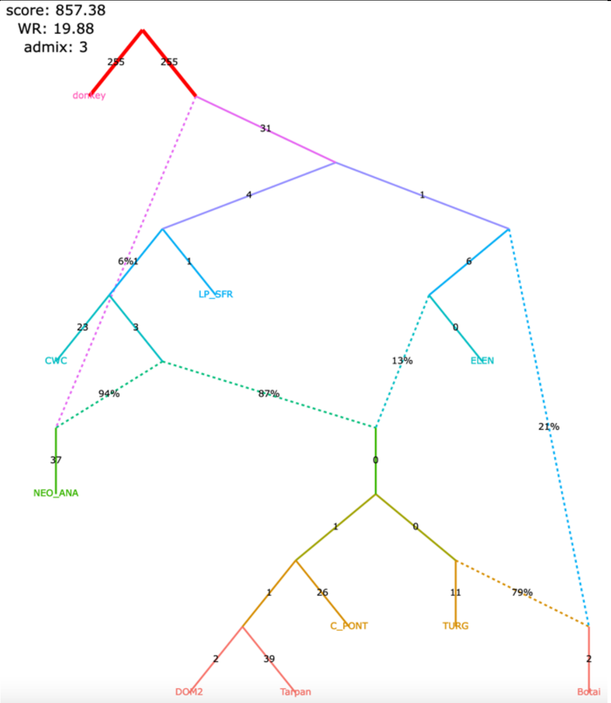

</td>
</tr>
</table>

Figure S8

#### **c**, published model, 4 admixture events

| | |
|---|---|
| **claim 1** yes<br>**claim 2** yes<br>**claim 3** yes<br>**claim 4** yes<br>**claim 5** yes | **claim 1** yes<br>**claim 2** yes<br>**claim 3** yes<br>**claim 4** yes<br>**claim 5** yes |
| The modified set of individuals and the corrected algorithm for calculating $f_3$-statistics (1,767,419 variable site with no missing data at the group level) | The modified set of individuals and the original algorithm for calculating $f_3$-statistics (6,343,116 variable site with no missing data at the group level) |
| 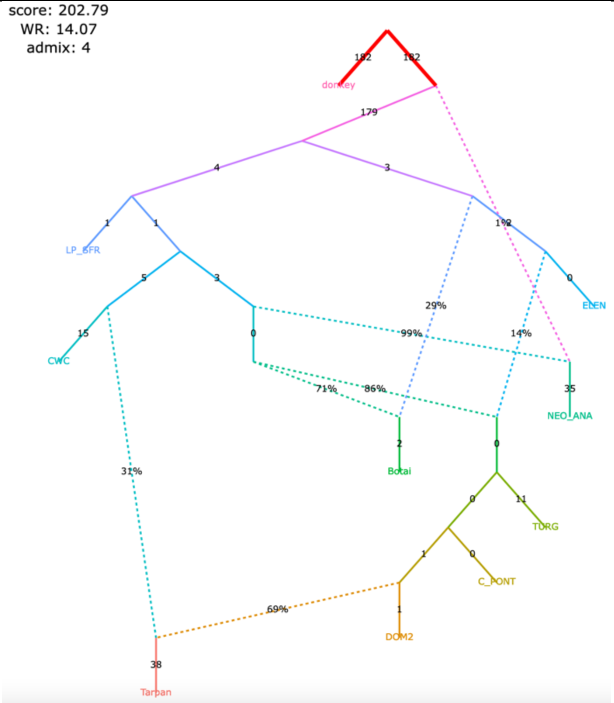 | 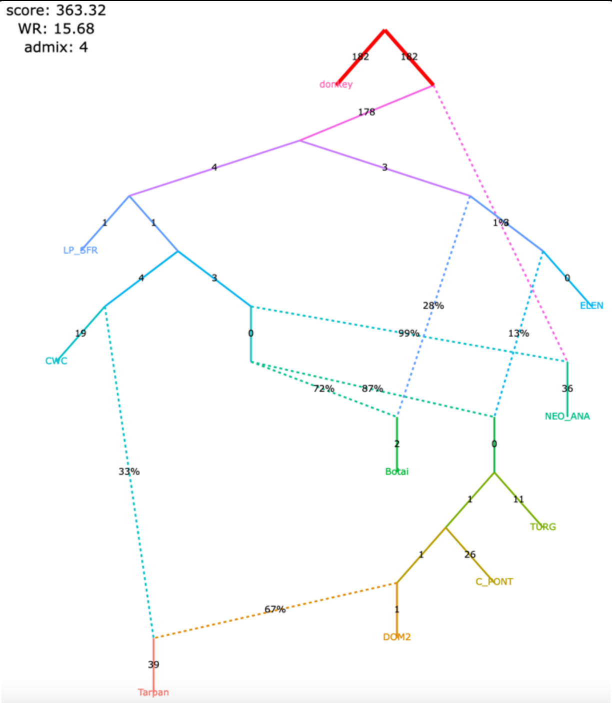 |

Figure S8

**d**, an alternative model with 4 admixture events fitting not significantly worse than the published one

| | |
|---|---|
| claim 1 yes<br>claim 2 yes<br>claim 3 yes<br>claim 4 yes<br>claim 5 yes | claim 1 yes<br>claim 2 yes<br>claim 3 yes<br>claim 4 yes<br>claim 5 yes |
| The modified set of individuals and the corrected algorithm for calculating *f*$_3$-statistics (1,767,419 variable site with no missing data at the group level) | The modified set of individuals and the original algorithm for calculating *f*$_3$-statistics (6,343,116 variable site with no missing data at the group level) |

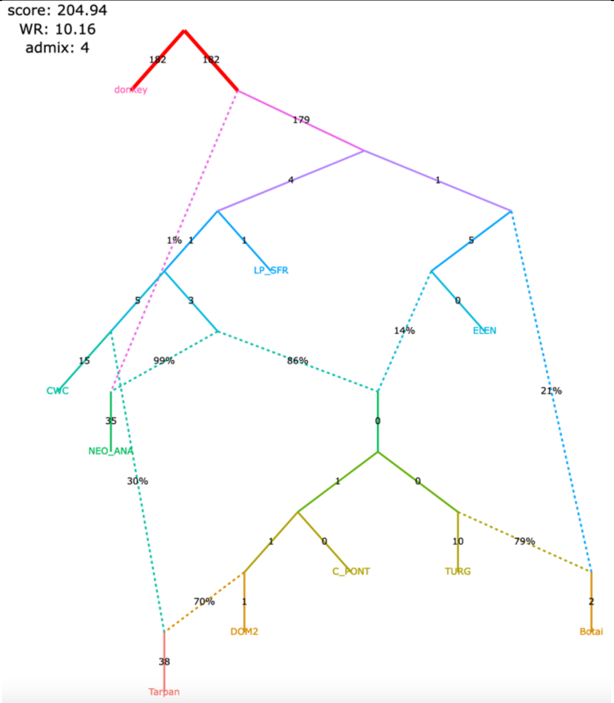

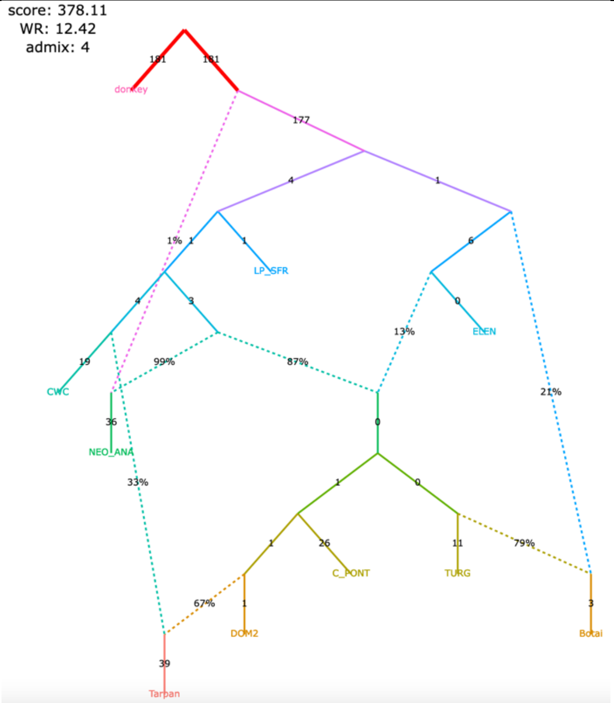

Figure S8

#### **e**, published model, 5 admixture events

| | |
|---|---|
| **claim 1** yes<br>**claim 2** yes<br>**claim 3** yes<br>**claim 4** yes<br>**claim 5** yes | **claim 1** yes<br>**claim 2** yes<br>**claim 3** yes<br>**claim 4** yes<br>**claim 5** yes |
| The modified set of individuals and the corrected algorithm for calculating *f*$_3$-statistics (1,767,419 variable site with no missing data at the group level) | The modified set of individuals and the original algorithm for calculating *f*$_3$-statistics (6,343,116 variable site with no missing data at the group level) |

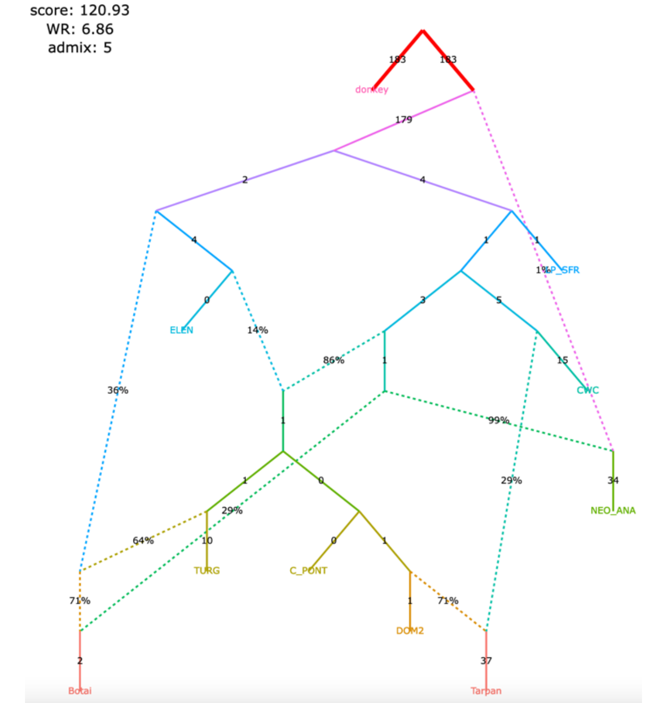

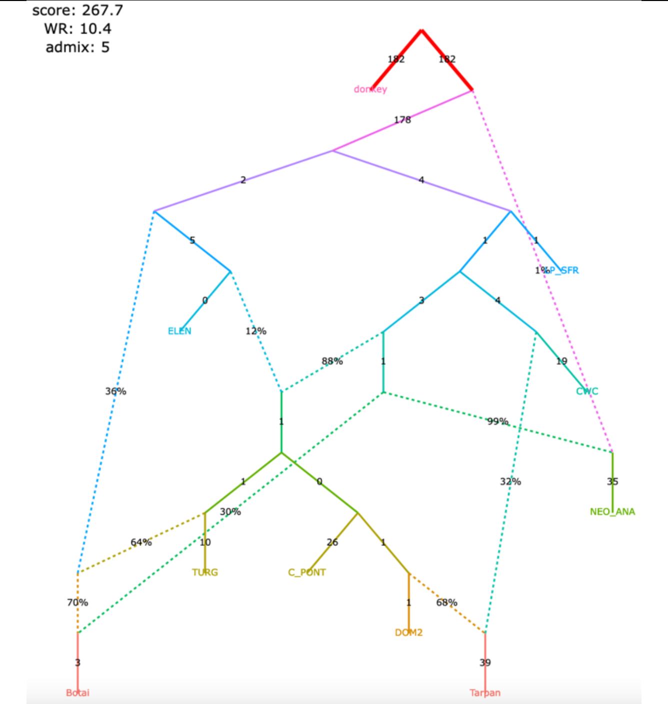

Figure S8

**f**, an alternative model with 5 admixture events fitting not significantly worse than the published one

| | |
|---|---|
| **claim 1** yes<br>**claim 2** yes<br>**claim 3** no<br>**claim 4** yes<br>**claim 5** yes | **claim 1** yes<br>**claim 2** yes<br>**claim 3** no<br>**claim 4** yes<br>**claim 5** yes |
| The modified set of individuals and the corrected algorithm for calculating $f_3$-statistics (1,767,419 variable site with no missing data at the group level) | The modified set of individuals and the original algorithm for calculating $f_3$-statistics (6,343,116 variable site with no missing data at the group level) |

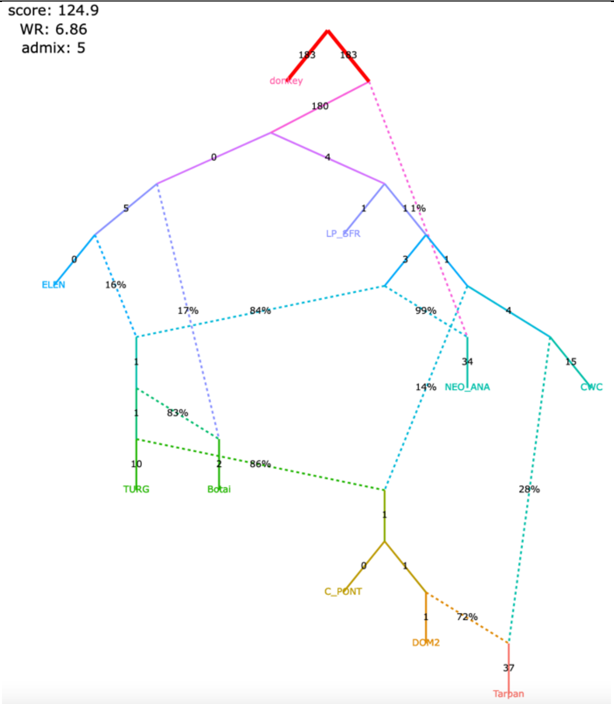

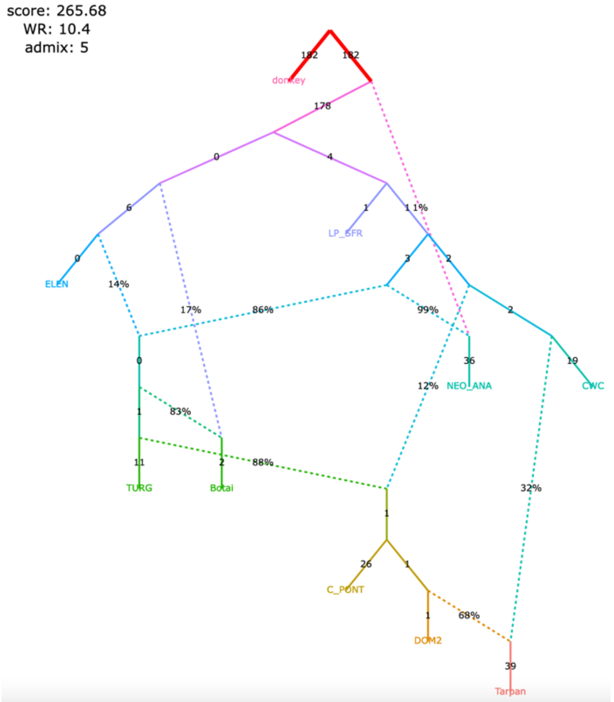

Figure S8

#### **g**, selected models with 6 admixture events

| | |
|---|---|
| **claim 1** yes<br>**claim 2** yes<br>**claim 3** no<br>**claim 4** yes<br>**claim 5** yes | **claim 1** yes<br>**claim 2** yes<br>**claim 3** no<br>**claim 4** yes<br>**claim 5** yes |
| The modified set of individuals and the corrected algorithm for calculating $f_3$-statistics (1,767,419 variable site with no missing data at the group level) | The modified set of individuals and the original algorithm for calculating $f_3$-statistics (6,343,116 variable site with no missing data at the group level) |
| 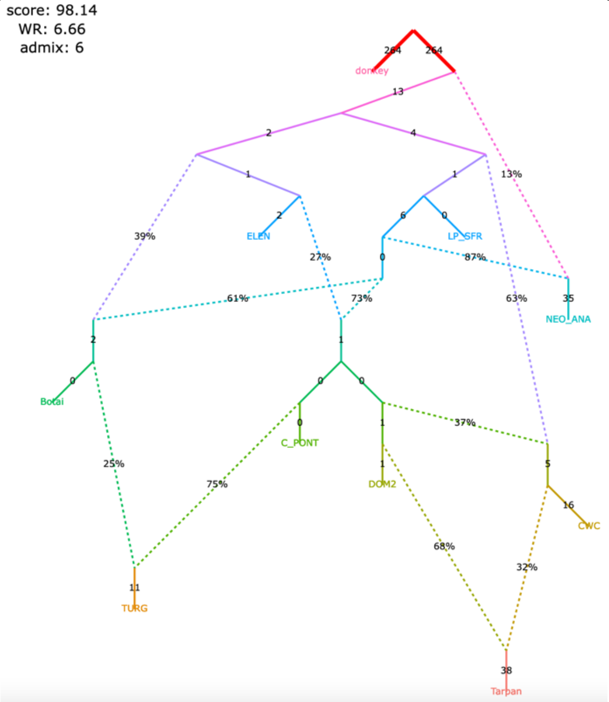 | 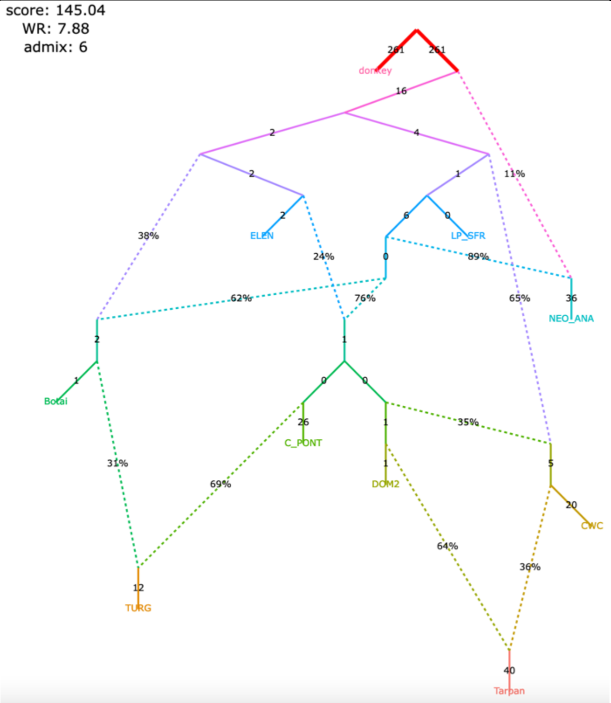 |
| **claim 1** no<br>**claim 2** yes<br>**claim 3** no<br>**claim 4** no<br>**claim 5** yes | **claim 1** no<br>**claim 2** yes<br>**claim 3** no<br>**claim 4** no<br>**claim 5** yes |
| 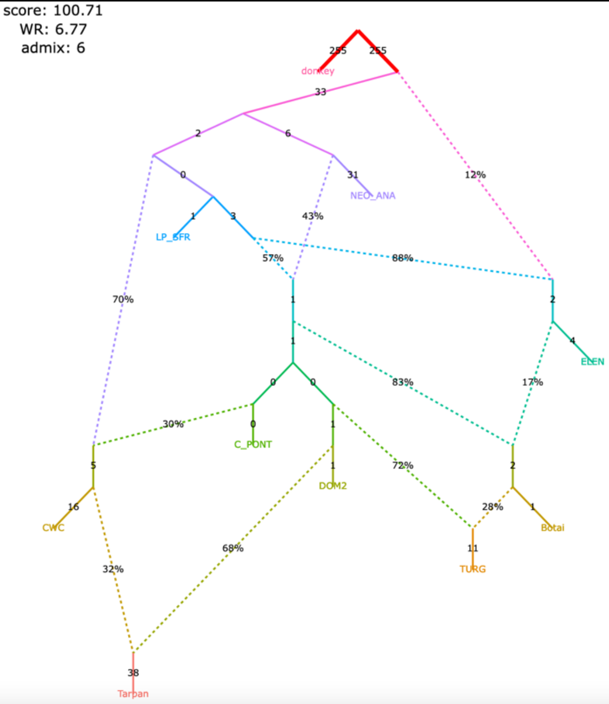 | 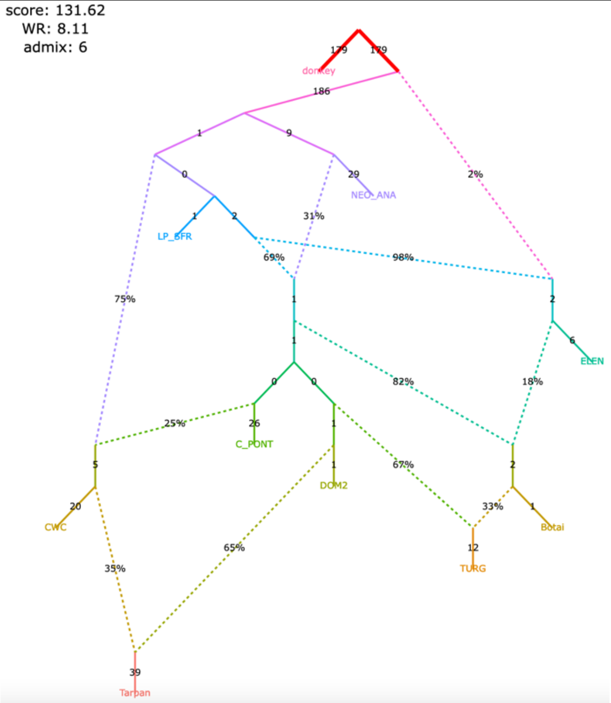 |

Figure S8

#### **h**, selected models with 7 admixture events (all plausible models with WR < 5 SE)

| | |
|---|---|
| **claim 1** yes<br>**claim 2** yes<br>**claim 3** no<br>**claim 4** yes<br>**claim 5** no | **claim 1** yes<br>**claim 2** yes<br>**claim 3** no<br>**claim 4** yes<br>**claim 5** no |
| The modified set of individuals and the corrected algorithm for calculating $f_3$-statistics (1,767,419 variable site with no missing data at the group level) | The modified set of individuals and the original algorithm for calculating $f_3$-statistics (6,343,116 variable site with no missing data at the group level) |

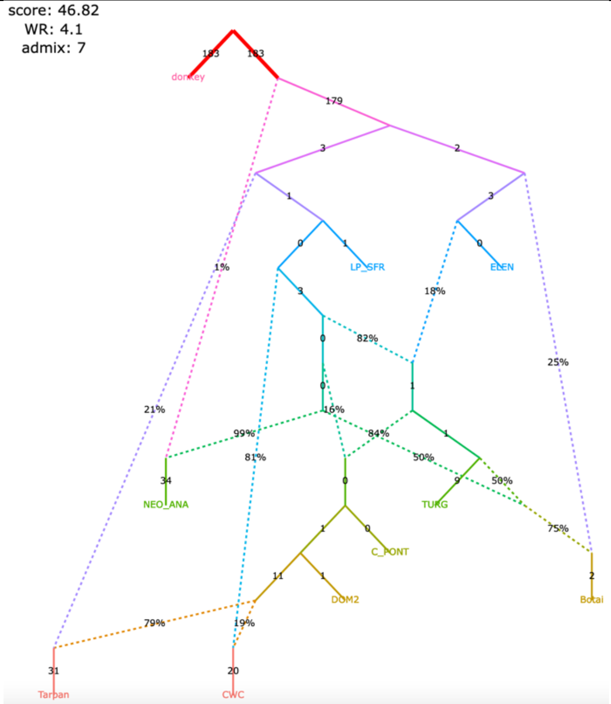

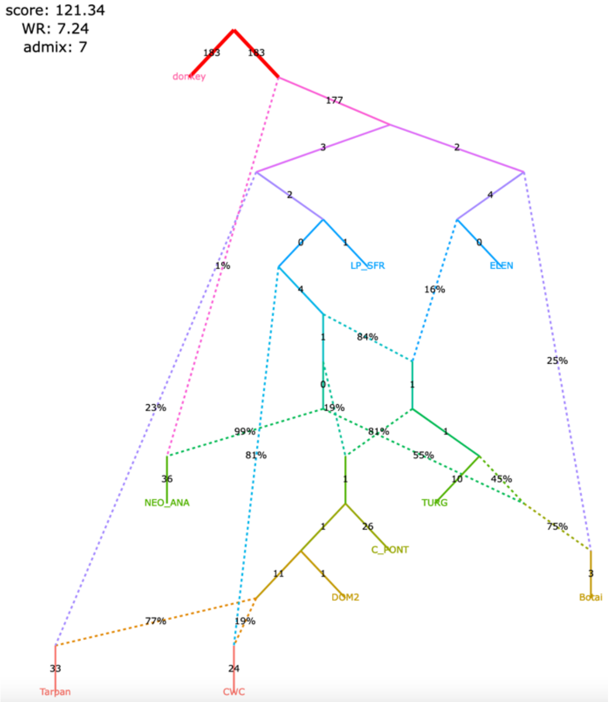

| | |
|---|---|
| **claim 1** yes<br>**claim 2** yes<br>**claim 3** no<br>**claim 4** yes<br>**claim 5** yes | **claim 1** yes<br>**claim 2** yes<br>**claim 3** no<br>**claim 4** yes<br>**claim 5** yes |

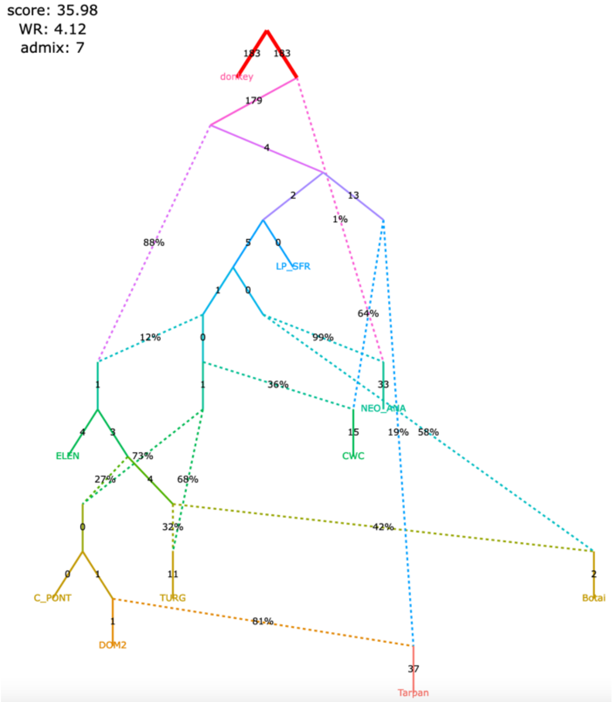

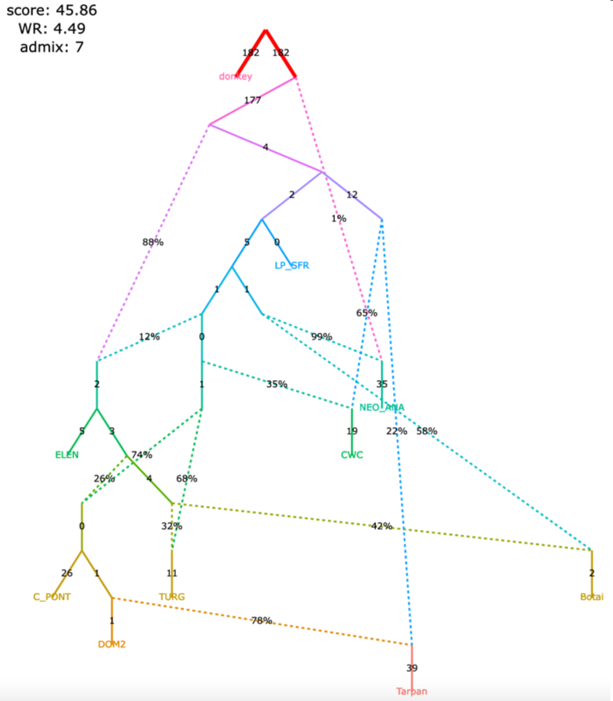

Figure S8

#### **i**, selected models with 7 admixture events (all plausible models with WR < 5 SE)

| | |
|---|---|
| **claim 1** yes<br>**claim 2** yes<br>**claim 3** no<br>**claim 4** yes<br>**claim 5** yes | **claim 1** yes<br>**claim 2** yes<br>**claim 3** no<br>**claim 4** yes<br>**claim 5** yes |
| The modified set of individuals and the corrected algorithm for calculating $f_3$-statistics (1,767,419 variable site with no missing data at the group level) | The modified set of individuals and the original algorithm for calculating $f_3$-statistics (6,343,116 variable site with no missing data at the group level) |

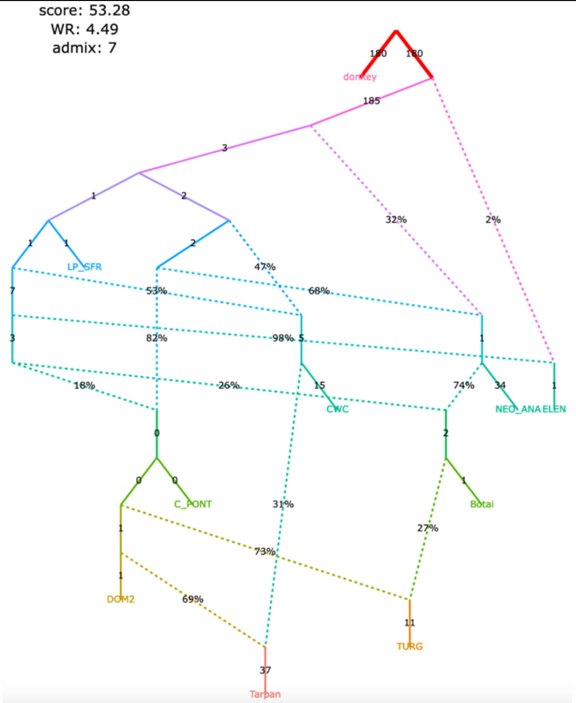

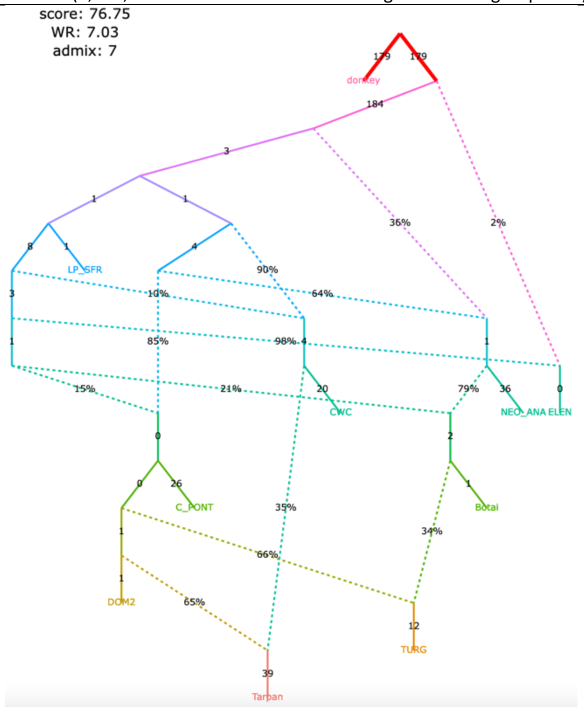

| | |
|---|---|
| **claim 1** no<br>**claim 2** yes<br>**claim 3** no<br>**claim 4** yes<br>**claim 5** no | **claim 1** no<br>**claim 2** yes<br>**claim 3** no<br>**claim 4** yes<br>**claim 5** no |

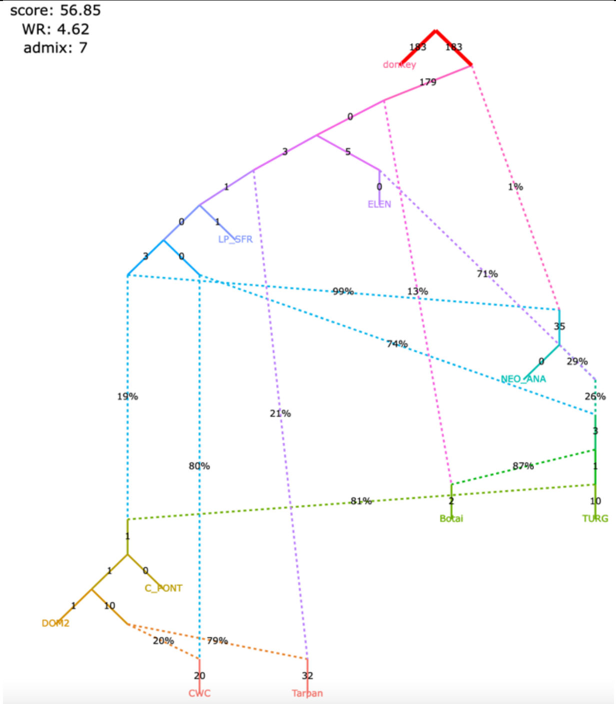

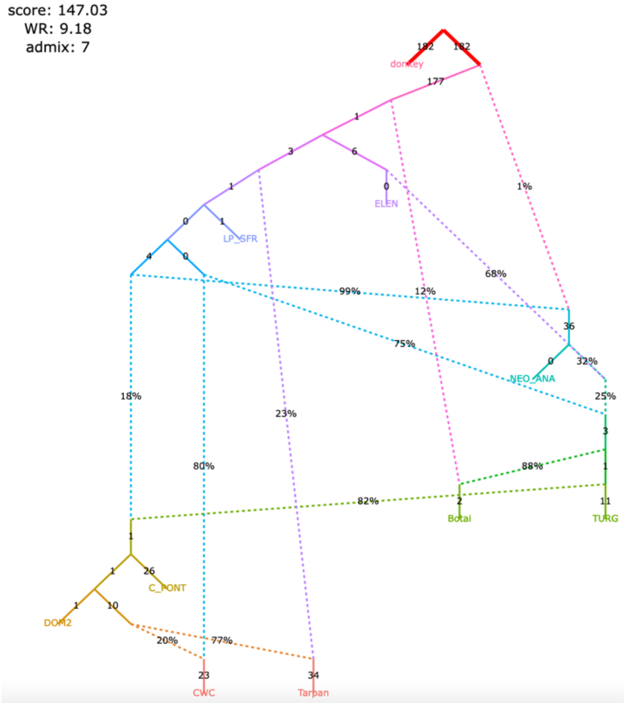

Figure S8

#### **j**, selected models with 8 admixture events (all plausible models with WR < 4 SE)

**claim 1** no
**claim 2** yes
**claim 3** no
**claim 4** yes
**claim 5** no

The modified set of individuals and the corrected algorithm for calculating *f*$_3$-statistics (1,767,419 variable site with no missing data at the group level)

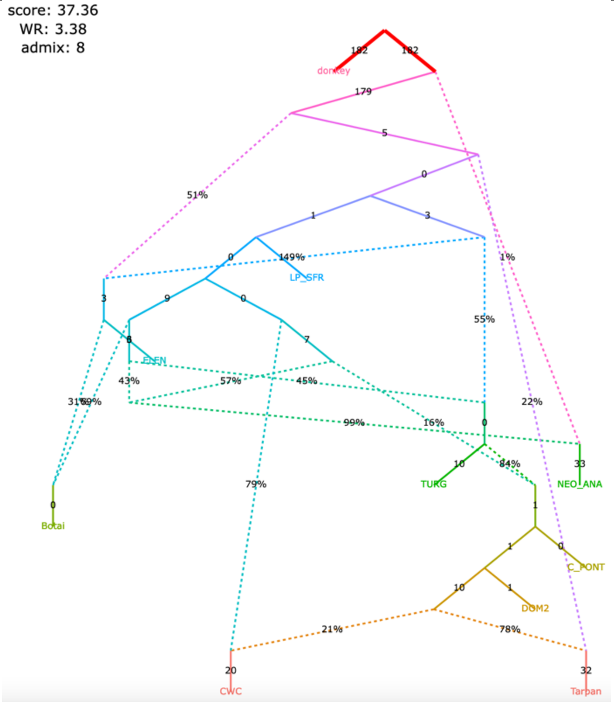

**claim 1** no
**claim 2** yes
**claim 3** no
**claim 4** yes
**claim 5** no

The modified set of individuals and the original algorithm for calculating *f*$_3$-statistics (6,343,116 variable site with no missing data at the group level)

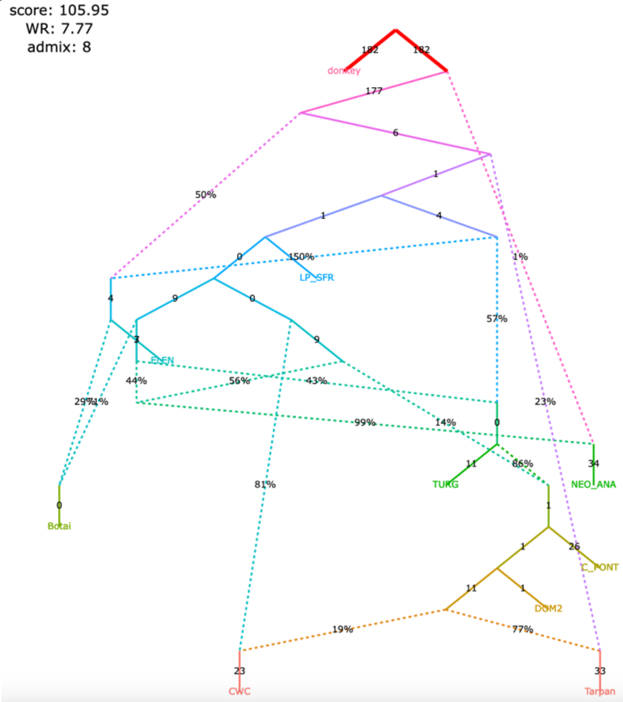

**claim 1** no
**claim 2** yes
**claim 3** yes
**claim 4** yes
**claim 5** no

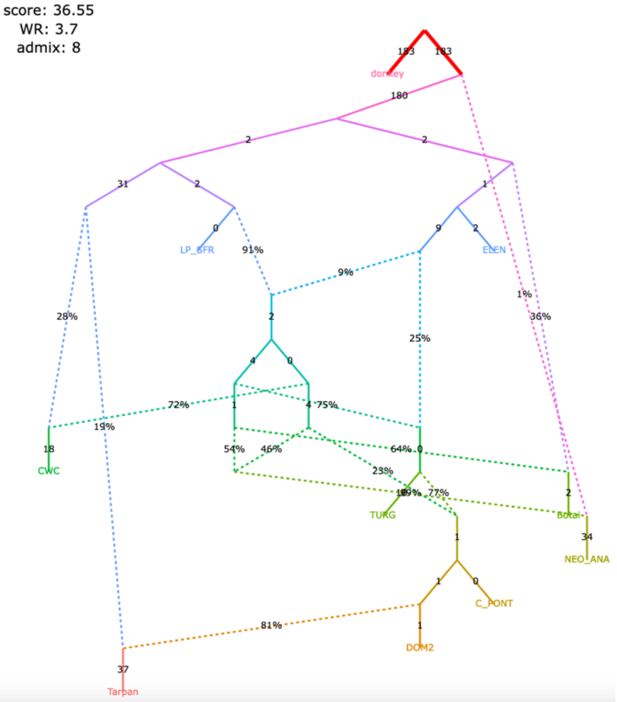

**claim 1** no
**claim 2** yes
**claim 3** yes
**claim 4** yes
**claim 5** no

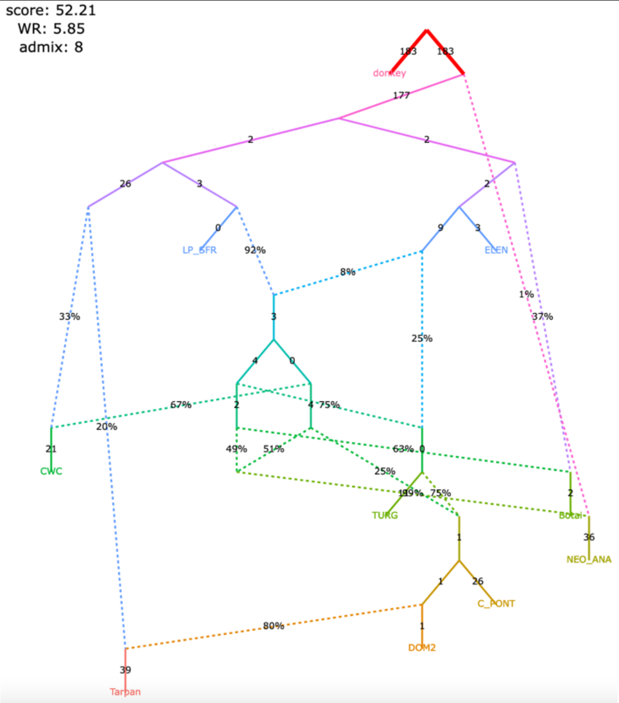

Figure S8

#### **k**, selected models with 8 admixture events (all plausible models with WR < 4 SE)

| claim | |
|---|---|
| **claim 1** | yes |
| **claim 2** | yes |
| **claim 3** | no |
| **claim 4** | yes |
| **claim 5** | yes |

The modified set of individuals and the corrected algorithm for calculating $f_3$-statistics (1,767,419 variable site with no missing data at the group level)

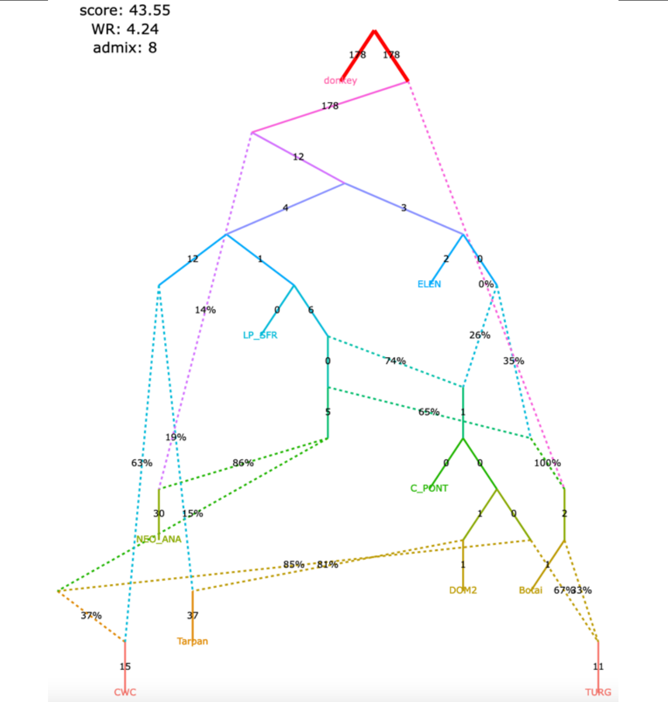

| claim | |
|---|---|
| **claim 1** | yes |
| **claim 2** | yes |
| **claim 3** | no |
| **claim 4** | yes |
| **claim 5** | yes |

The modified set of individuals and the original algorithm for calculating $f_3$-statistics (6,343,116 variable site with no missing data at the group level)

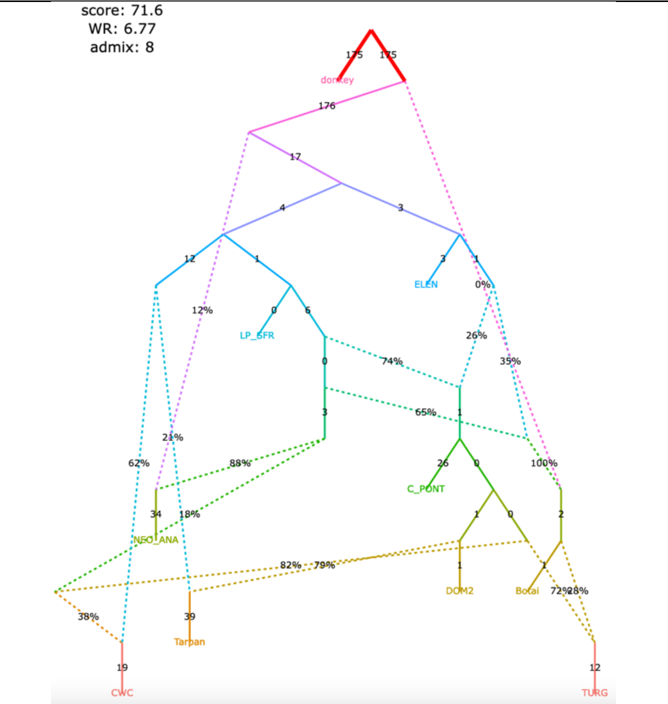

| claim | |
|---|---|
| **claim 1** | yes |
| **claim 2** | yes |
| **claim 3** | yes |
| **claim 4** | yes |
| **claim 5** | yes |

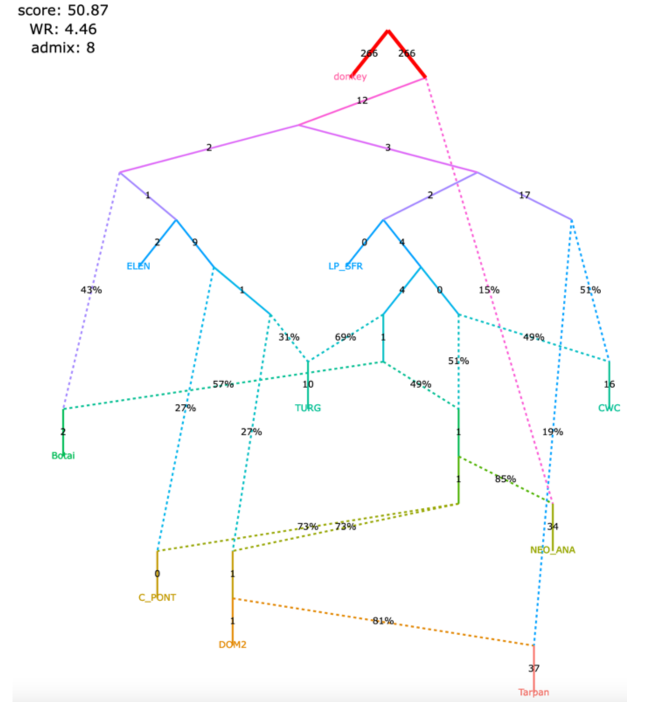

| claim | |
|---|---|
| **claim 1** | yes |
| **claim 2** | yes |
| **claim 3** | yes |
| **claim 4** | yes |
| **claim 5** | yes |

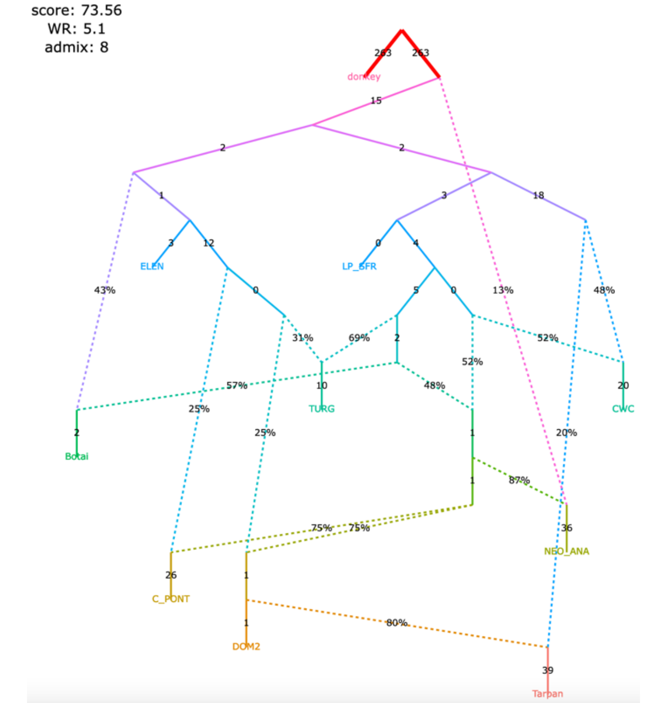

Figure S8

#### I, selected models with 8 admixture events (all plausible models with WR < 4 SE)

The modified set of individuals and the corrected algorithm for calculating $f_3$-statistics (1,767,419 variable site with no missing data at the group level)

| | |
|---|---|
| **claim 1** | yes |
| **claim 2** | yes |
| **claim 3** | no |
| **claim 4** | yes |
| **claim 5** | yes |

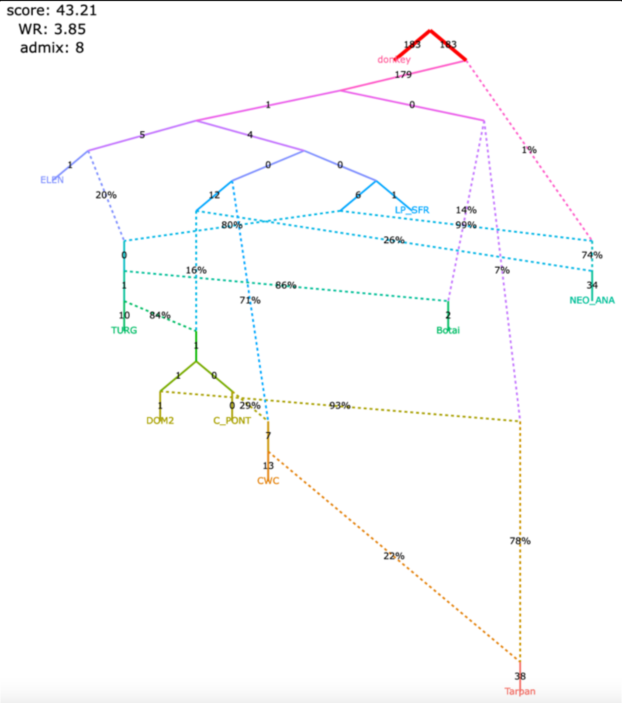

The modified set of individuals and the original algorithm for calculating $f_3$-statistics (6,343,116 variable site with no missing data at the group level)

| | |
|---|---|
| **claim 1** | yes |
| **claim 2** | yes |
| **claim 3** | no |
| **claim 4** | yes |
| **claim 5** | yes |

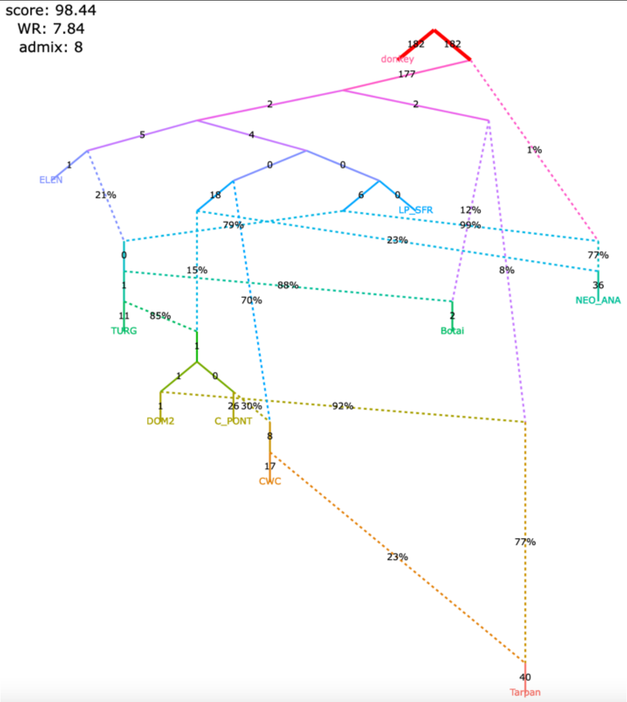

Figure S8

**m**, selected models with 9 admixture events (all plausible models with WR < 4 SE)

| | |
|---|---|
| **claim 1** no<br>**claim 2** yes<br>**claim 3** no<br>**claim 4** no<br>**claim 5** yes | **claim 1** no<br>**claim 2** yes<br>**claim 3** no<br>**claim 4** no<br>**claim 5** yes |
| The modified set of individuals and the corrected algorithm for calculating $f_3$-statistics (1,767,419 variable site with no missing data at the group level) | The modified set of individuals and the original algorithm for calculating $f_3$-statistics (6,343,116 variable site with no missing data at the group level) |
| 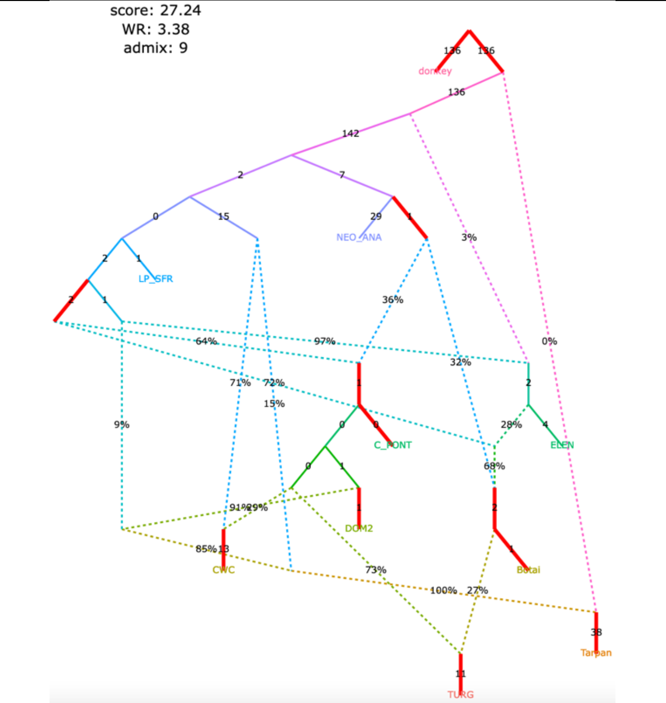 | 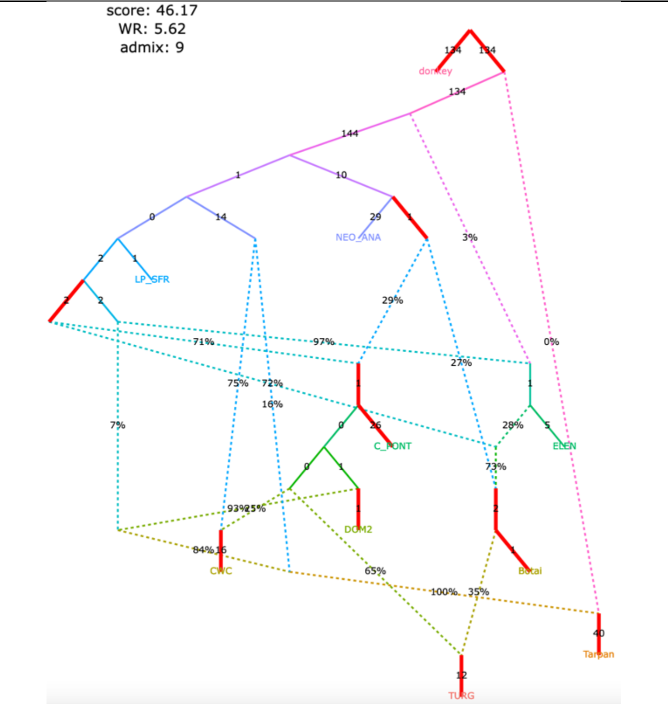 |
| **claim 1** no<br>**claim 2** yes<br>**claim 3** no<br>**claim 4** yes<br>**claim 5** no | **claim 1** no<br>**claim 2** yes<br>**claim 3** no<br>**claim 4** yes<br>**claim 5** no |
| 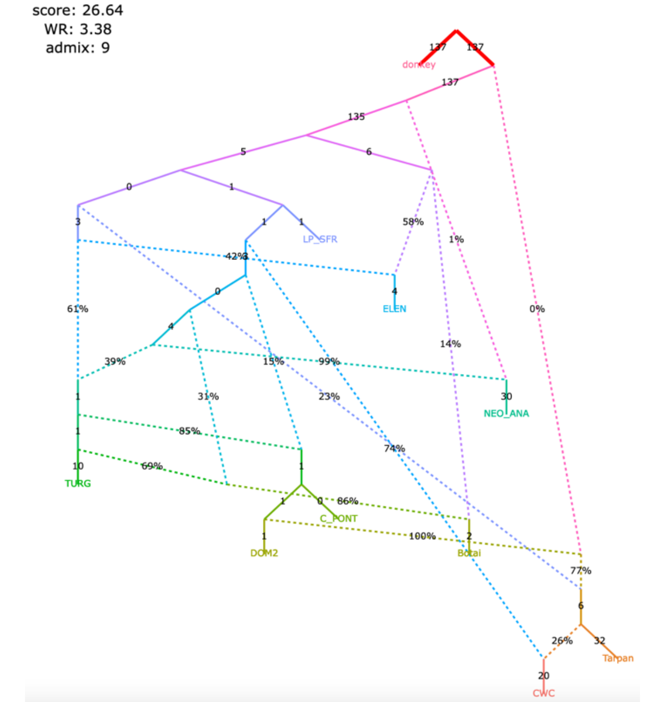 | 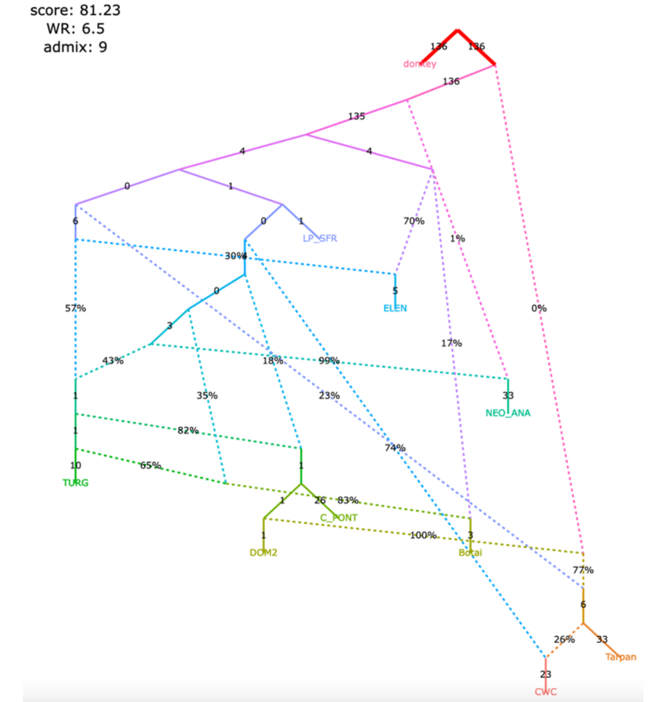 |

Figure S8

#### **n**, selected models with 9 admixture events (all plausible models with WR < 4 SE)

| |
|---|
| **claim 1** yes |
| **claim 2** yes |
| **claim 3** no |
| **claim 4** yes |
| **claim 5** no |

| |
|---|
| **claim 1** yes |
| **claim 2** yes |
| **claim 3** no |
| **claim 4** yes |
| **claim 5** no |

The modified set of individuals and the corrected algorithm for calculating $f_3$-statistics (1,767,419 variable site with no missing data at the group level)

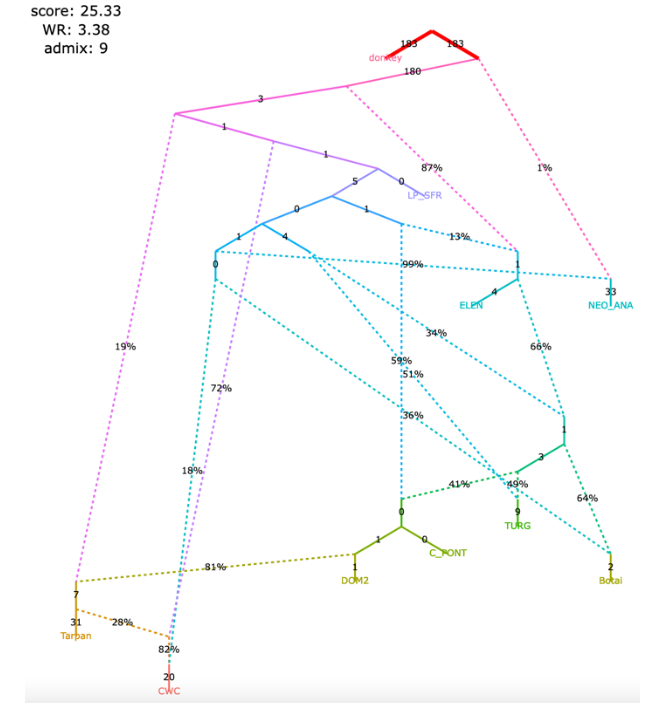

The modified set of individuals and the original algorithm for calculating $f_3$-statistics (6,343,116 variable site with no missing data at the group level)

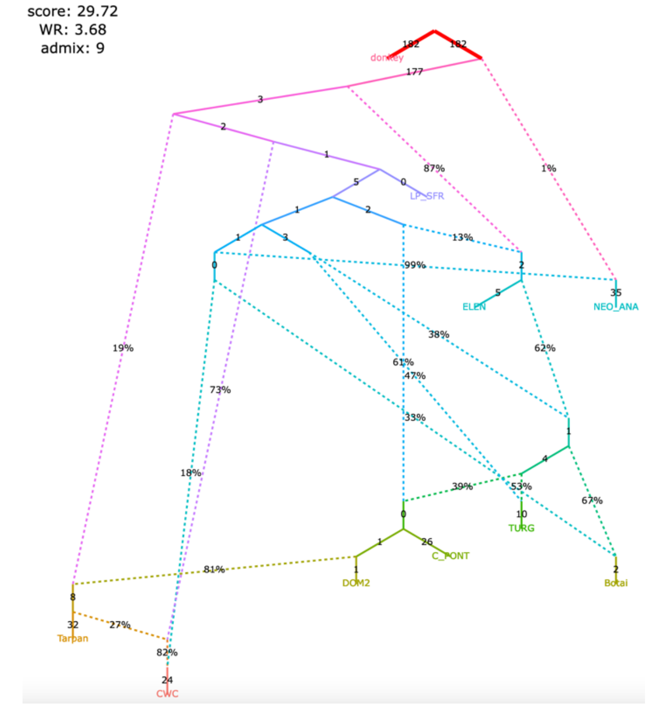

| |
|---|
| **claim 1** no |
| **claim 2** yes |
| **claim 3** no |
| **claim 4** yes |
| **claim 5** no |

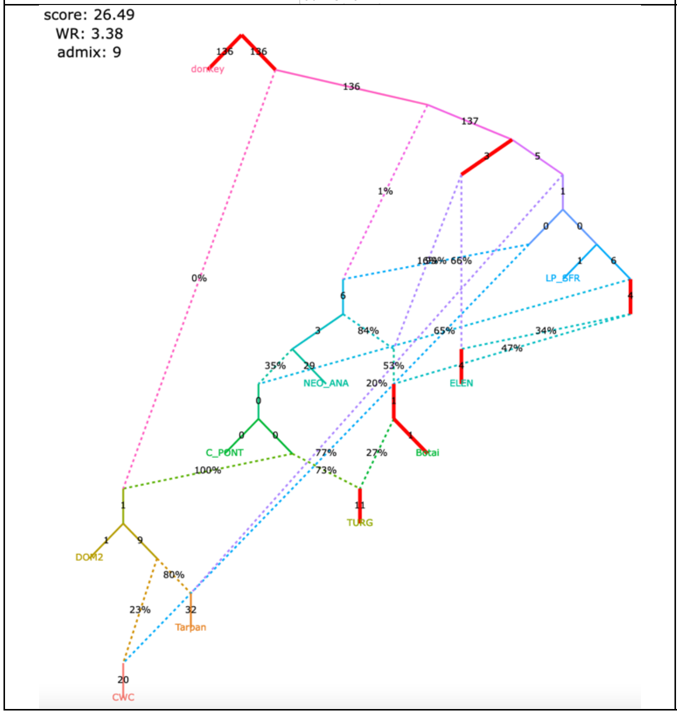

| |
|---|
| **claim 1** no |
| **claim 2** yes |
| **claim 3** no |
| **claim 4** yes |
| **claim 5** no |

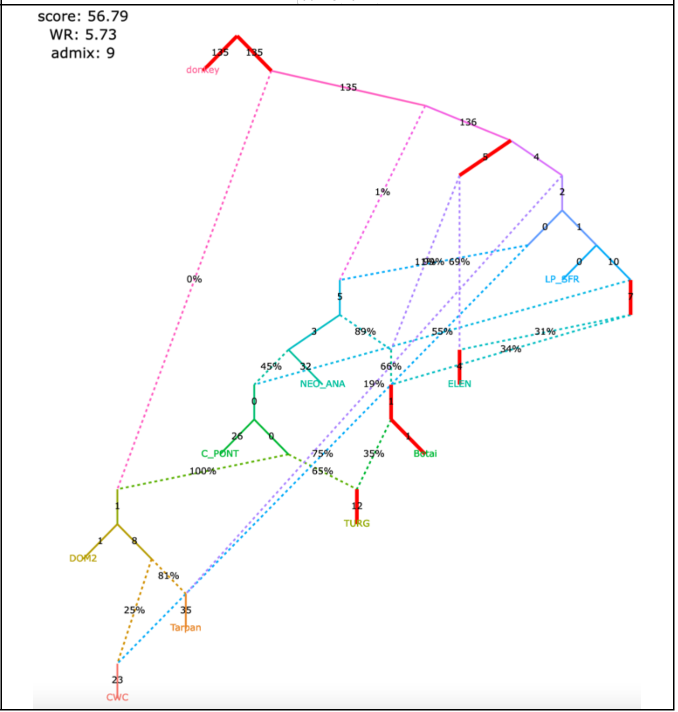

Figure S8

**o**, selected models with 9 admixture events (all plausible models with WR < 4 SE)

| | |
|---|---|
| **claim 1** yes<br>**claim 2** yes<br>**claim 3** no<br>**claim 4** no<br>**claim 5** yes | **claim 1** yes<br>**claim 2** yes<br>**claim 3** no<br>**claim 4** no<br>**claim 5** yes |
| The modified set of individuals and the corrected algorithm for calculating *f*$_3$-statistics (1,767,419 variable site with no missing data at the group level) | The modified set of individuals and the original algorithm for calculating *f*$_3$-statistics (6,343,116 variable site with no missing data at the group level) |

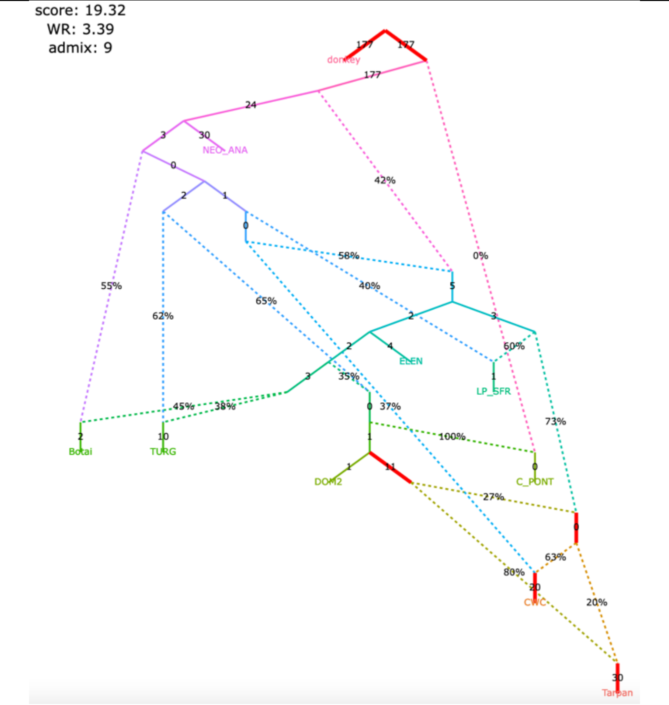

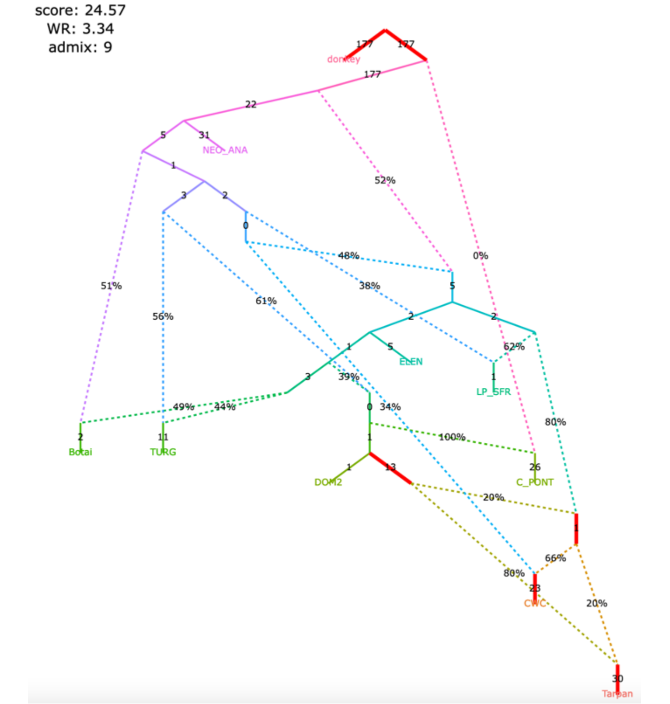

| | |
|---|---|
| **claim 1** yes<br>**claim 2** yes<br>**claim 3** no<br>**claim 4** yes<br>**claim 5** no | **claim 1** yes<br>**claim 2** yes<br>**claim 3** no<br>**claim 4** yes<br>**claim 5** no |

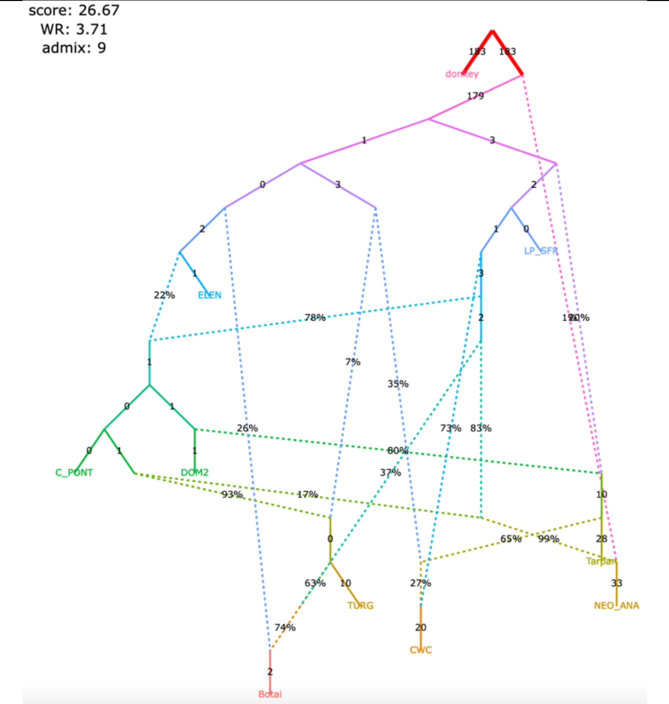

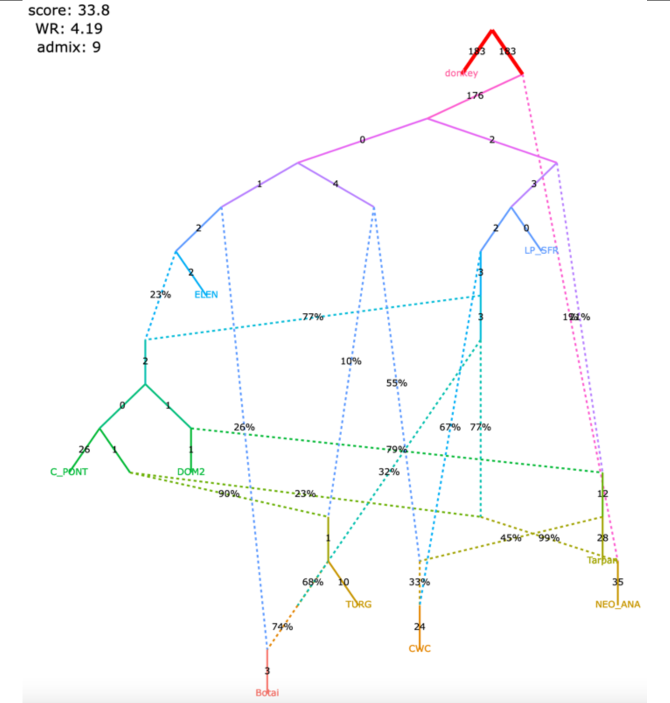

Figure S8

**p**, selected models with 9 admixture events (all plausible models with WR < 4 SE)

| | |
|---|---|
| claim 1 yes<br>claim 2 yes<br>claim 3 yes<br>claim 4 yes<br>claim 5 no | claim 1 yes<br>claim 2 yes<br>claim 3 yes<br>claim 4 yes<br>claim 5 no |
| The modified set of individuals and the corrected algorithm for calculating $f_3$-statistics (1,767,419 variable site with no missing data at the group level) | The modified set of individuals and the original algorithm for calculating $f_3$-statistics (6,343,116 variable site with no missing data at the group level) |

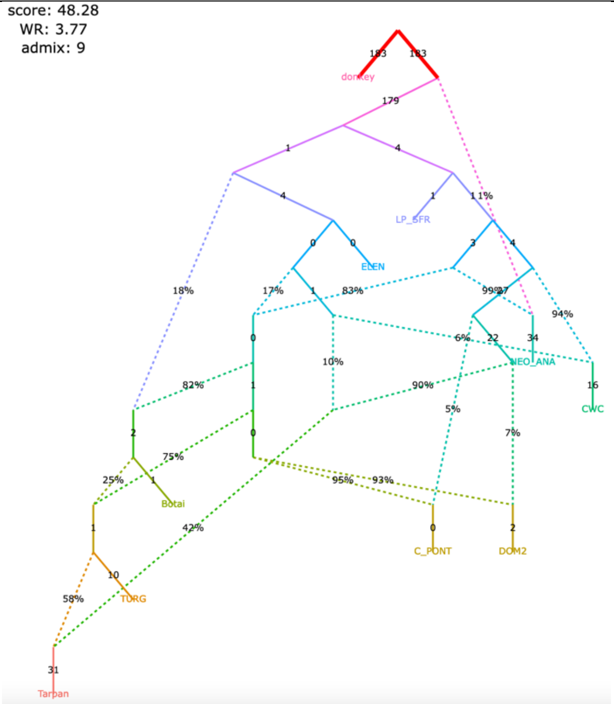

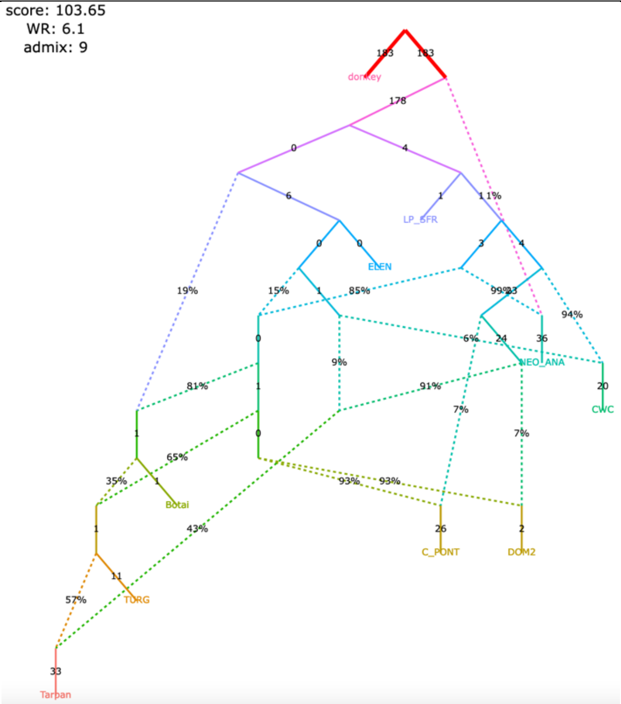

| | |
|---|---|
| claim 1 yes<br>claim 2 yes<br>claim 3 no<br>claim 4 yes<br>claim 5 yes | claim 1 yes<br>claim 2 yes<br>claim 3 no<br>claim 4 yes<br>claim 5 yes |

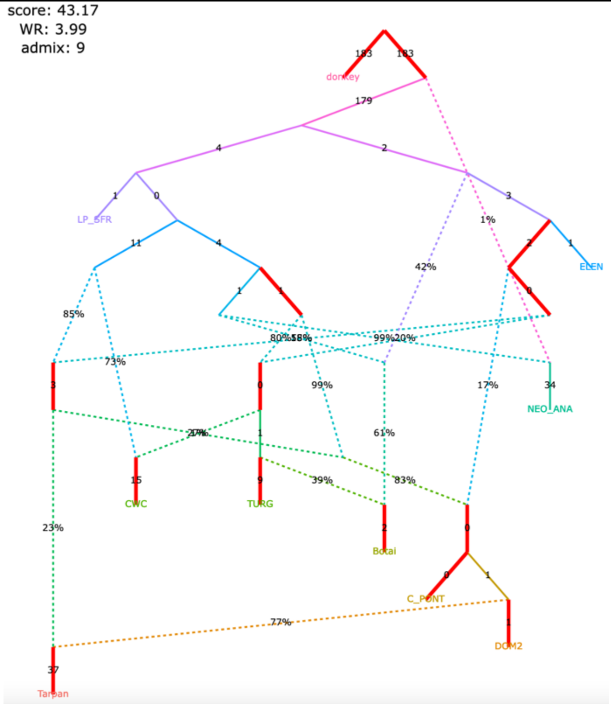

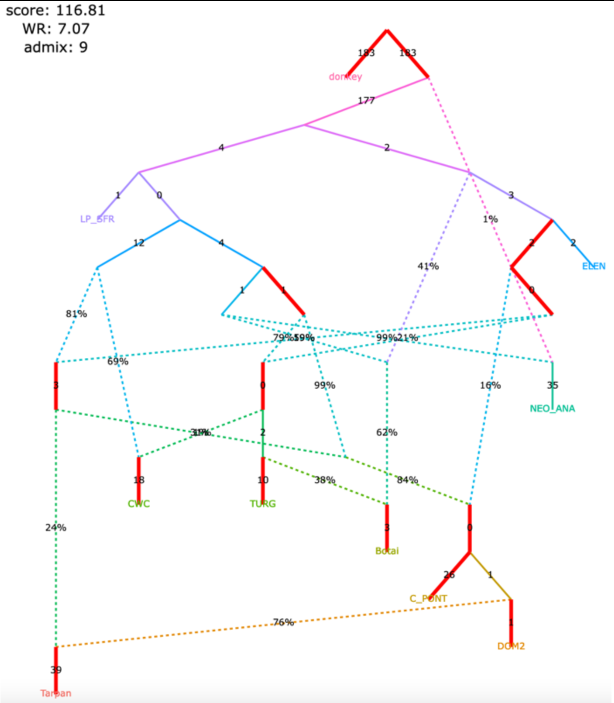

Figure S8

#### **q**, selected models with 9 admixture events (all plausible models with WR < 4 SE)

| claim 1 yes<br>claim 2 yes<br>claim 3 no<br>claim 4 yes<br>claim 5 yes | claim 1 yes<br>claim 2 yes<br>claim 3 no<br>claim 4 yes<br>claim 5 yes |
|---|---|
| The modified set of individuals and the corrected algorithm for calculating $f_3$-statistics (1,767,419 variable site with no missing data at the group level) | The modified set of individuals and the original algorithm for calculating $f_3$-statistics (6,343,116 variable site with no missing data at the group level) |
| 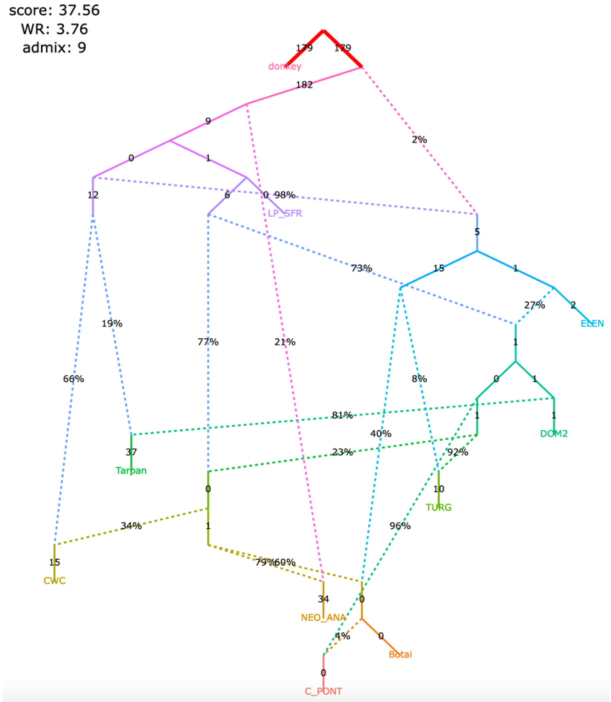 | 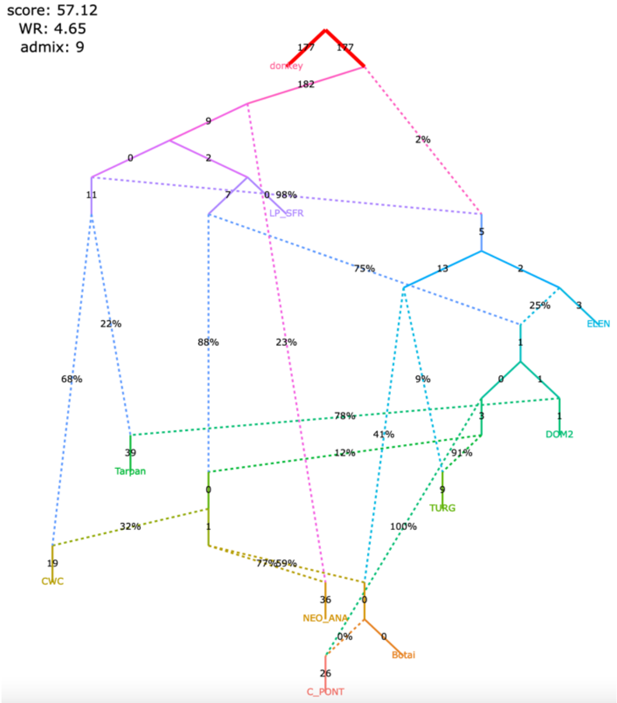 |
| claim 1 yes<br>claim 2 yes<br>claim 3 no<br>claim 4 yes<br>claim 5 no | claim 1 yes<br>claim 2 yes<br>claim 3 no<br>claim 4 yes<br>claim 5 no |
| 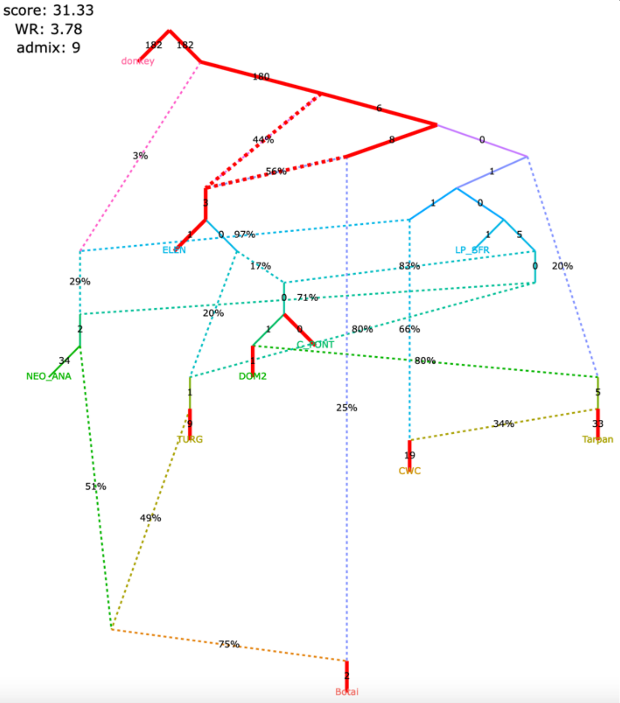 | 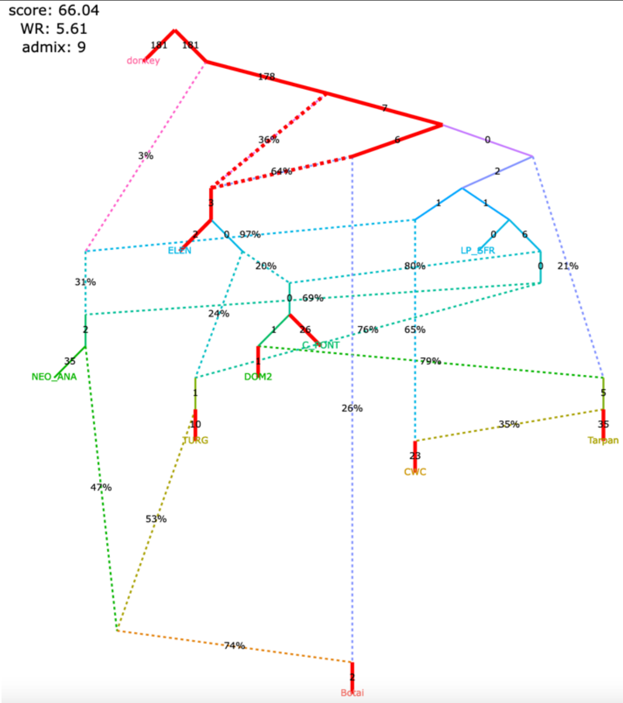 |

Figure S8

**r**, selected models with 9 admixture events (all plausible models with WR < 4 SE)

| | |
|---|---|
| **claim 1** no<br>**claim 2** no<br>**claim 3** no<br>**claim 4** no<br>**claim 5** yes | **claim 1** no<br>**claim 2** no<br>**claim 3** no<br>**claim 4** no<br>**claim 5** yes |
| The modified set of individuals and the corrected algorithm for calculating *f*$_3$-statistics (1,767,419 variable site with no missing data at the group level) | The modified set of individuals and the original algorithm for calculating *f*$_3$-statistics (6,343,116 variable site with no missing data at the group level) |
| 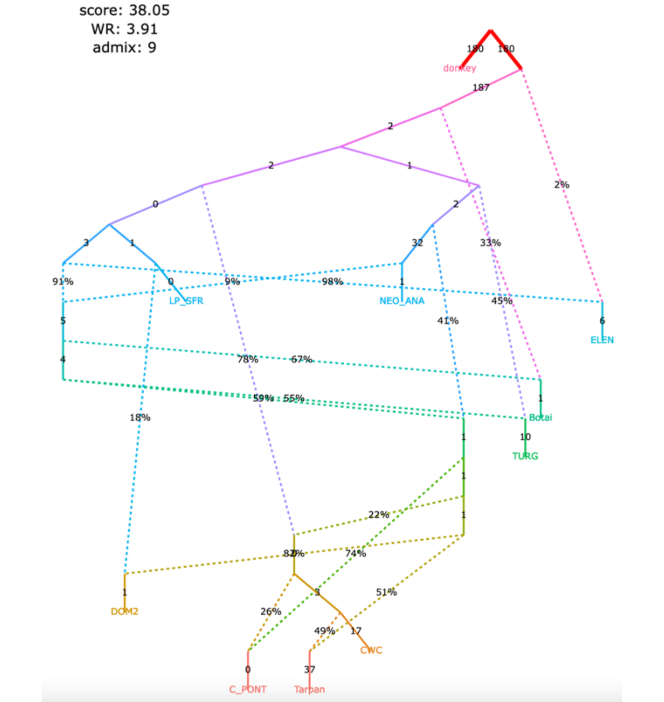  | 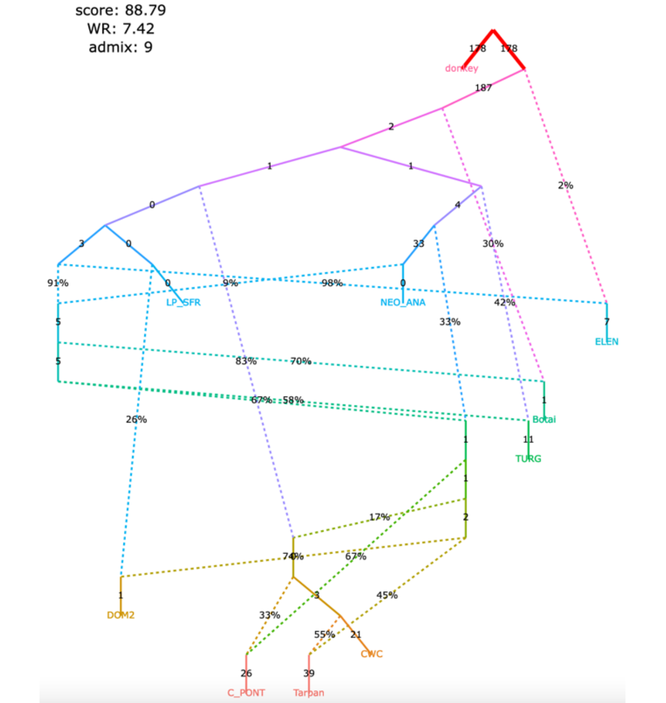  |

Figure S9

**Figure S9.** Published admixture graph from Hajdinjak *et al*. (2021) and alternative graphs found with *findGraphs* (12 populations, 8 admixture events).

**a**, published model

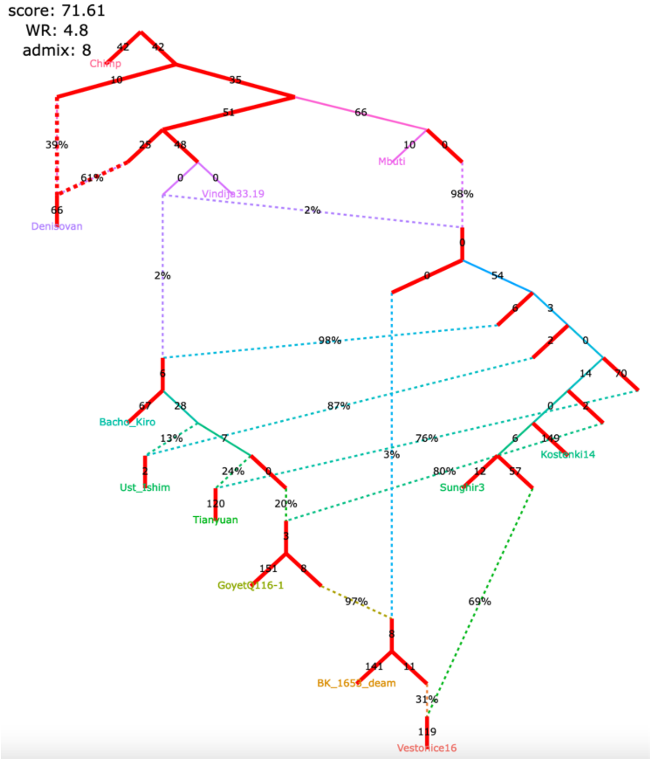

**b**, published model, simplified

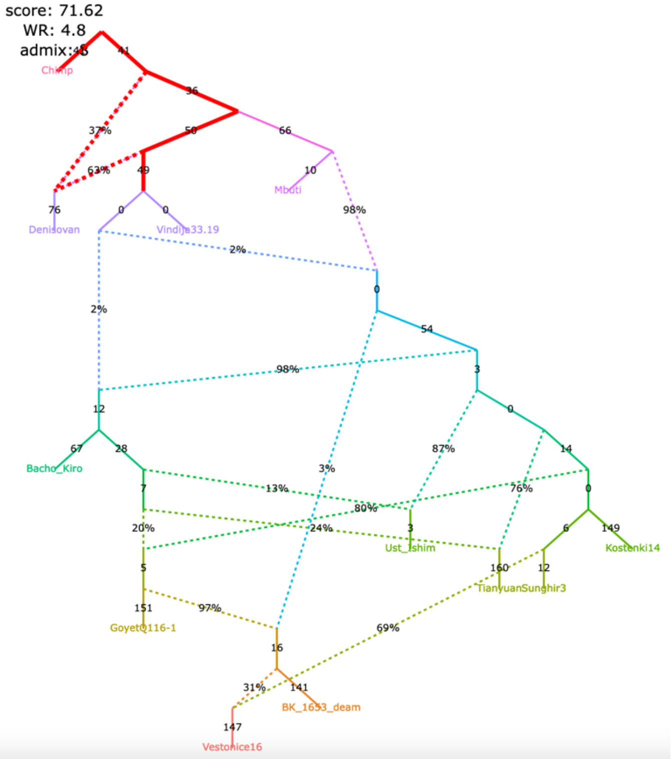

Claims by Hajdinjak et al. 2021 relying on the admixture graph:

1) Gene flow from the Bacho Kiro lineage to Ust'-Ishim, Tianyuan, and GoyetQ116-1;
2) the BK1653 individual belonged to a population that was related, but not identical, to that of the GoyetQ116-1 individual;
3) Vestonice is a mixture of a Sunghir-related and a BK1653-related lineage.

Figure S9

**c**, selected alternative models fitting significantly better than the published one

claim 1 partially supported, claims 2&3 supported

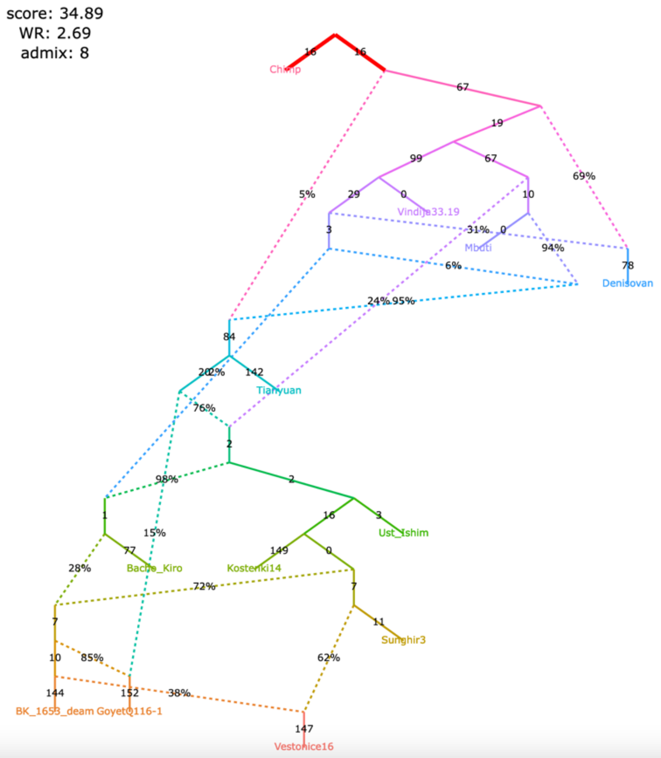

claim 1 not supported, claims 2&3 supported

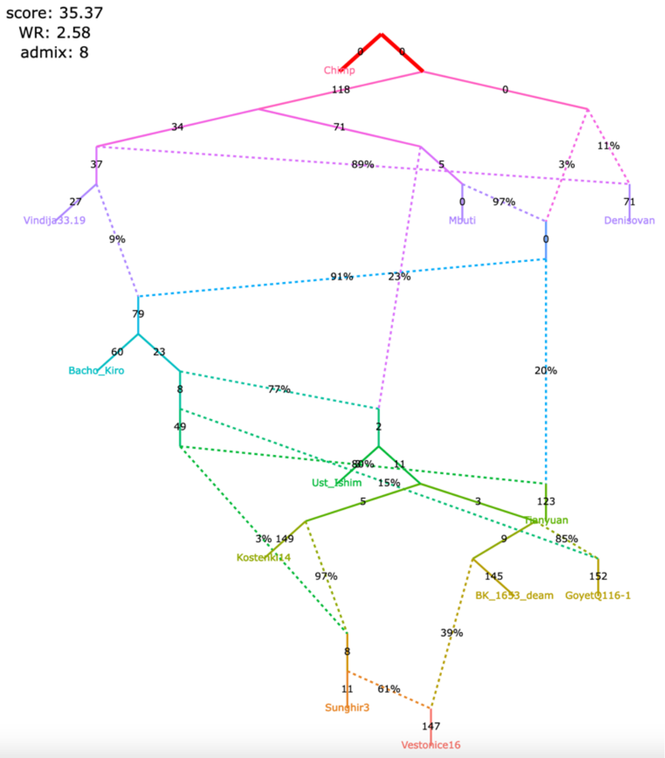

claim 1 not supported, claims 2&3 supported

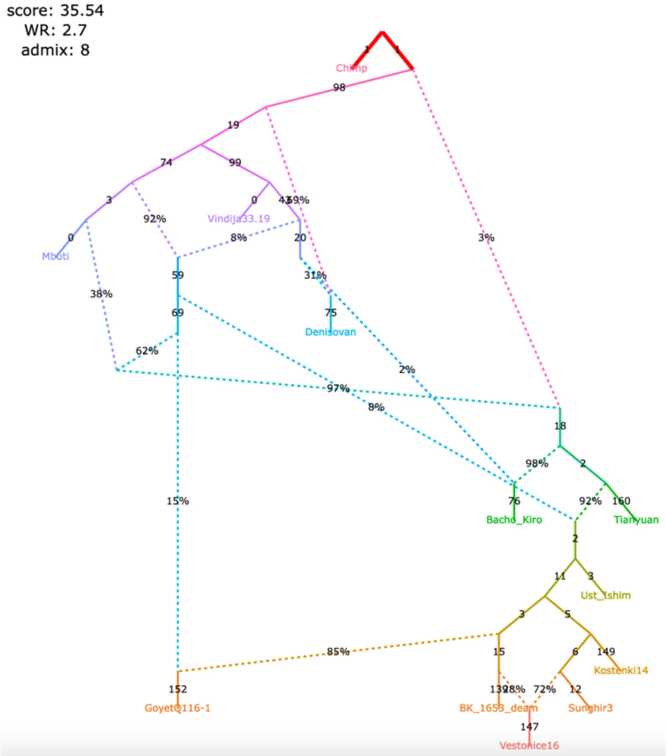

claim 1 not supported, claims 2&3 supported

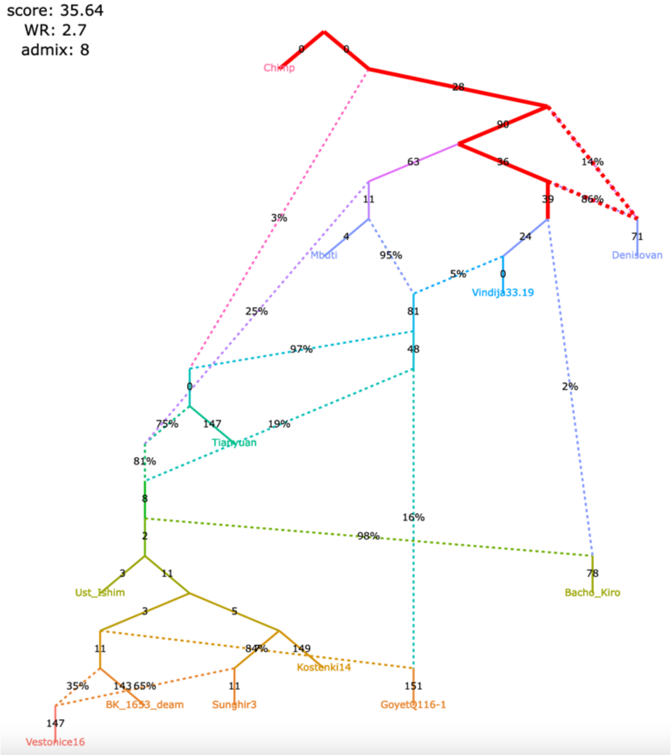

Figure S9

**d**, selected alternative models fitting significantly better than the published one

claim 1 not supported, claims 2&3 supported

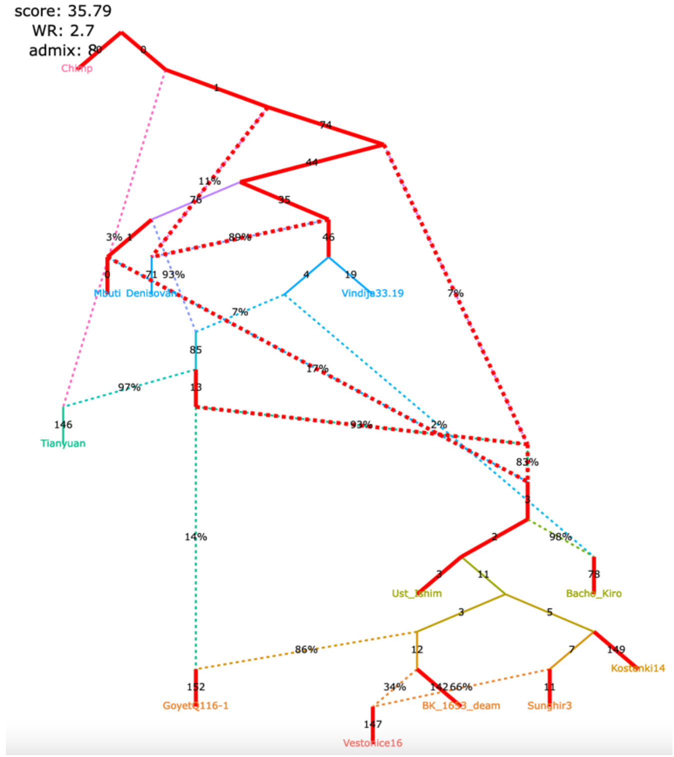

claim 1 not supported, claims 2&3 supported

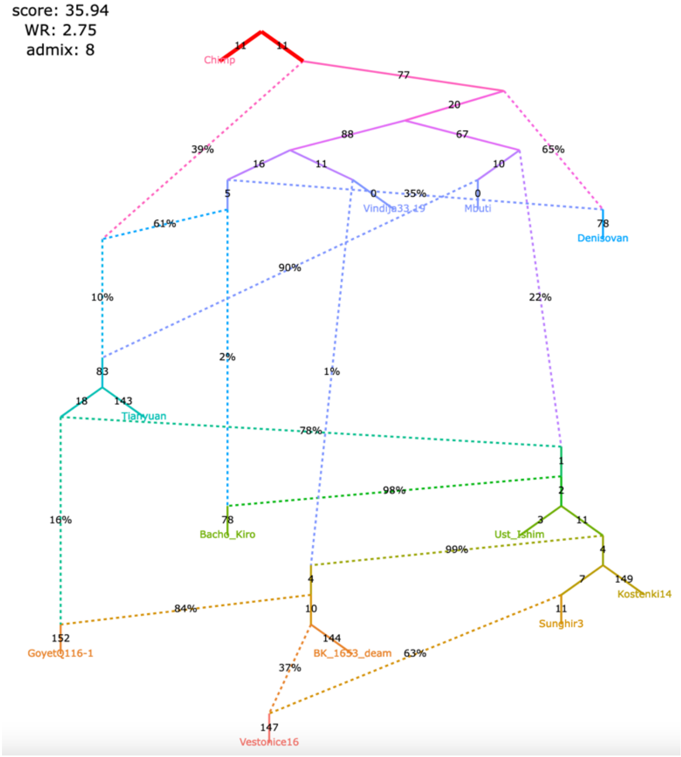

claims 1&2 not supported, claim 3 supported

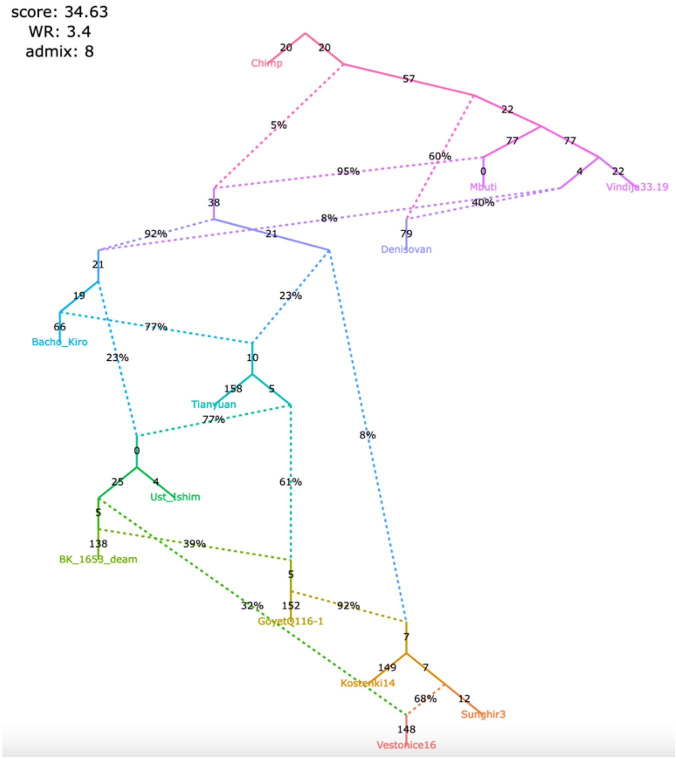

claims 1&2 not supported, claim 3 supported

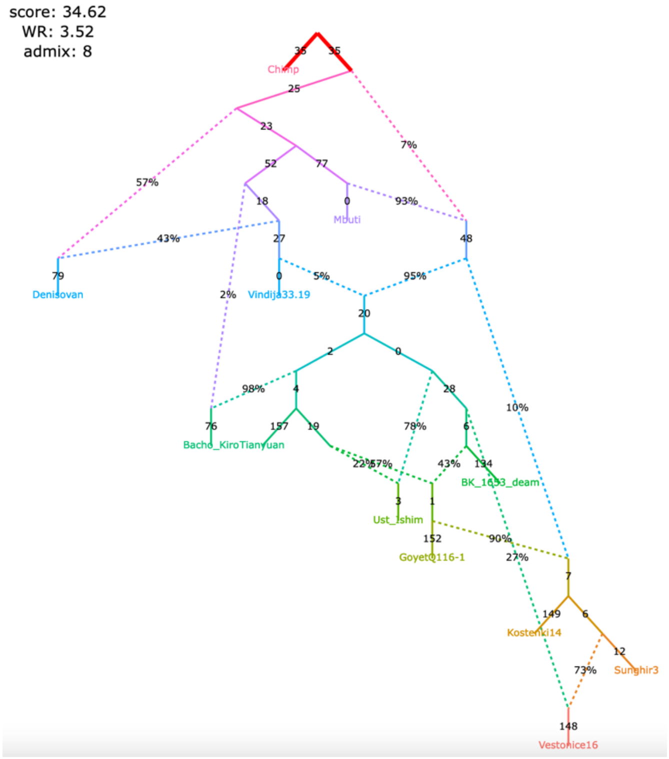

Figure S9

**e**, selected alternative models fitting significantly better than the published one

claims 1&2 not supported, claim 3 supported

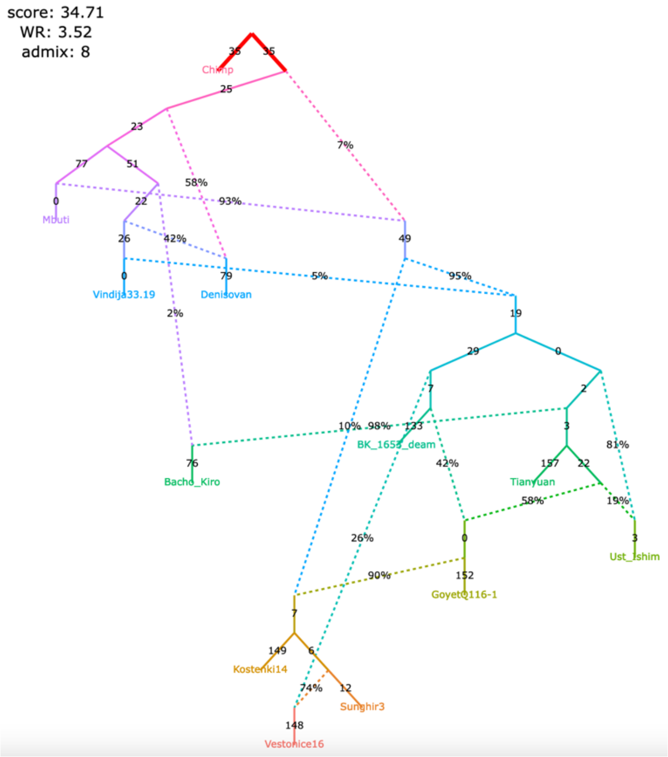

claims 1&2 not supported, claim 3 supported

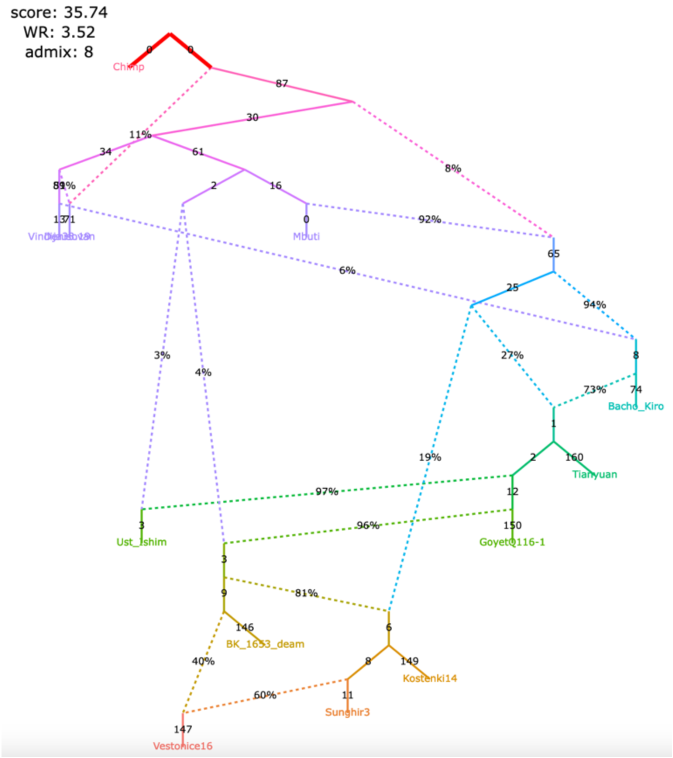

claims 1&2 not supported, claim 3 supported

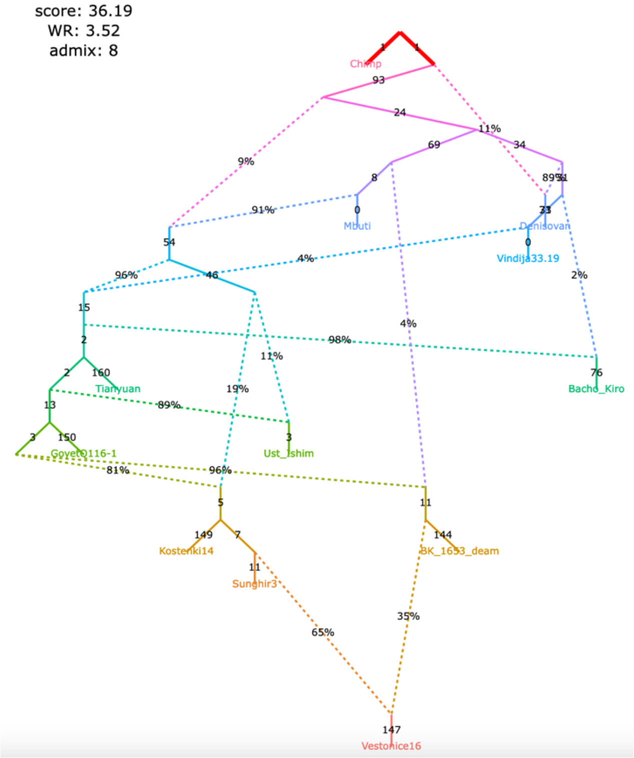

Figure S10

**Figure S10.** Published admixture graph from Lipson *et al*. (2020) and alternative graphs found with *findGraphs* (12 populations, 11 admixture events) using the updated algorithm for calculating *f*-statistics. The graphs were also re-fitted using the original algorithm for calculating *f*-statistics.

**a**, published model

| | |
|---|---|
| no. of East Asian groups with Onge-related ancestry | 6 |
| no. of East Asian groups with European-related ancestry | 1 |
| direction of gene flow China_Island_EN > Taiwan_IA | forward |
| Mongolia_East is basal with respect to China_Upper_YR and Nepal_Chokhopani | yes |
| Taiwan_IA is basal with respect to Mongolia_East and Nepal_Chokhopani | yes |

The original set of SNPs and individuals and the corrected algorithm for calculating $f_3$-statistics (211,738 variable site with no missing data at the group level)

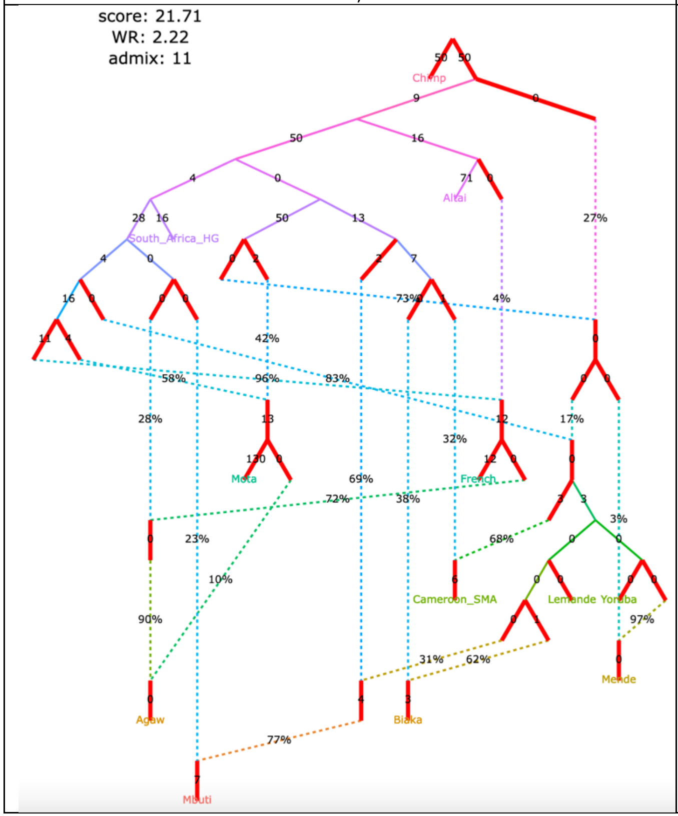

| | |
|---|---|
| no. of East Asian groups with Onge-related ancestry | 6 |
| no. of East Asian groups with European-related ancestry | 1 |
| direction of gene flow China_Island_EN > Taiwan_IA | forward |
| Mongolia_East is basal with respect to China_Upper_YR and Nepal_Chokhopani | yes |
| Taiwan_IA is basal with respect to Mongolia_East and Nepal_Chokhopani | yes |

The original set of SNPs and individuals and the original algorithm for calculating $f_3$-statistics (363,131 variable site with no missing data at the group level)

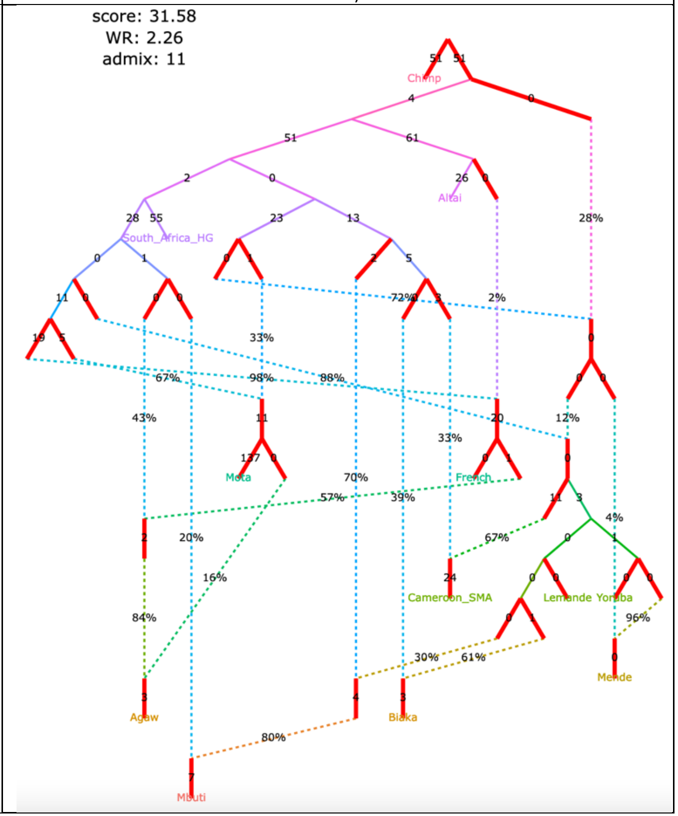

Figure S10

**b**, published model, simplified

| | |
|---|---|
| no. of East Asian groups with Onge-related ancestry 6<br>no. of East Asian groups with European-related ancestry 1<br>direction of gene flow China_Island_EN > Taiwan_IA forward<br>Mongolia_East is basal with respect to China_Upper_YR and Nepal_Chokhopani yes<br>Taiwan_IA is basal with respect to Mongolia_East and Nepal_Chokhopani yes | no. of East Asian groups with Onge-related ancestry 6<br>no. of East Asian groups with European-related ancestry 1<br>direction of gene flow China_Island_EN > Taiwan_IA forward<br>Mongolia_East is basal with respect to China_Upper_YR and Nepal_Chokhopani yes<br>Taiwan_IA is basal with respect to Mongolia_East and Nepal_Chokhopani yes |
| The original set of SNPs and individuals and the corrected algorithm for calculating $f_3$-statistics (211,738 variable site with no missing data at the group level) | The original set of SNPs and individuals and the original algorithm for calculating $f_3$-statistics (363,131 variable site with no missing data at the group level) |

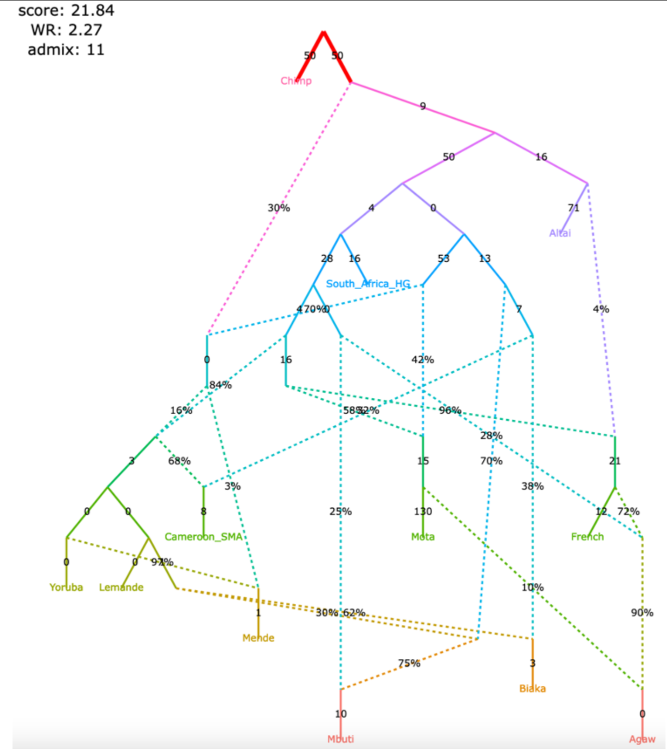

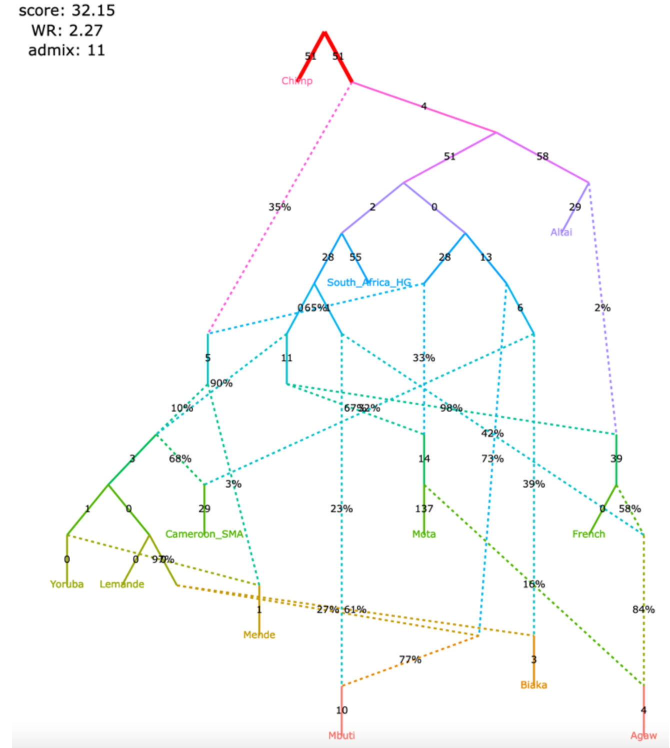

Selected claims by Lipson et al. 2020 relying on the admixture graph:

1) A lineage maximized in present-day West African groups (Lemande, Mende, and Yoruba) also contributed some ancestry to the ancient Shum Laka individual, and present-day Biaka and Mbuti;
2) another ancestry component in Shum Laka is a deep-branching lineage maximized in rainforest hunter-gatherers Biaka and Mbuti;
3) "super-archaic" ancestry (i.e., diverging at the modern human/Neanderthal split point or deeper) contributed to Biaka, Shum Laka, Mbuti, Lemande, Mende, and Yoruba;
4) a ghost modern human lineage (or lineages) contributed to Agaw, Mota, Biaka, Shum Laka, Mbuti, Lemande, Mende, and Yoruba.

Figure S10

#### **c**, selected alternative models fitting nominally better than the published one

| | |
|---|---|
| **support for claim 1** partial<br>**support for claim 2** not supported<br>**support for claim 3** not supported<br>**support for claim 4** not supported | **support for claim 1** partial<br>**support for claim 2** not supported<br>**support for claim 3** not supported<br>**support for claim 4** not supported |
| The original set of SNPs and individuals and the corrected algorithm for calculating $f_3$-statistics (211,738 variable site with no missing data at the group level) | The original set of SNPs and individuals and the original algorithm for calculating $f_3$-statistics (363,131 variable site with no missing data at the group level) |
| 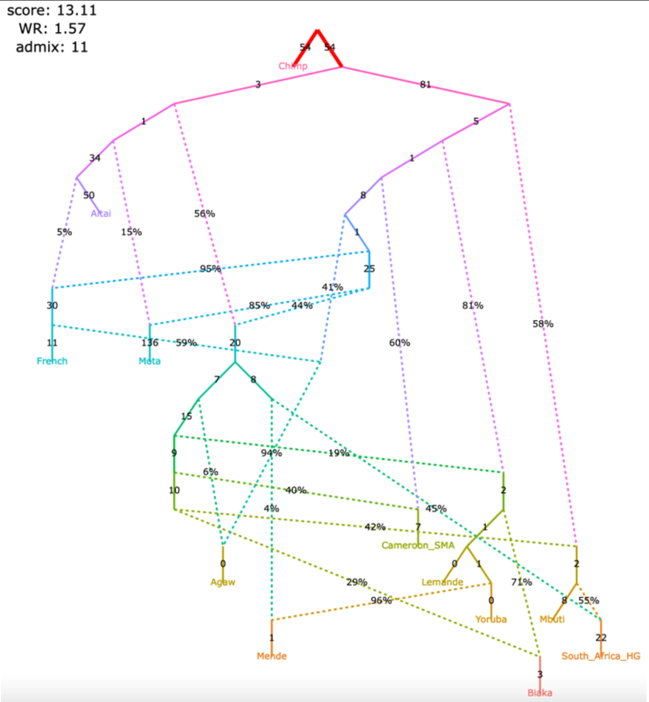  | 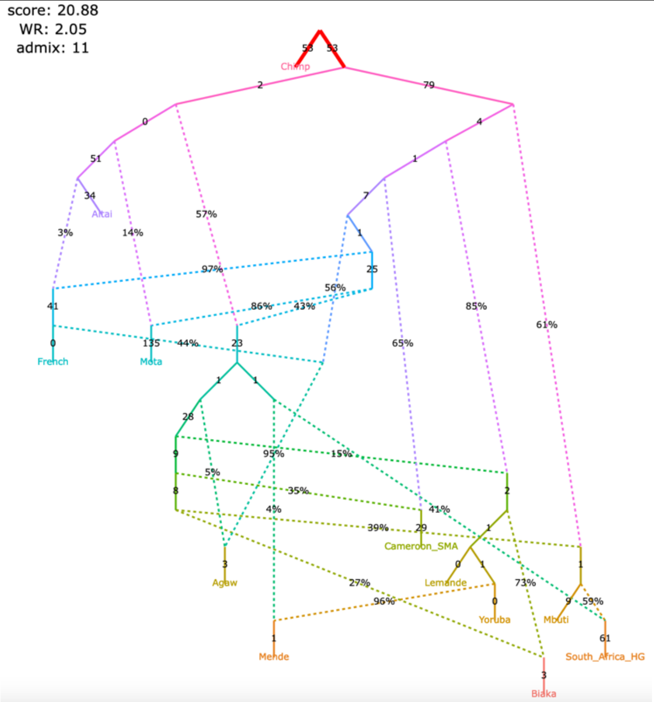  |
| **support for claim 1** not supported<br>**support for claim 2** not supported<br>**support for claim 3** partial<br>**support for claim 4** not supported | **support for claim 1** not supported<br>**support for claim 2** not supported<br>**support for claim 3** partial<br>**support for claim 4** not supported |
| 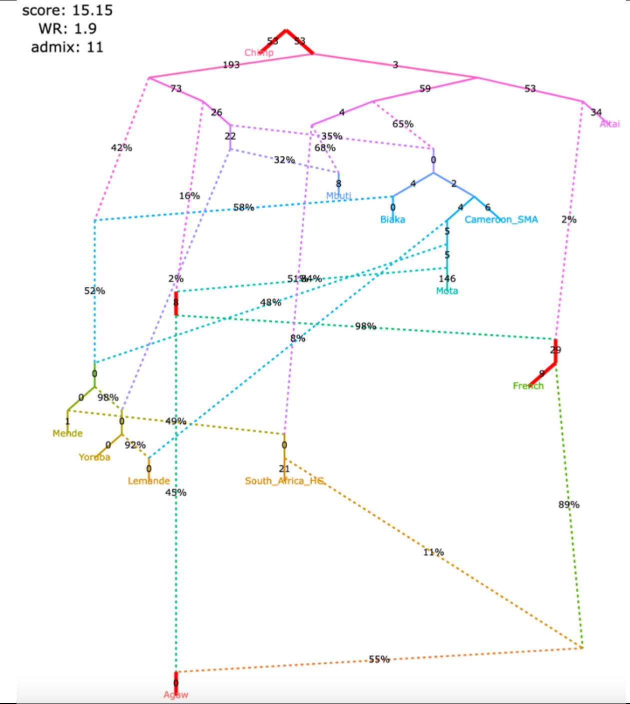  | 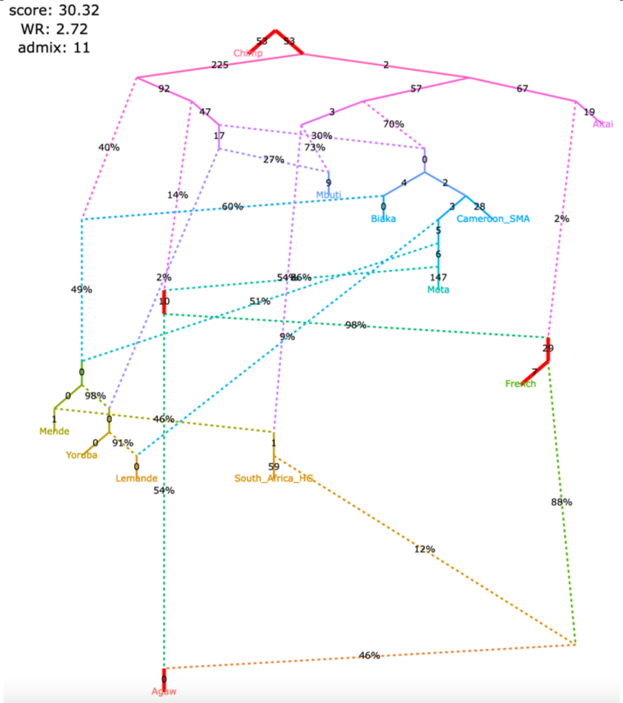  |

Figure S10

#### **d**, selected alternative models fitting nominally better than the published one

| | |
|---|---|
| **support for claim 1** | not supported |
| **support for claim 2** | partial |
| **support for claim 3** | partial |
| **support for claim 4** | not supported |

The original set of SNPs and individuals and the corrected algorithm for calculating $f_3$-statistics (211,738 variable site with no missing data at the group level)

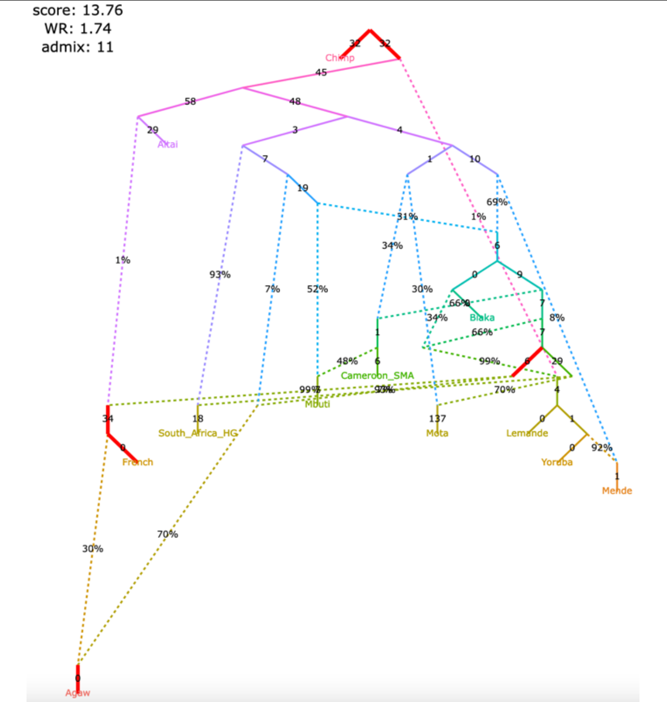

| | |
|---|---|
| **support for claim 1** | not supported |
| **support for claim 2** | partial |
| **support for claim 3** | partial |
| **support for claim 4** | not supported |

The original set of SNPs and individuals and the original algorithm for calculating $f_3$-statistics (363,131 variable site with no missing data at the group level)

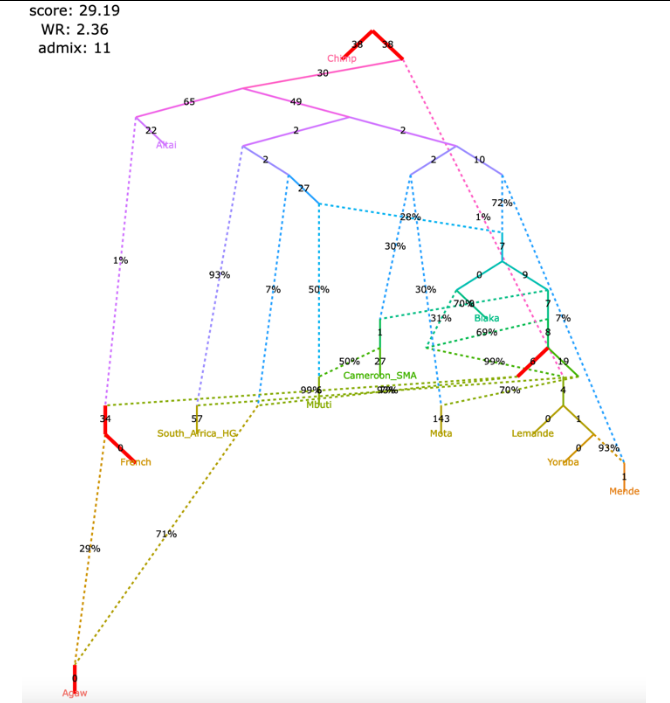

| | |
|---|---|
| **support for claim 1** | partial |
| **support for claim 2** | not supported |
| **support for claim 3** | not supported |
| **support for claim 4** | partial |

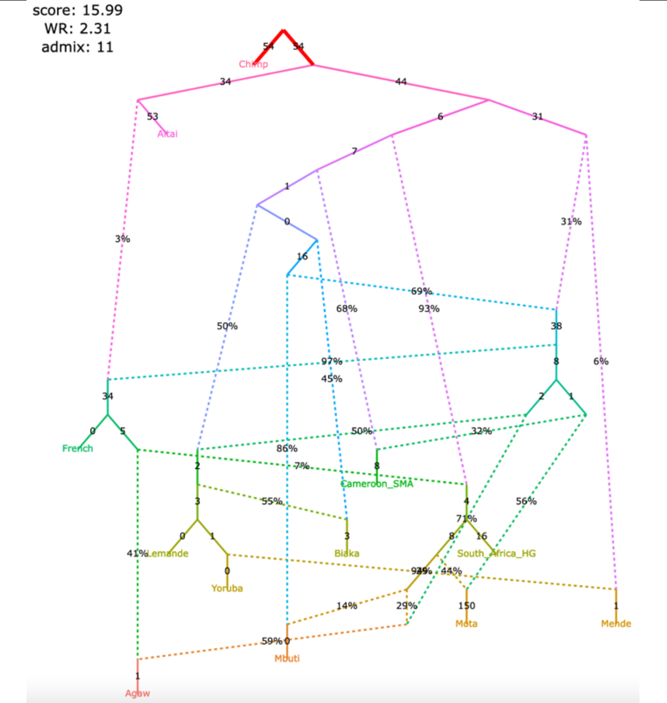

| | |
|---|---|
| **support for claim 1** | partial |
| **support for claim 2** | not supported |
| **support for claim 3** | not supported |
| **support for claim 4** | partial |

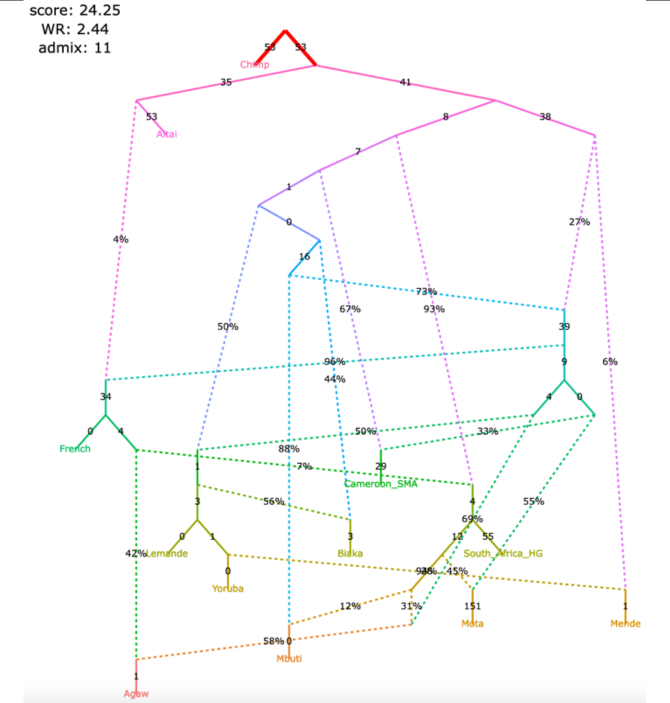

Figure S10

#### **e**, selected alternative models fitting nominally better than the published one

| | |
|---|---|
| **support for claim 1** supported<br>**support for claim 2** not supported<br>**support for claim 3** partial<br>**support for claim 4** not supported | **support for claim 1** supported<br>**support for claim 2** not supported<br>**support for claim 3** partial<br>**support for claim 4** not supported |
| The original set of SNPs and individuals and the corrected algorithm for calculating $f_3$-statistics (211,738 variable site with no missing data at the group level) | The original set of SNPs and individuals and the original algorithm for calculating $f_3$-statistics (363,131 variable site with no missing data at the group level) |
| score: 13.97<br>WR: 1.67<br>admix: 11 | score: 19.95<br>WR: 2.22<br>admix: 11 |

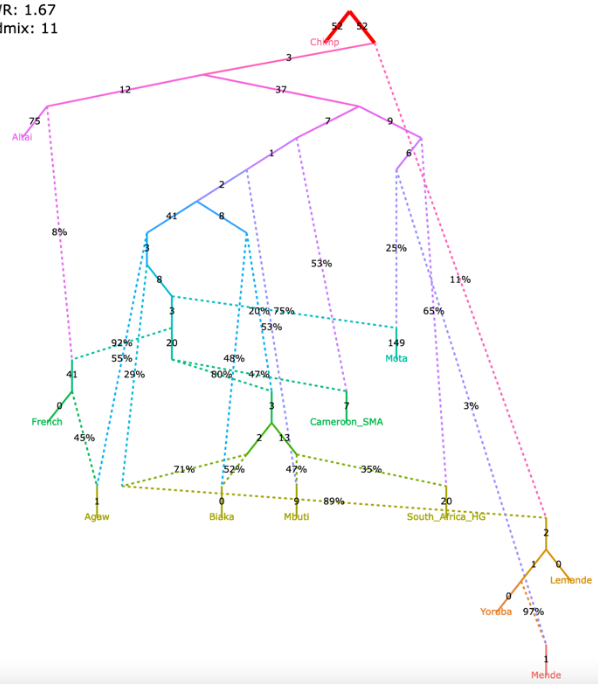

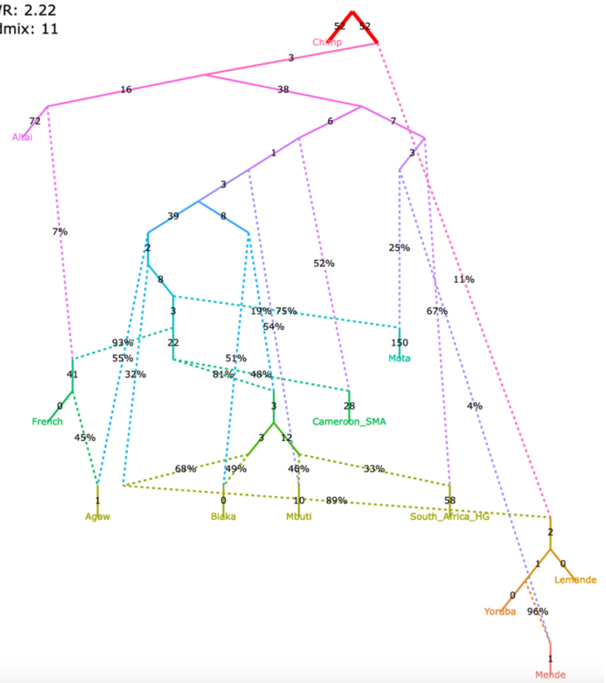

| | |
|---|---|
| **support for claim 1** not supported<br>**support for claim 2** partial<br>**support for claim 3** partial<br>**support for claim 4** partial | **support for claim 1** not supported<br>**support for claim 2** partial<br>**support for claim 3** partial<br>**support for claim 4** partial |
| score: 14.62<br>WR: 1.69<br>admix: 11 | score: 28.87<br>WR: 2.45<br>admix: 11 |

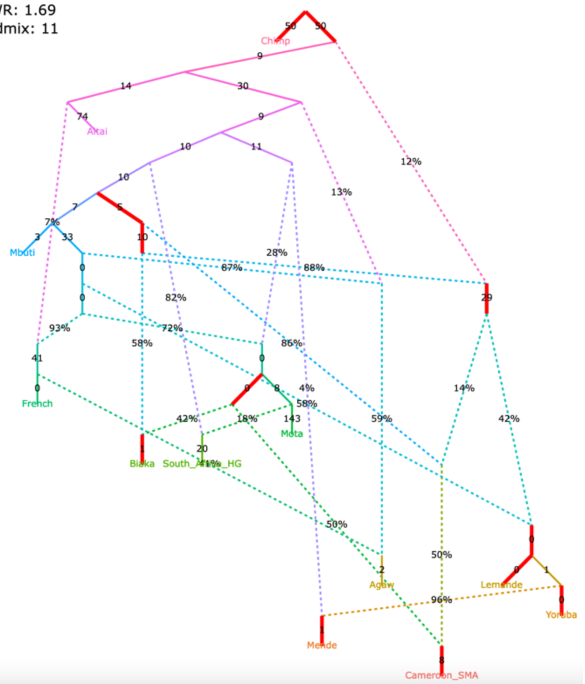

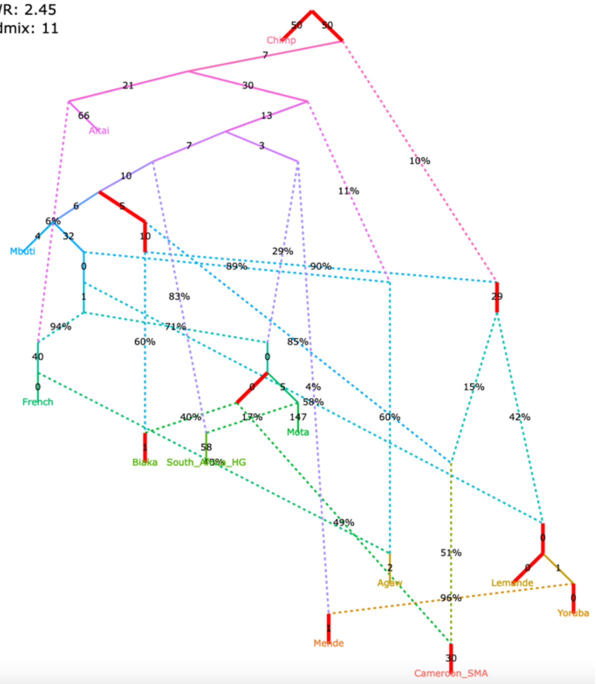

Figure S10

#### **f**, selected alternative models fitting nominally better than the published one

| | |
|---|---|
| **support for claim 1** partial<br>**support for claim 2** partial<br>**support for claim 3** not supported<br>**support for claim 4** partial | **support for claim 1** partial<br>**support for claim 2** partial<br>**support for claim 3** not supported<br>**support for claim 4** partial |
| The original set of SNPs and individuals and the corrected algorithm for calculating $f_3$-statistics (211,738 variable site with no missing data at the group level) | The original set of SNPs and individuals and the original algorithm for calculating $f_3$-statistics (363,131 variable site with no missing data at the group level) |

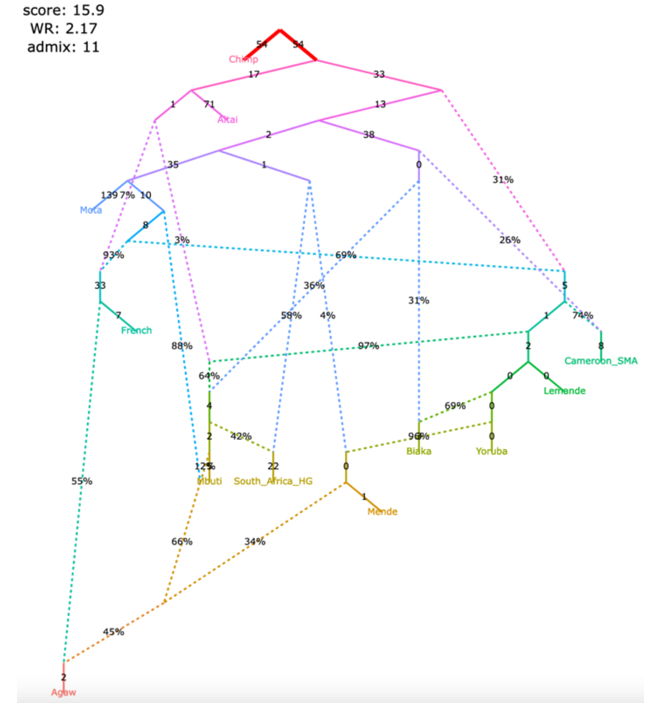

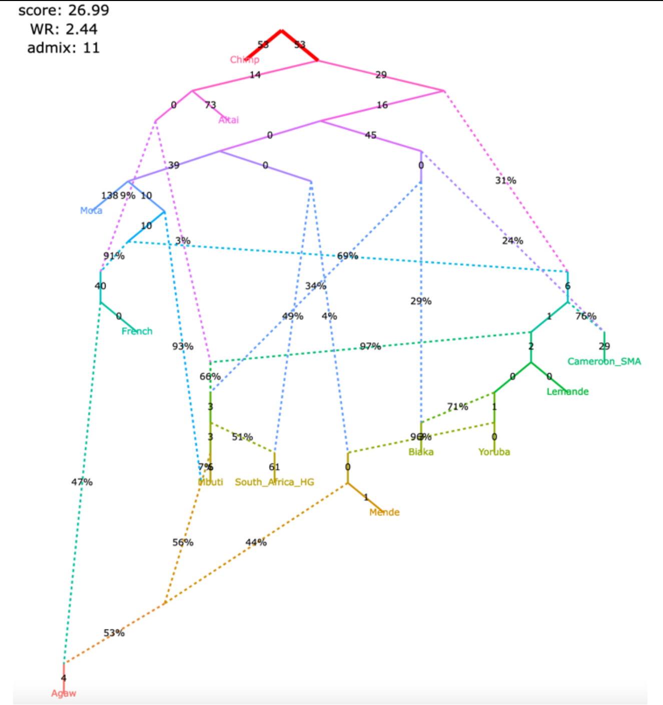

| | |
|---|---|
| **support for claim 1** partial<br>**support for claim 2** not supported<br>**support for claim 3** partial<br>**support for claim 4** partial | **support for claim 1** partial<br>**support for claim 2** not supported<br>**support for claim 3** partial<br>**support for claim 4** partial |

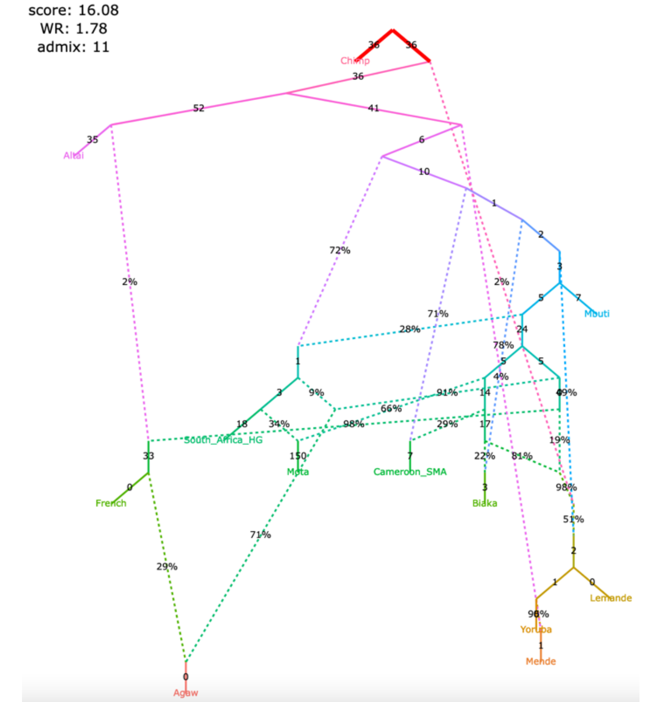

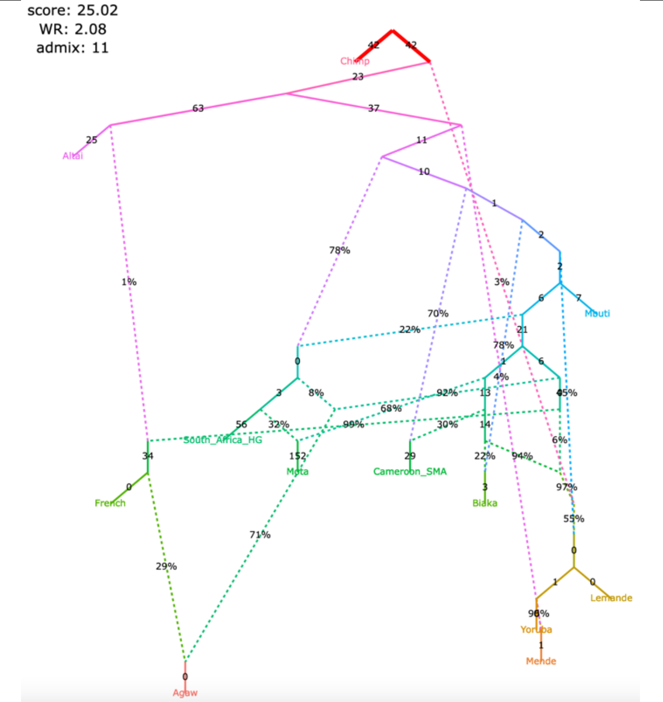

Figure S10

#### **g**, selected alternative models fitting nominally better than the published one

<table>
<tr>
<td>
support for claim 1 supported<br>
support for claim 2 partial<br>
support for claim 3 not supported<br>
support for claim 4 partial
</td>
<td>
support for claim 1 supported<br>
support for claim 2 partial<br>
support for claim 3 not supported<br>
support for claim 4 partial
</td>
</tr>
<tr>
<td>The original set of SNPs and individuals and the corrected algorithm for calculating $f_3$-statistics (211,738 variable site with no missing data at the group level)</td>
<td>The original set of SNPs and individuals and the original algorithm for calculating $f_3$-statistics (363,131 variable site with no missing data at the group level)</td>
</tr>
<tr>
<td>
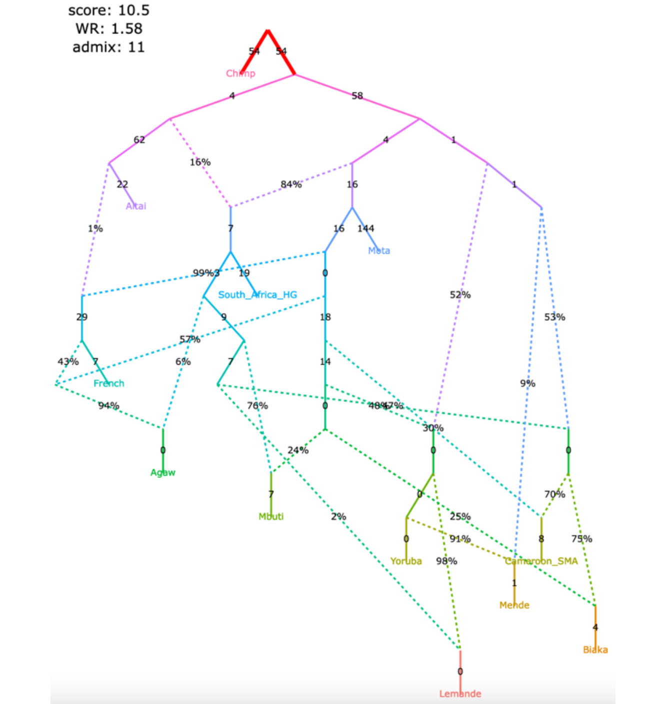

</td>
<td>
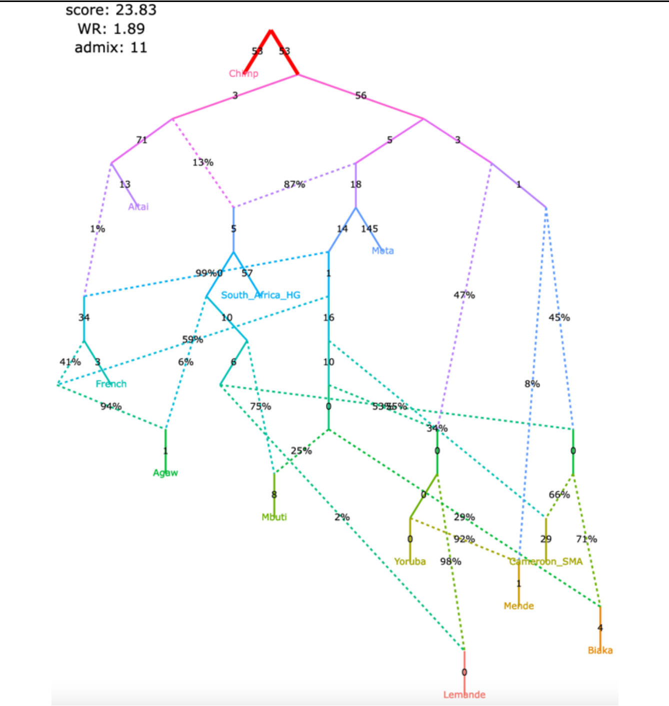

</td>
</tr>
<tr>
<td>
support for claim 1 partial<br>
support for claim 2 partial<br>
support for claim 3 supported<br>
support for claim 4 not supported
</td>
<td>
support for claim 1 partial<br>
support for claim 2 partial<br>
support for claim 3 supported<br>
support for claim 4 not supported
</td>
</tr>
<tr>
<td>
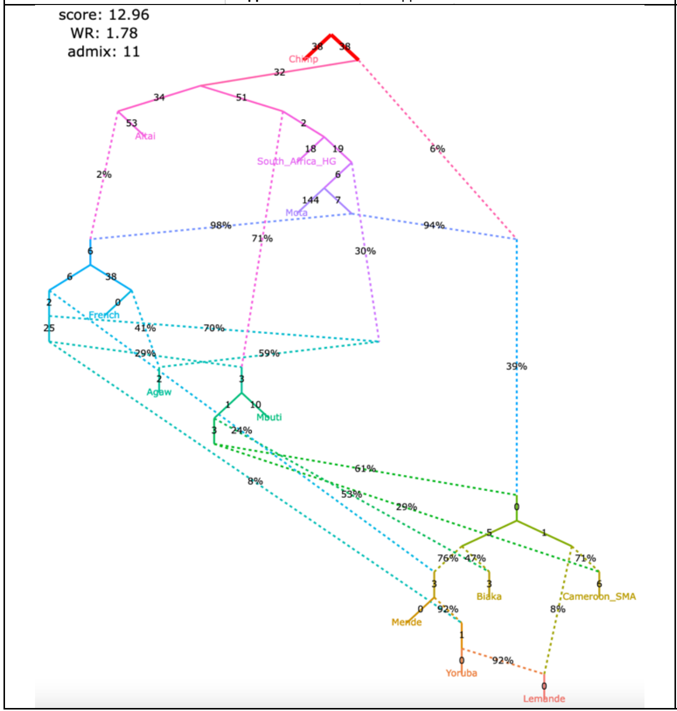

</td>
<td>
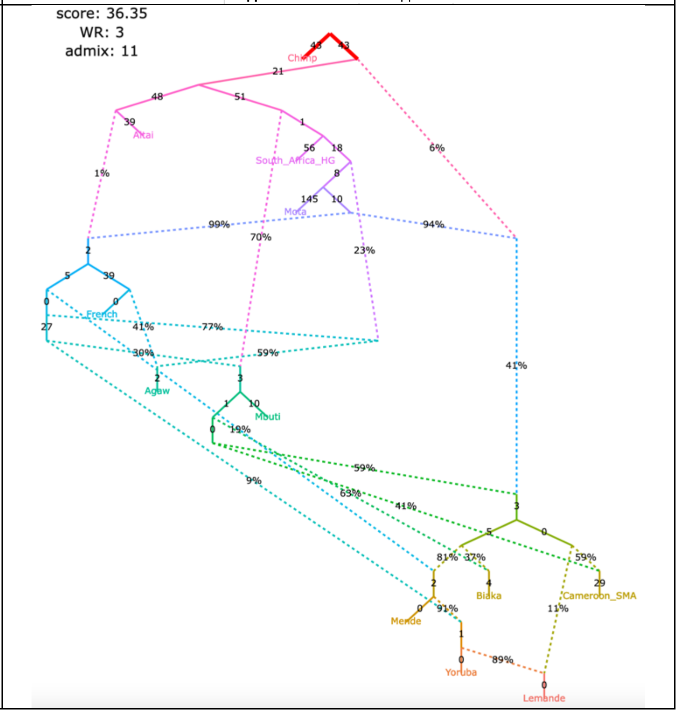

</td>
</tr>
</table>

Figure S10

#### **h**, selected alternative models fitting nominally better than the published one

| | |
|---|---|
| **support for claim 1** partial<br>**support for claim 2** partial<br>**support for claim 3** supported<br>**support for claim 4** not supported | **support for claim 1** partial<br>**support for claim 2** partial<br>**support for claim 3** supported<br>**support for claim 4** not supported |
| The original set of SNPs and individuals and the corrected algorithm for calculating $f_3$-statistics (211,738 variable site with no missing data at the group level) | The original set of SNPs and individuals and the original algorithm for calculating $f_3$-statistics (363,131 variable site with no missing data at the group level) |
| 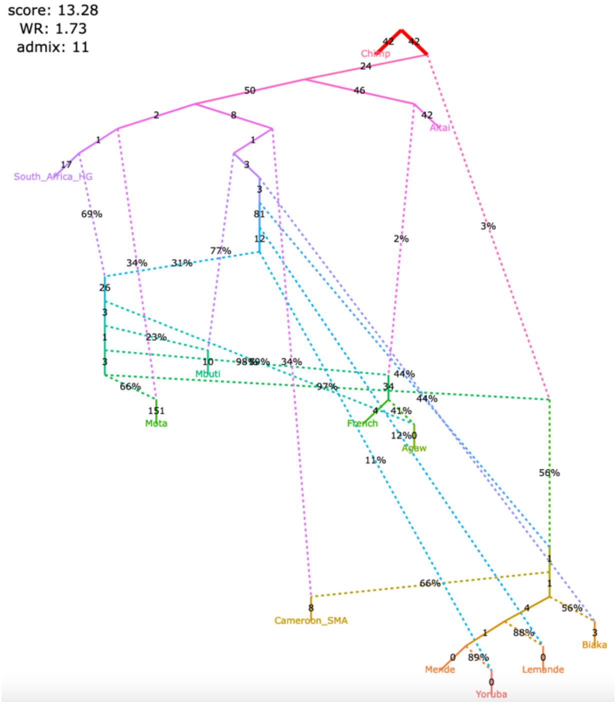 | 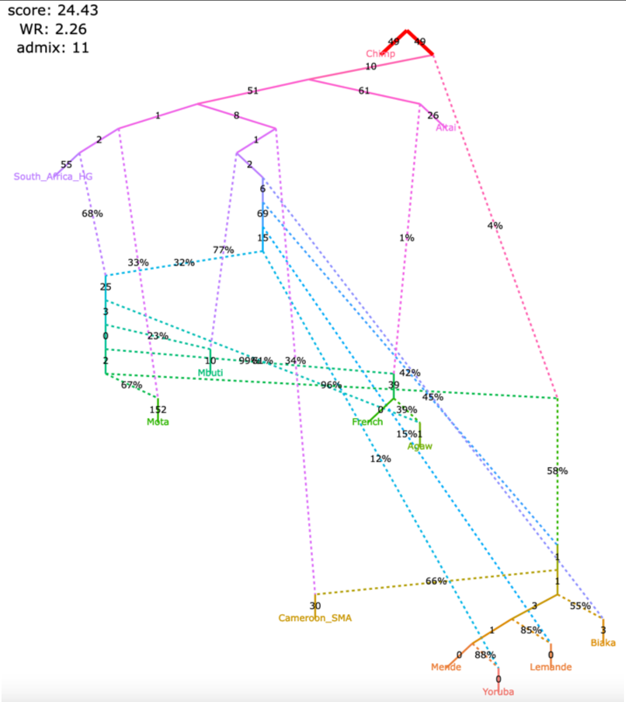 |
| **support for claim 1** partial<br>**support for claim 2** partial<br>**support for claim 3** supported<br>**support for claim 4** not supported | **support for claim 1** partial<br>**support for claim 2** partial<br>**support for claim 3** supported<br>**support for claim 4** not supported |
| 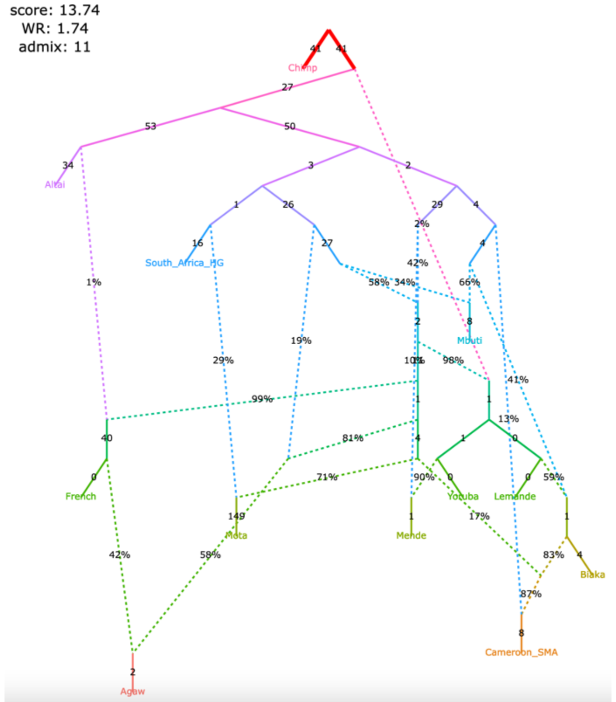 | 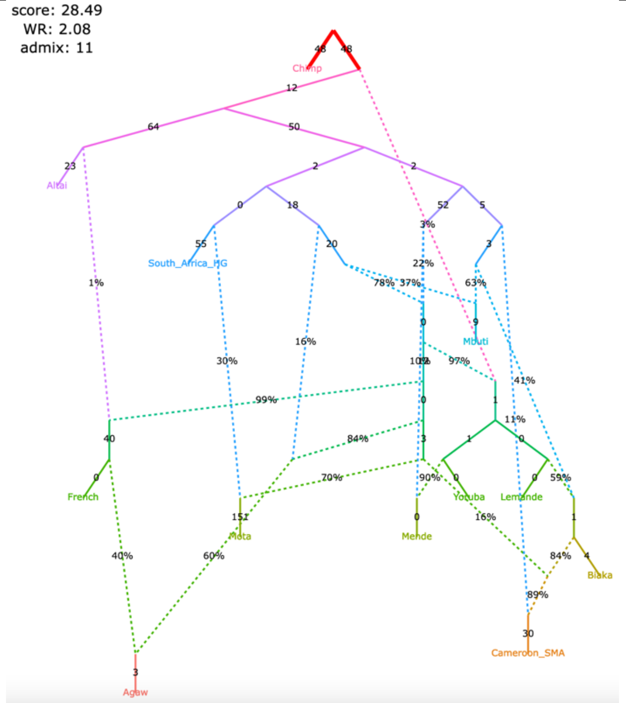 |

Figure S10

#### **i**, selected alternative models fitting nominally better than the published one

| **support for claim 1** | not supported |
|---|---|
| **support for claim 2** | partial |
| **support for claim 3** | partial |
| **support for claim 4** | supported |

The original set of SNPs and individuals and the corrected algorithm for calculating $f_3$-statistics (211,738 variable site with no missing data at the group level)

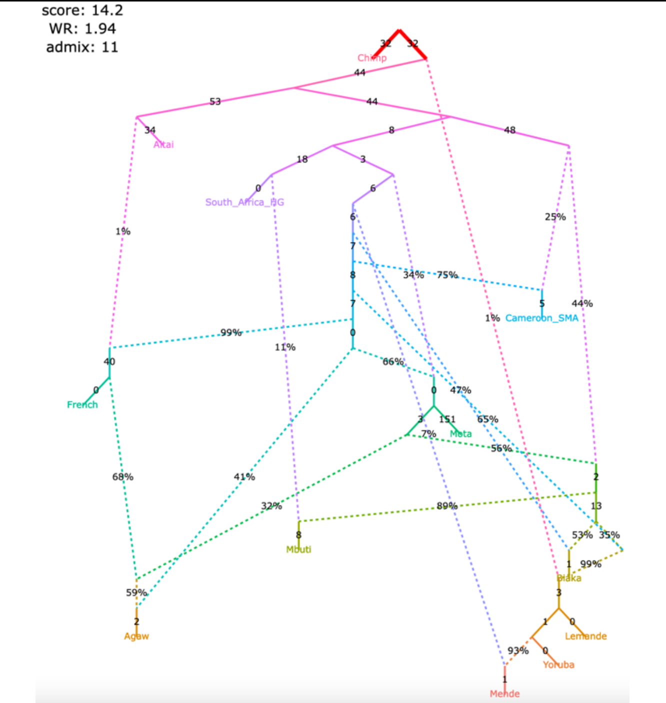

| **support for claim 1** | not supported |
|---|---|
| **support for claim 2** | partial |
| **support for claim 3** | partial |
| **support for claim 4** | supported |

The original set of SNPs and individuals and the original algorithm for calculating $f_3$-statistics (363,131 variable site with no missing data at the group level)

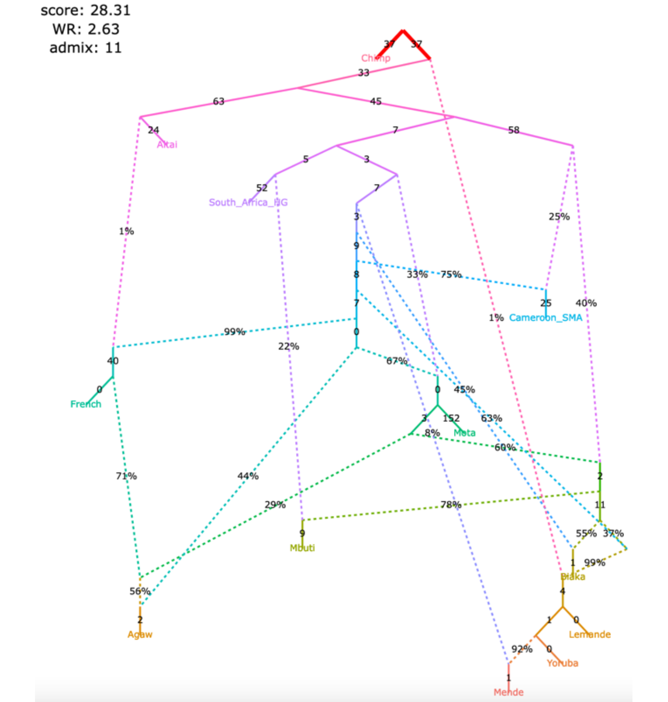

| **support for claim 1** | supported |
|---|---|
| **support for claim 2** | partial |
| **support for claim 3** | not supported |
| **support for claim 4** | partial |

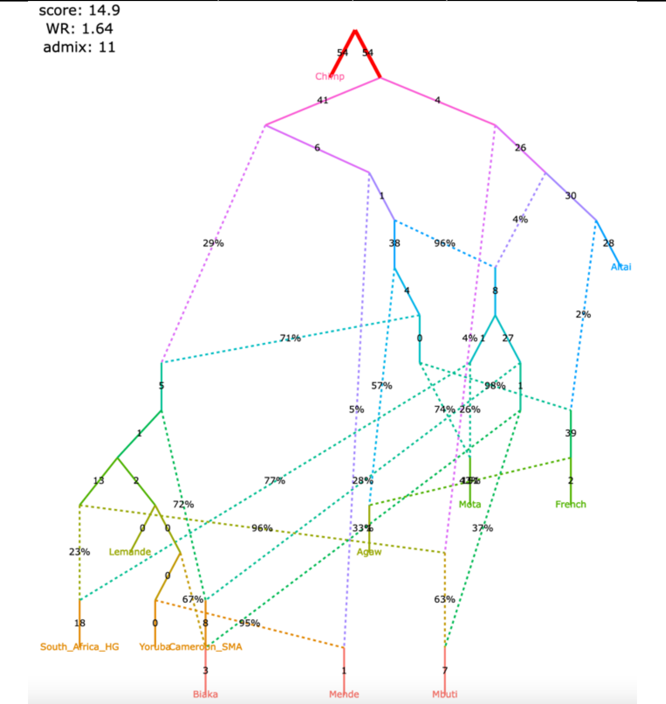

| **support for claim 1** | supported |
|---|---|
| **support for claim 2** | partial |
| **support for claim 3** | not supported |
| **support for claim 4** | partial |

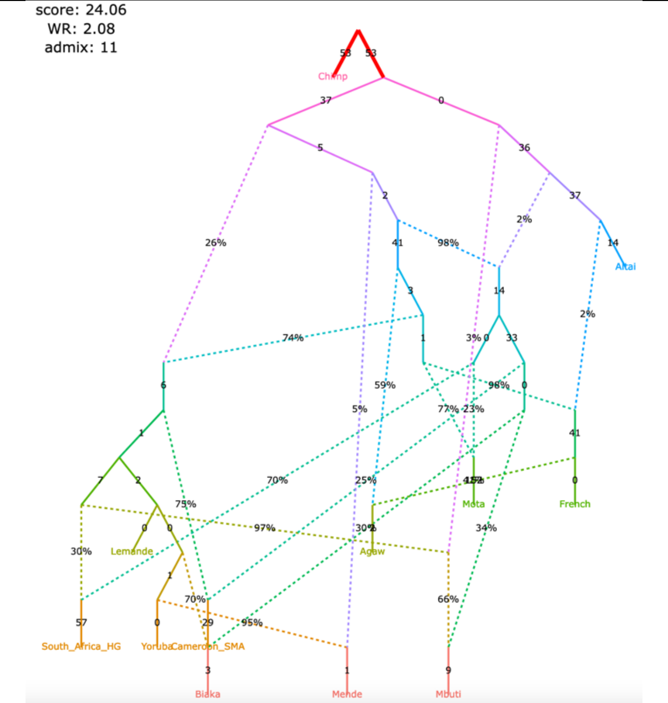

Figure S10

#### **j**, selected alternative models fitting nominally better than the published one

| | | | |
|---|---|---|---|
| **support for claim 1** | supported | **support for claim 1** | supported |
| **support for claim 2** | partial | **support for claim 2** | partial |
| **support for claim 3** | partial | **support for claim 3** | partial |
| **support for claim 4** | not supported | **support for claim 4** | not supported |

| The original set of SNPs and individuals and the corrected algorithm for calculating $f_3$-statistics (211,738 variable site with no missing data at the group level) | The original set of SNPs and individuals and the original algorithm for calculating $f_3$-statistics (363,131 variable site with no missing data at the group level) |
|---|---|

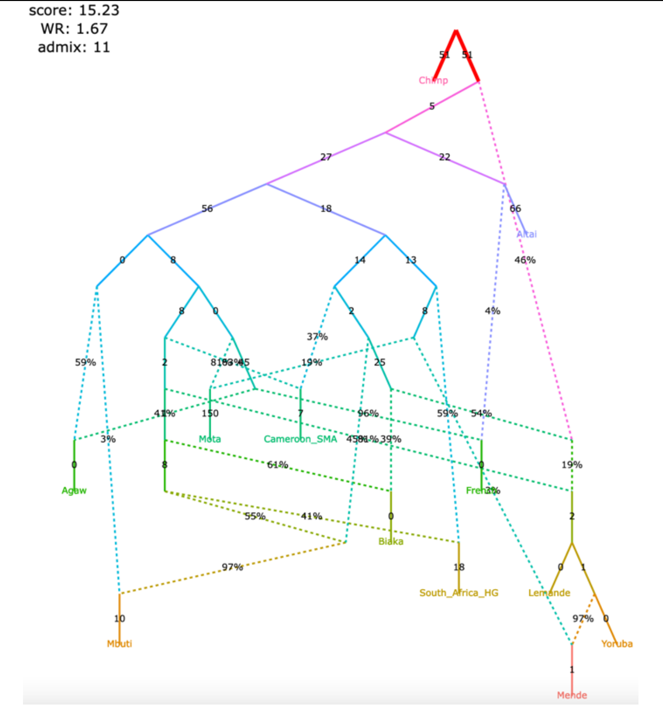

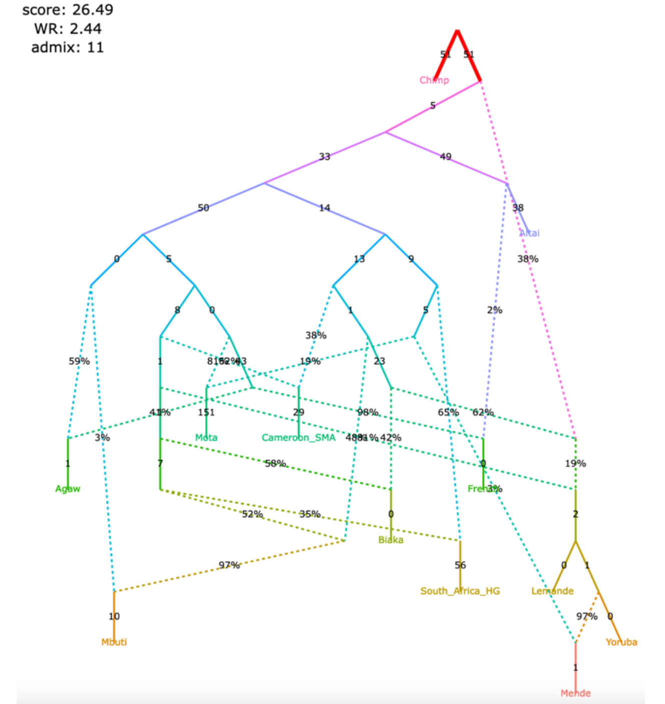

| | | | |
|---|---|---|---|
| **support for claim 1** | supported | **support for claim 1** | supported |
| **support for claim 2** | not supported | **support for claim 2** | not supported |
| **support for claim 3** | partial | **support for claim 3** | partial |
| **support for claim 4** | partial | **support for claim 4** | partial |

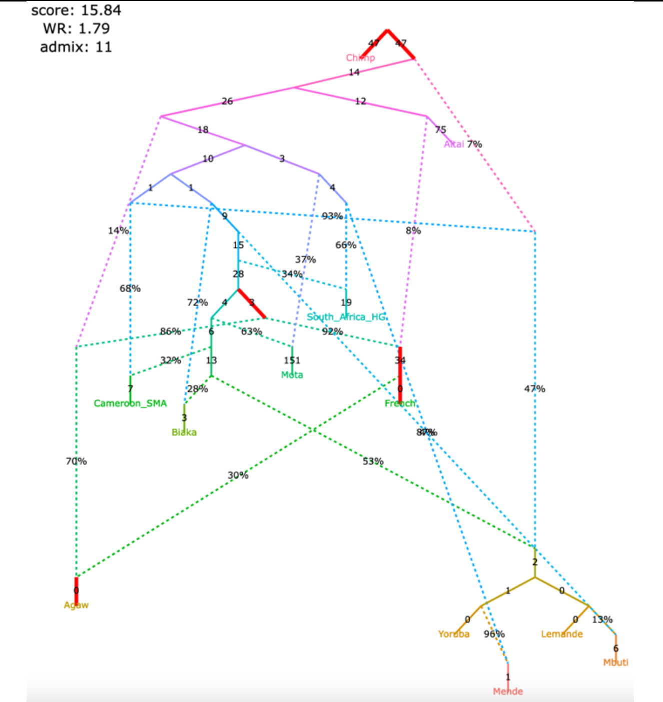

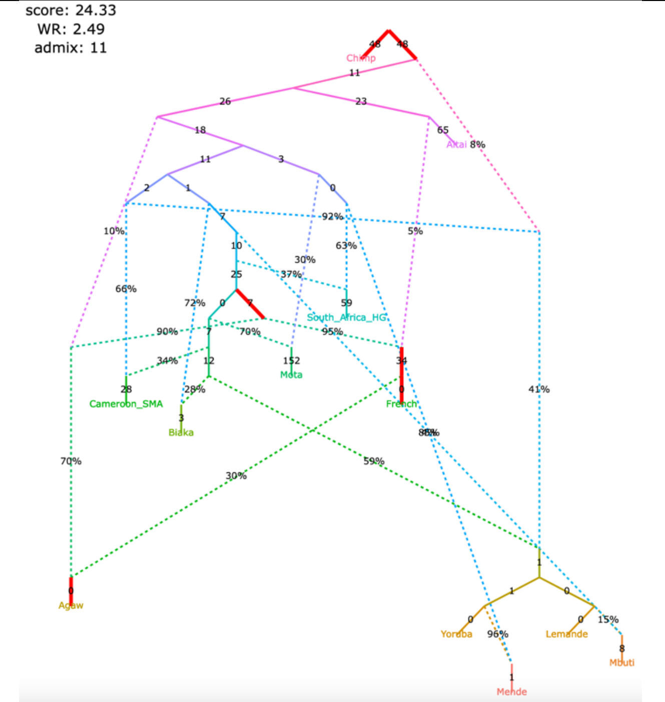

Figure S10

#### **k**, selected alternative models fitting nominally better than the published one

The original set of SNPs and individuals and the corrected algorithm for calculating $f_3$-statistics (211,738 variable site with no missing data at the group level)

| | |
|---|---|
| **support for claim 1** | supported |
| **support for claim 2** | supported |
| **support for claim 3** | supported |
| **support for claim 4** | not supported |

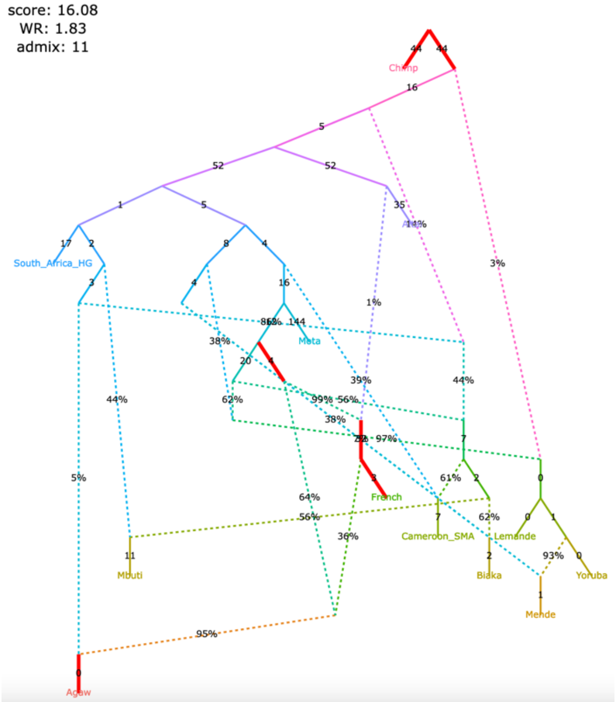

The original set of SNPs and individuals and the original algorithm for calculating $f_3$-statistics (363,131 variable site with no missing data at the group level)

| | |
|---|---|
| **support for claim 1** | supported |
| **support for claim 2** | supported |
| **support for claim 3** | supported |
| **support for claim 4** | not supported |

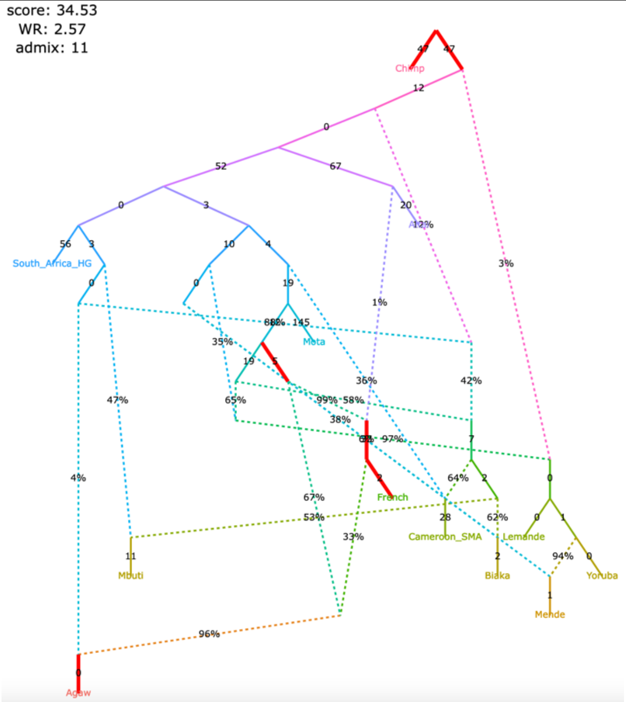

| | |
|---|---|
| **support for claim 1** | partial |
| **support for claim 2** | partial |
| **support for claim 3** | partial |
| **support for claim 4** | partial |

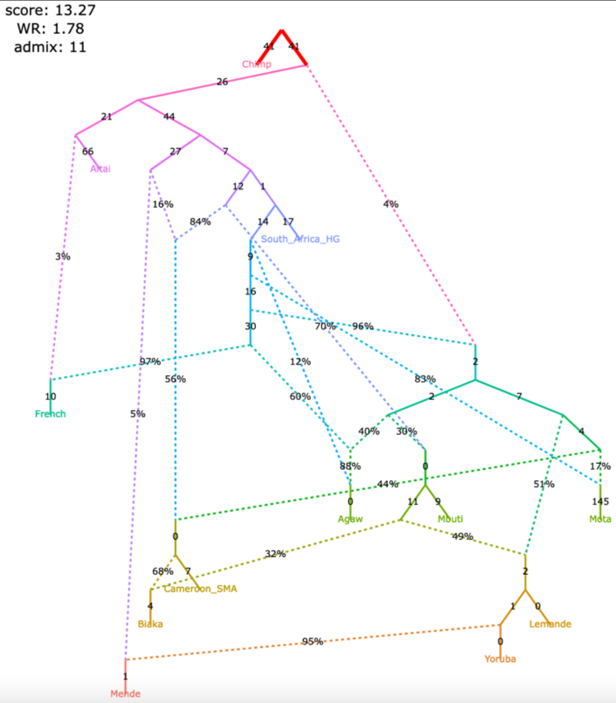

| | |
|---|---|
| **support for claim 1** | partial |
| **support for claim 2** | partial |
| **support for claim 3** | partial |
| **support for claim 4** | partial |

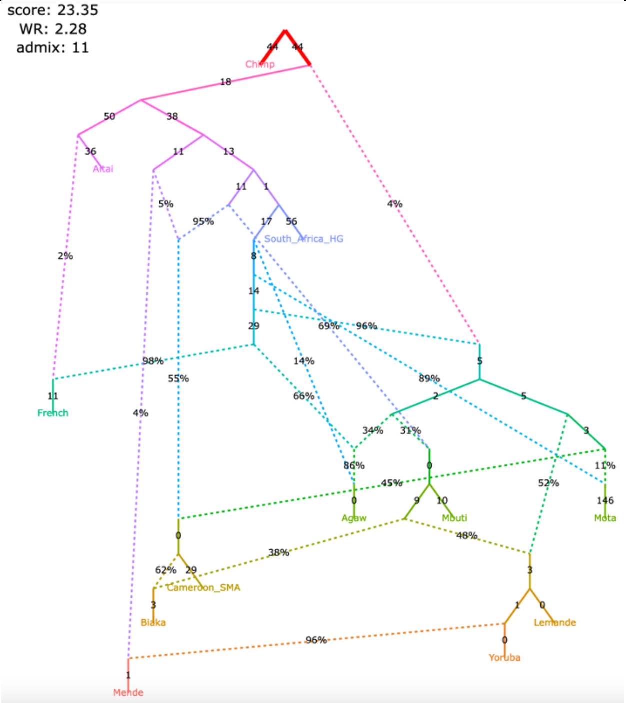

Figure S10

#### **I**, selected alternative models fitting nominally better than the published one

| | | | |
|---|---|---|---|
| **support for claim 1** | partial | **support for claim 1** | partial |
| **support for claim 2** | partial | **support for claim 2** | partial |
| **support for claim 3** | partial | **support for claim 3** | partial |
| **support for claim 4** | partial | **support for claim 4** | partial |

| The original set of SNPs and individuals and the corrected algorithm for calculating $f_3$-statistics (211,738 variable site with no missing data at the group level) | The original set of SNPs and individuals and the original algorithm for calculating $f_3$-statistics (363,131 variable site with no missing data at the group level) |
|---|---|

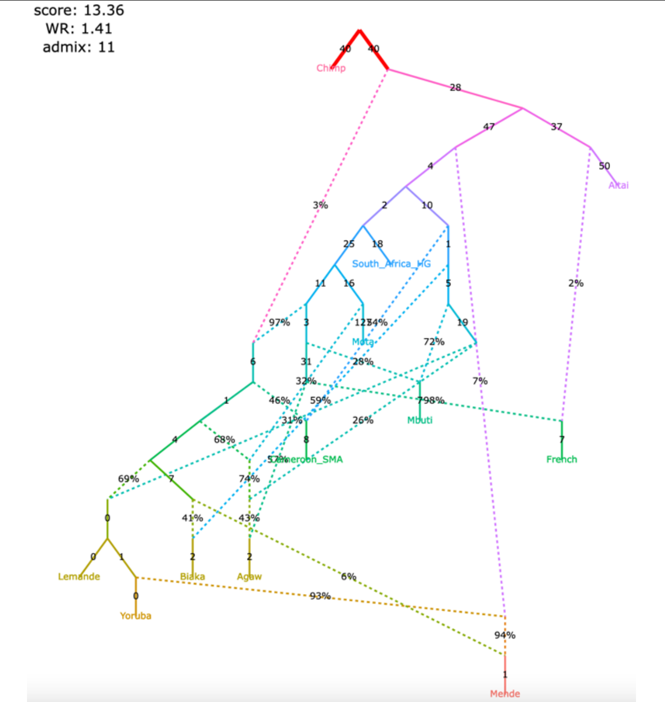

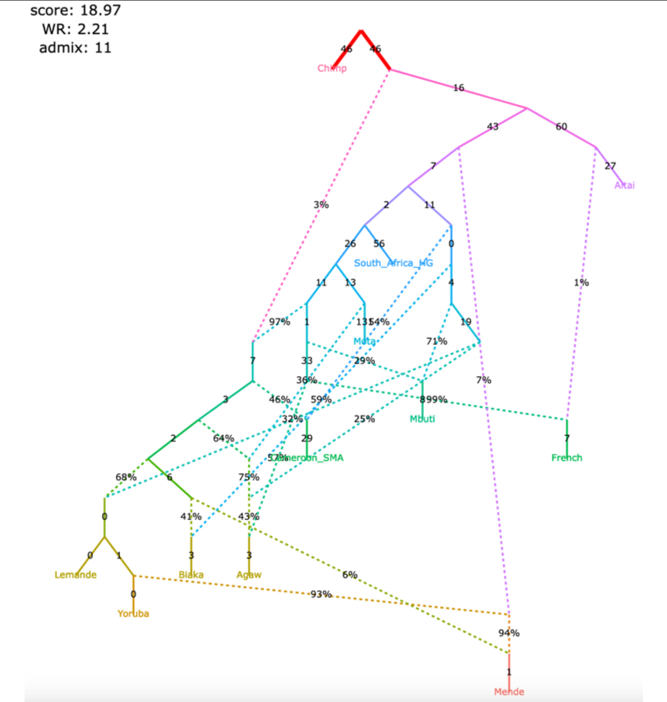

| | | | |
|---|---|---|---|
| **support for claim 1** | partial | **support for claim 1** | partial |
| **support for claim 2** | partial | **support for claim 2** | partial |
| **support for claim 3** | partial | **support for claim 3** | partial |
| **support for claim 4** | partial | **support for claim 4** | partial |

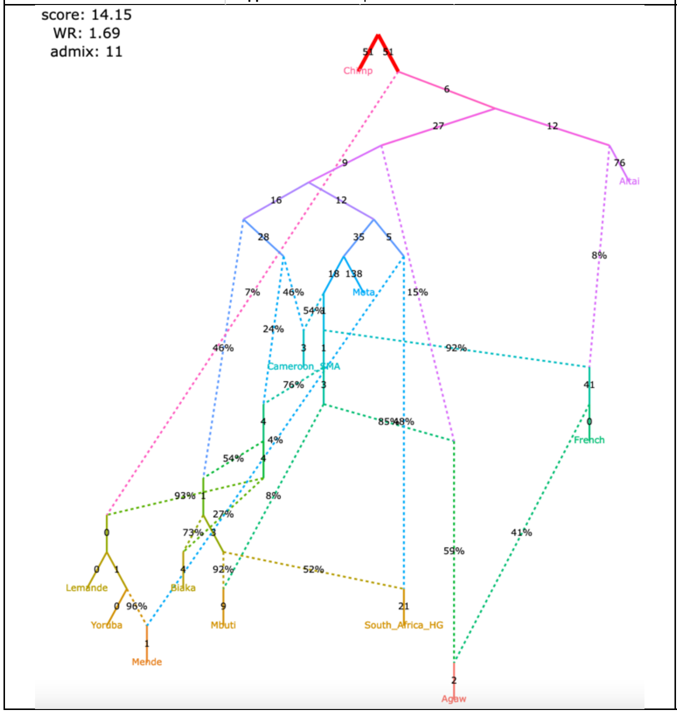

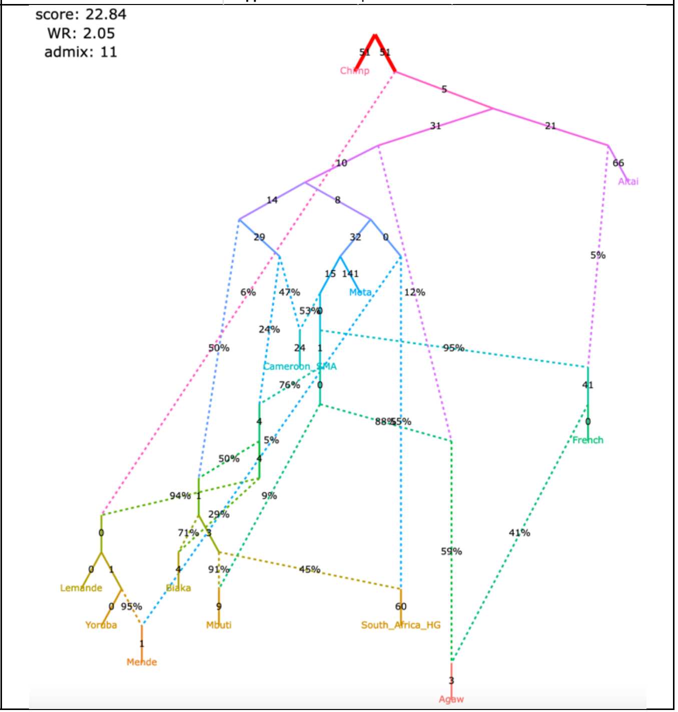

Figure S10

**m**, selected alternative models fitting nominally better than the published one

| | |
|---|---|
| **support for claim 1** | partial |
| **support for claim 2** | partial |
| **support for claim 3** | partial |
| **support for claim 4** | partial |

The original set of SNPs and individuals and the corrected algorithm for calculating $f_3$-statistics (211,738 variable site with no missing data at the group level)

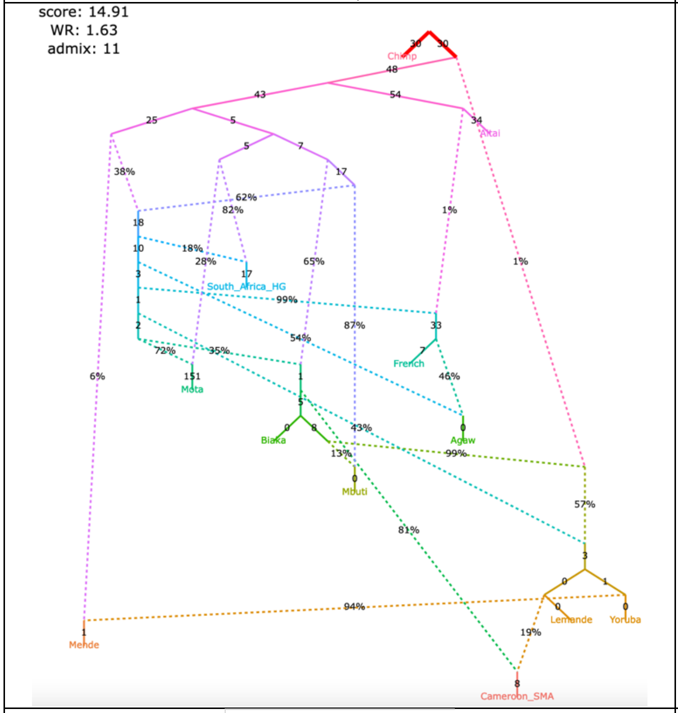

| | |
|---|---|
| **support for claim 1** | partial |
| **support for claim 2** | partial |
| **support for claim 3** | partial |
| **support for claim 4** | partial |

The original set of SNPs and individuals and the original algorithm for calculating $f_3$-statistics (363,131 variable site with no missing data at the group level)

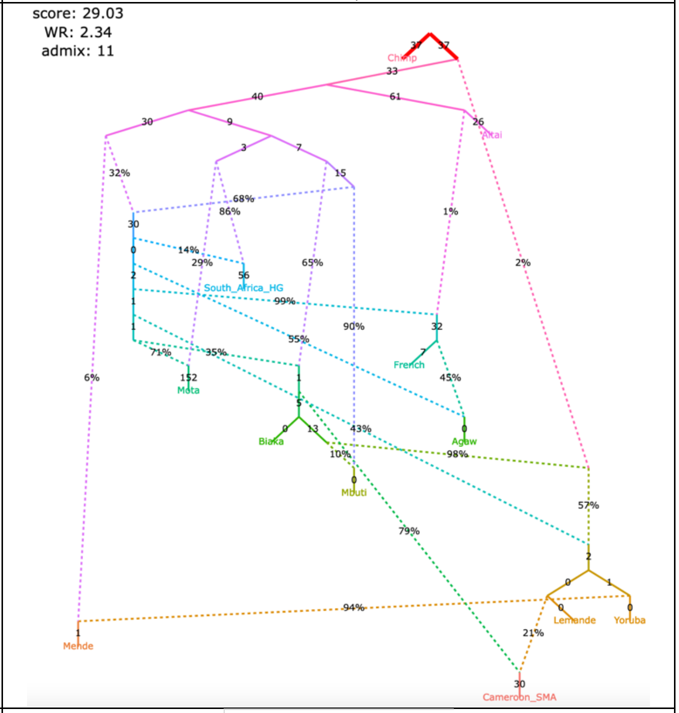

| | |
|---|---|
| **support for claim 1** | partial |
| **support for claim 2** | partial |
| **support for claim 3** | partial |
| **support for claim 4** | partial |

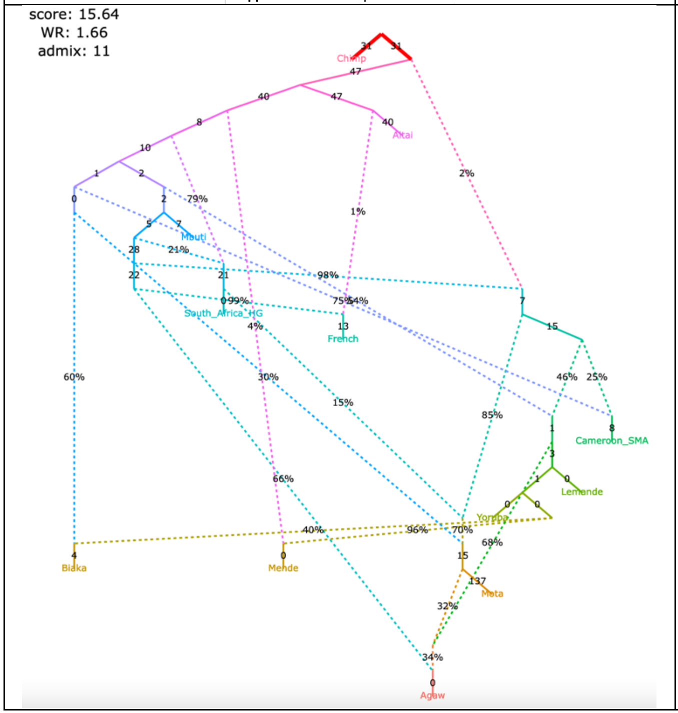

| | |
|---|---|
| **support for claim 1** | partial |
| **support for claim 2** | partial |
| **support for claim 3** | partial |
| **support for claim 4** | partial |

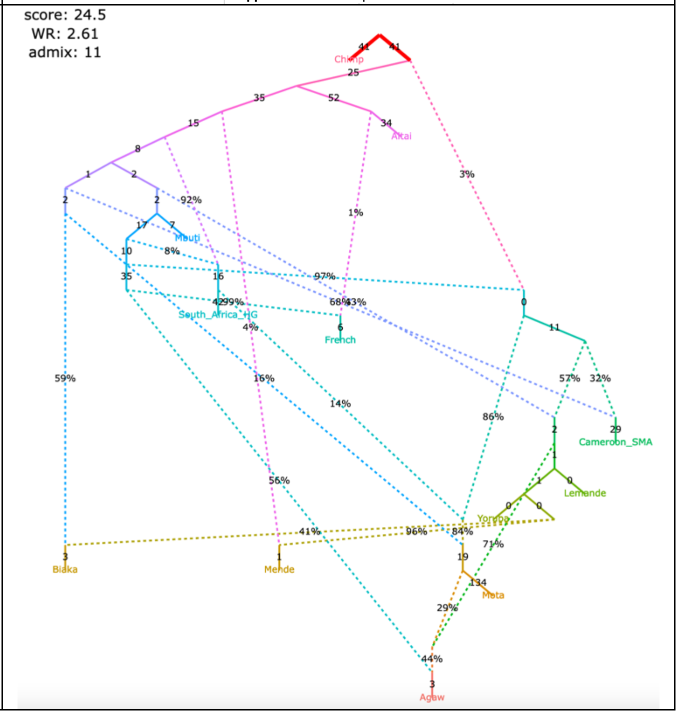

Figure S10

**n**, selected alternative models fitting nominally better than the published one

| | |
|---|---|
| **support for claim 1** partial<br>**support for claim 2** partial<br>**support for claim 3** supported<br>**support for claim 4** partial | **support for claim 1** partial<br>**support for claim 2** partial<br>**support for claim 3** supported<br>**support for claim 4** partial |
| The original set of SNPs and individuals and the corrected algorithm for calculating $f_3$-statistics (211,738 variable site with no missing data at the group level) | The original set of SNPs and individuals and the original algorithm for calculating $f_3$-statistics (363,131 variable site with no missing data at the group level) |
| 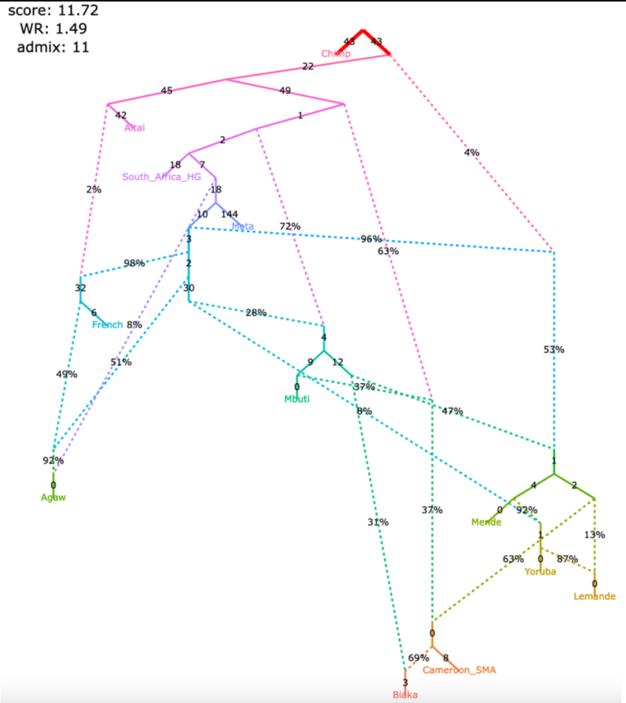 | 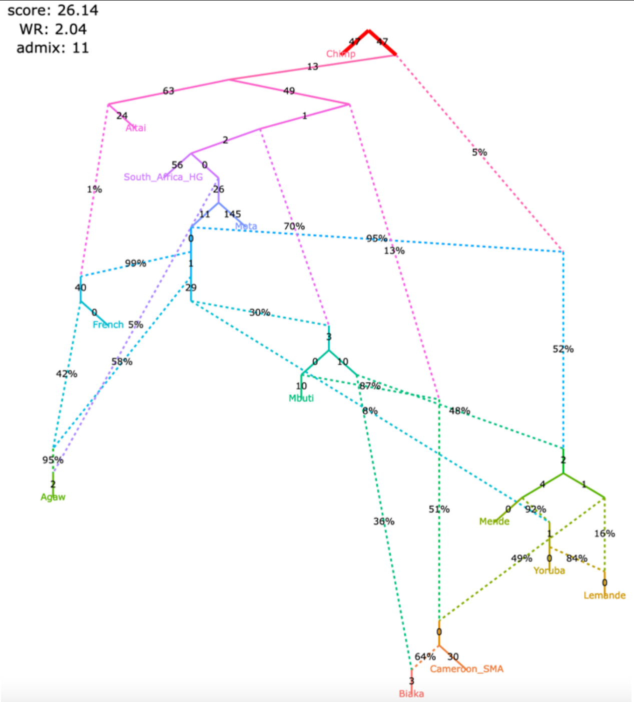 |
| **support for claim 1** supported<br>**support for claim 2** partial<br>**support for claim 3** partial<br>**support for claim 4** partial | **support for claim 1** supported<br>**support for claim 2** partial<br>**support for claim 3** partial<br>**support for claim 4** partial |
| 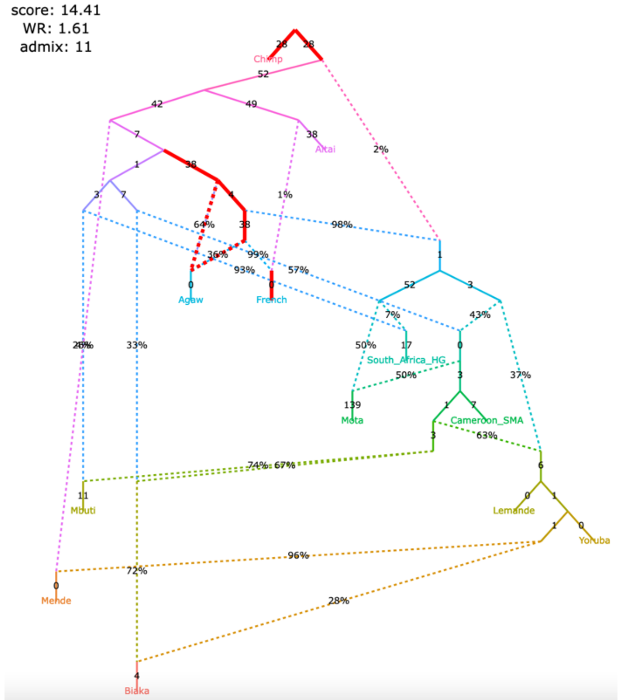 | 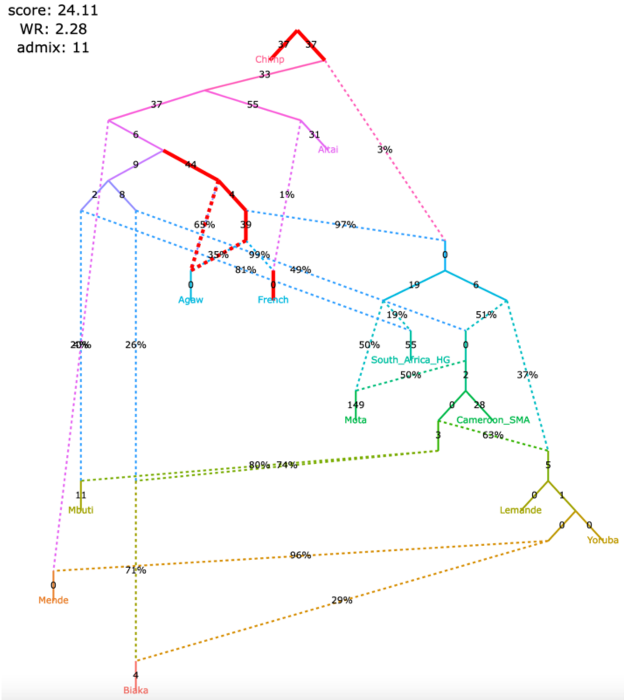 |

Figure S10

**o**, selected alternative models fitting nominally better than the published one

| | |
|---|---|
| **support for claim 1** partial<br>**support for claim 2** partial<br>**support for claim 3** supported<br>**support for claim 4** partial | **support for claim 1** partial<br>**support for claim 2** partial<br>**support for claim 3** supported<br>**support for claim 4** partial |
| The original set of SNPs and individuals and the corrected algorithm for calculating $f_3$-statistics (211,738 variable site with no missing data at the group level) | The original set of SNPs and individuals and the original algorithm for calculating $f_3$-statistics (363,131 variable site with no missing data at the group level) |

score: 16.13
WR: 2.05
admix: 11

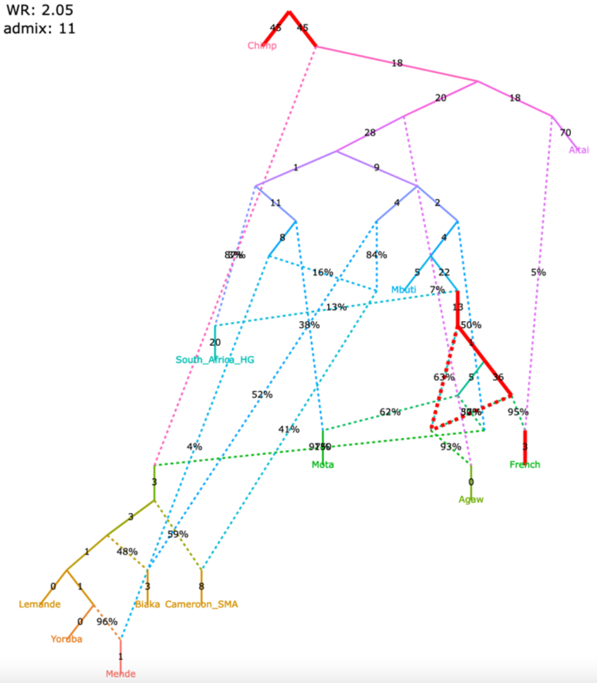

score: 31.26
WR: 2.72
admix: 11

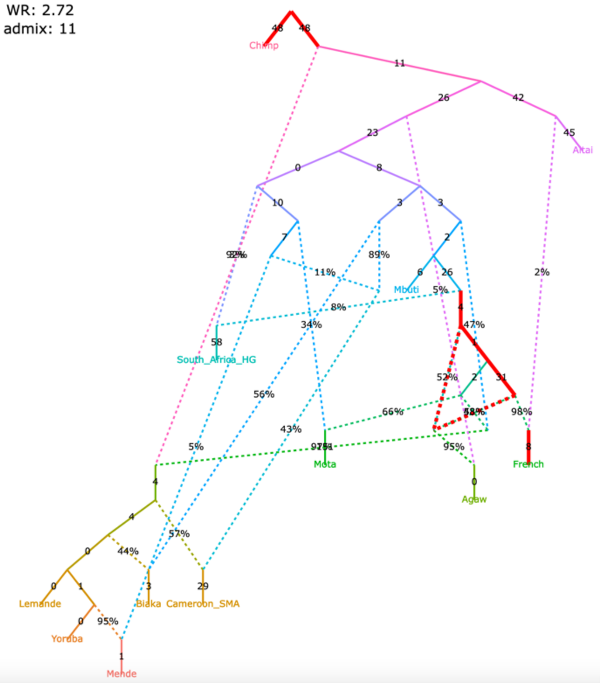

| | |
|---|---|
| **support for claim 1** supported<br>**support for claim 2** partial<br>**support for claim 3** supported<br>**support for claim 4** partial | **support for claim 1** supported<br>**support for claim 2** partial<br>**support for claim 3** supported<br>**support for claim 4** partial |

score: 15.03
WR: 1.77
admix: 11

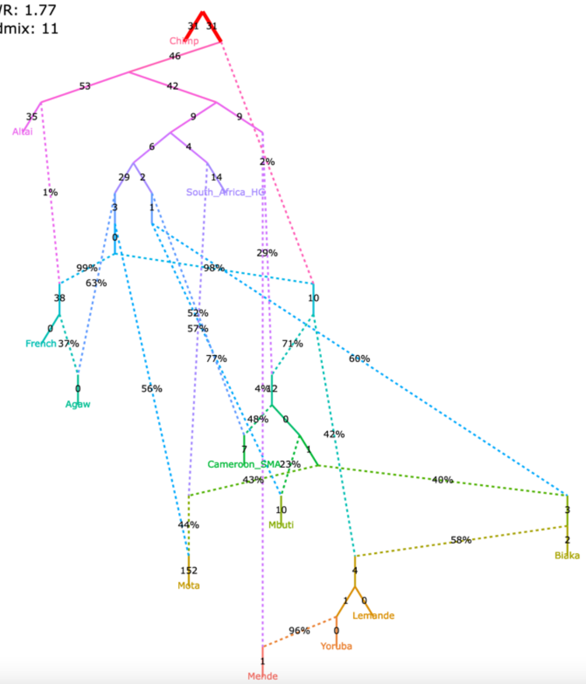

score: 29.02
WR: 2.19
admix: 11

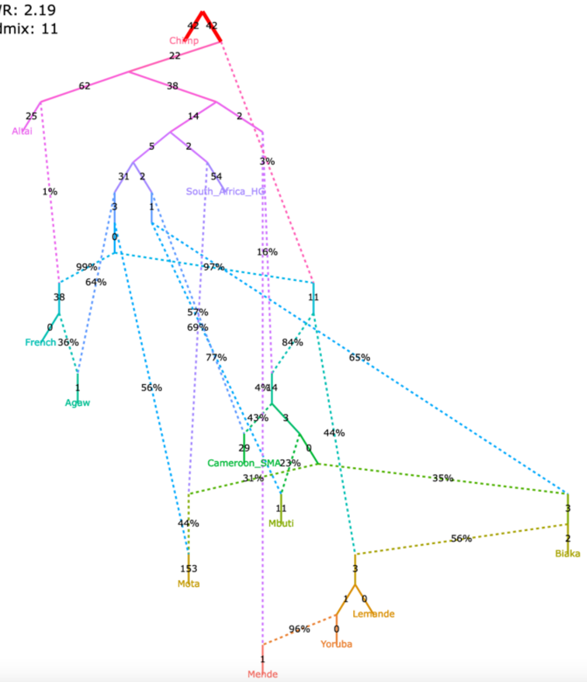

Figure S10

#### **p**, selected alternative models fitting nominally better than the published one

| | |
|---|---|
| **support for claim 1** | supported |
| **support for claim 2** | partial |
| **support for claim 3** | supported |
| **support for claim 4** | partial |

The original set of SNPs and individuals and the corrected algorithm for calculating *f*$_3$-statistics (211,738 variable site with no missing data at the group level)

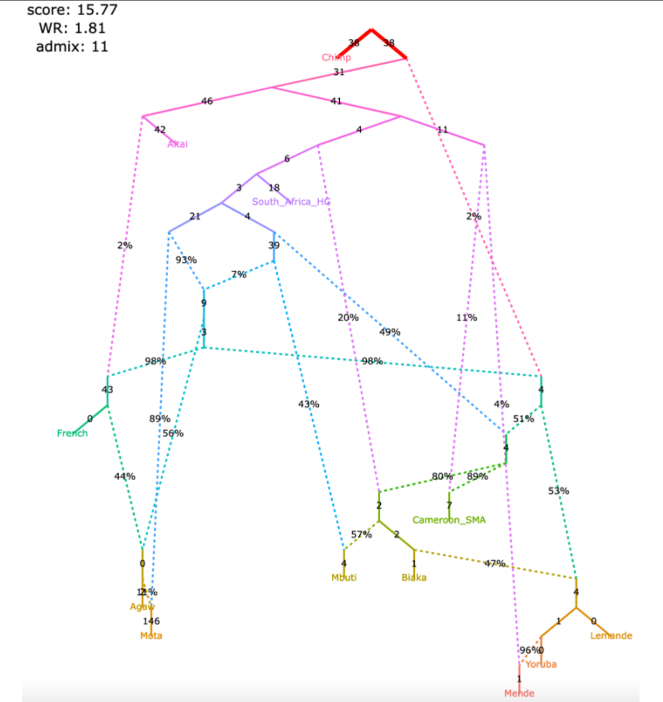

| | |
|---|---|
| **support for claim 1** | supported |
| **support for claim 2** | partial |
| **support for claim 3** | supported |
| **support for claim 4** | partial |

The original set of SNPs and individuals and the original algorithm for calculating *f*$_3$-statistics (363,131 variable site with no missing data at the group level)

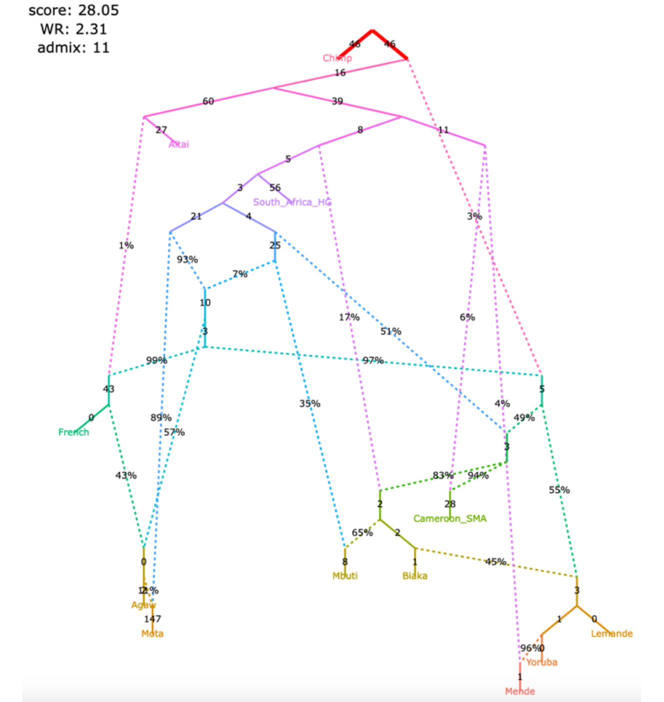

| | |
|---|---|
| **support for claim 1** | supported |
| **support for claim 2** | partial |
| **support for claim 3** | supported |
| **support for claim 4** | partial |

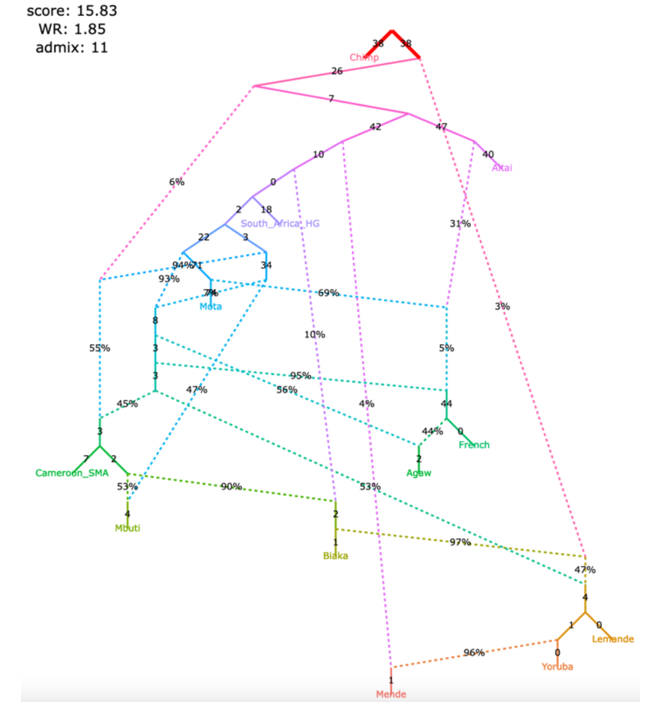

| | |
|---|---|
| **support for claim 1** | supported |
| **support for claim 2** | partial |
| **support for claim 3** | supported |
| **support for claim 4** | partial |

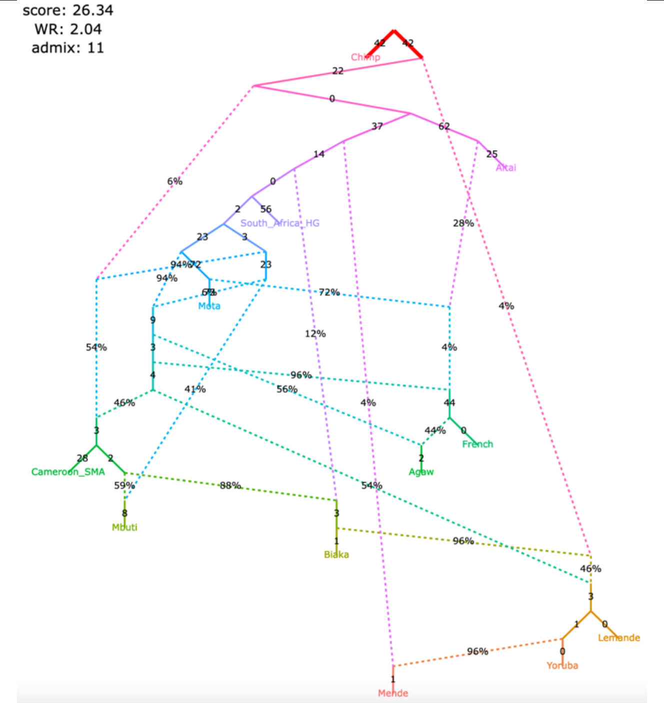

Figure S10

#### **q**, selected alternative models fitting nominally better than the published one

| support for claim 1 | supported | support for claim 1 | supported |
|---|---|---|---|
| support for claim 2 | partial | support for claim 2 | partial |
| support for claim 3 | supported | support for claim 3 | supported |
| support for claim 4 | supported | support for claim 4 | supported |

The original set of SNPs and individuals and the corrected algorithm for calculating $f_3$-statistics (211,738 variable site with no missing data at the group level)

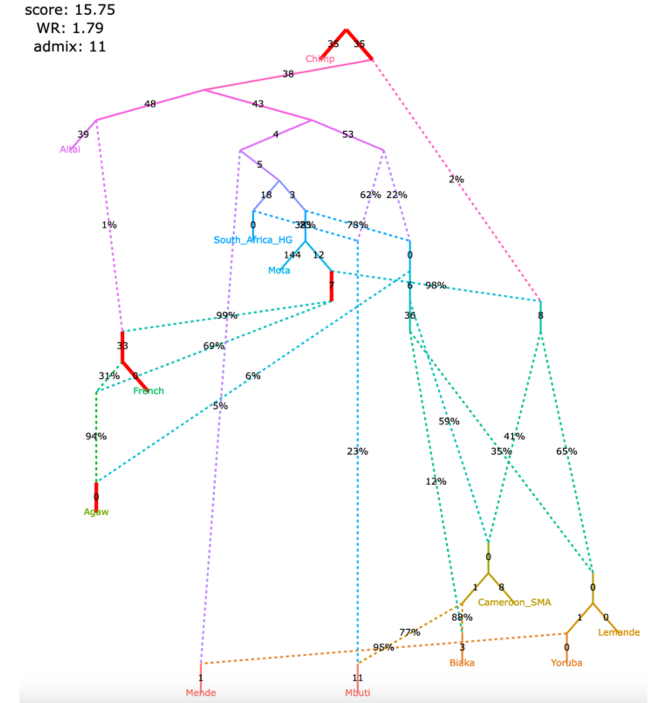

The original set of SNPs and individuals and the original algorithm for calculating $f_3$-statistics (363,131 variable site with no missing data at the group level)

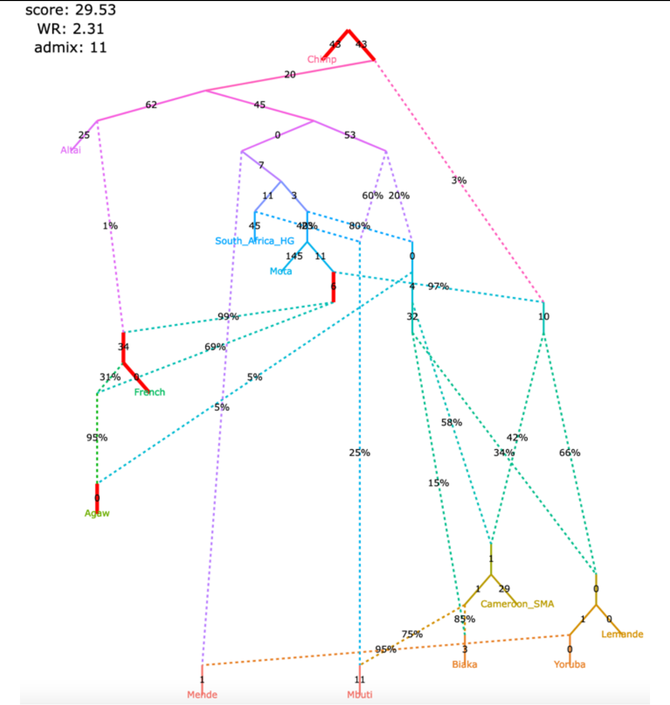

| support for claim 1 | supported | support for claim 1 | supported |
|---|---|---|---|
| support for claim 2 | supported | support for claim 2 | supported |
| support for claim 3 | supported | support for claim 3 | supported |
| support for claim 4 | supported | support for claim 4 | supported |

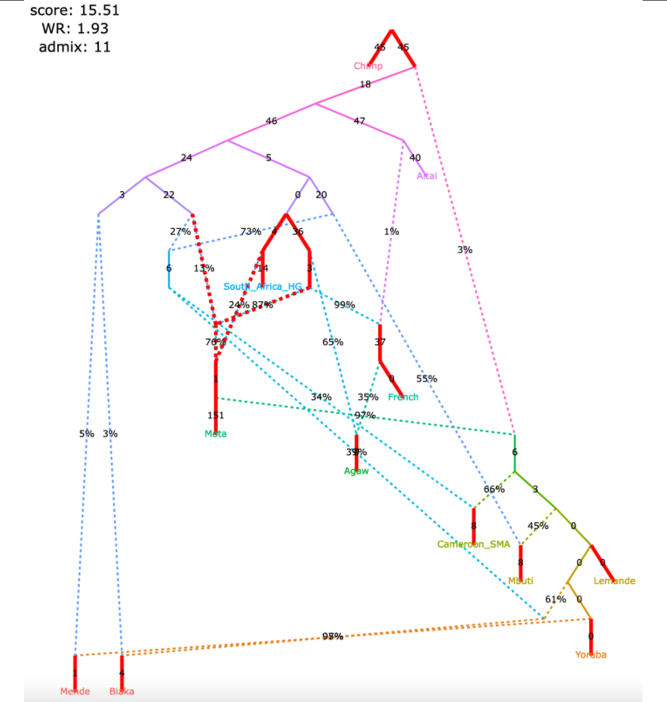

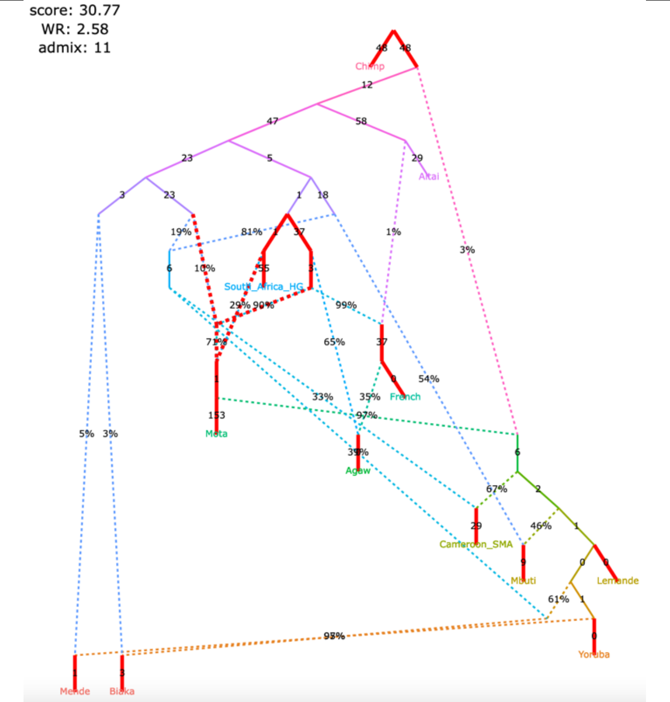

Figure S11

**Figure S11.** Published admixture graph from Wang *et al*. (2021) and alternative graphs found with *findGraphs* (12 populations, 8 admixture events) using the updated algorithm for calculating *f*-statistics. The graphs were also re-fitted using the original algorithm for calculating *f*-statistics.

**a**, published model

| | |
|---|---|
| no. of East Asian groups with Onge-related ancestry | 6 |
| no. of East Asian groups with European-related ancestry | 1 |
| direction of gene flow China_Island_EN > Taiwan_IA | forward |
| Mongolia_East is basal with respect to China_Upper_YR and Nepal_Chokhopani | yes |
| Taiwan_IA is basal with respect to Mongolia_East and Nepal_Chokhopani | yes |

The original set of SNPs and individuals and the corrected algorithm for calculating $f_3$-statistics (203,753 variable site with no missing data at the group level)

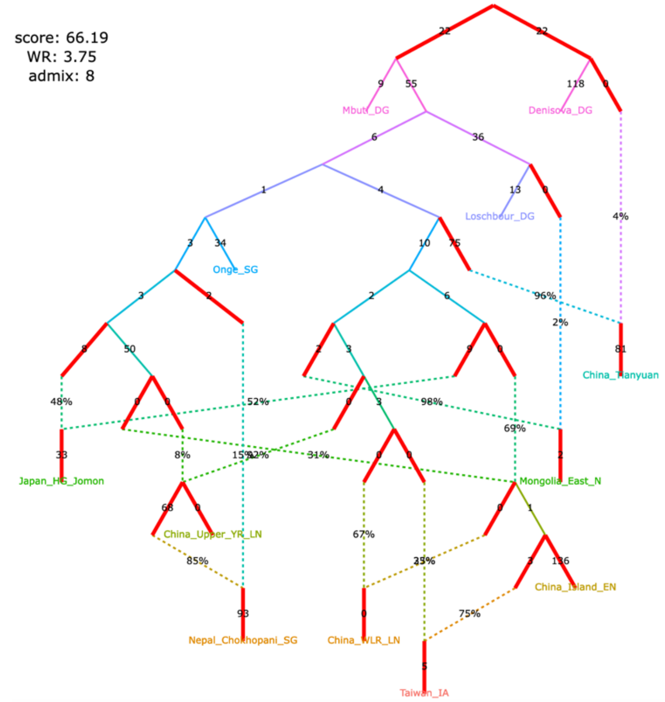

| | |
|---|---|
| no. of East Asian groups with Onge-related ancestry | 6 |
| no. of East Asian groups with European-related ancestry | 1 |
| direction of gene flow China_Island_EN > Taiwan_IA | forward |
| Mongolia_East is basal with respect to China_Upper_YR and Nepal_Chokhopani | yes |
| Taiwan_IA is basal with respect to Mongolia_East and Nepal_Chokhopani | yes |

The original set of SNPs and individuals and the original algorithm for calculating $f_3$-statistics (496,233 variable site with no missing data at the group level)

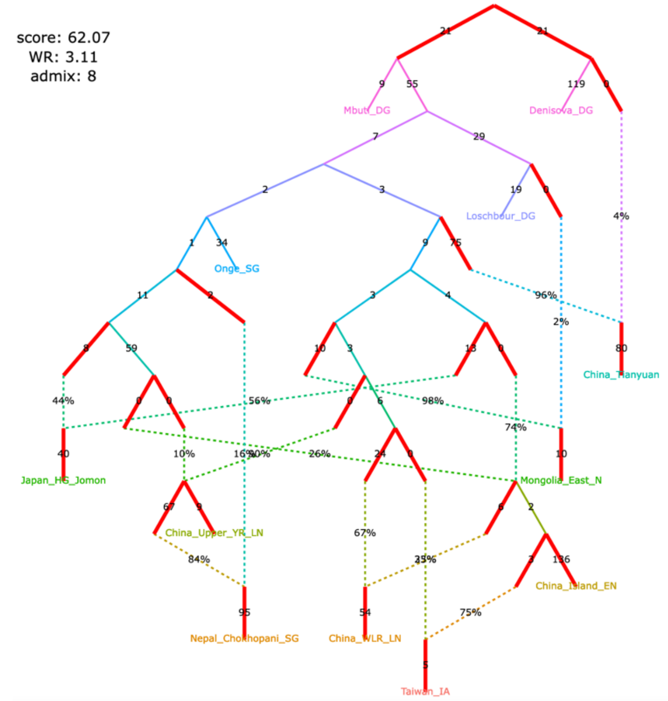

Figure S11

**b**, published model, simplified

| | |
|---|---|
| no. of East Asian groups with Onge-related ancestry | 6 |
| no. of East Asian groups with European-related ancestry | 1 |
| direction of gene flow China_Island_EN > Taiwan_IA | forward |
| Mongolia_East is basal with respect to China_Upper_YR and Nepal_Chokhopani | yes |
| Taiwan_IA is basal with respect to Mongolia_East and Nepal_Chokhopani | yes |

The original set of SNPs and individuals and the corrected algorithm for calculating $f_3$-statistics (203,753 variable site with no missing data at the group level)

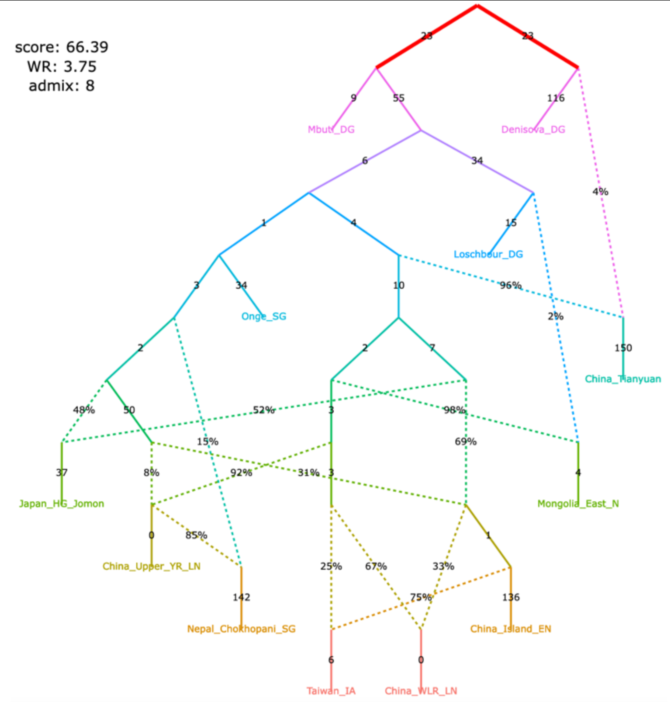

| | |
|---|---|
| no. of East Asian groups with Onge-related ancestry | 6 |
| no. of East Asian groups with European-related ancestry | 1 |
| direction of gene flow China_Island_EN > Taiwan_IA | forward |
| Mongolia_East is basal with respect to China_Upper_YR and Nepal_Chokhopani | yes |
| Taiwan_IA is basal with respect to Mongolia_East and Nepal_Chokhopani | yes |

The original set of SNPs and individuals and the original algorithm for calculating $f_3$-statistics (496,233 variable site with no missing data at the group level)

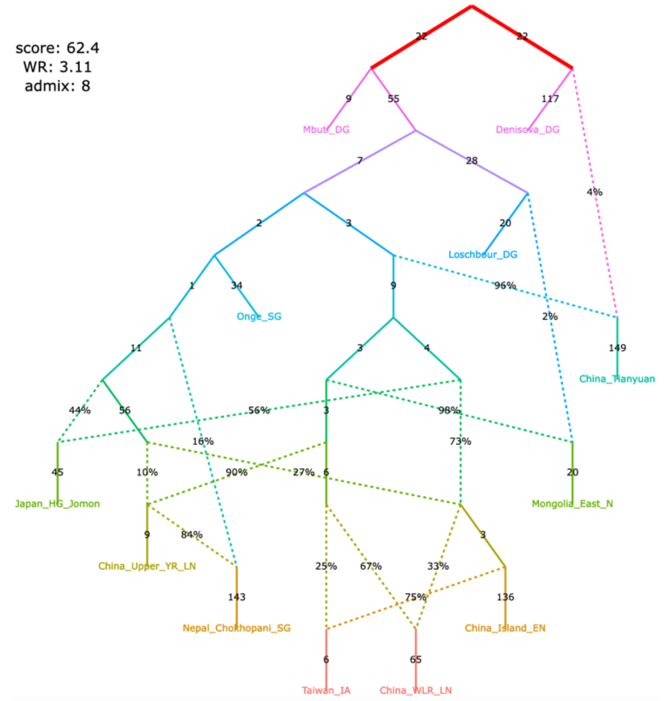

A claim by Wang et al. 2021 relying on the admixture graph:

There is Onge-related admixture in the following lineages: Jomon (Japan_HG_Jomon), Tibetan (Nepal_Chokhopani_SG), Upper Yellow River Late Neolithic (China_Upper_YR_LN), West Liao River Late Neolithic (China_WLR_LN), Taiwan Iron Age (Taiwan_IA), China Island Early Neolithic (Liangdao, China_Island_EN).

Figure S11

#### **c**, alternative models fitting significantly better than the published one

| | |
|---|---|
| no. of East Asian groups with Onge-related ancestry | 1 |
| no. of East Asian groups with European-related ancestry | 8 |
| direction of gene flow China_Island_EN > Taiwan_IA | forward |
| Mongolia_East is basal with respect to China_Upper_YR and Nepal_Chokhopani | yes |
| Taiwan_IA is basal with respect to Mongolia_East and Nepal_Chokhopani | yes |

The original set of SNPs and individuals and the corrected algorithm for calculating $f_3$-statistics (203,753 variable site with no missing data at the group level)

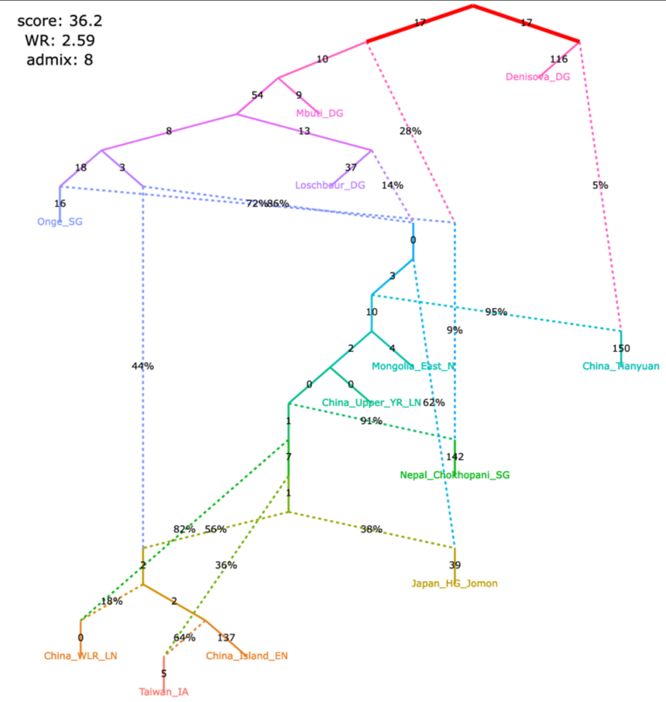

| | |
|---|---|
| no. of East Asian groups with Onge-related ancestry | 1 |
| no. of East Asian groups with European-related ancestry | 8 |
| direction of gene flow China_Island_EN > Taiwan_IA | forward |
| Mongolia_East is basal with respect to China_Upper_YR and Nepal_Chokhopani | yes |
| Taiwan_IA is basal with respect to Mongolia_East and Nepal_Chokhopani | yes |

The original set of SNPs and individuals and the original algorithm for calculating $f_3$-statistics (496,233 variable site with no missing data at the group level)

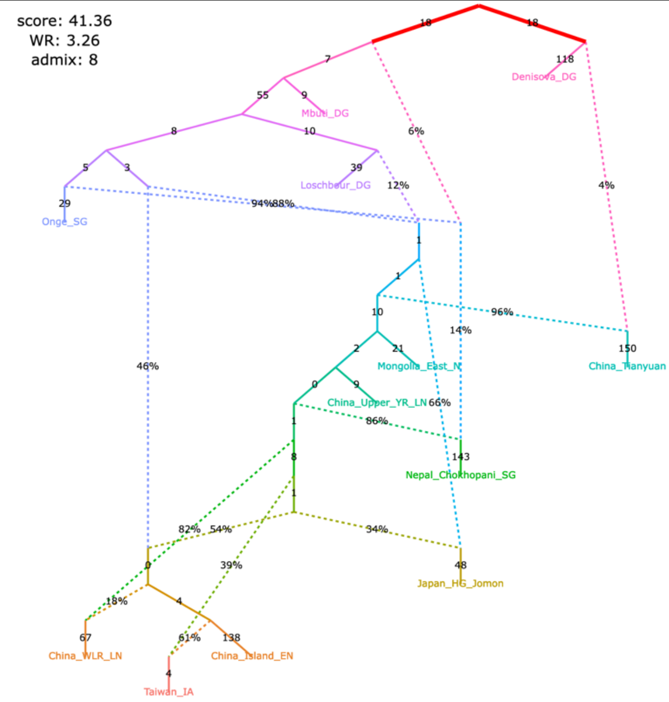

| | |
|---|---|
| no. of East Asian groups with Onge-related ancestry | 2 |
| no. of East Asian groups with European-related ancestry | 8 |
| direction of gene flow China_Island_EN > Taiwan_IA | forward |
| Mongolia_East is basal with respect to China_Upper_YR and Nepal_Chokhopani | yes |
| Taiwan_IA is basal with respect to Mongolia_East and Nepal_Chokhopani | yes |

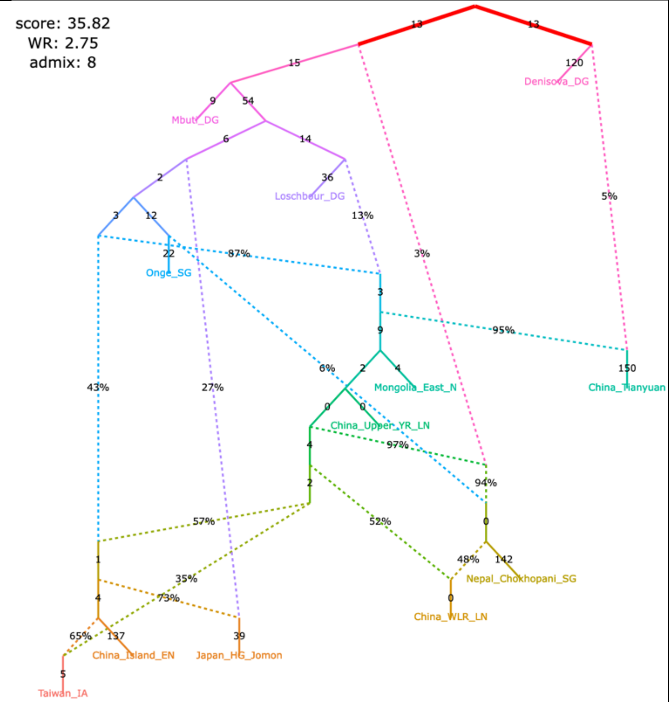

| | |
|---|---|
| no. of East Asian groups with Onge-related ancestry | 2 |
| no. of East Asian groups with European-related ancestry | 8 |
| direction of gene flow China_Island_EN > Taiwan_IA | forward |
| Mongolia_East is basal with respect to China_Upper_YR and Nepal_Chokhopani | yes |
| Taiwan_IA is basal with respect to Mongolia_East and Nepal_Chokhopani | yes |

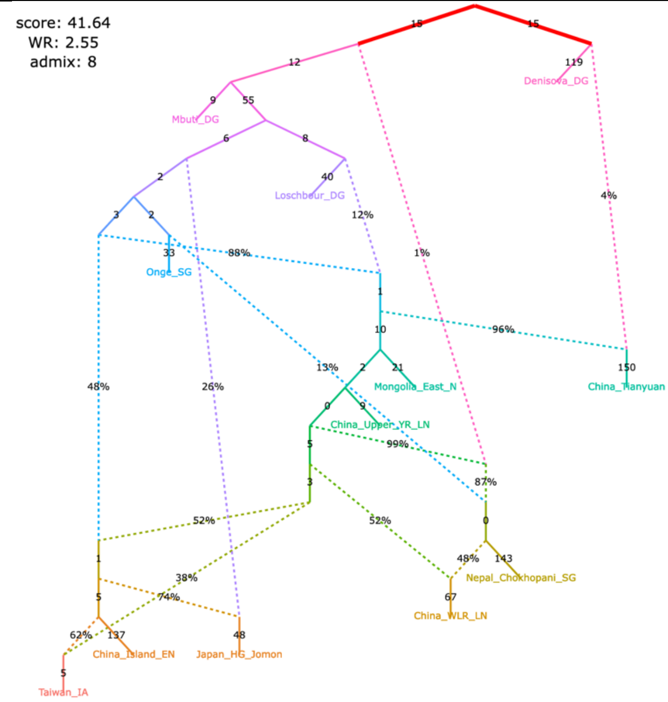

Figure S11

#### **d**, alternative models fitting significantly better than the published one

| | |
|---|---|
| no. of East Asian groups with Onge-related ancestry | 1 |
| no. of East Asian groups with European-related ancestry | 8 |
| direction of gene flow China_Island_EN > Taiwan_IA | no gene flow |
| Mongolia_East is basal with respect to China_Upper_YR and Nepal_Chokhopani | yes |
| Taiwan_IA is basal with respect to Mongolia_East and Nepal_Chokhopani | yes |

The original set of SNPs and individuals and the corrected algorithm for calculating $f_3$-statistics (203,753 variable site with no missing data at the group level)

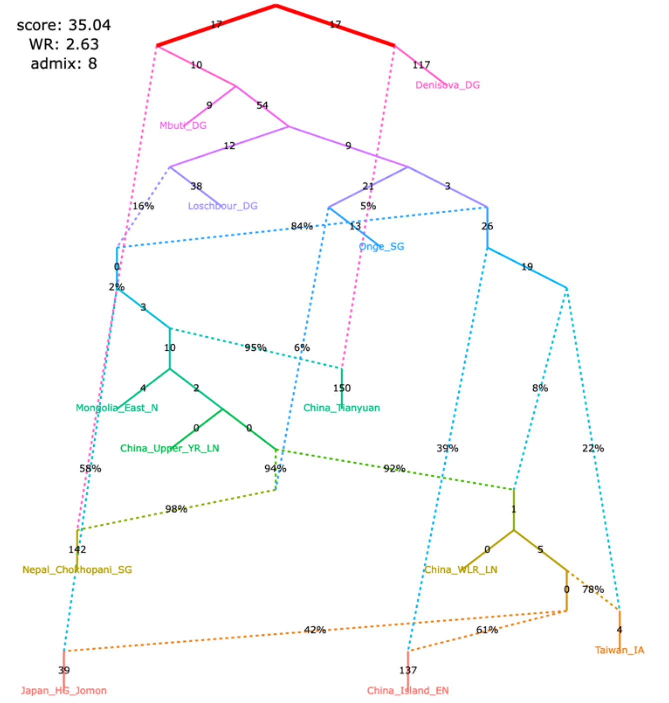

| | |
|---|---|
| no. of East Asian groups with Onge-related ancestry | 1 |
| no. of East Asian groups with European-related ancestry | 8 |
| direction of gene flow China_Island_EN > Taiwan_IA | no gene flow |
| Mongolia_East is basal with respect to China_Upper_YR and Nepal_Chokhopani | yes |
| Taiwan_IA is basal with respect to Mongolia_East and Nepal_Chokhopani | yes |

The original set of SNPs and individuals and the original algorithm for calculating $f_3$-statistics (496,233 variable site with no missing data at the group level)

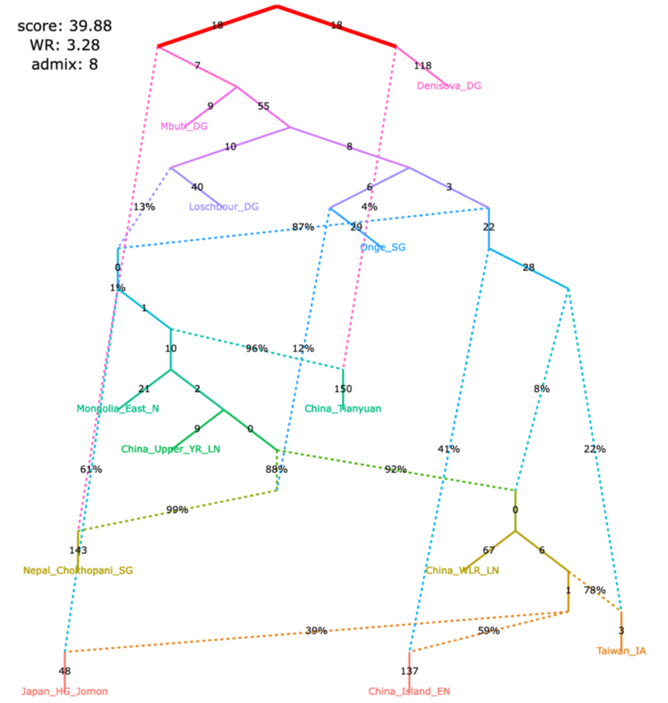

| | |
|---|---|
| no. of East Asian groups with Onge-related ancestry | 1 |
| no. of East Asian groups with European-related ancestry | 8 |
| direction of gene flow China_Island_EN > Taiwan_IA | reverse |
| Mongolia_East is basal with respect to China_Upper_YR and Nepal_Chokhopani | yes |
| Taiwan_IA is basal with respect to Mongolia_East and Nepal_Chokhopani | yes |

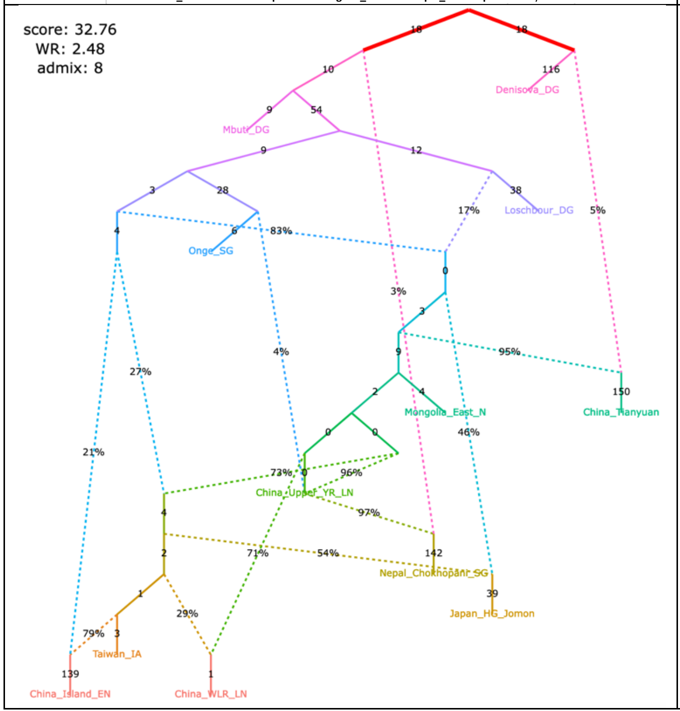

| | |
|---|---|
| no. of East Asian groups with Onge-related ancestry | 1 |
| no. of East Asian groups with European-related ancestry | 8 |
| direction of gene flow China_Island_EN > Taiwan_IA | reverse |
| Mongolia_East is basal with respect to China_Upper_YR and Nepal_Chokhopani | yes |
| Taiwan_IA is basal with respect to Mongolia_East and Nepal_Chokhopani | yes |

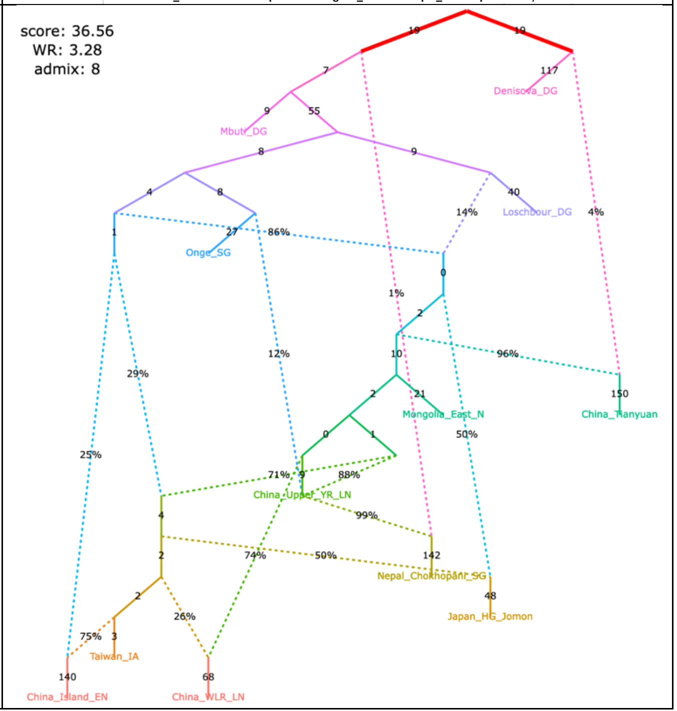

Figure S11

#### **e**, alternative models fitting significantly better than the published one

| | |
|---|---|
| no. of East Asian groups with Onge-related ancestry | 1 |
| no. of East Asian groups with European-related ancestry | 8 |
| direction of gene flow China_Island_EN > Taiwan_IA | reverse |
| Mongolia_East is basal with respect to China_Upper_YR and Nepal_Chokhopani | yes |
| Taiwan_IA is basal with respect to Mongolia_East and Nepal_Chokhopani | yes |

| | |
|---|---|
| no. of East Asian groups with Onge-related ancestry | 1 |
| no. of East Asian groups with European-related ancestry | 8 |
| direction of gene flow China_Island_EN > Taiwan_IA | reverse |
| Mongolia_East is basal with respect to China_Upper_YR and Nepal_Chokhopani | yes |
| Taiwan_IA is basal with respect to Mongolia_East and Nepal_Chokhopani | yes |

The original set of SNPs and individuals and the corrected algorithm for calculating $f_3$-statistics (203,753 variable site with no missing data at the group level)

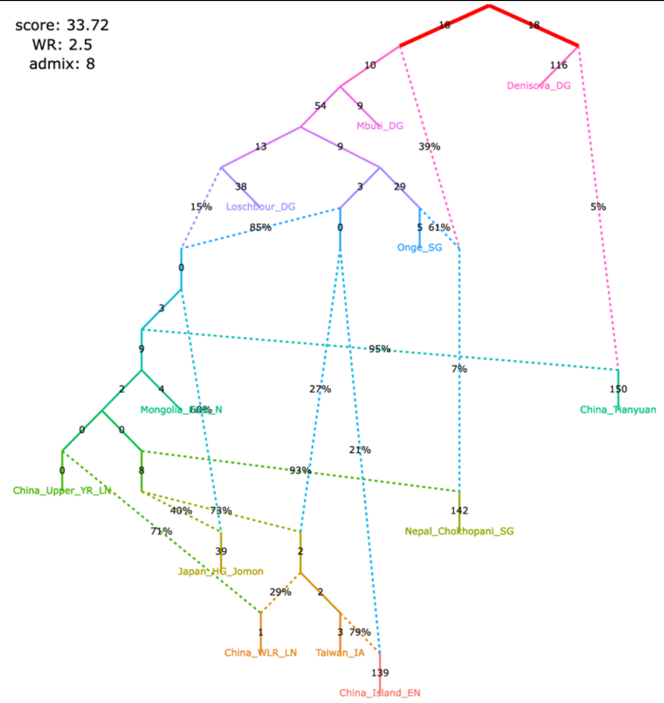

The original set of SNPs and individuals and the original algorithm for calculating $f_3$-statistics (496,233 variable site with no missing data at the group level)

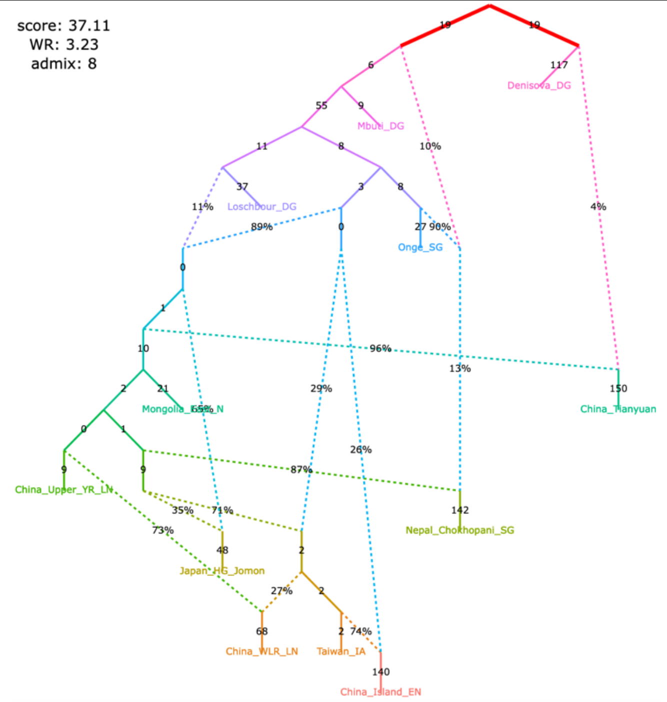

| | |
|---|---|
| no. of East Asian groups with Onge-related ancestry | 1 |
| no. of East Asian groups with European-related ancestry | 8 |
| direction of gene flow China_Island_EN > Taiwan_IA | reverse |
| Mongolia_East is basal with respect to China_Upper_YR and Nepal_Chokhopani | yes |
| Taiwan_IA is basal with respect to Mongolia_East and Nepal_Chokhopani | yes |

| | |
|---|---|
| no. of East Asian groups with Onge-related ancestry | 1 |
| no. of East Asian groups with European-related ancestry | 8 |
| direction of gene flow China_Island_EN > Taiwan_IA | reverse |
| Mongolia_East is basal with respect to China_Upper_YR and Nepal_Chokhopani | yes |
| Taiwan_IA is basal with respect to Mongolia_East and Nepal_Chokhopani | yes |

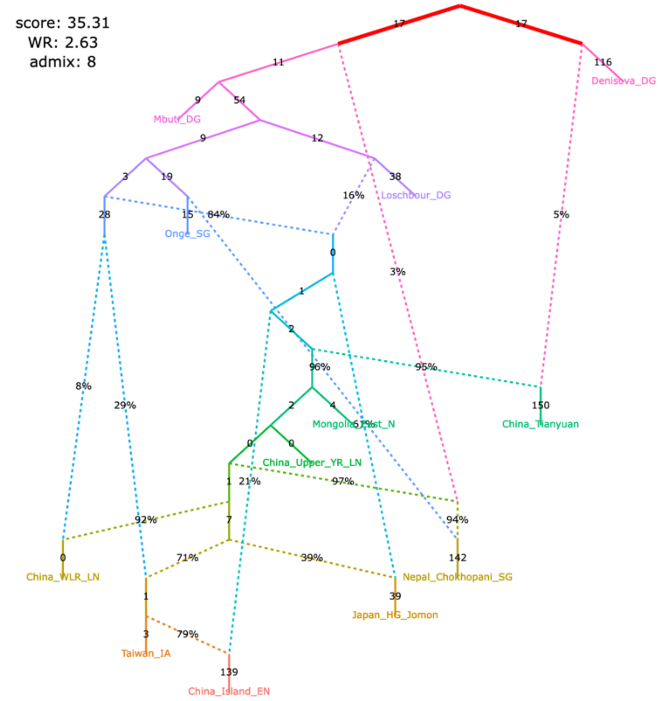

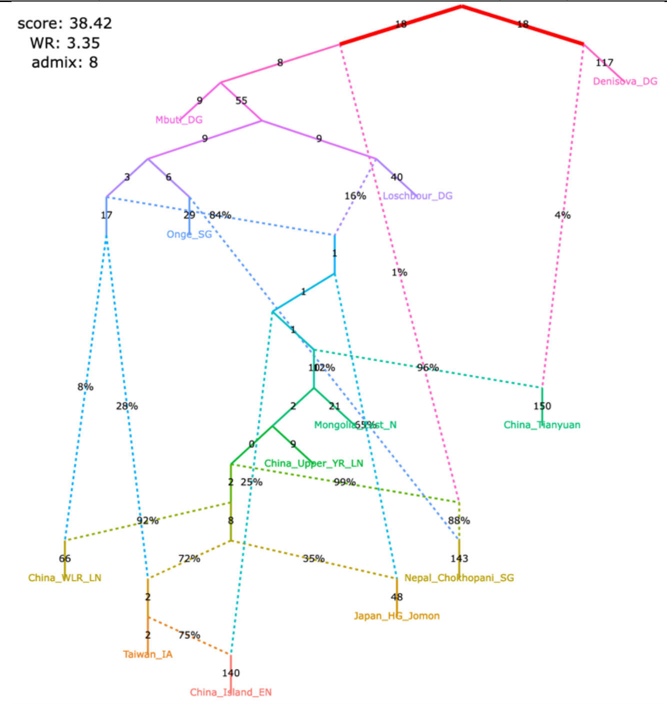

Figure S11

#### **f**, alternative models fitting significantly better than the published one

| | |
|---|---|
| no. of East Asian groups with Onge-related ancestry | 1 |
| no. of East Asian groups with European-related ancestry | 8 |
| direction of gene flow China_Island_EN > Taiwan_IA | reverse |
| Mongolia_East is basal with respect to China_Upper_YR and Nepal_Chokhopani | yes |
| Taiwan_IA is basal with respect to Mongolia_East and Nepal_Chokhopani | yes |

| | |
|---|---|
| no. of East Asian groups with Onge-related ancestry | 1 |
| no. of East Asian groups with European-related ancestry | 8 |
| direction of gene flow China_Island_EN > Taiwan_IA | reverse |
| Mongolia_East is basal with respect to China_Upper_YR and Nepal_Chokhopani | yes |
| Taiwan_IA is basal with respect to Mongolia_East and Nepal_Chokhopani | yes |

| The original set of SNPs and individuals and the corrected algorithm for calculating $f_3$-statistics (203,753 variable site with no missing data at the group level) | The original set of SNPs and individuals and the original algorithm for calculating $f_3$-statistics (496,233 variable site with no missing data at the group level) |
|---|---|

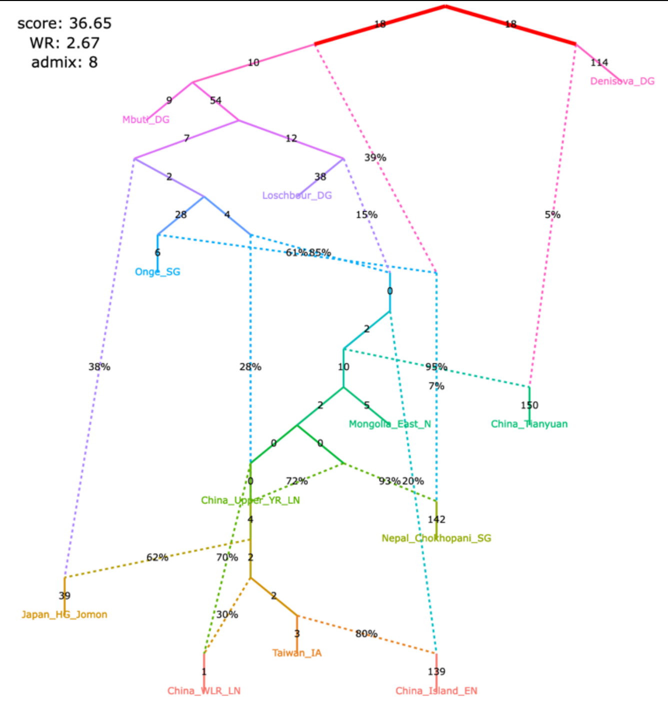

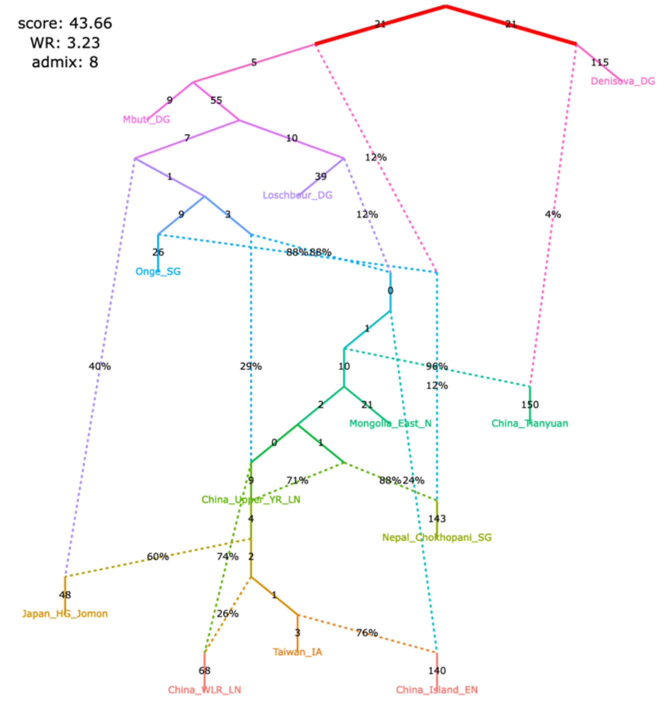

| | |
|---|---|
| no. of East Asian groups with Onge-related ancestry | 1 |
| no. of East Asian groups with European-related ancestry | 8 |
| direction of gene flow China_Island_EN > Taiwan_IA | reverse |
| Mongolia_East is basal with respect to China_Upper_YR and Nepal_Chokhopani | yes |
| Taiwan_IA is basal with respect to Mongolia_East and Nepal_Chokhopani | yes |

| | |
|---|---|
| no. of East Asian groups with Onge-related ancestry | 1 |
| no. of East Asian groups with European-related ancestry | 8 |
| direction of gene flow China_Island_EN > Taiwan_IA | reverse |
| Mongolia_East is basal with respect to China_Upper_YR and Nepal_Chokhopani | yes |
| Taiwan_IA is basal with respect to Mongolia_East and Nepal_Chokhopani | yes |

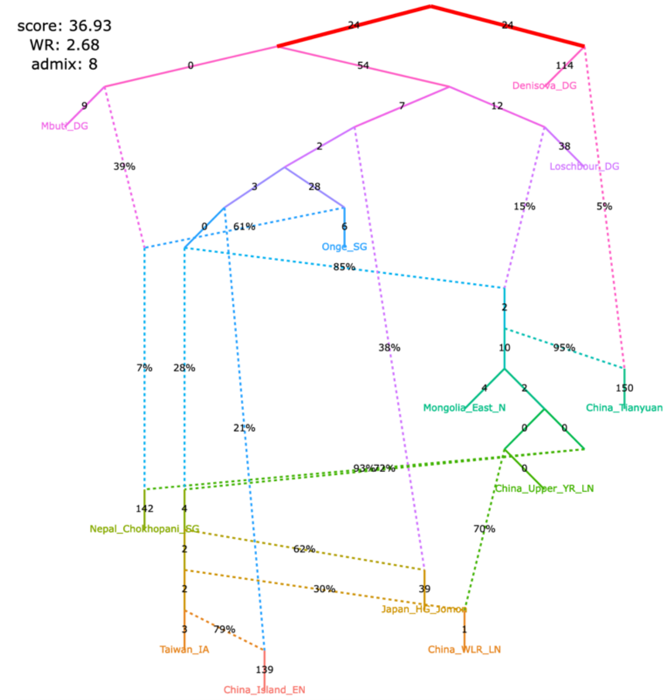

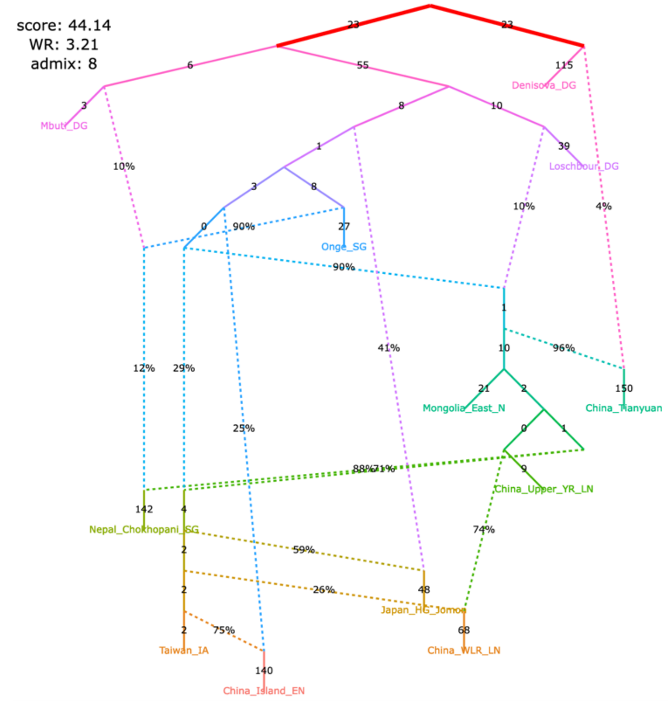

Figure S11

#### **g**, alternative models fitting significantly better than the published one

| | |
|---|---|
| no. of East Asian groups with Onge-related ancestry | 0 |
| no. of East Asian groups with European-related ancestry | 8 |
| direction of gene flow China_Island_EN > Taiwan_IA | no gene flow |
| Mongolia_East is basal with respect to China_Upper_YR and Nepal_Chokhopani | yes |
| Taiwan_IA is basal with respect to Mongolia_East and Nepal_Chokhopani | yes |

The original set of SNPs and individuals and the corrected algorithm for calculating $f_3$-statistics (203,753 variable site with no missing data at the group level)

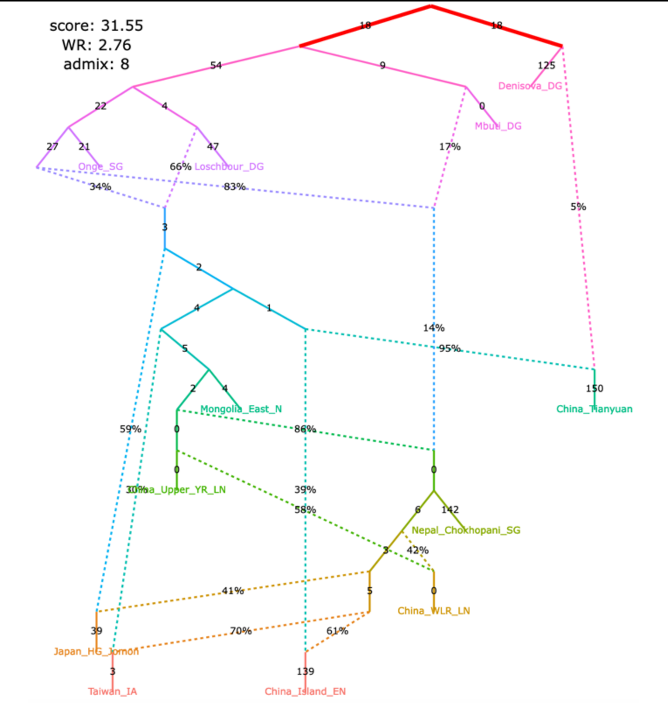

| | |
|---|---|
| no. of East Asian groups with Onge-related ancestry | 0 |
| no. of East Asian groups with European-related ancestry | 8 |
| direction of gene flow China_Island_EN > Taiwan_IA | no gene flow |
| Mongolia_East is basal with respect to China_Upper_YR and Nepal_Chokhopani | yes |
| Taiwan_IA is basal with respect to Mongolia_East and Nepal_Chokhopani | yes |

The original set of SNPs and individuals and the original algorithm for calculating $f_3$-statistics (496,233 variable site with no missing data at the group level)

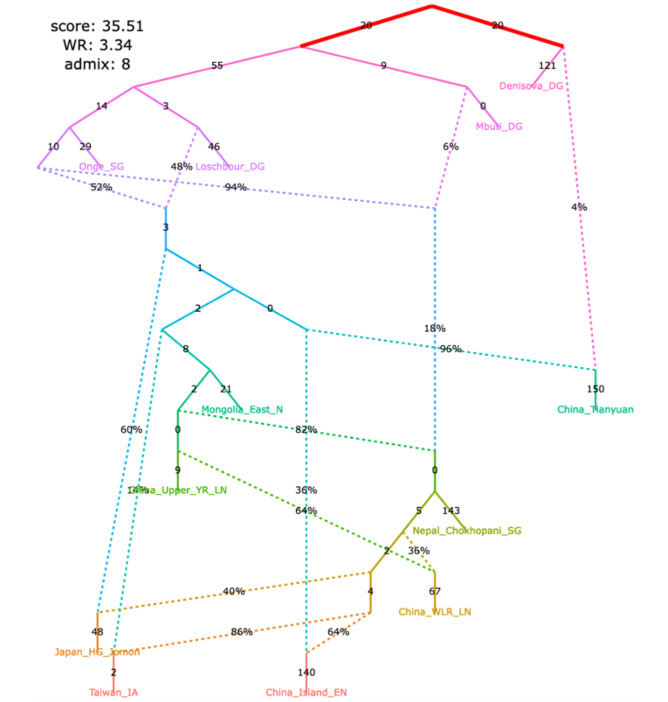

| | |
|---|---|
| no. of East Asian groups with Onge-related ancestry | 0 |
| no. of East Asian groups with European-related ancestry | 8 |
| direction of gene flow China_Island_EN > Taiwan_IA | no gene flow |
| Mongolia_East is basal with respect to China_Upper_YR and Nepal_Chokhopani | yes |
| Taiwan_IA is basal with respect to Mongolia_East and Nepal_Chokhopani | yes |

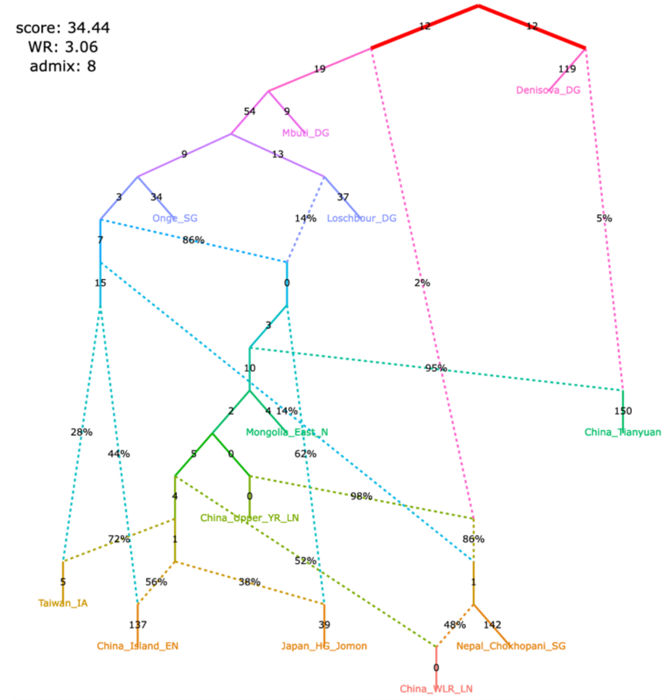

| | |
|---|---|
| no. of East Asian groups with Onge-related ancestry | 0 |
| no. of East Asian groups with European-related ancestry | 8 |
| direction of gene flow China_Island_EN > Taiwan_IA | no gene flow |
| Mongolia_East is basal with respect to China_Upper_YR and Nepal_Chokhopani | yes |
| Taiwan_IA is basal with respect to Mongolia_East and Nepal_Chokhopani | yes |

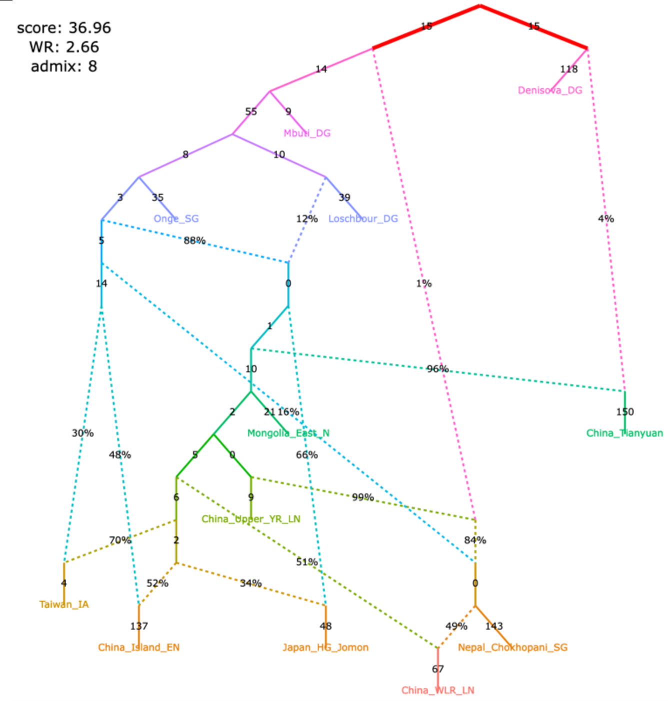

Figure S11

#### **h**, alternative models fitting significantly better than the published one

| | |
|---|---|
| no. of East Asian groups with Onge-related ancestry | 0 |
| no. of East Asian groups with European-related ancestry | 8 |
| direction of gene flow China_Island_EN > Taiwan_IA | reverse |
| Mongolia_East is basal with respect to China_Upper_YR and Nepal_Chokhopani | yes |
| Taiwan_IA is basal with respect to Mongolia_East and Nepal_Chokhopani | unclear |

The original set of SNPs and individuals and the corrected algorithm for calculating $f_3$-statistics (203,753 variable site with no missing data at the group level)

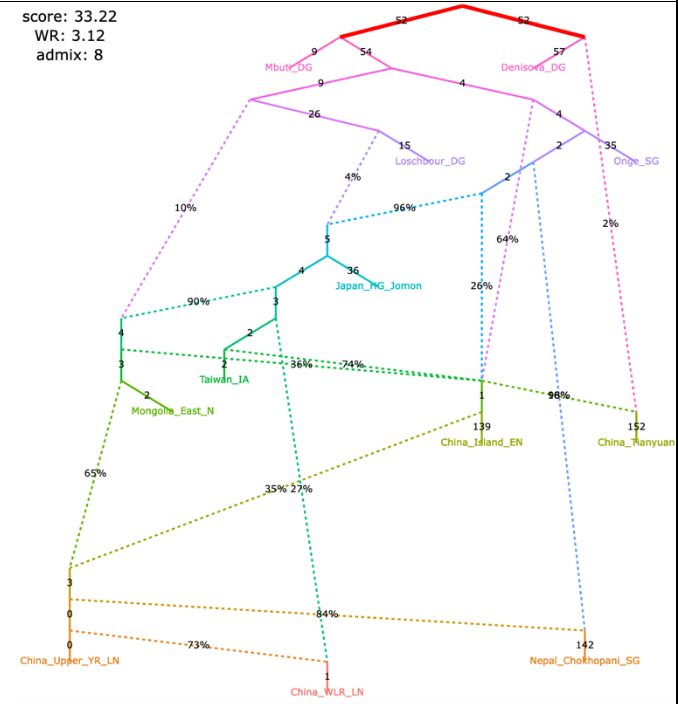

| | |
|---|---|
| no. of East Asian groups with Onge-related ancestry | 0 |
| no. of East Asian groups with European-related ancestry | 8 |
| direction of gene flow China_Island_EN > Taiwan_IA | reverse |
| Mongolia_East is basal with respect to China_Upper_YR and Nepal_Chokhopani | yes |
| Taiwan_IA is basal with respect to Mongolia_East and Nepal_Chokhopani | unclear |

The original set of SNPs and individuals and the original algorithm for calculating $f_3$-statistics (496,233 variable site with no missing data at the group level)

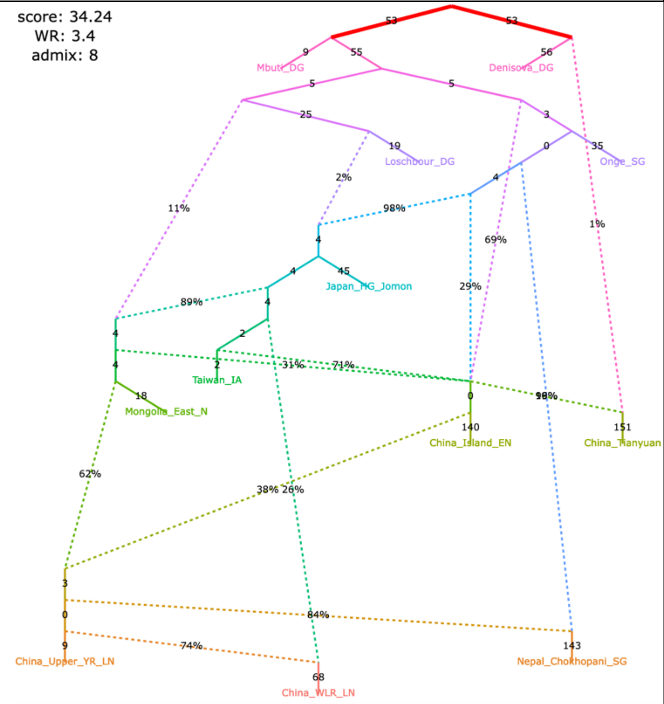

| | |
|---|---|
| no. of East Asian groups with Onge-related ancestry | 0 |
| no. of East Asian groups with European-related ancestry | 8 |
| direction of gene flow China_Island_EN > Taiwan_IA | reverse |
| Mongolia_East is basal with respect to China_Upper_YR and Nepal_Chokhopani | yes |
| Taiwan_IA is basal with respect to Mongolia_East and Nepal_Chokhopani | yes |

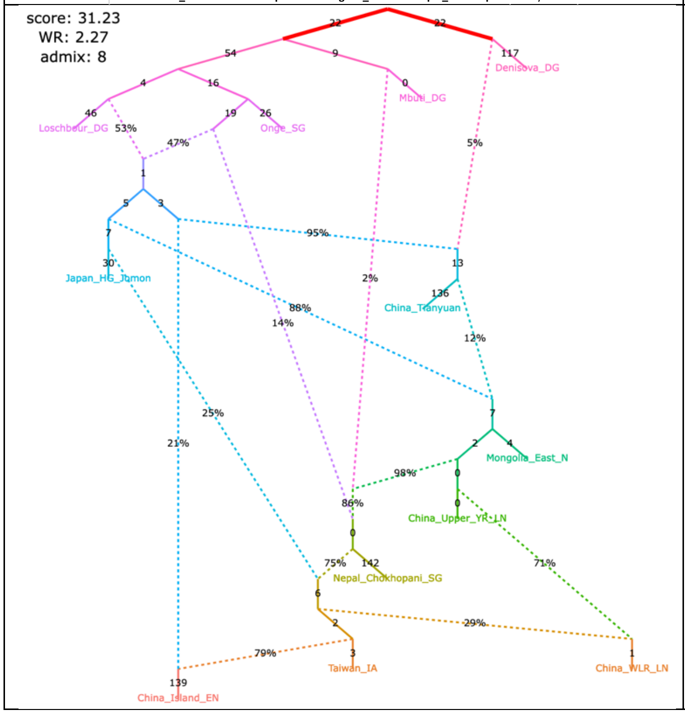

| | |
|---|---|
| no. of East Asian groups with Onge-related ancestry | 0 |
| no. of East Asian groups with European-related ancestry | 8 |
| direction of gene flow China_Island_EN > Taiwan_IA | reverse |
| Mongolia_East is basal with respect to China_Upper_YR and Nepal_Chokhopani | yes |
| Taiwan_IA is basal with respect to Mongolia_East and Nepal_Chokhopani | yes |

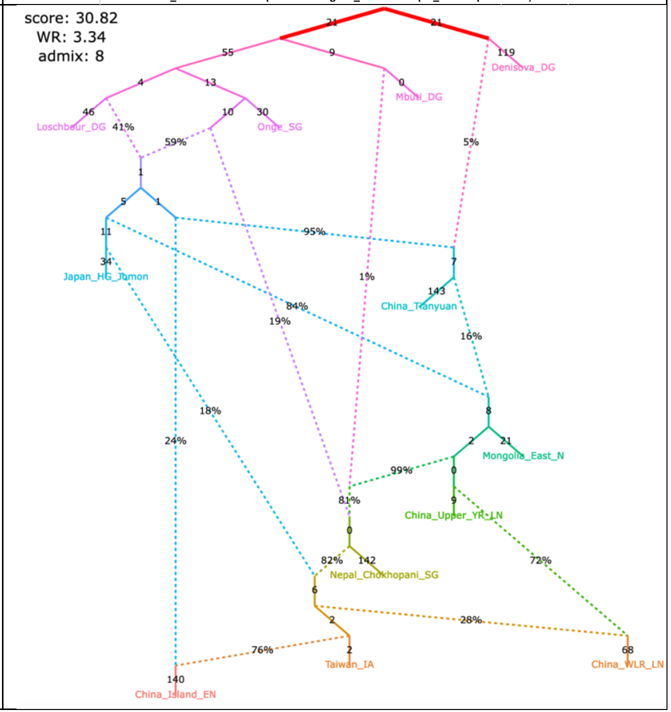

Figure S11

#### **i**, alternative models fitting significantly better than the published one

| | |
|---|---|
| no. of East Asian groups with Onge-related ancestry | 0 |
| no. of East Asian groups with European-related ancestry | 8 |
| direction of gene flow China_Island_EN > Taiwan_IA | reverse |
| Mongolia_East is basal with respect to China_Upper_YR and Nepal_Chokhopani | yes |
| Taiwan_IA is basal with respect to Mongolia_East and Nepal_Chokhopani | yes |

The original set of SNPs and individuals and the corrected algorithm for calculating $f_3$-statistics (203,753 variable site with no missing data at the group level)

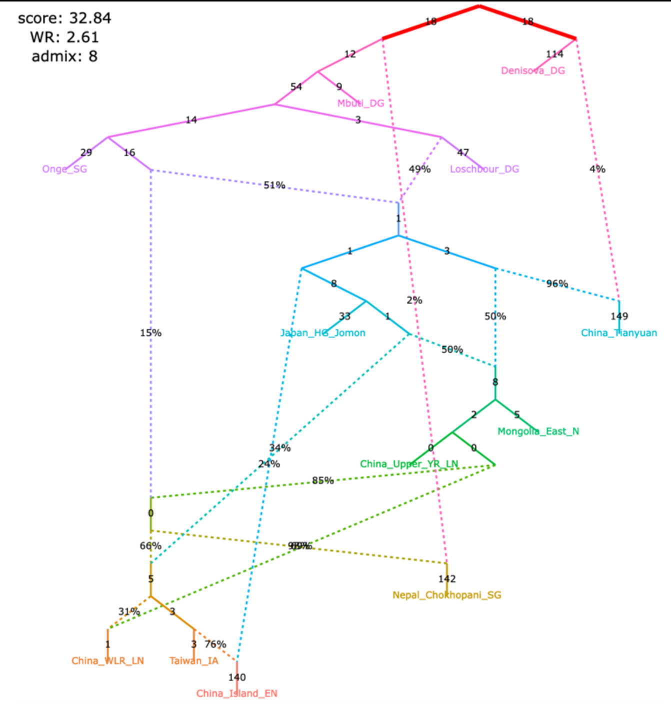

| | |
|---|---|
| no. of East Asian groups with Onge-related ancestry | 0 |
| no. of East Asian groups with European-related ancestry | 8 |
| direction of gene flow China_Island_EN > Taiwan_IA | reverse |
| Mongolia_East is basal with respect to China_Upper_YR and Nepal_Chokhopani | yes |
| Taiwan_IA is basal with respect to Mongolia_East and Nepal_Chokhopani | yes |

The original set of SNPs and individuals and the original algorithm for calculating $f_3$-statistics (496,233 variable site with no missing data at the group level)

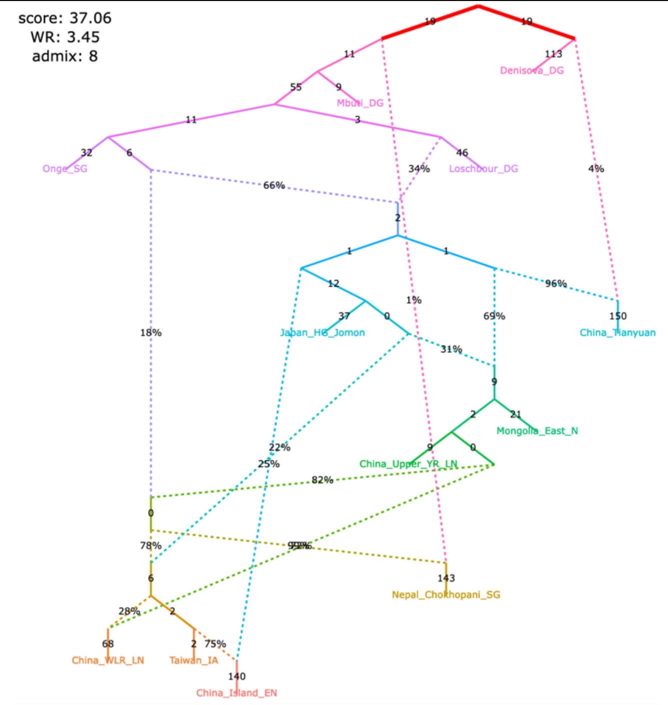

| | |
|---|---|
| no. of East Asian groups with Onge-related ancestry | 0 |
| no. of East Asian groups with European-related ancestry | 8 |
| direction of gene flow China_Island_EN > Taiwan_IA | reverse |
| Mongolia_East is basal with respect to China_Upper_YR and Nepal_Chokhopani | yes |
| Taiwan_IA is basal with respect to Mongolia_East and Nepal_Chokhopani | yes |

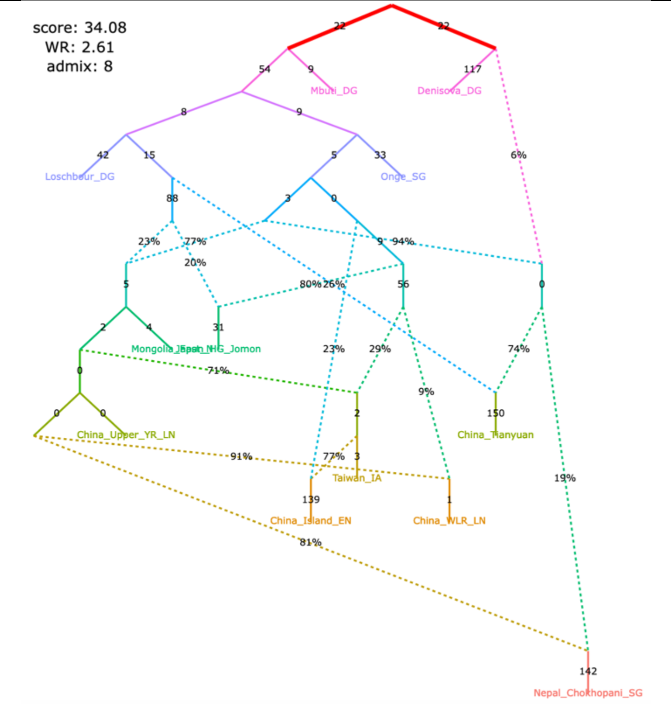

| | |
|---|---|
| no. of East Asian groups with Onge-related ancestry | 0 |
| no. of East Asian groups with European-related ancestry | 8 |
| direction of gene flow China_Island_EN > Taiwan_IA | reverse |
| Mongolia_East is basal with respect to China_Upper_YR and Nepal_Chokhopani | yes |
| Taiwan_IA is basal with respect to Mongolia_East and Nepal_Chokhopani | yes |

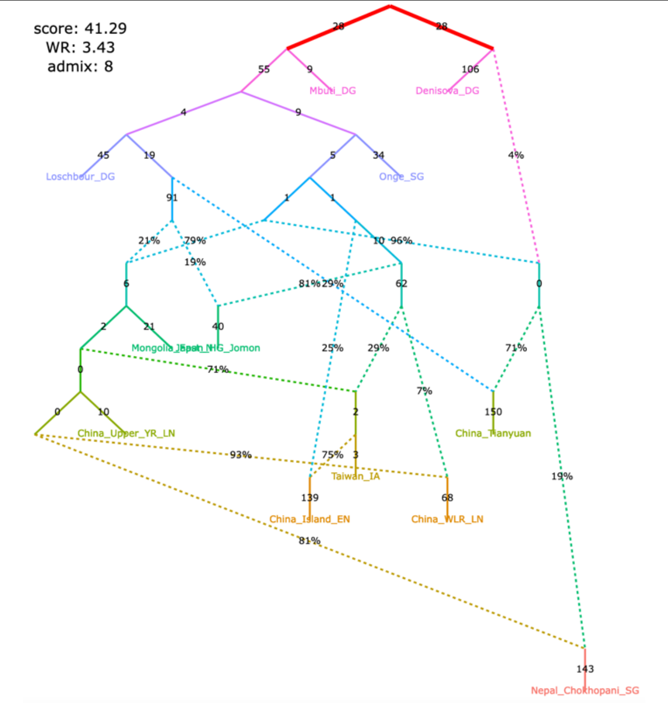

Figure S11

#### **j**, alternative models fitting significantly better than the published one

| | |
|---|---|
| no. of East Asian groups with Onge-related ancestry | 0 |
| no. of East Asian groups with European-related ancestry | 8 |
| direction of gene flow China_Island_EN > Taiwan_IA | reverse |
| Mongolia_East is basal with respect to China_Upper_YR and Nepal_Chokhopani | yes |
| Taiwan_IA is basal with respect to Mongolia_East and Nepal_Chokhopani | yes |

The original set of SNPs and individuals and the corrected algorithm for calculating $f_3$-statistics (203,753 variable site with no missing data at the group level)

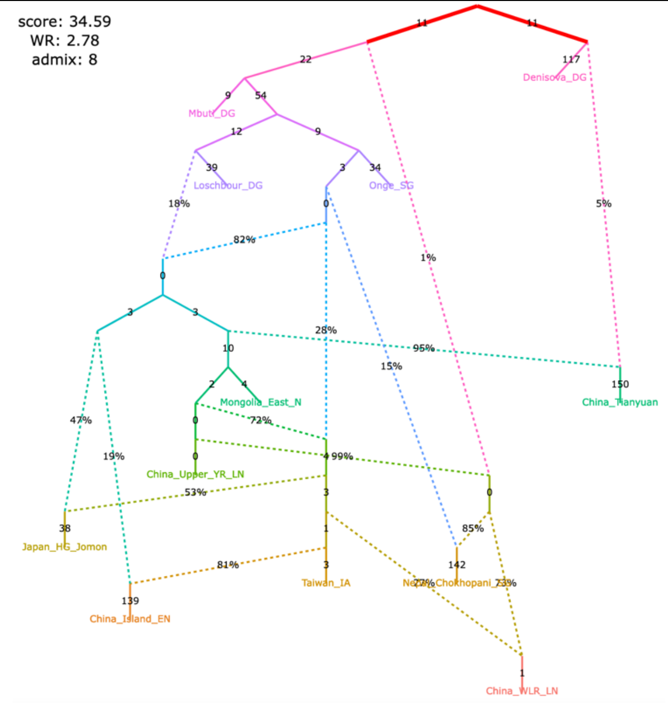

| | |
|---|---|
| no. of East Asian groups with Onge-related ancestry | 0 |
| no. of East Asian groups with European-related ancestry | 8 |
| direction of gene flow China_Island_EN > Taiwan_IA | reverse |
| Mongolia_East is basal with respect to China_Upper_YR and Nepal_Chokhopani | yes |
| Taiwan_IA is basal with respect to Mongolia_East and Nepal_Chokhopani | yes |

The original set of SNPs and individuals and the original algorithm for calculating $f_3$-statistics (496,233 variable site with no missing data at the group level)

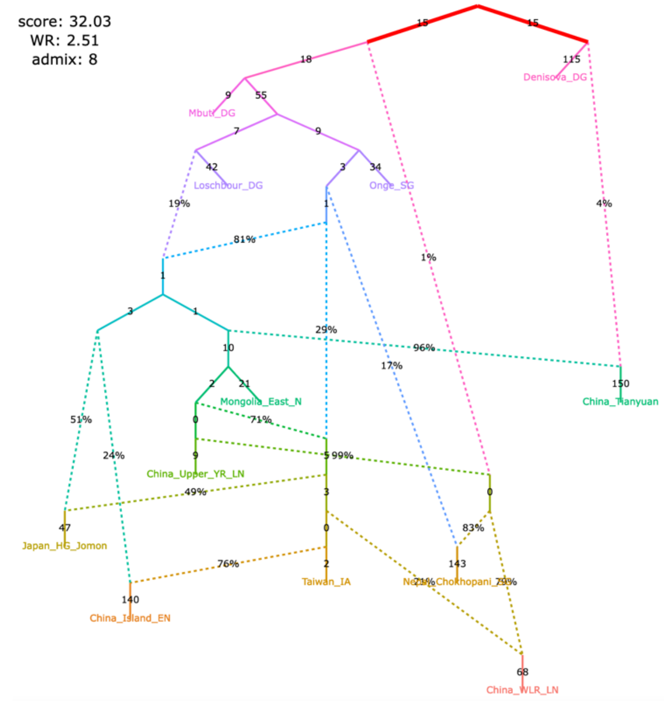

| | |
|---|---|
| no. of East Asian groups with Onge-related ancestry | 0 |
| no. of East Asian groups with European-related ancestry | 8 |
| direction of gene flow China_Island_EN > Taiwan_IA | reverse |
| Mongolia_East is basal with respect to China_Upper_YR and Nepal_Chokhopani | yes |
| Taiwan_IA is basal with respect to Mongolia_East and Nepal_Chokhopani | yes |

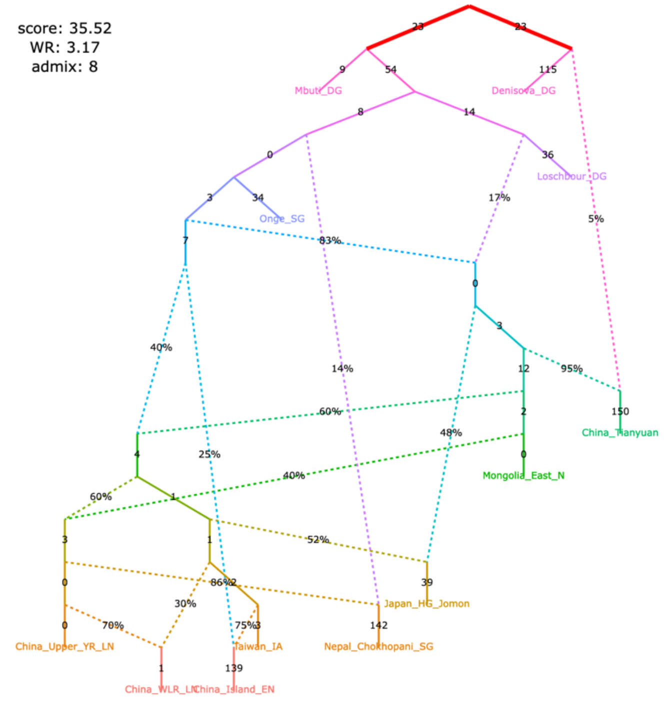

| | |
|---|---|
| no. of East Asian groups with Onge-related ancestry | 0 |
| no. of East Asian groups with European-related ancestry | 8 |
| direction of gene flow China_Island_EN > Taiwan_IA | reverse |
| Mongolia_East is basal with respect to China_Upper_YR and Nepal_Chokhopani | yes |
| Taiwan_IA is basal with respect to Mongolia_East and Nepal_Chokhopani | yes |

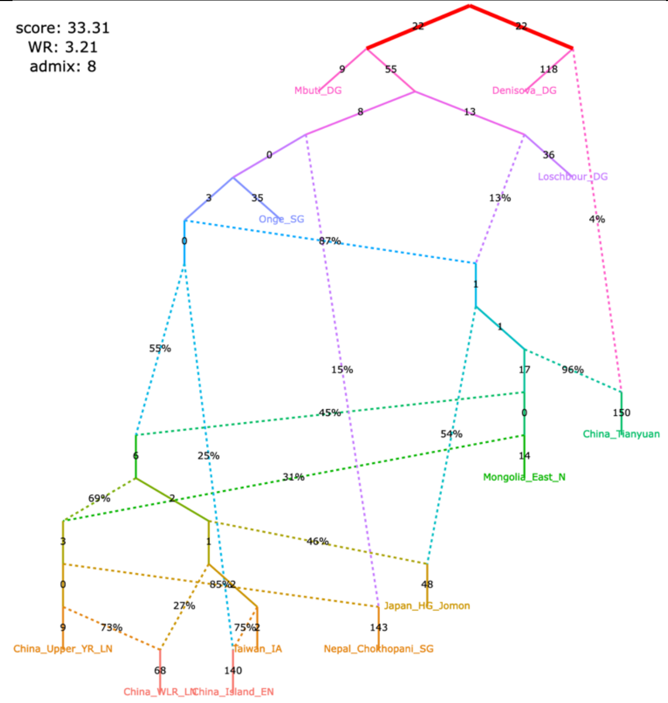

Figure S11

#### **k**, alternative models fitting significantly better than the published one

| | |
|---|---|
| no. of East Asian groups with Onge-related ancestry | 0 |
| no. of East Asian groups with European-related ancestry | 8 |
| direction of gene flow China_Island_EN > Taiwan_IA | reverse |
| Mongolia_East is basal with respect to China_Upper_YR and Nepal_Chokhopani | yes |
| Taiwan_IA is basal with respect to Mongolia_East and Nepal_Chokhopani | yes |

The original set of SNPs and individuals and the corrected algorithm for calculating $f_3$-statistics (203,753 variable site with no missing data at the group level)

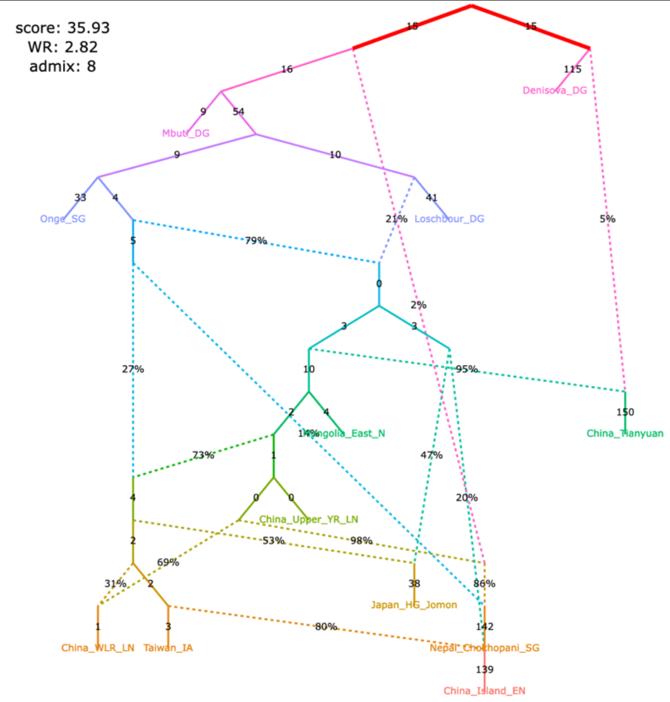

| | |
|---|---|
| no. of East Asian groups with Onge-related ancestry | 0 |
| no. of East Asian groups with European-related ancestry | 8 |
| direction of gene flow China_Island_EN > Taiwan_IA | reverse |
| Mongolia_East is basal with respect to China_Upper_YR and Nepal_Chokhopani | yes |
| Taiwan_IA is basal with respect to Mongolia_East and Nepal_Chokhopani | yes |

The original set of SNPs and individuals and the original algorithm for calculating $f_3$-statistics (496,233 variable site with no missing data at the group level)

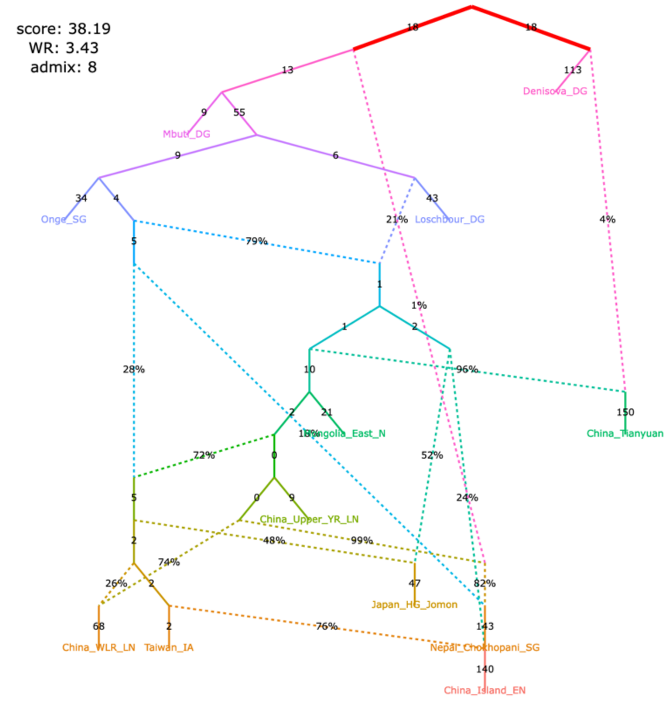

| | |
|---|---|
| no. of East Asian groups with Onge-related ancestry | 0 |
| no. of East Asian groups with European-related ancestry | 8 |
| direction of gene flow China_Island_EN > Taiwan_IA | reverse |
| Mongolia_East is basal with respect to China_Upper_YR and Nepal_Chokhopani | yes |
| Taiwan_IA is basal with respect to Mongolia_East and Nepal_Chokhopani | yes |

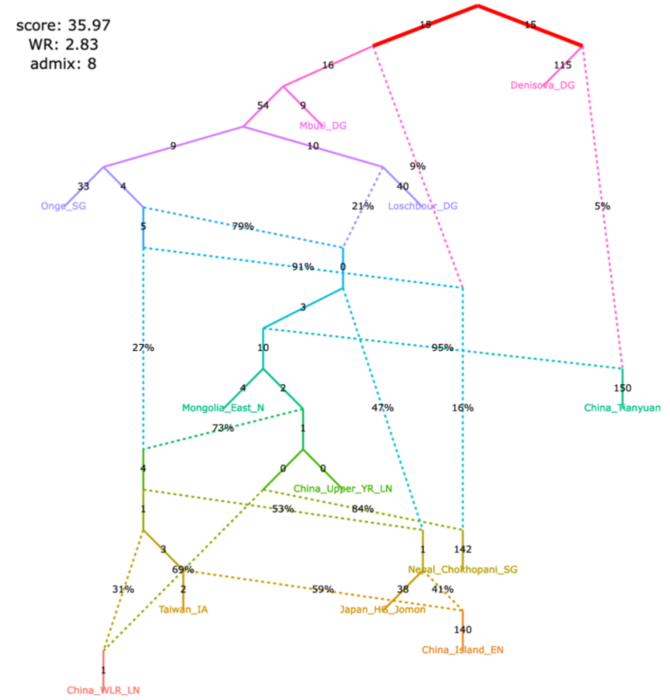

| | |
|---|---|
| no. of East Asian groups with Onge-related ancestry | 0 |
| no. of East Asian groups with European-related ancestry | 8 |
| direction of gene flow China_Island_EN > Taiwan_IA | reverse |
| Mongolia_East is basal with respect to China_Upper_YR and Nepal_Chokhopani | yes |
| Taiwan_IA is basal with respect to Mongolia_East and Nepal_Chokhopani | yes |

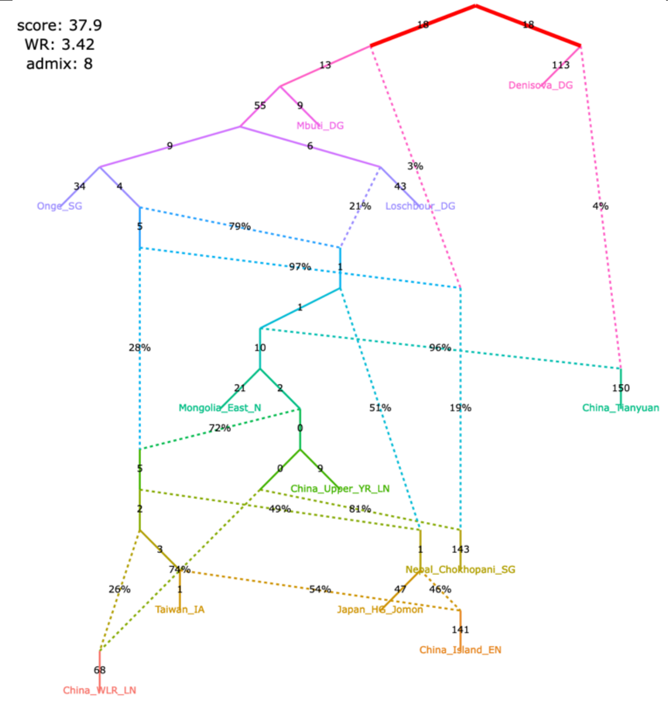

Figure S11

#### **I**, alternative models fitting significantly better than the published one

| | Left model | Right model |
|---|---|---|
| no. of East Asian groups with Onge-related ancestry | 0 | 0 |
| no. of East Asian groups with European-related ancestry | 8 | 8 |
| direction of gene flow China_Island_EN > Taiwan_IA | reverse | reverse |
| Mongolia_East is basal with respect to China_Upper_YR and Nepal_Chokhopani | yes | yes |
| Taiwan_IA is basal with respect to Mongolia_East and Nepal_Chokhopani | yes | yes |

The original set of SNPs and individuals and the corrected algorithm for calculating $f_3$-statistics (203,753 variable site with no missing data at the group level)

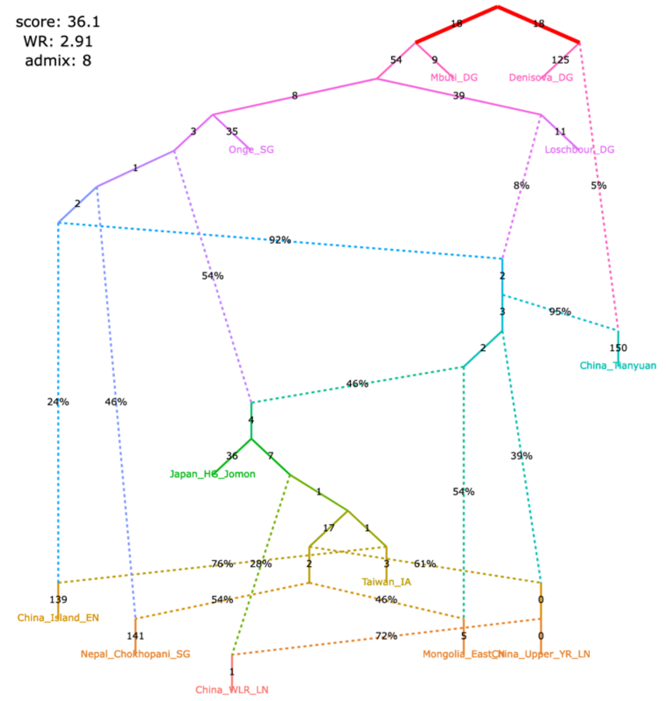

The original set of SNPs and individuals and the original algorithm for calculating $f_3$-statistics (496,233 variable site with no missing data at the group level)

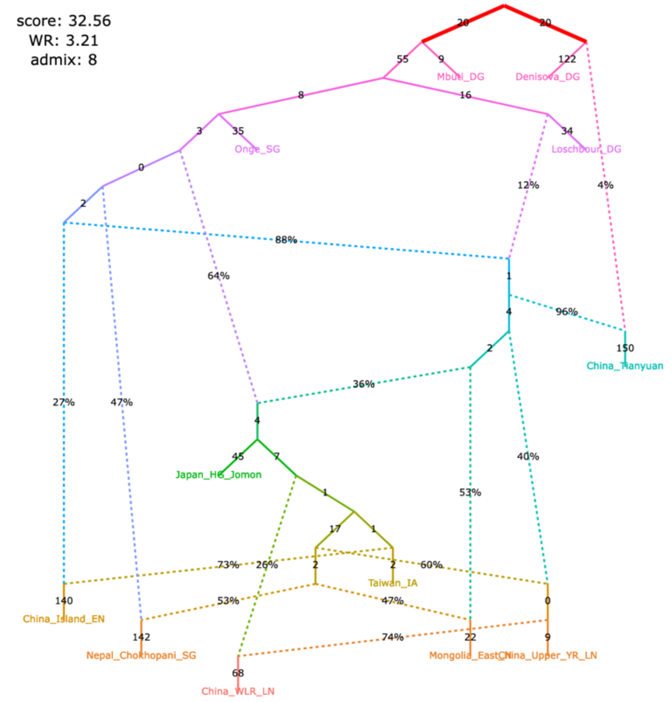

| | Left model | Right model |
|---|---|---|
| no. of East Asian groups with Onge-related ancestry | 0 | 0 |
| no. of East Asian groups with European-related ancestry | 5 | 5 |
| direction of gene flow China_Island_EN > Taiwan_IA | no gene flow | no gene flow |
| Mongolia_East is basal with respect to China_Upper_YR and Nepal_Chokhopani | yes | yes |
| Taiwan_IA is basal with respect to Mongolia_East and Nepal_Chokhopani | unclear | unclear |

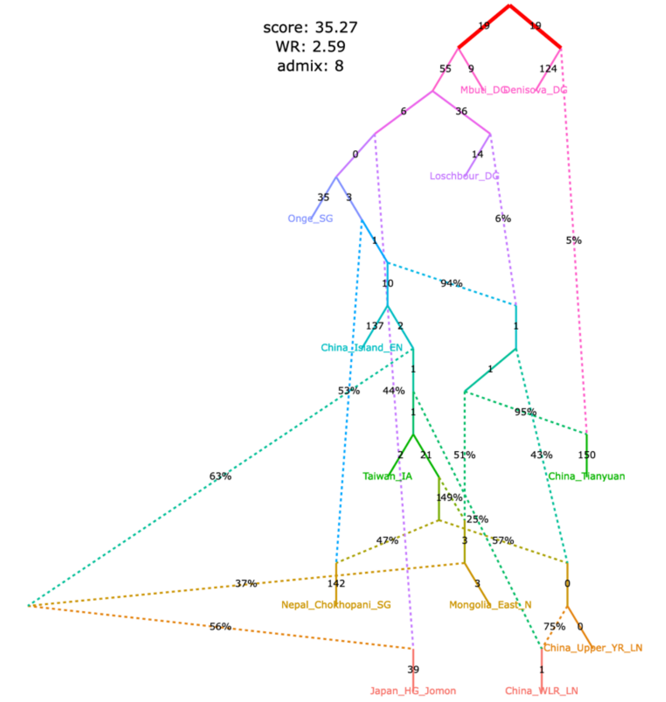

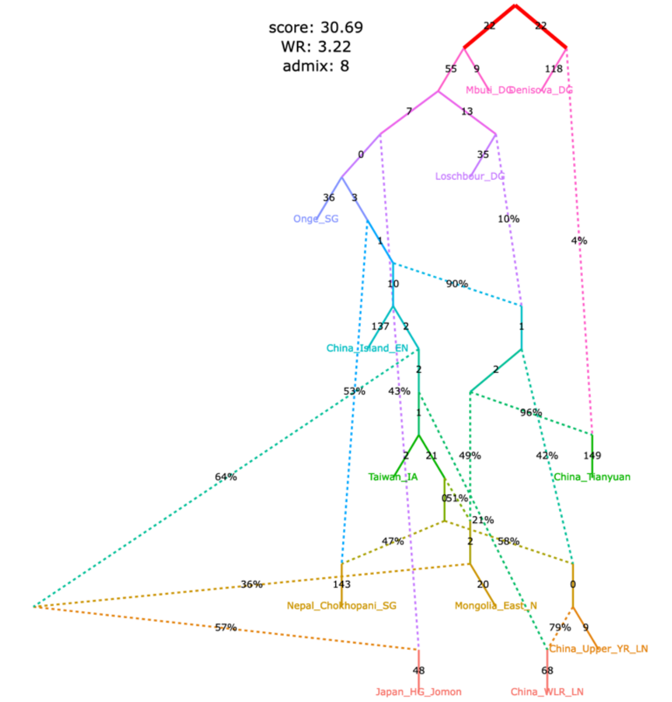

Figure S12

**Figure S12.** Simplified published admixture graph for West Eurasian groups from Sikora *et al.* (2019) and alternative graphs found with *findGraphs* (13 populations, 6 admixture events).

**a**, published model, with chimpanzee added as an outgroup and 4 low-level gene flows from the Neanderthal lineage dropped

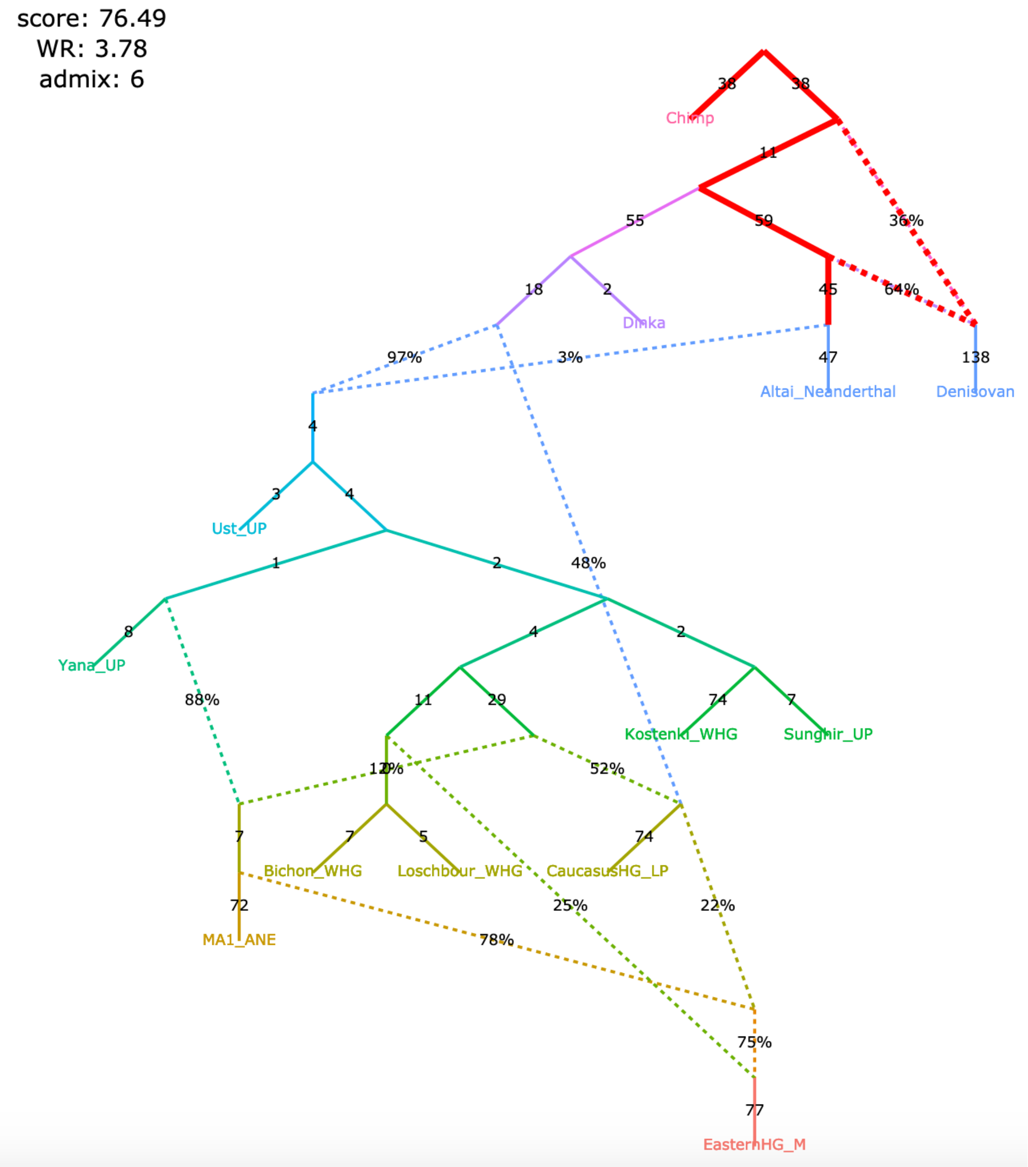

A claim by Sikora et al. 2019 relying on the admixture graph above:

The Mal'ta (MA1_ANE) lineage receives a gene flow from the Caucasus hunter-gatherer (CaucasusHG_LP) lineage.

Figure S12

**b**, models fitting nominally better than the published one and not supporting the claim

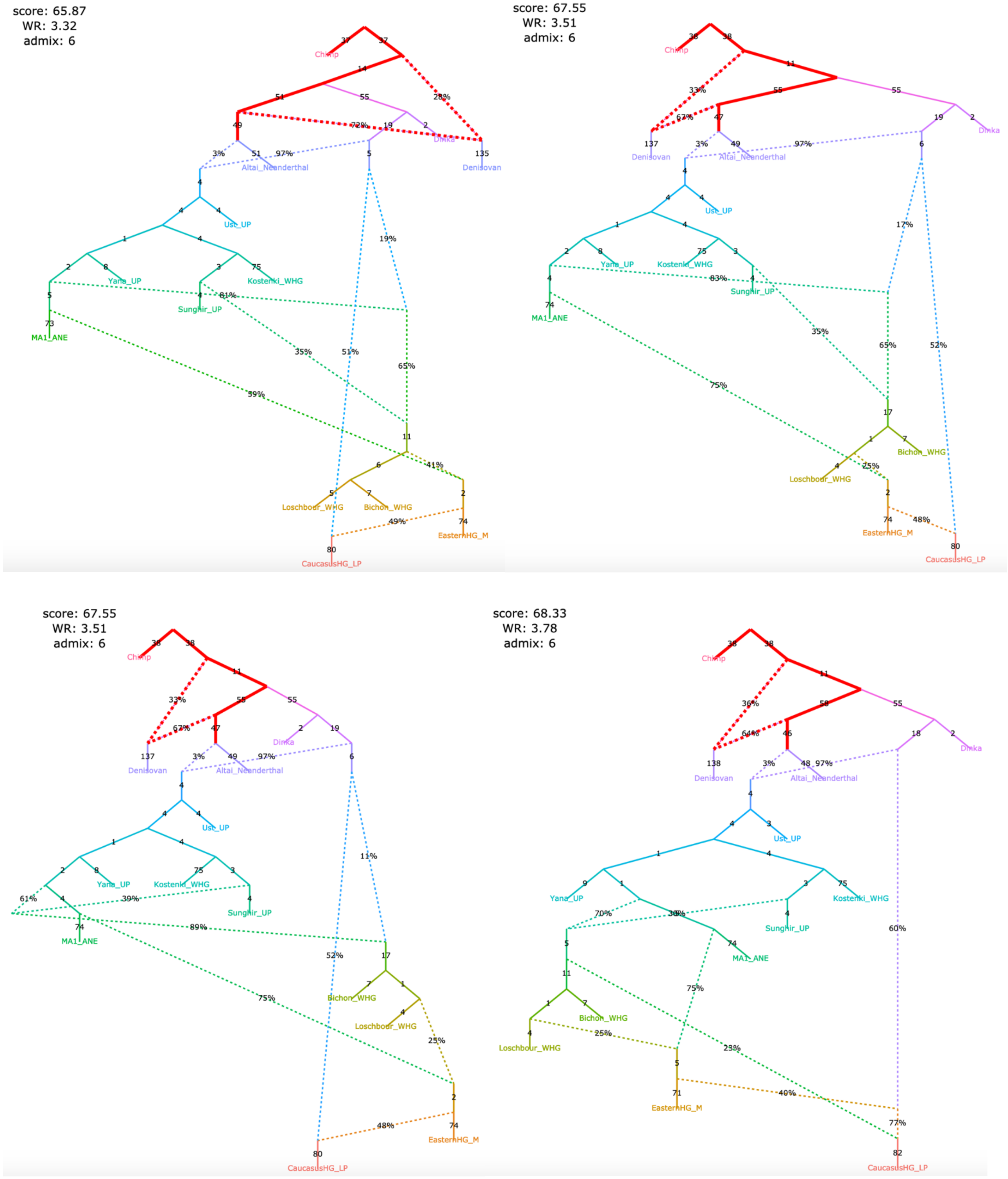

Figure S12

**c**, models fitting nominally better than the published one and not supporting the claim

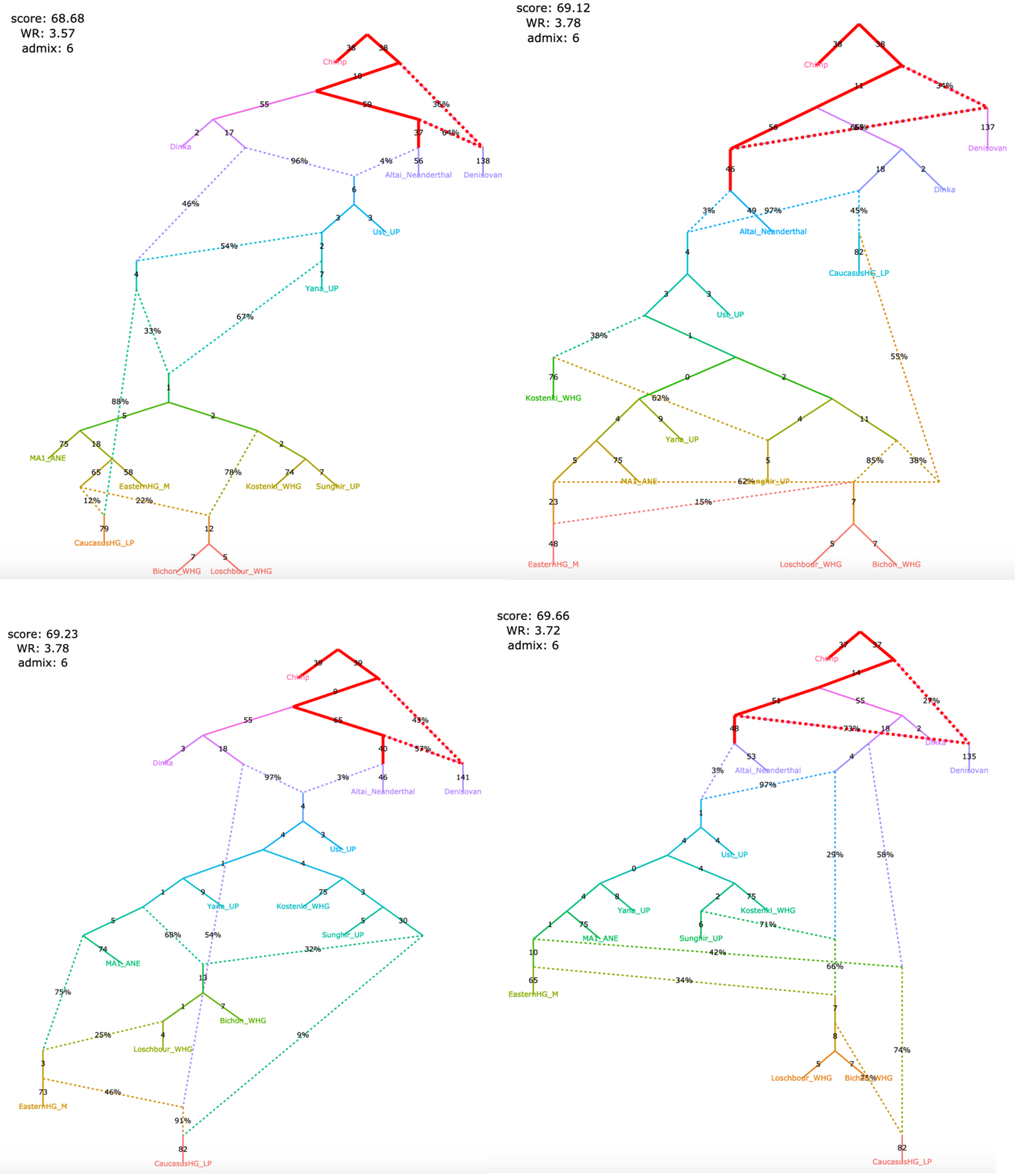

Figure S12

**d**, models fitting nominally better than the published one and not supporting the claim

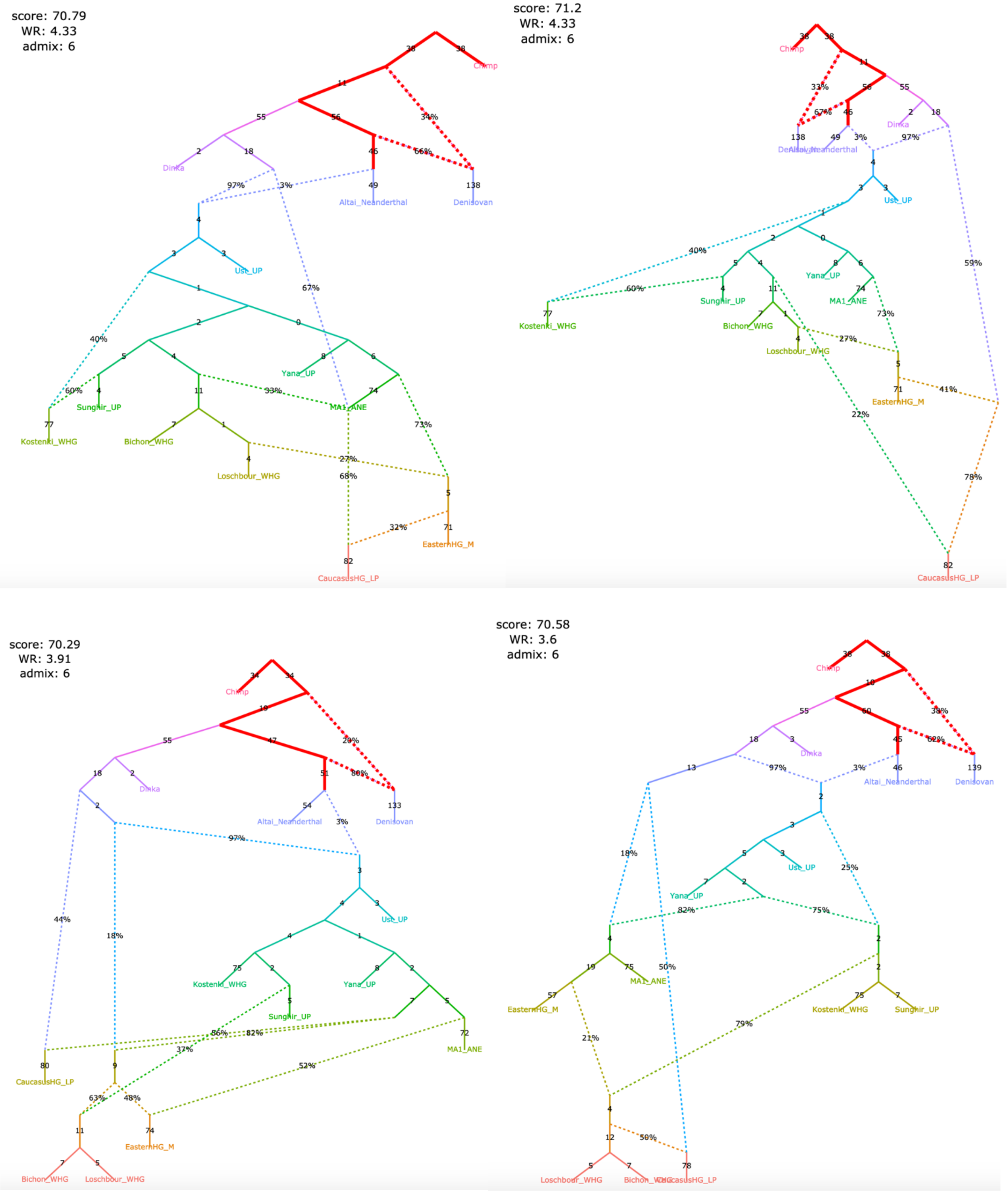

Figure S12

**e**, models fitting nominally better than the published one and not supporting the claim

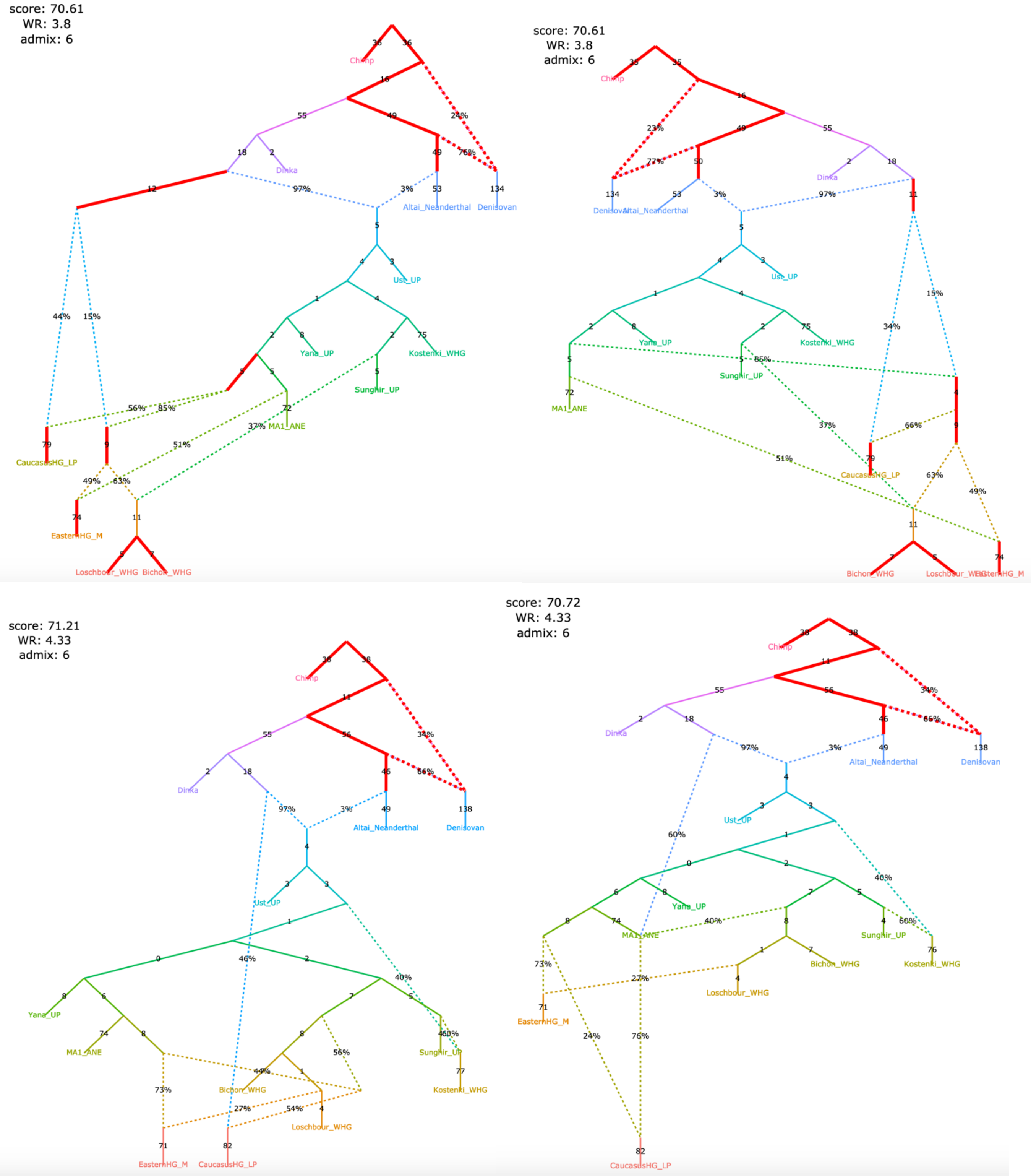

Figure S12

**f**, models fitting nominally better than the published one and not supporting the claim

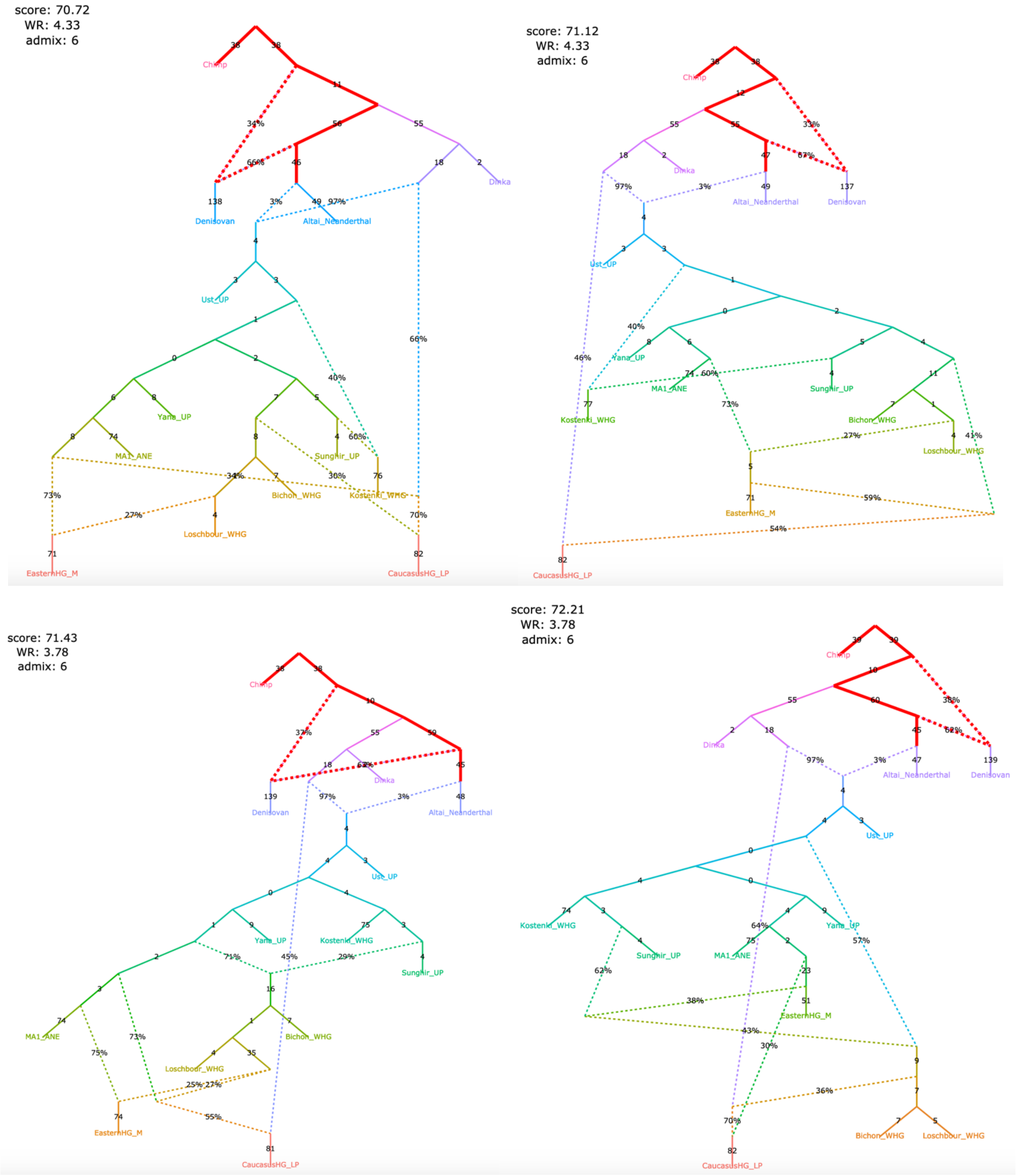

Figure S12

**g**, models fitting nominally better than the published one and not supporting the claim

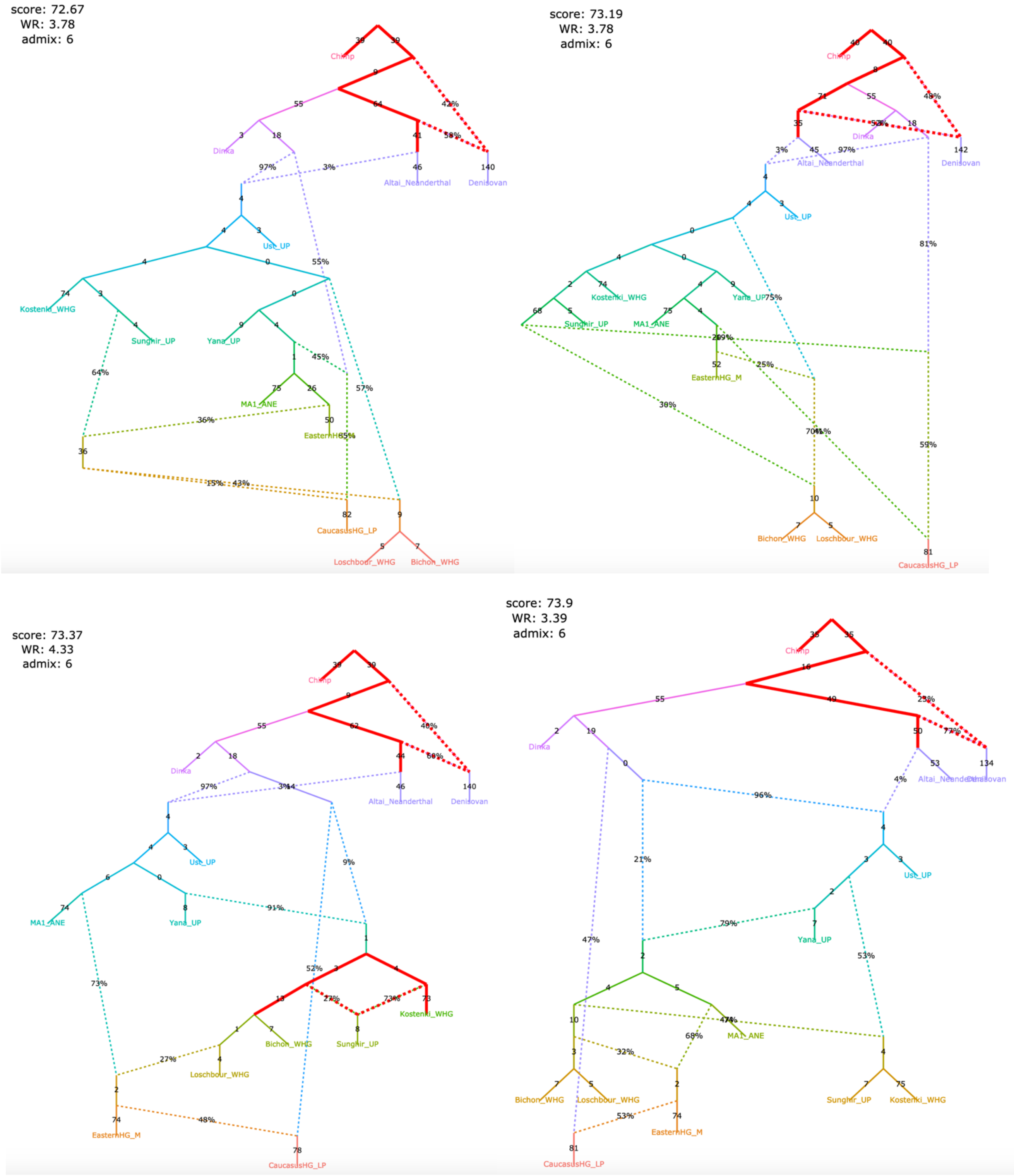

Figure S12

**h**, models fitting nominally better than the published one and not supporting the claim

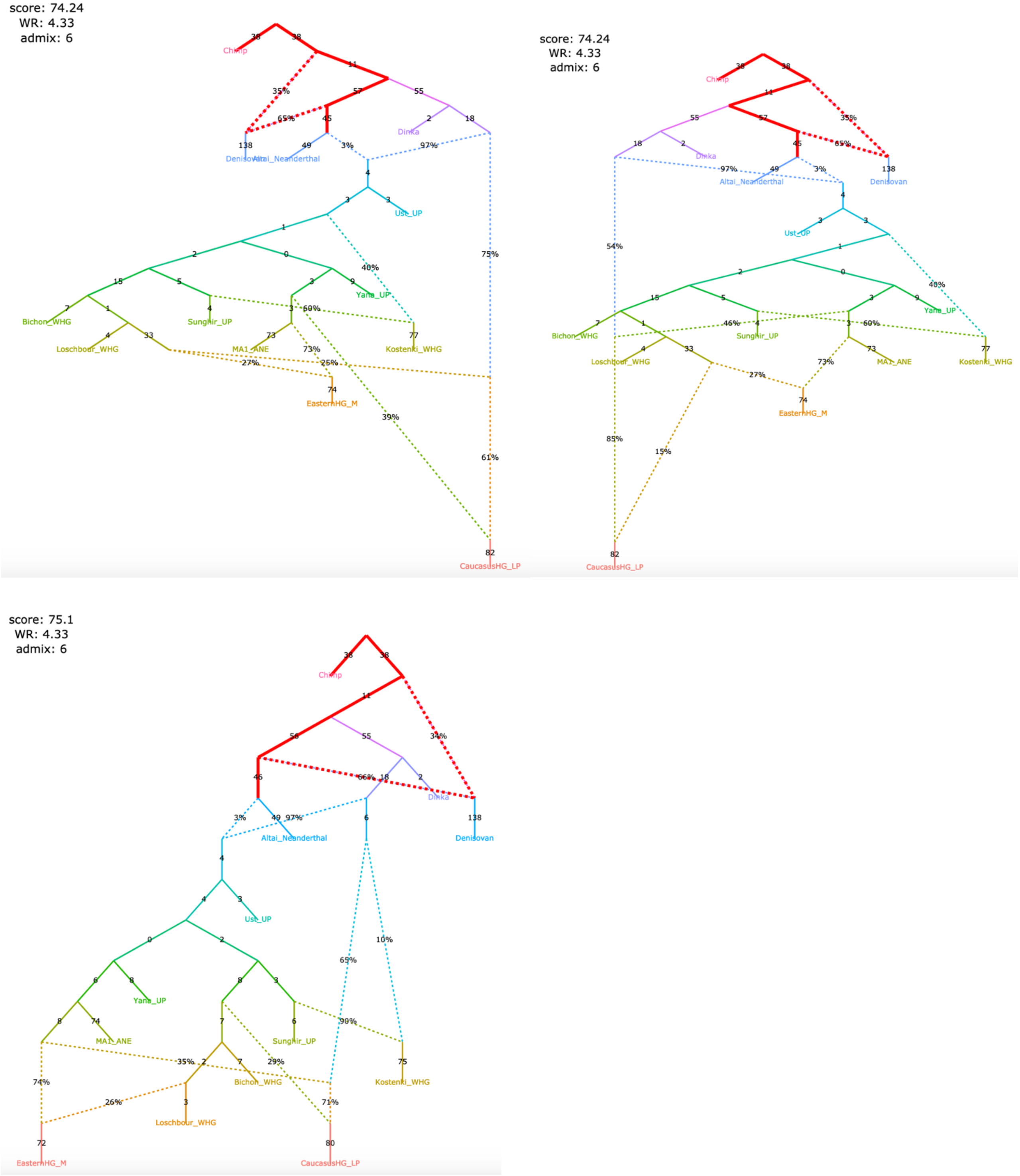

Figure S12

**i**, models fitting nominally better than the published one and <u>supporting</u> the claim

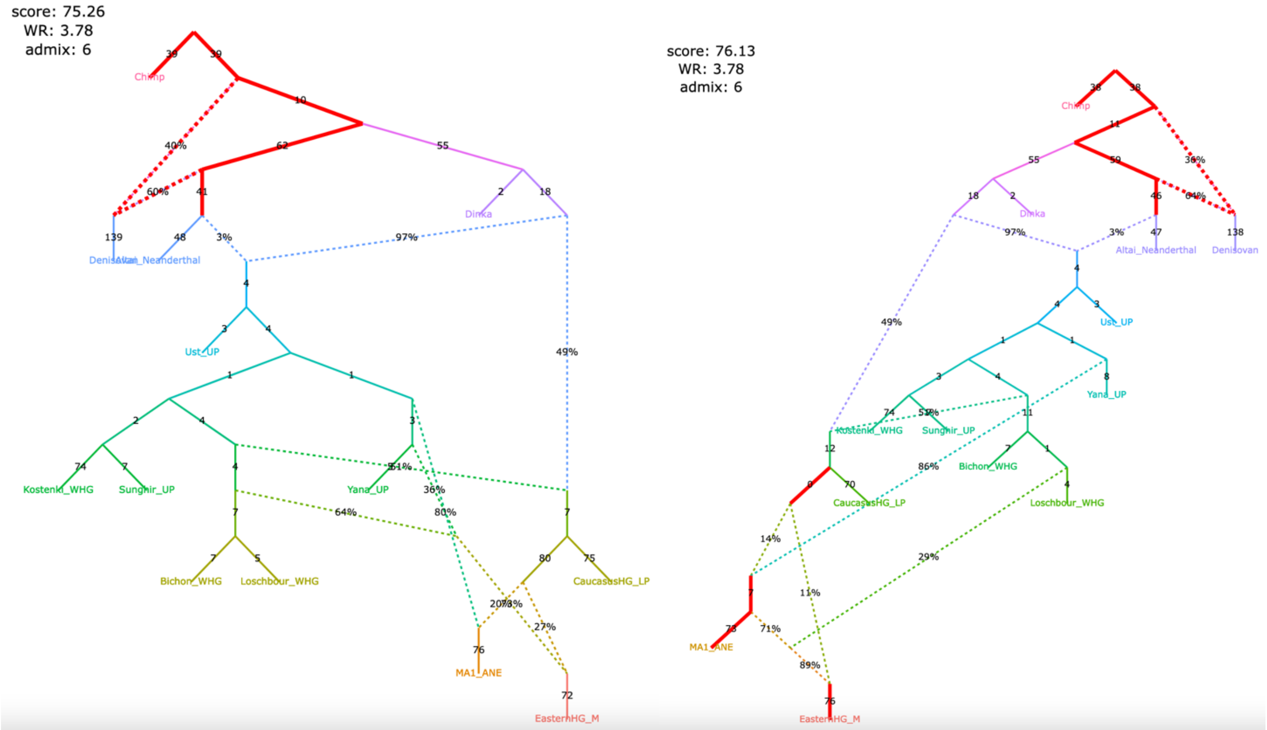

Figure S13

**Figure S13.** Simplified published admixture graph for East Eurasian groups from Sikora *et al*. (2019) and alternative graphs found with *findGraphs* (14 populations, 6 admixture events).

**a**, published model, with chimpanzee added as an outgroup and 4 low-level gene flows from the Neanderthal lineage dropped. **b**, the same model as above, simplified by dropping unidentifiable edges

**a**

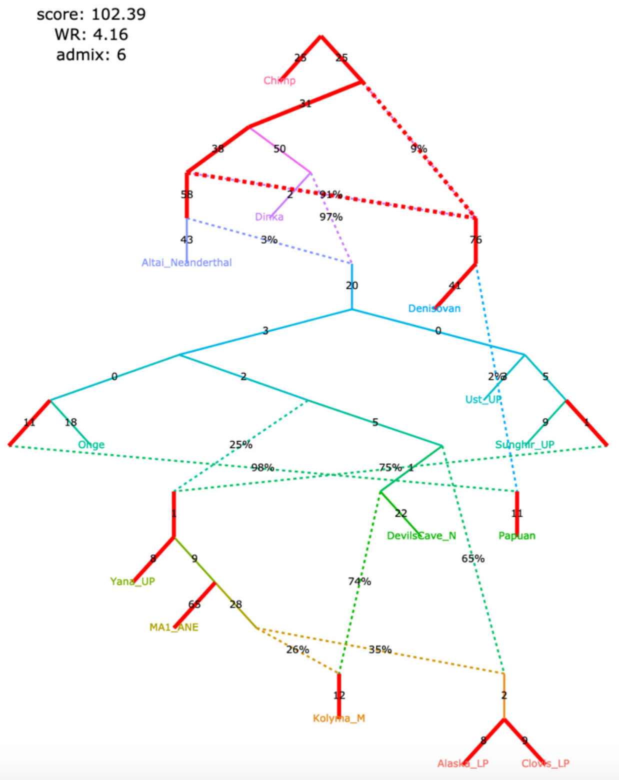

**b**

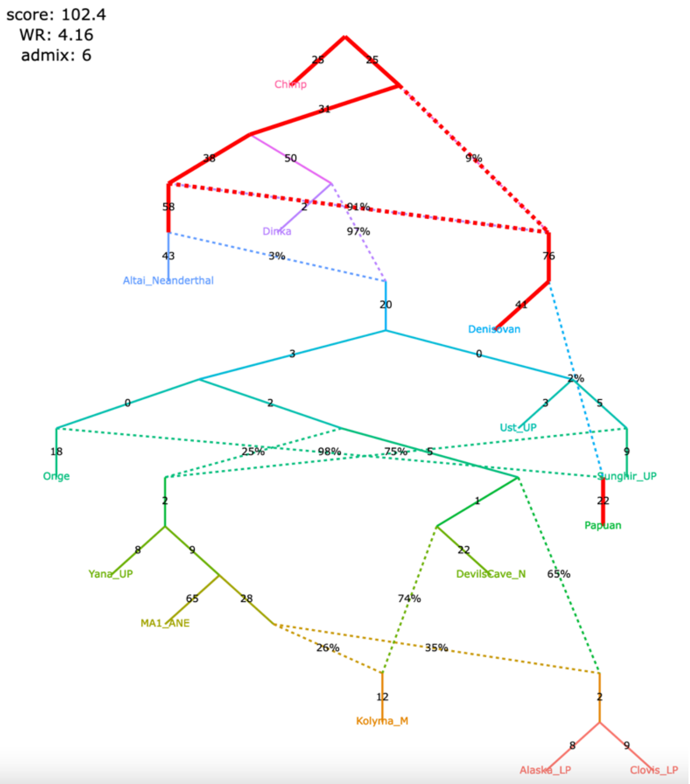

Claims by Sikora et al. 2019 relying on the admixture graph above:

1) The Mal'ta (MA1_ANE) and Yana (Yana_UP) lineages receive a gene flow from an Asian source diverging before the Devil's Cave (DevilsCave_N), Kolyma (Kolyma_M), USR1 (Alaska_LP), and Clovis (Clovis_LP) lineages;
2) European ancestry in the Kolyma, USR1, and Clovis lineages is closer to Mal'ta than to Yana;
3) The Devil's Cave lineage receives no European-related gene flows, and Kolyma has less European-related ancestry than ancient Americans (USR1 and Clovis).

Figure S13

**c**, an alternative model fitting nominally better than the published one and supporting all 3 claims

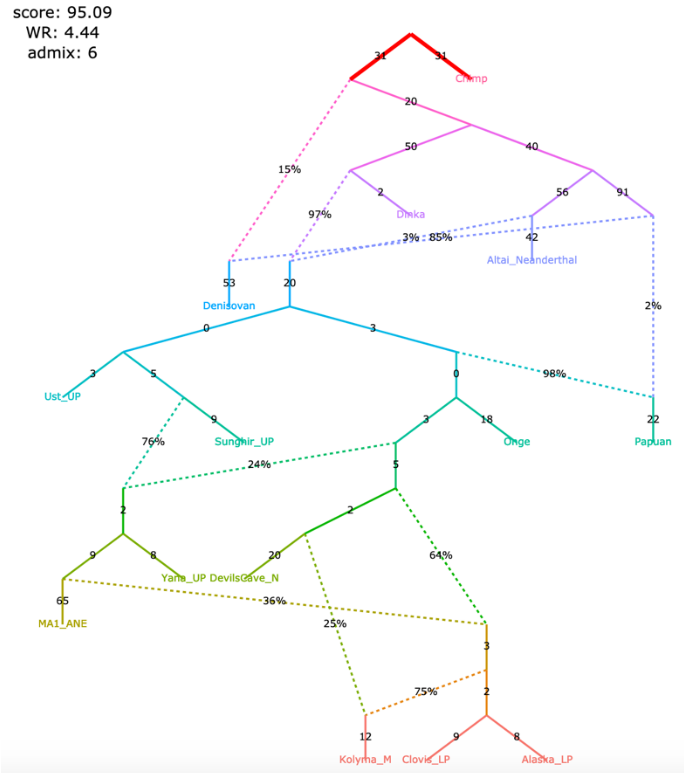

Figure S13

**d**, alternative models fitting not significantly worse than the published model

claims 1&3 not supported, claim 2 supported

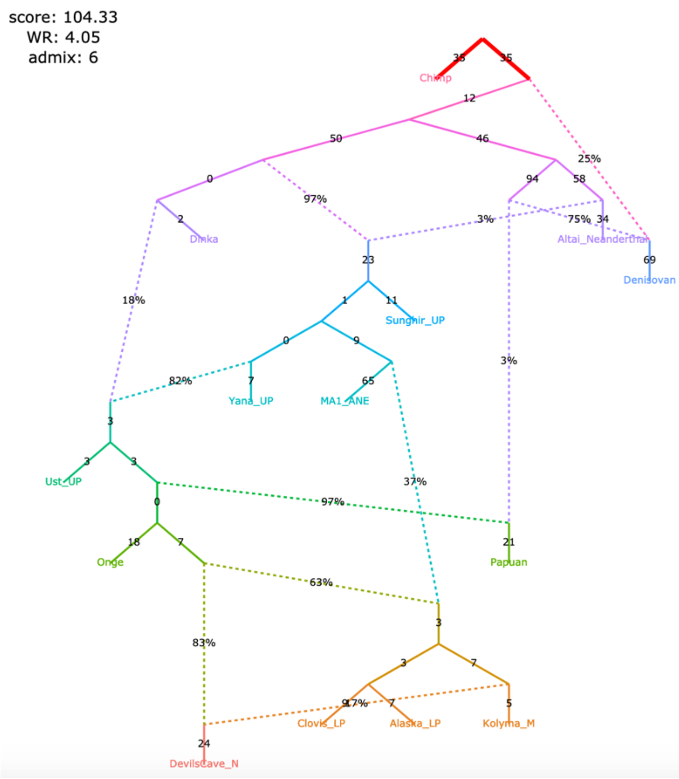

claims 1 not supported, claims 2&3 supported

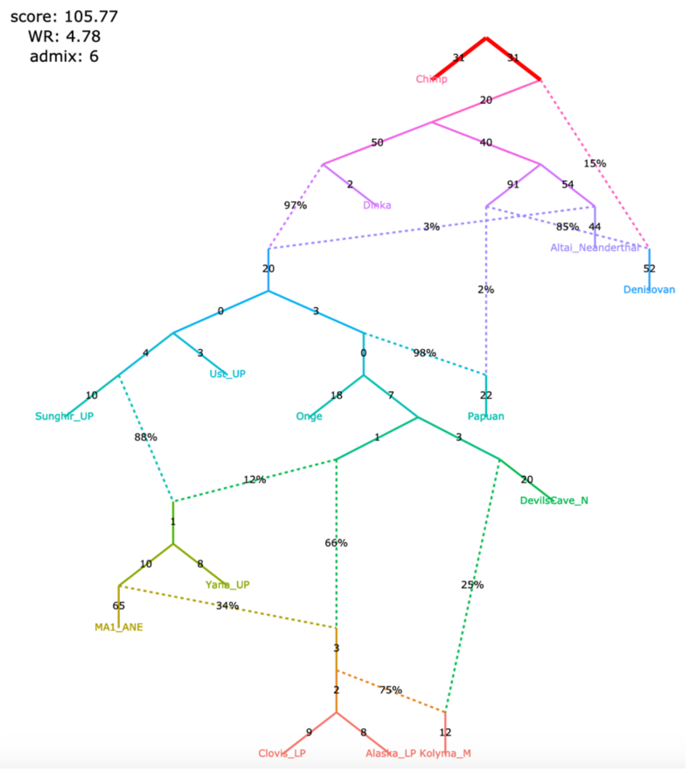

claim 1 not supported, claims 2&3 supported

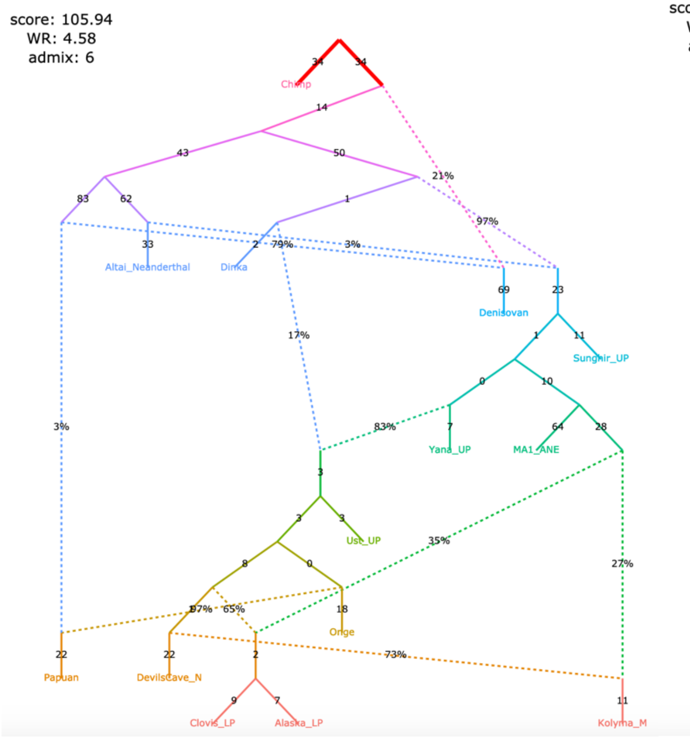

claims 1&2 supported, claim 3 not supported

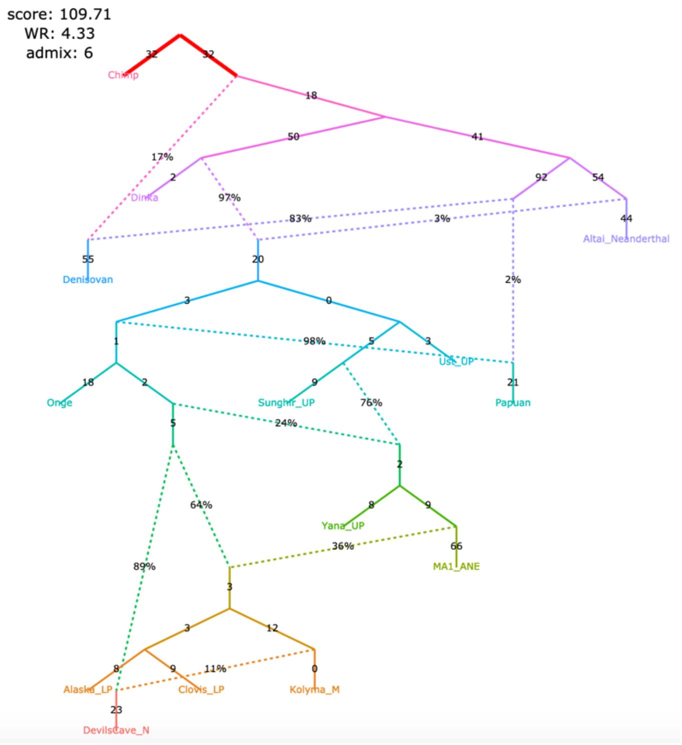

Figure S13

**e**, alternative models fitting not significantly worse than the published model

claim 1 not supported, claims 2&3 supported

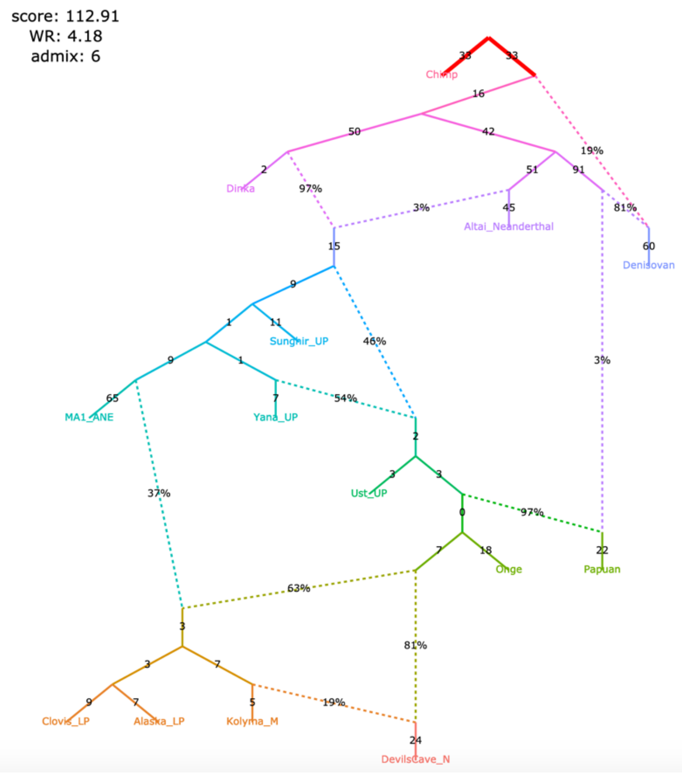

claims 1&3 not supported, claim 2 supported

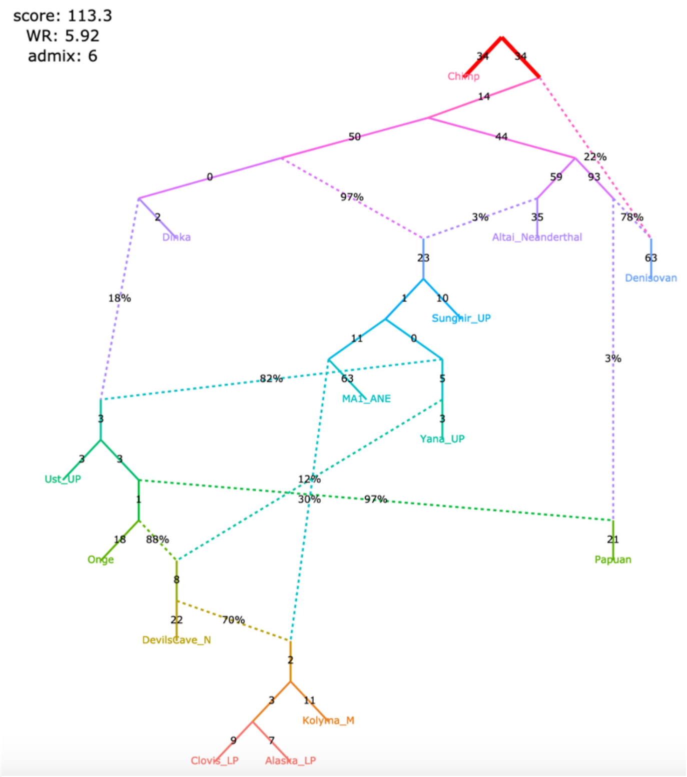

claim 1 not supported, claims 2&3 supported

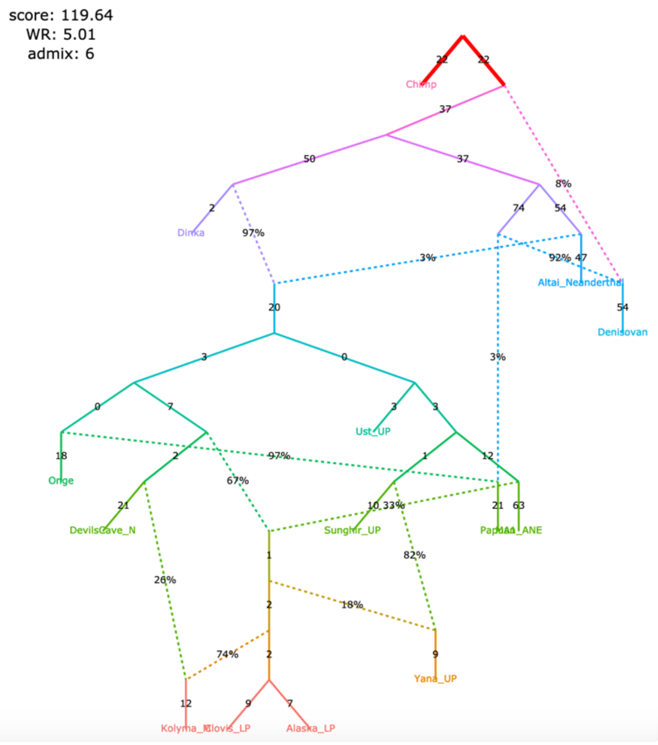

claim 1 not supported, claims 2&3 supported

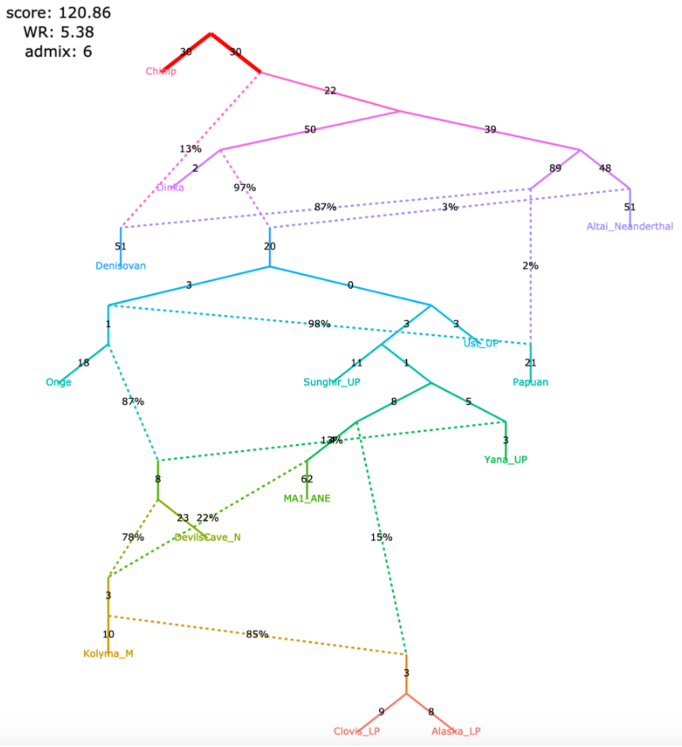

Figure S13

**f**, alternative models fitting not significantly worse than the published model

claims 1&3 not supported, claim 2 supported

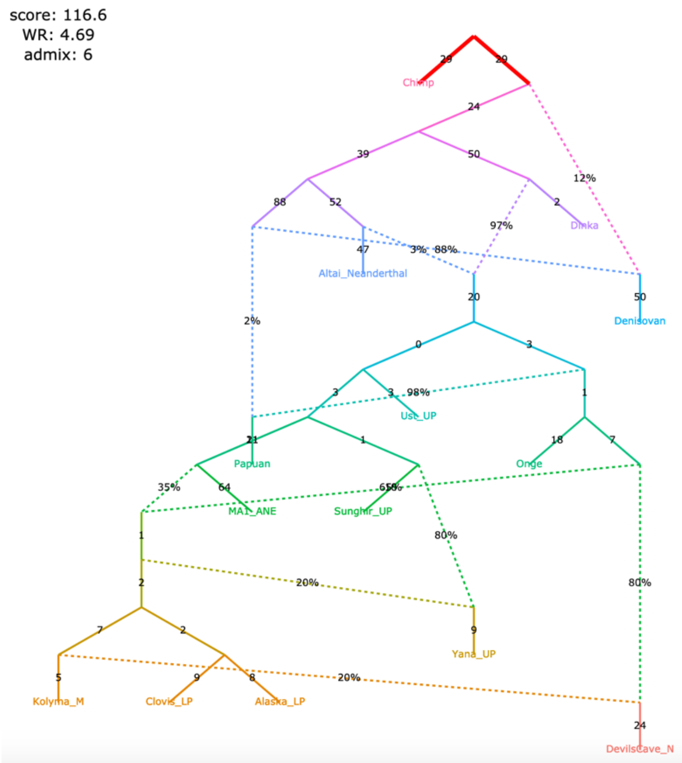

claims 1&3 not supported, claim 2 supported

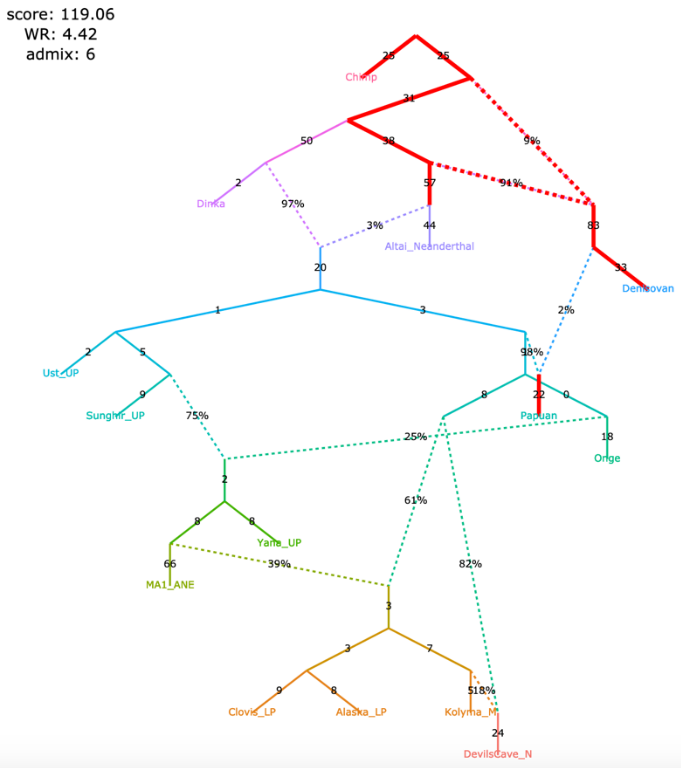

claims 1&2 supported, claim 3 not supported

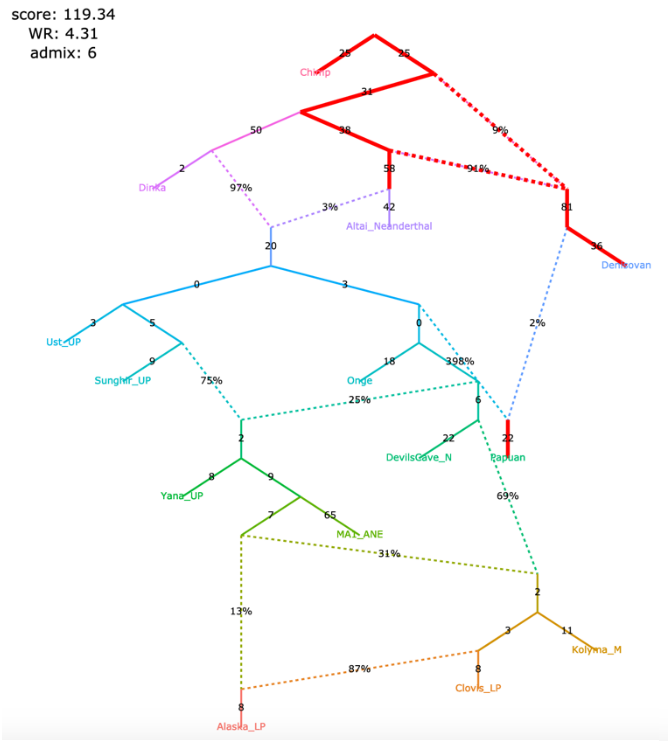

claims 1&2 supported, claim 3 not supported

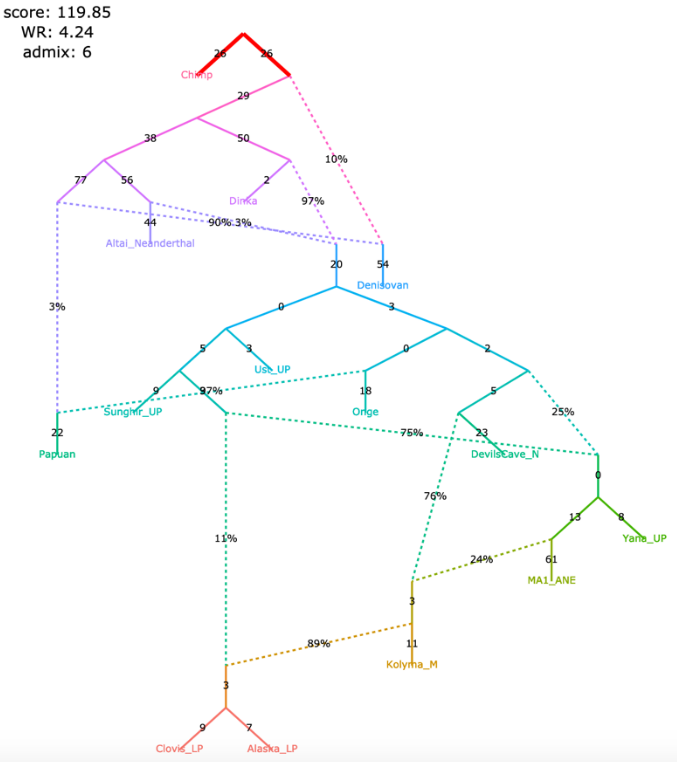

Figure S13

**g**, alternative models fitting not significantly worse than the published model

claims 1&3 not supported, claim 2 supported

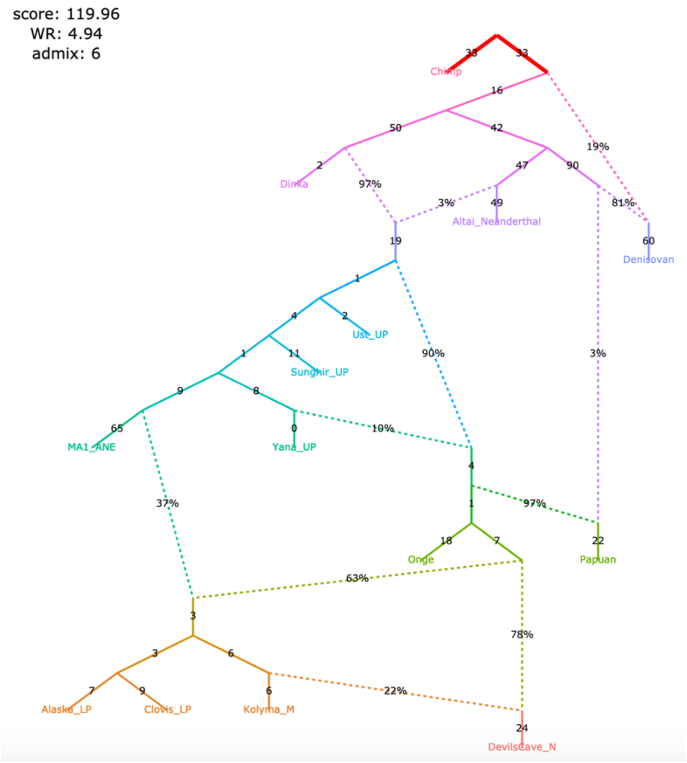
