## Supplementary material for "On the limits of fitting complex models of population history to genetic data": Source file for main manuscript (without rasterization that happens in the PDF)

**Figure 1**

**a**

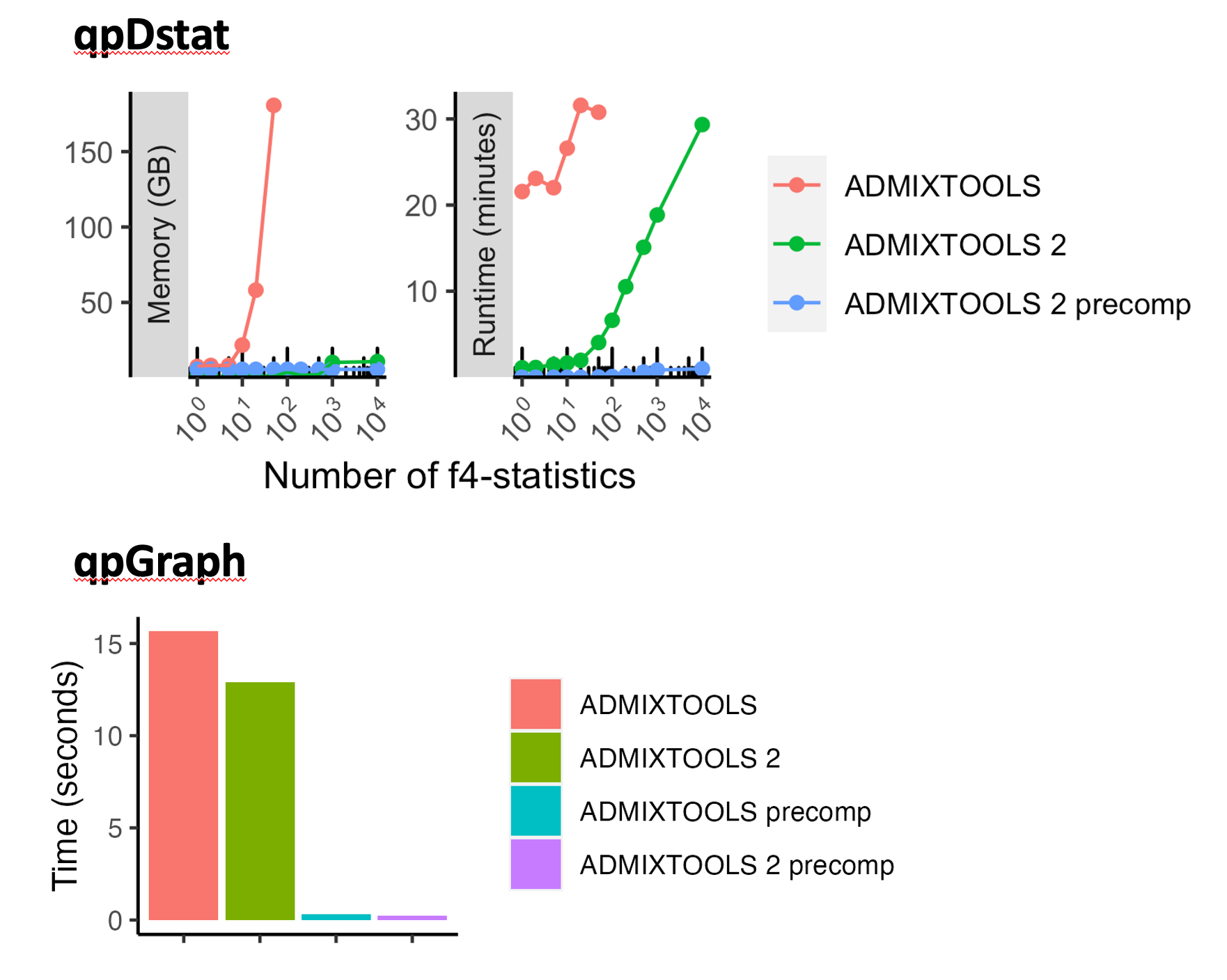

**b**

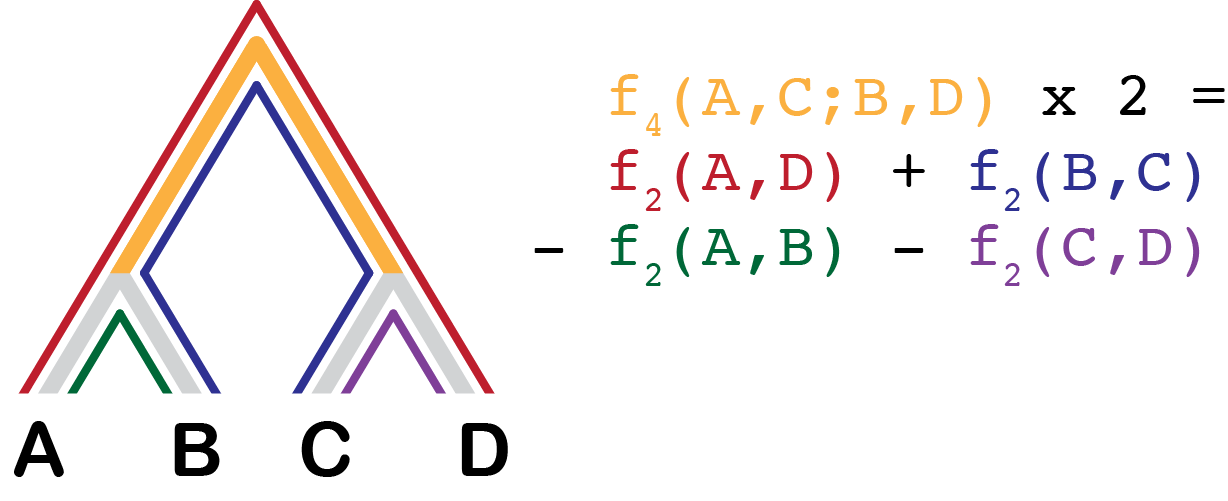

**c**

| **Program** | **f-statistic** | **Primary use** |
| --- | --- | --- |
| qp3pop | *f_3_* | Test for admixture |
| qpDstat | *f_4_* | Test for admixture |
| qpF4ratio | *f_4_* | Estimating admixture proportions |
| qpWave | *f_4_* | Finding how many gene flows connect two sets of populations |
| qpAdm | *f_4_* | Estimating admixture proportions |
| qpGraph | *f_3_* | Fitting admixture graph models of history |

**d**

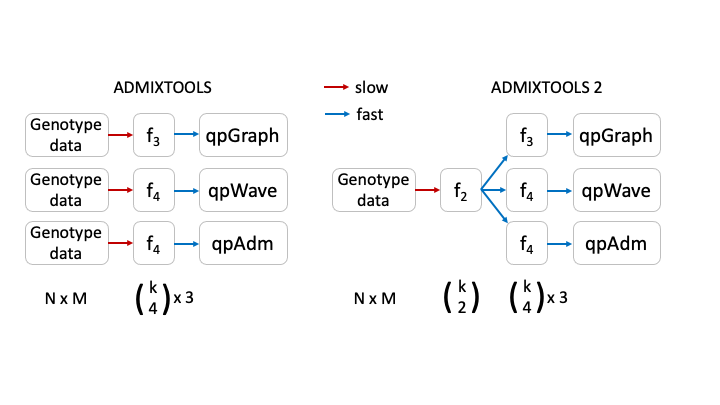

**Figure 1:**

**a** Performance comparison of *f*-statistic computation and admixture graph fitting. Top: Memory usage and runtime for computing *f*-statistics using (1) the *qpDstat* program in *ADMIXTOOLS* v7.0.2 released in 06/2021 (2) the $f_{4}$ function in *ADMIXTOOLS 2* without precomputing $f_{2}$-statistics, and (3) the $f_{4}$ function in *ADMIXTOOLS 2* with precomputed $f_{2}$-statistics. (1) and (2) give identical results, whereas (3) only gives identical results in the absence of missing data, which limits usefulness beyond a moderate number of populations. Bottom: Runtime comparison of *qpGraph* with and without precomputed *f*-statistics.

**b** Illustration of $f_{2}$-and $f_{4}$-statistics. $f_{2}$ measures the amount of drift separating any two populations, while $f_{4}$ measures the amount of drift shared between two population pairs. Every *f_4_*-statistic is a linear combination of four *f_2_*-statistics.

**c** Overview of the major *ADMIXTOOLS* programs, their primary use cases, and their associated *f*-statistics.

**d** Schematic representation of the computations behind the *ADMIXTOOLS* programs *qpGraph*, *qpWave*, and *qpAdm*. *ADMIXTOOLS 2* separates the computation of $f_{2}$-statistics from the later steps in the pipeline. Shown below are the number of data points for $N$ individuals, $M$ SNPs, and $k$populations. The exact number of all possible non-redundant $f_{2}$, $f_{3}$, and $f_{4}$-statistics for $k$ populations are $\binom{k}{2}$, $\frac{1}{2}\binom{k}{3}$, and $\frac{1}{3}\binom{k}{4}$. A small number of $f_{2}$-statistics can be used to obtain a much larger number of $f_{3}$- and $f_{4}$-statistics and requires much less space than the raw genotype data.

**a**

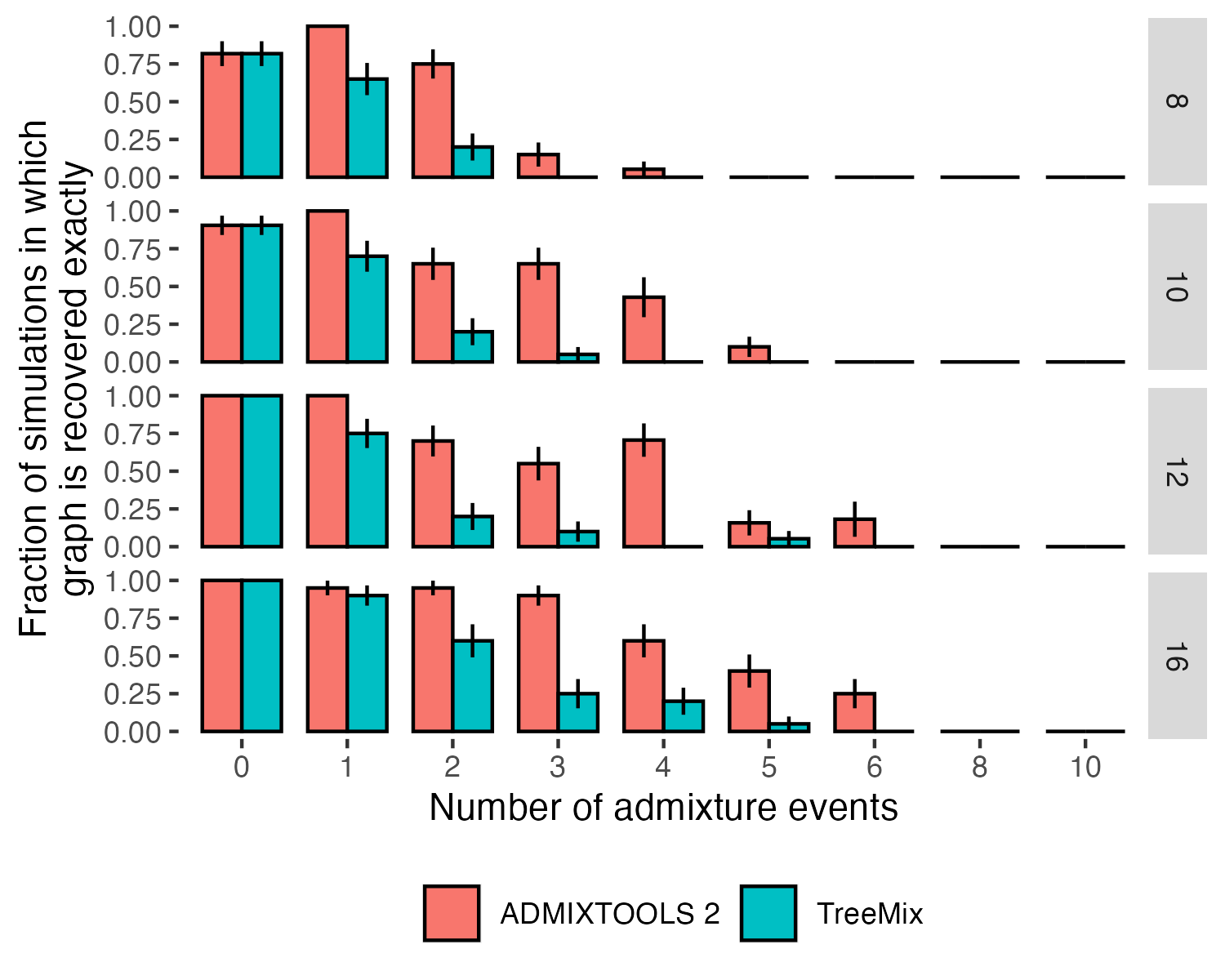

**b**

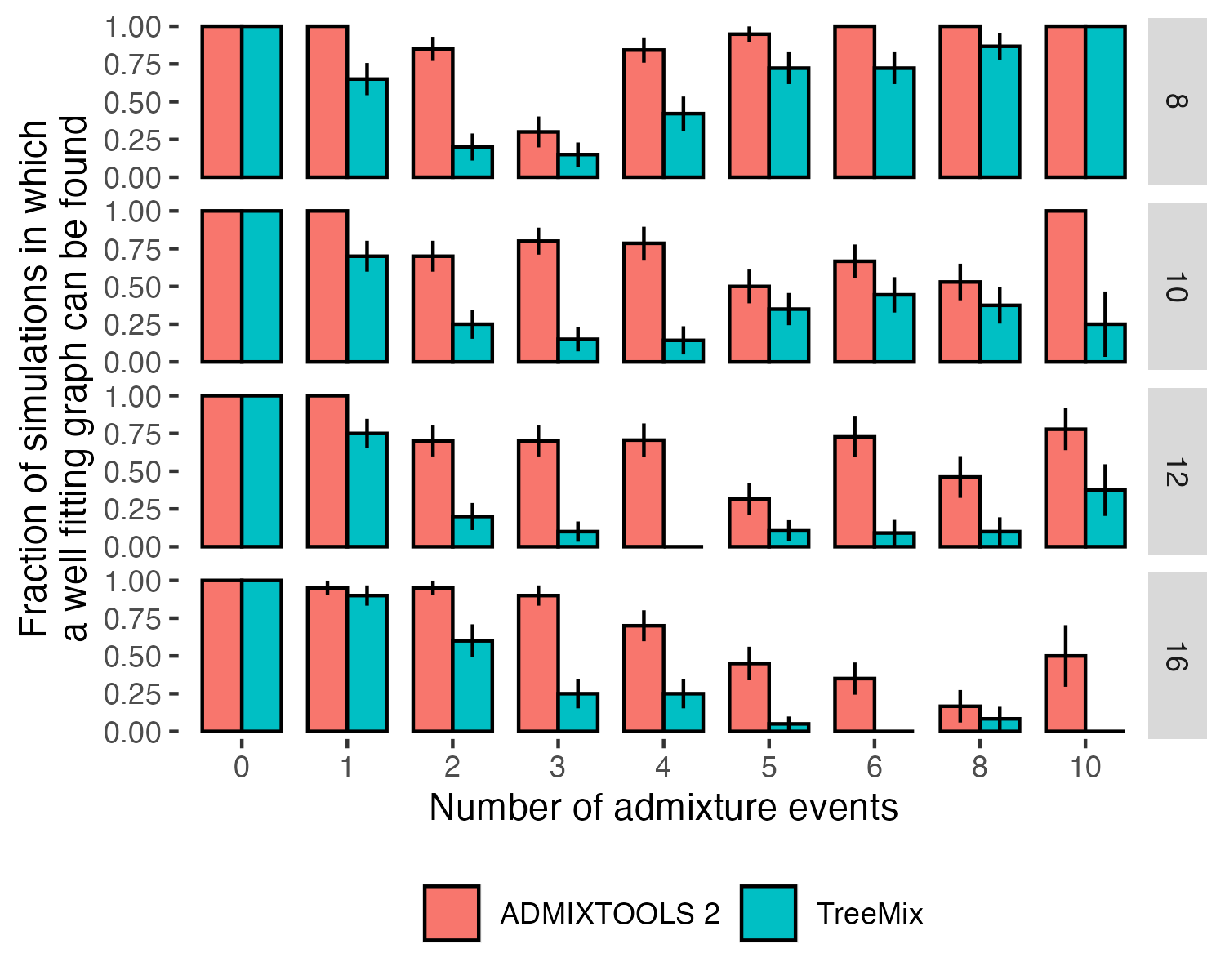

**Figure 2:**

Comparison of accuracy of automated search for optimal topology in the *findGraphs* function of *ADMIXTOOLS 2* and *TreeMix* using simulated graphs with 8, 10, 12, and 16 populations, and 0 to 10 admixture events. Error bars show standard errors calculated as ${SE^{2}=p(1-p)/n}$ where $p$ is the fraction on the y-axis and $n$ is the number of simulations in each group (typically 20). In the case of *ADMIXTOOLS 2*, we applied *findGraphs* three times on each simulated data set and picked a result with the best fit score. More details are provided in the Methods section. **a** Fraction of simulations where the simulated graph is recovered exactly. **b** Fraction of simulations where the simulated graph is either recovered exactly, or the score is at least as good as the score of the simulated graph, when both graphs are evaluated by *ADMIXTOOLS 2*. More admixture edges greatly increase the search space and make it more difficult to recover the simulated graph, but they do make it easier to find alternative graphs with good fits.

A second challenge in comparing different admixture graph models arises when comparing models of different complexity (i.e., with a different number of admixture events). Established methods such as AIC and BIC are applicable and also can account for different model complexity if the number of independent parameters in a model is known. However, the number of independent parameters estimated in an admixture graph is not simply determined by the number of groups, drift edges, or admixture events, as it also depends on the graph topology in a complex way. We implement a method to compare admixture graph models of different complexity by using a new scoring function, which uses different blocks of SNPs for deriving fitted and estimated *f*-statistics. This ensures that our model comparison test does not favor more complex models by allowing them to overfit the data. This cross-validation approach can also be used to rank alternative models of the same complexity and deal with overfitting. We note that the calculation of cross-validated log-likelihood scores is not turned on in *findGraphs* by default, and to make our results more comparable to those of the published studies we revisited, we relied on standard log-likelihood scores below.

To test if our method is well calibrated, we simulated 100,000 SNPs under the same graph in 1000 replicates. We then created two new topologies by removing one out of two symmetric edges from the first graph (**Figure 3c**). These new incorrect models are symmetrically related to the first graph and can be used to test the null hypothesis that the true difference in log-likelihood scores of these two graphs is zero. The uniform distribution of *p*-values confirms that our method is well calibrated (**Figure 3d**). A caveat is that only one symmetric topology was explored in this way.

**a, b**

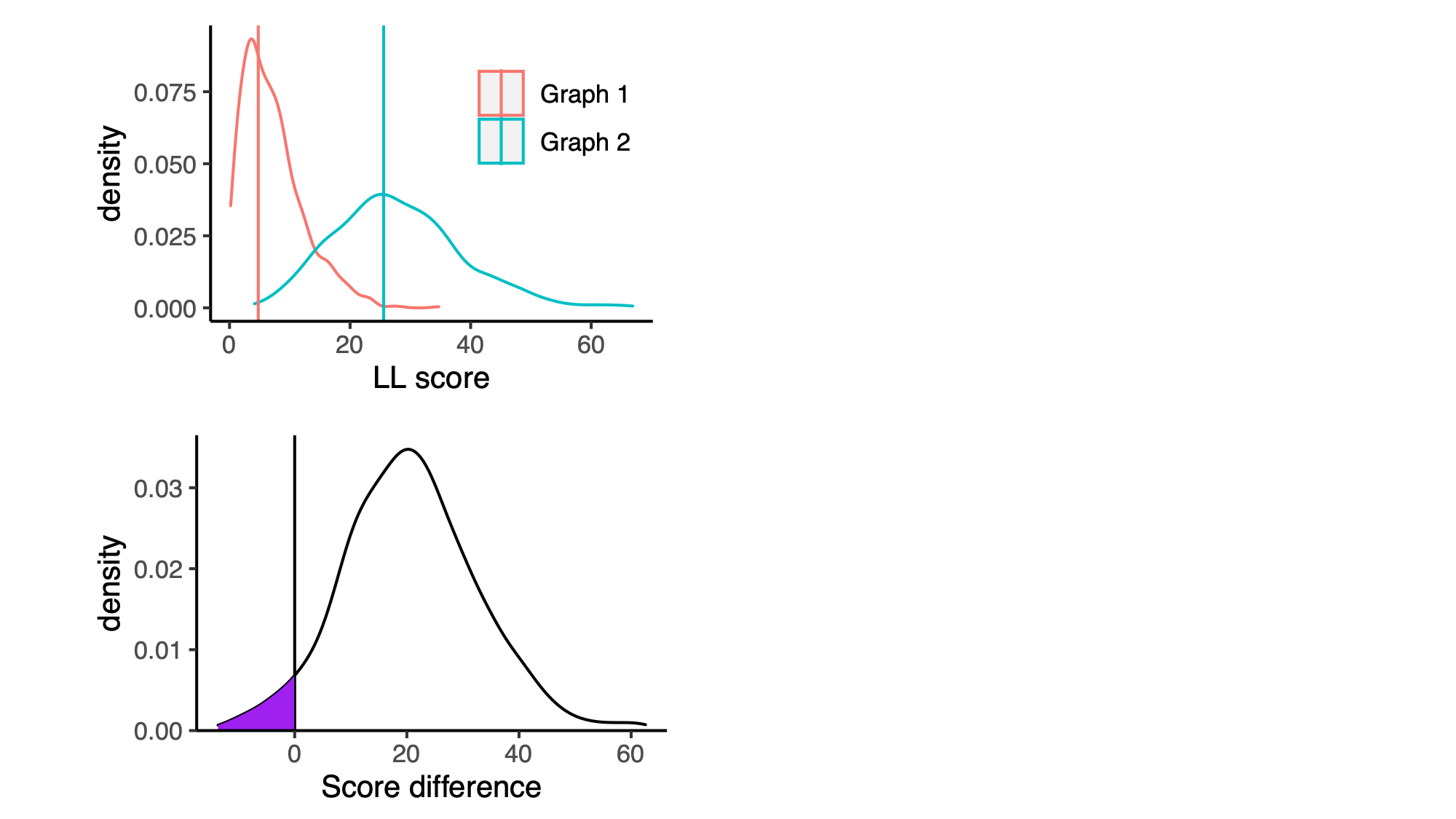

**c**

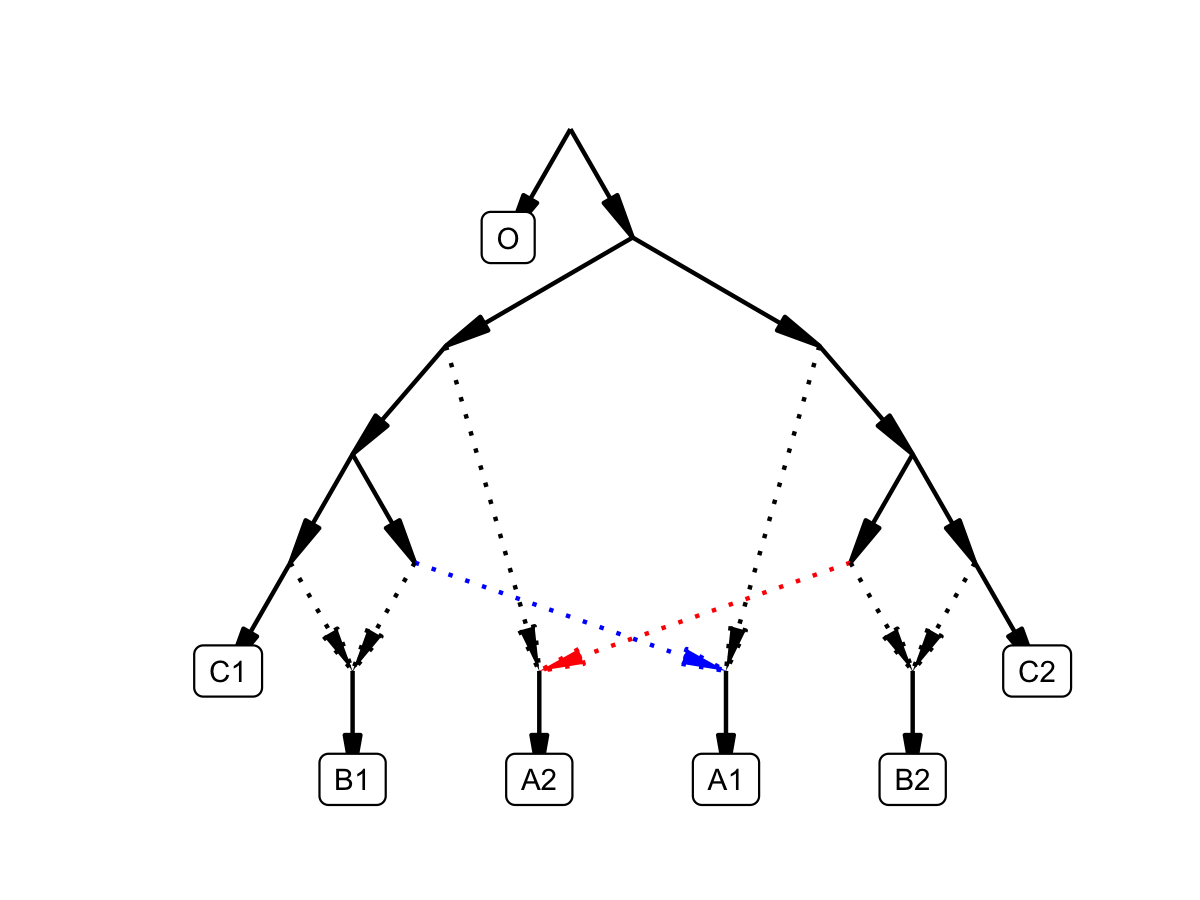

**d**

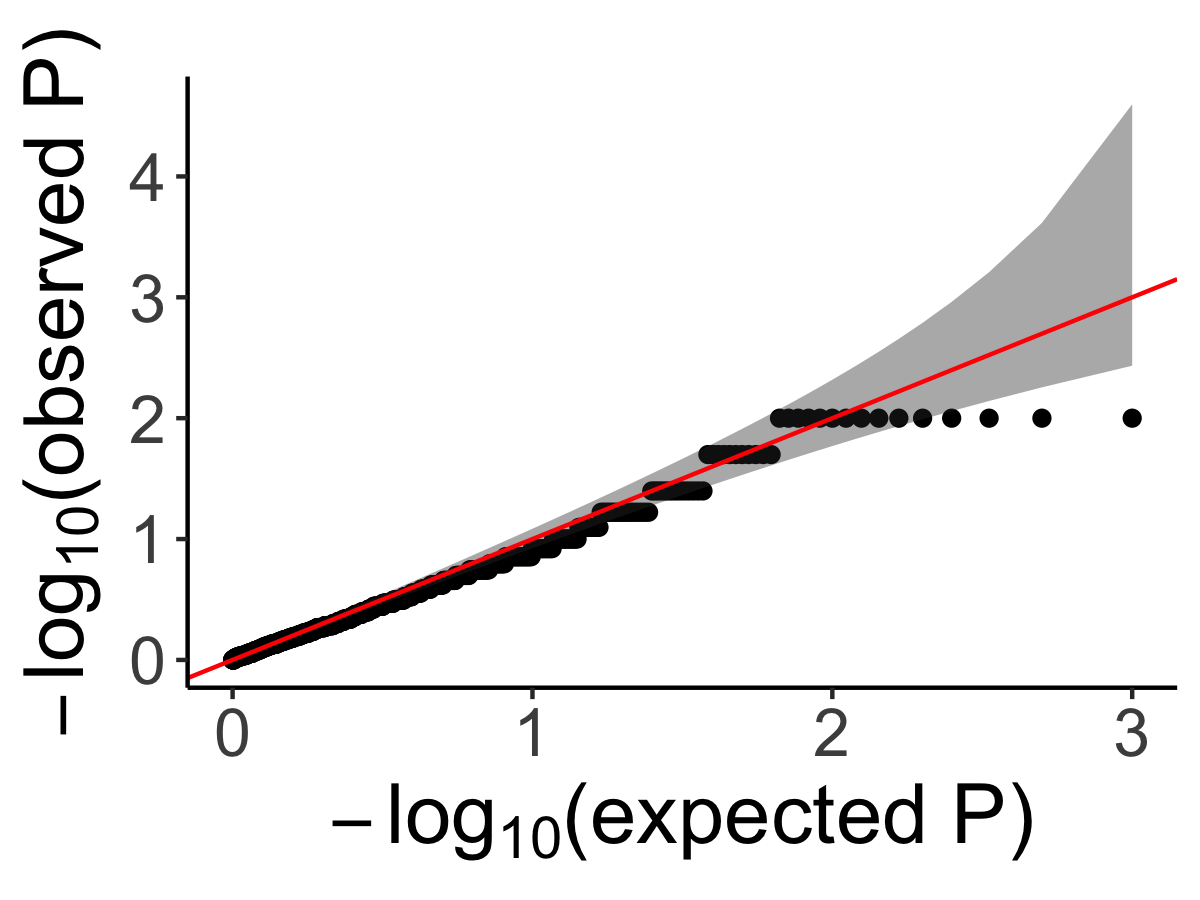

**Figure 3: Calibrating the bootstrap model comparison approach**

**a** Bootstrap sampling distributions of the log-likelihood scores for two admixture graphs (shown in **Figure S1**) of the same populations fitted using real data. Vertical lines show the log-likelihood scores computed on all SNP blocks.

**b** Distribution of differences of the bootstrap log-likelihood scores for both graphs (same data as in **a**). The purple area shows the proportion of resamplings in which the first graph has a higher score than the second graph. The two-sided *p*-value for the null hypothesis of no difference is equivalent to twice that area (or one over the number of bootstrap iterations if all values fall on one side of zero). In this case it is 0.078.

**c** The admixture graph which was used to evaluate our method for testing the significance of the difference of two graph fits on simulated data. We simulated under the full graph and fitted two graphs that result from deleting either the red admixture edge or the blue admixture edge. These two graphs have the same expected *qpGraph* fit score but can have different scores in any one simulation iteration.

**d** QQ plot of *p*-values testing for a score difference between the two graphs (on simulated data) under the null hypothesis of no difference, confirming that the method is well calibrated.

(4) Determining identifiability of admixture graph parameters

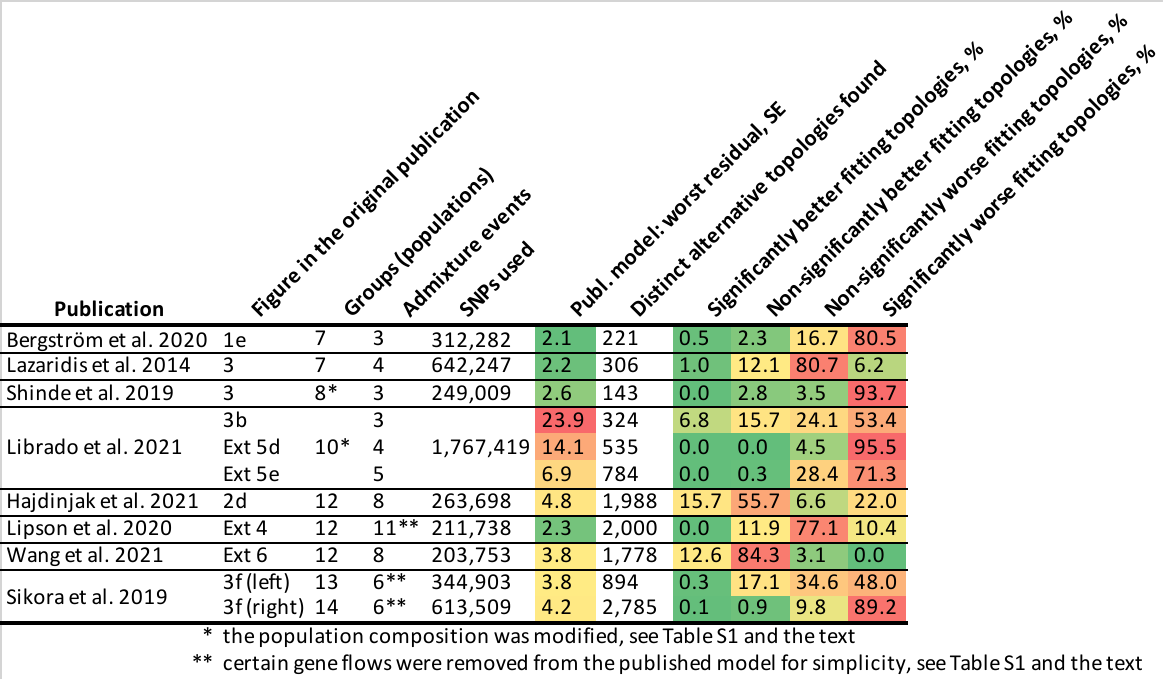

**Table 1: Published graphs in the context of automatically found graphs.**

We compared graphs from 8 publications to alternative graphs inferred on the same or very similar data (see **Table S1** for details).

**Publication**: Last name of the first author and year of the relevant publication.

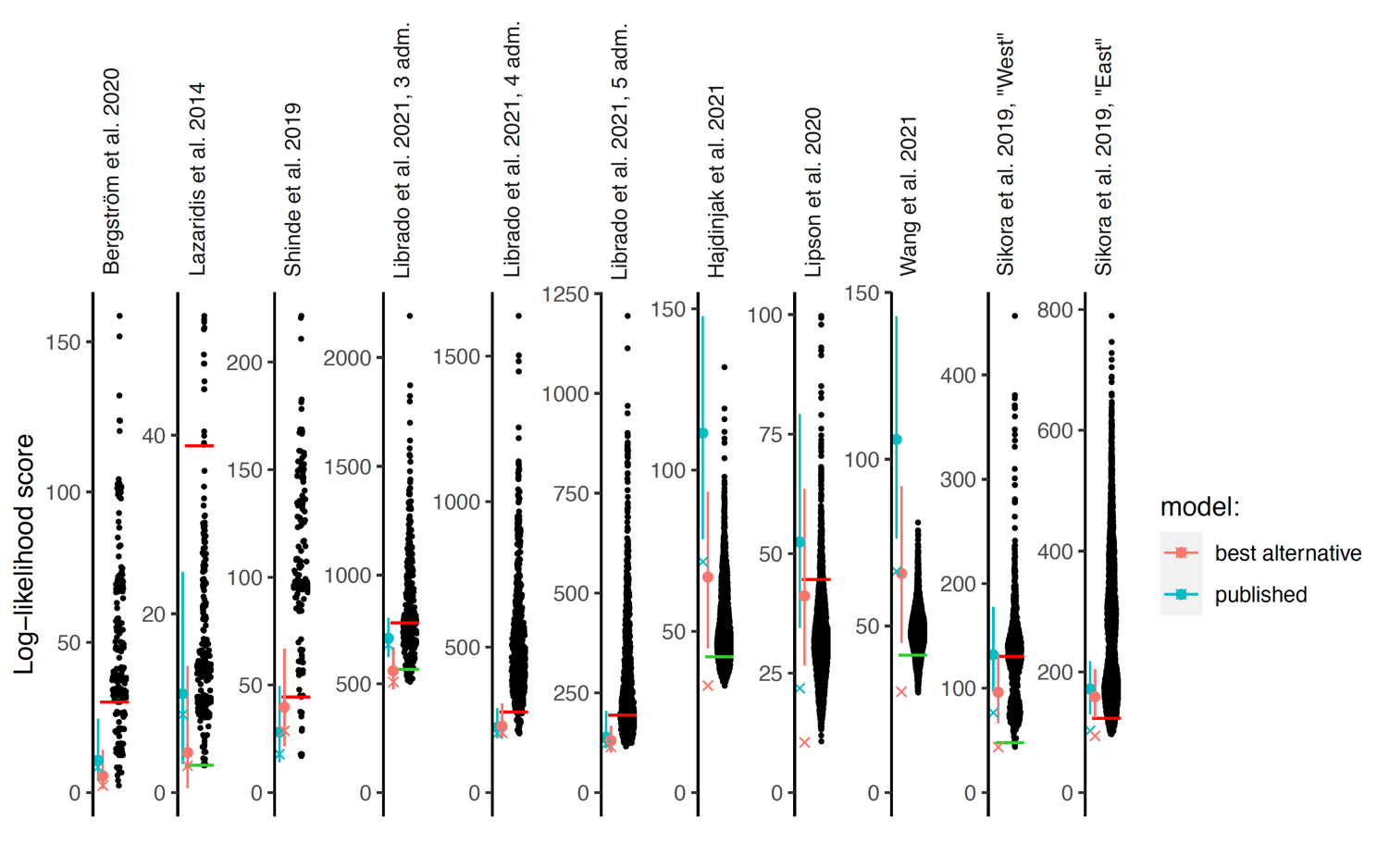

**Figure 4**:

Log-likelihood scores of published graphs (those shown in **Table 1**) and automatically inferred graphs. Each dot represents the log likelihood score of a best-fitting graph from one *findGraphs* iteration (low values of the score indicate a better fit); only topologically distinct graphs are shown. Log-likelihood scores for the published models and best-fitting alternative models found are shown by blue and pink x’s, respectively. Bootstrap distributions of log-likelihood scores for these models (vertical lines, 90% CI) and their medians (solid dots) are also shown. Lower scores of the fits obtained using all SNPs, relative to the bootstrap distribution, indicate overfitting to the full data set. Green and red horizontal lines show the approximate locations where newly found models consistently have fits significantly better or worse, respectively, than those of the published model. In the case of the Bergström *et al*., Lazaridis et al. and Hajdinjak et al. studies, one or more worst-fitting models were removed for improving the visualization. The setups shown here (population composition, number of groups and admixture events, topology search constraints) match those shown in **Table 1**.

**Figure 5**:

Published graphs from six studies for which we explored alternative admixture graph fits. In all cases, we selected a temporally plausible alternative model that fits nominally or significantly better than the published model and has important qualitative differences compared to the published model with respect to the interpretation about population relationships. In all but one case, the model has the same complexity as the published model shown on the left with respect to the number of admixture events; the exception is the reanalysis of Librado *et al*. 2021 horse dataset since the published model with 3 admixture events is a poor fit (worst Z-score comparing the observed and expected *f*-statistics has an absolute value of 23.9 even when changing the composition of the population groups to increase their homogeneity and improve the fit relative to the composition used in the published study). For this case, we show an alternative model with 8 admixture events that fits well and has important qualitative differences from the point of view of population history interpretation relative to the published model. The existence of well-fitting admixture graph models does not mean that the alternative models are the correct models; however, their identification is important because they prove that alternative reasonable scenarios exist to published models.

**a,** Bergström *et al*. 2020 published	Nominally better fitting graph for dogs that is more congruent to human history. For both species, Baikal and Native American groups are mixed between European- and East Asian-related lineages, and a “Basal Eurasian” lineage contributes to Near Eastern groups; these features are all characteristic of human history but absent in the published dog graph.

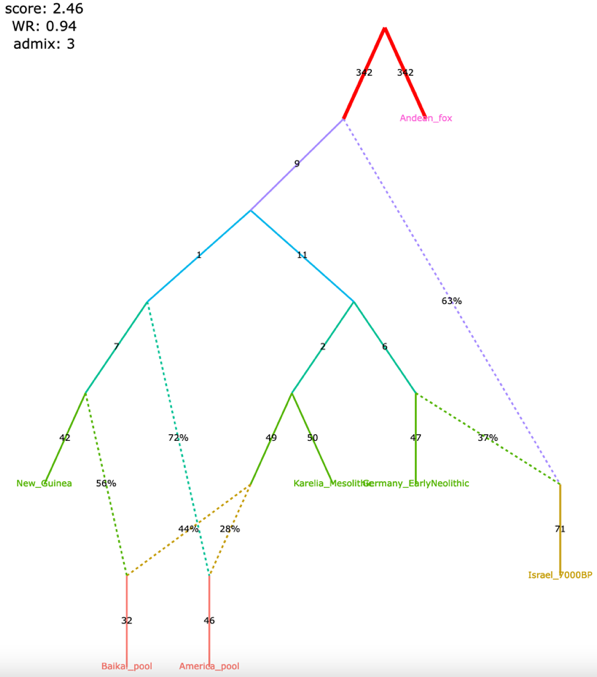

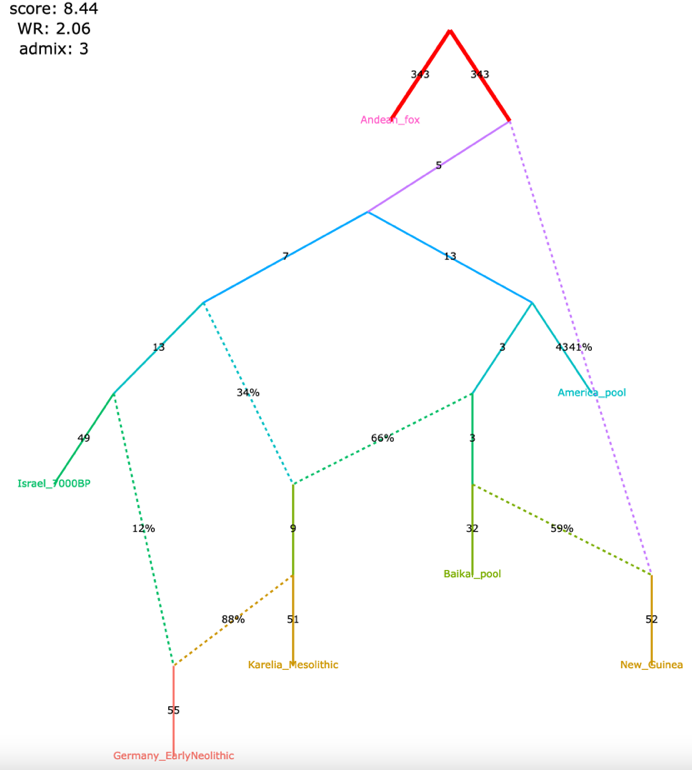

**b,** Librado *et al*. 2021 (modified	Significantly better fitting, temporally/geographically plausible.

population composition) published	In contrast to the published graph, in this graph with 8 mixture events (the minimum necessary to obtain an acceptable statistical fit to the data), a lineage maximized in horses associated with Yamnaya steppe pastoralists or their Sintashta descendants (C-PONT, TURG, or DOM2) contributes a substantial amount of ancestry to the horses from the Corded Ware archaeological context (CWC). Thus, in this model both CWC humans and horses are mixtures of Yamnaya and European farmer-associated lineages. This is qualitatively different from the suggestion that there was no Yamnaya-associated contribution to CWC horses which was a possibility raised in the paper. The eight-admixture graph also is different from the published model in that it shows a fitting model where the Tarpan horse does not have the history claimed in the study (as an admixture of the CWC and DOM2 horses).

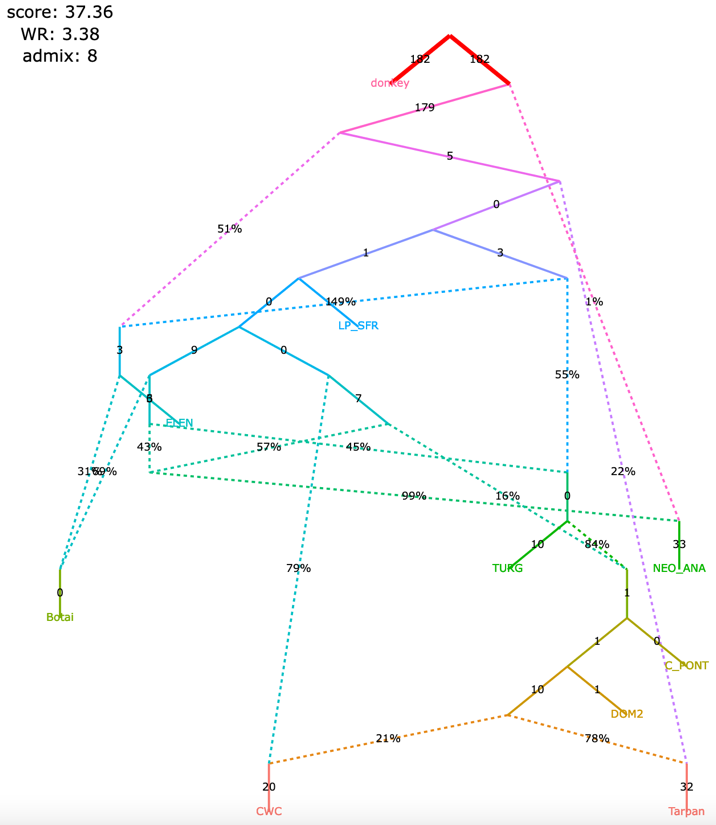

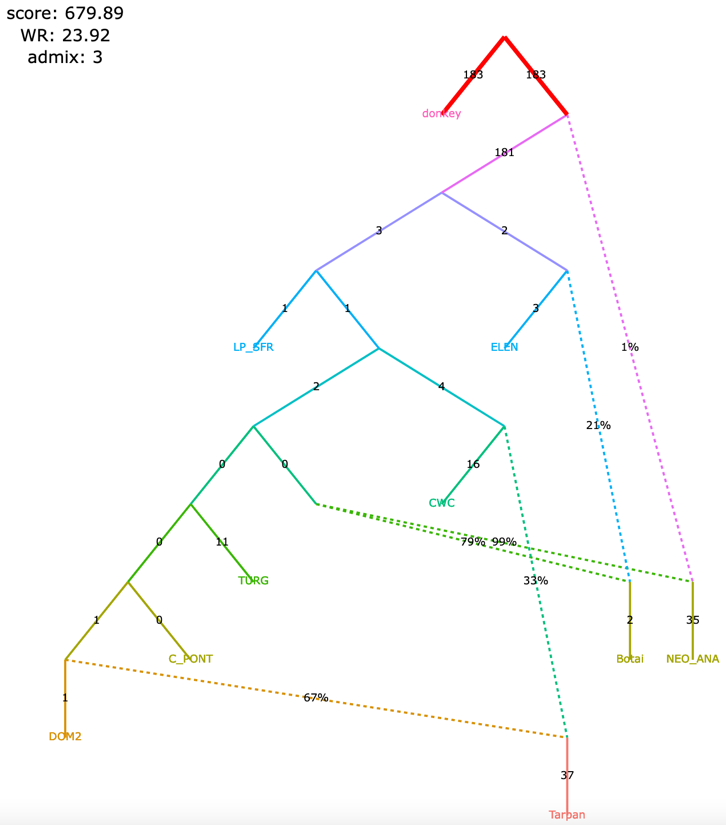

**c,** Hajdinjak *et al*. 2021 published	Significantly better fitting, but without a specific lineage shared between Bacho Kiro Initial Upper Paleolithic and East Asians. In this model, all the lineages shared between Bacho Kiro IUP and East Asians contributed a large fraction of the ancestry of later European hunter-gatherers as well, and thus this graph does not imply distinctive shared ancestry between the earliest modern humans in Europe and later people in East Asia, and instead could be explained by a quite different and also archaeologically plausible scenario of a primary modern human expansion out of the Near East contributing serially to the major lineages leading to Bacho Kiro, then later East Asians, then Ust’-Ishim, then the primary ancestry in later European hunter-gatherers.

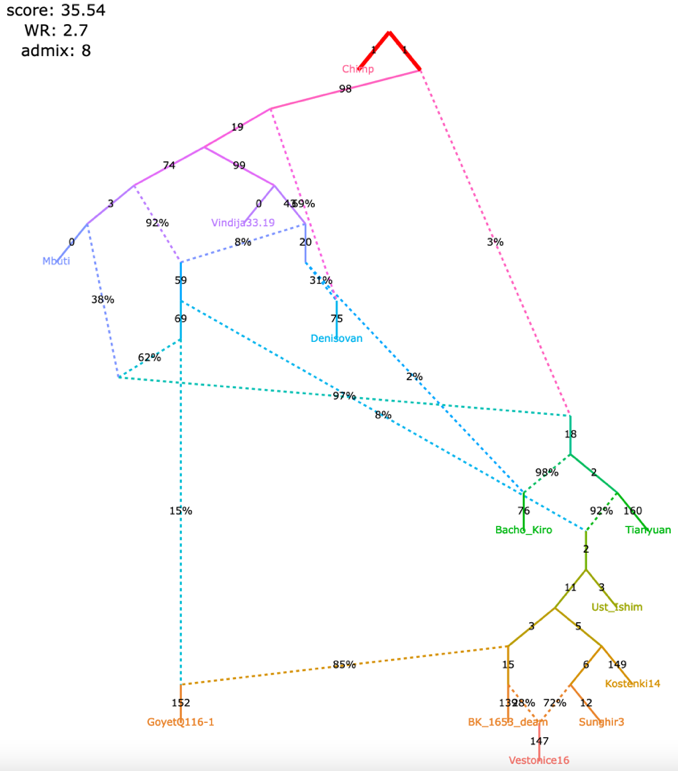

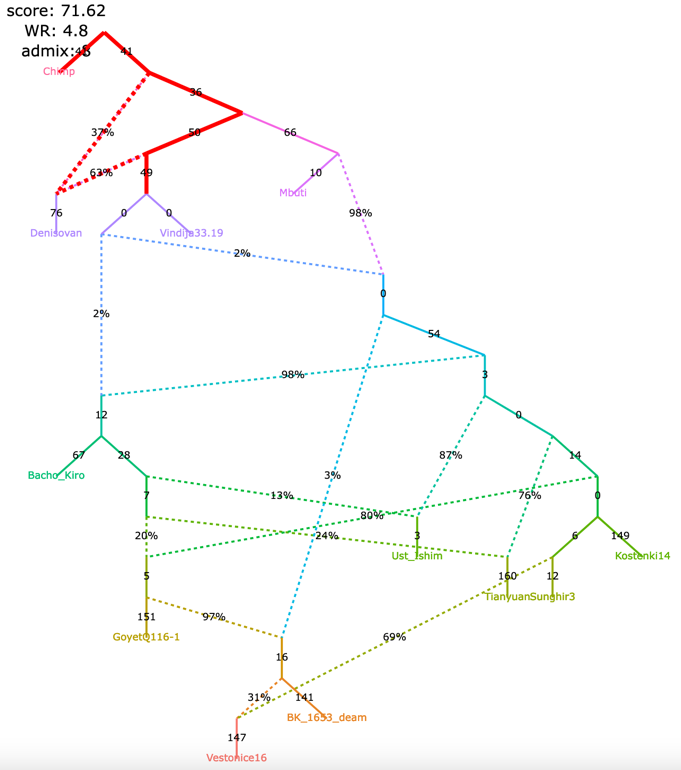

**d,** Lipson *et al*. 2020 published	Nominally better fitting. In contrast to the published graph, there is no single lineage specific to modern rainforest hunter-gatherers (Biaka and Mbuti) and Shum Laka (Cameroon_SMA). Rather, the primary ancestries in each group are separate deep-branching lineages (the deeper lineage they all share is also the source of the majority of ancestry in all anatomically modern humans modeled here). In contrast to the graph in the published paper, there is no West African-maximized ancestry present in mixed form in Biaka, Mbuti, and Shum Laka; archaic admixture is not limited to a subset of Africans, but is present in all anatomically modern humans in various proportions; and there is no ghost modern human ancestry in Agaw, Biaka, Lemande, Mbuti, Mende, Mota, Shum Laka, and Yoruba.

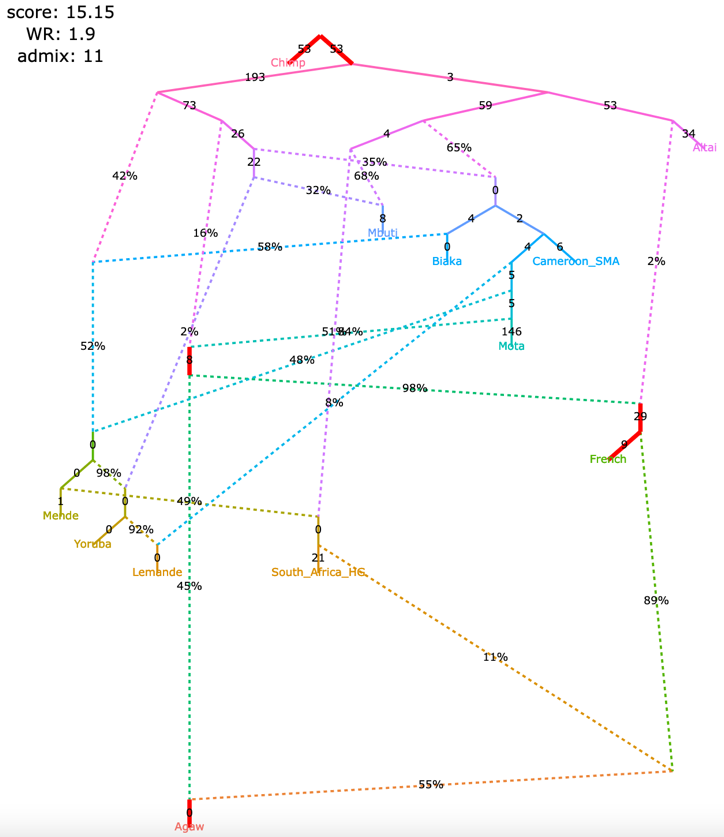

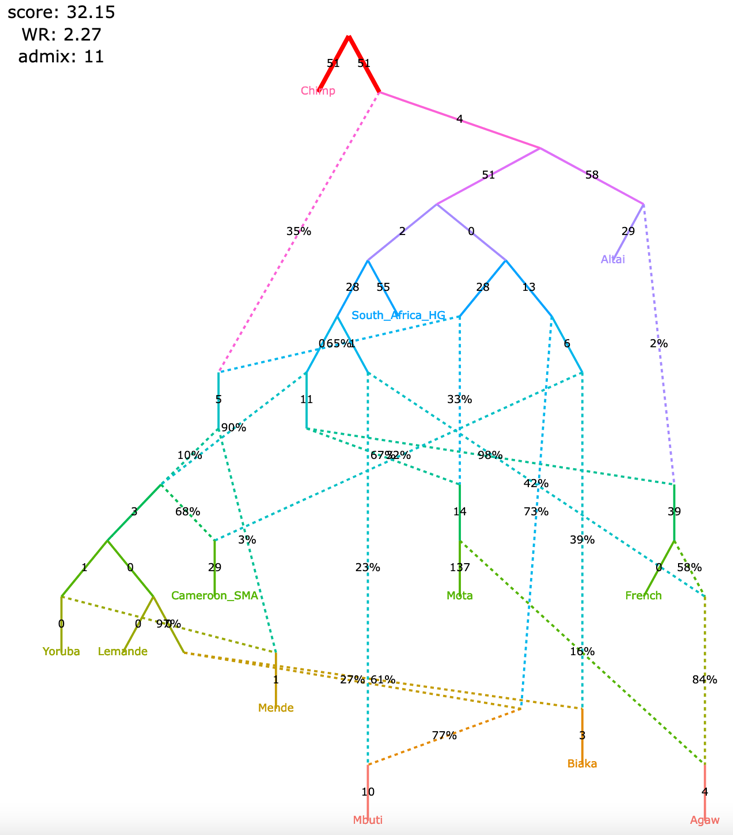

**e,** Wang *et al*. 2021 published	Significantly better fitting while meeting the constraints used to inform model-building in the published paper. The finding of Onge-related admixture that is widespread in East Asia suggesting an early peopling via a coastal route is not a feature of this model.

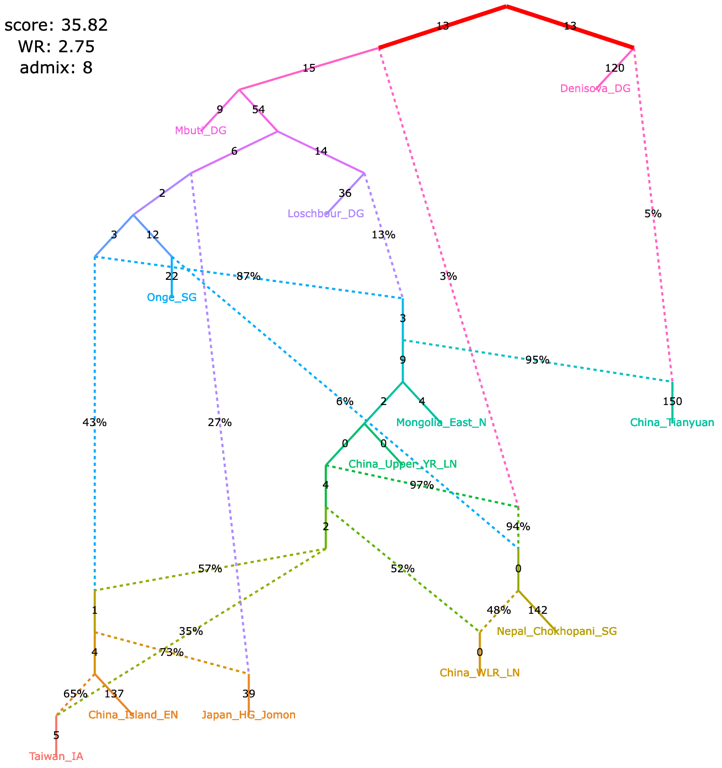

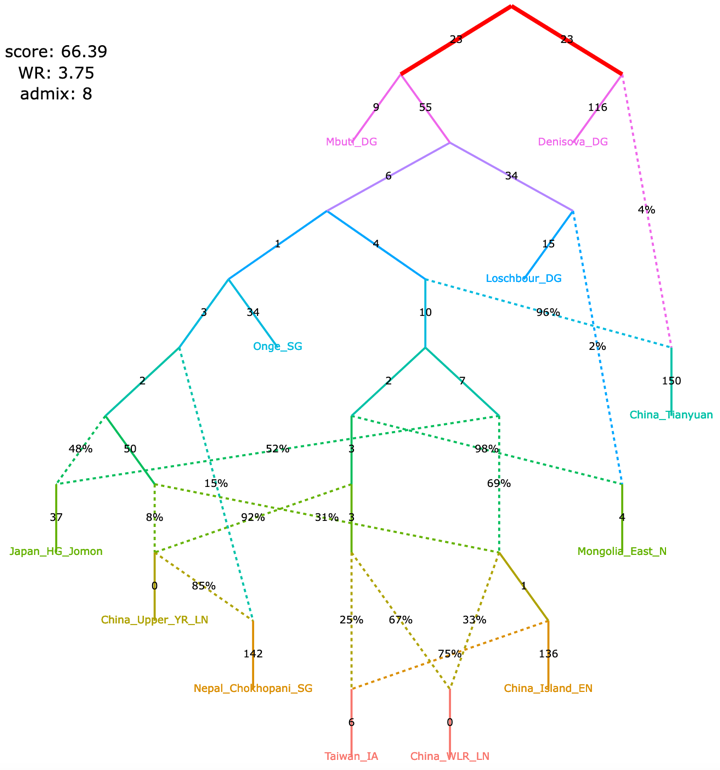

**f,** Sikora *et al*. 2019 (simplified Western graph)	Nominally better fitting. The striking feature of the admixture graph suggested in the paper whereby Mal’ta (MA1_ANE) derives some ancestry from the CHG-associated lineage is not a feature of this alternative model.

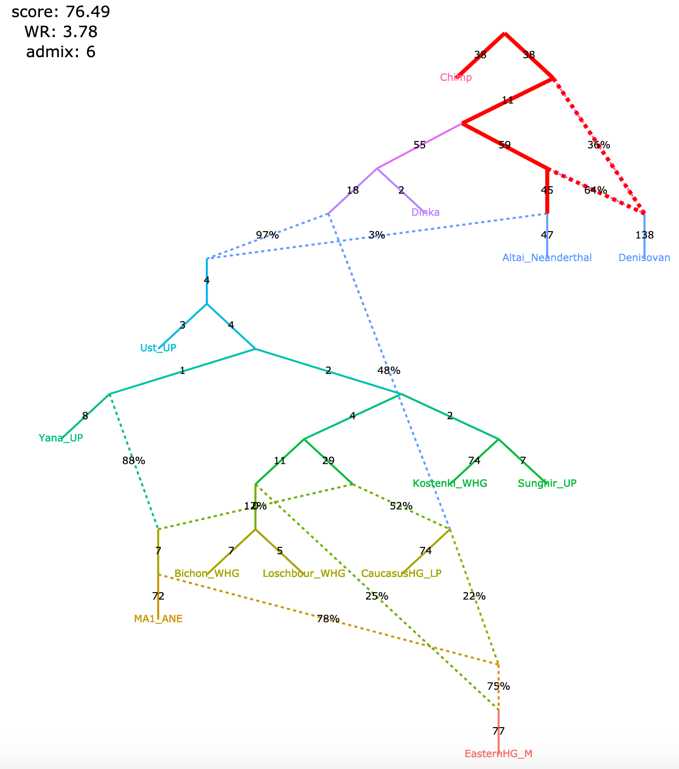

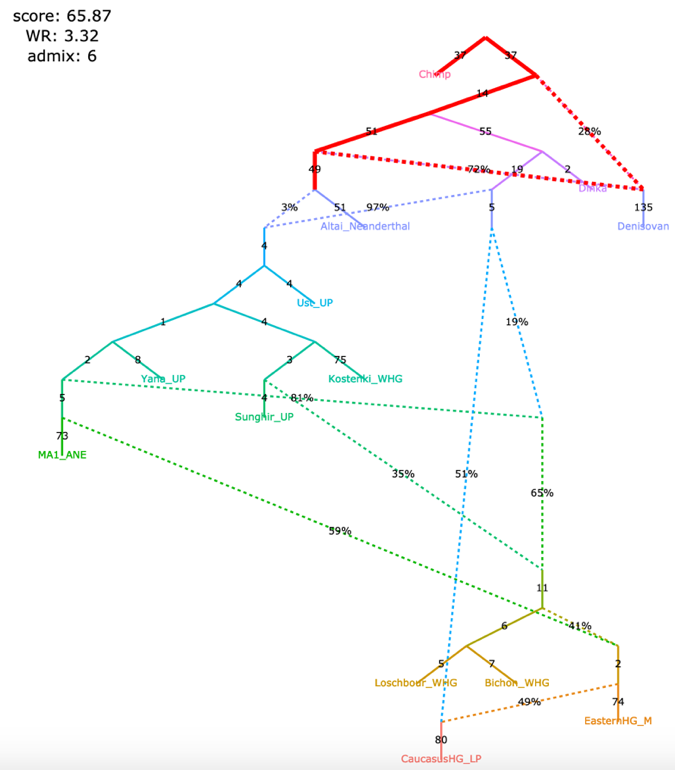

**Bergström et al. 2020.** The admixture graph for dogs in Figure 1e of Bergström et al. 2020 was inferred by exhaustively evaluating all graphs with two admixture events and outgroup ‘Andean fox’ for the six populations that remain in the graph after excluding an Early Neolithic dog from Germany. The only six-population graph with a worst residual (WR) below 3 standard errors (SE) was then chosen as a scaffold onto which the Early Neolithic dog genome from Germany was mapped, allowing for one more admixture event, and a seven-population graph with the lowest LL score was shown as the final model in the paper. Alternative six-population scaffolds were not explored in the original publication, although two six-population graphs with fits very similar to the best one were found. LL was not used as a ranking metric for alternative models in the original study; instead, the number of *f_4_*-statistics having residuals above 3 SE was considered. Since no *f_3_*-statistics were negative when all sites available for each population triplet were used (the “useallsnps: YES” option), we did not use the upgraded algorithm for calculating *f_3_*-statistics on pseudo-diploid data.

| **Study** | **Groups / admixture events** | **Features of the published model supported by temporally plausible alternative models generated by *findGraphs*** | **Features of the published model *not* supported by temporally plausible alternative models generated by *findGraphs*** |
| --- | --- | --- | --- |
| Bergström et al. 2020 | 7 / 3 * | Early divergence of domesticated dog lineages (prior to the date of the Karelian dog, 10,900 ya). | Siberian (Baikal), American, and Levantine dog lineages are unadmixed, and the West European (Germany Early Neolithic), East European (Karelia), and dogs of Southeast Asian origin (New Guinea singing dog) are admixed. |
| Lazaridis et al. 2014 | 7 / 4 | Present-day Europeans represent a mixture of three ancestral sources related to the following groups: Mal’ta (MA1), West European hunter-gatherers, and early European farmers. | N/A |
| Shinde et al. 2019 | 8 / 3 * | (1) Iranian farmer-related ancestry in the Indus Periphery group is not derived from the Hajji Firuz Neolithic or Tepe Hissar Chalcolithic groups.  (2) There is Asian-related ancestry in the Indus Periphery group. | N/A |
|  | 8 / 4 | (2) There is Asian ancestry in the Indus Periphery group. | (1) Iranian farmer-related ancestry in the Indus Periphery group is not derived from the Hajji Firuz Neolithic or Tepe Hissar Chalcolithic groups. |
| Librado et al. 2021 | 10 / 8 or 9 | (2) DOM2 and C-PONT are sister groups (they form a clade);  (4) there was gene flow from a deep-branching ghost group to the NEO-ANA group. | (1) NEO-ANA-related admixture is absent in the DOM2 group;  (3) there is no gene flow connecting the CWC group and the cluster associated with Yamnaya horses and horses of the later Sintashta culture whose ancestry is maximized in the Western Steppe (DOM2, C-PONT, TURG);  (5) Tarpan is a mixture of a CWC-related and a DOM2-related lineage. |
| Hajdinjak et al. 2021 | 12 / 8 | (3) the Vestonice16 lineage is a mixture of a Sunghir-related and a BK1653-related lineage. | (1) there are gene flows from the lineage found in the ~45,000-43,000-years-old Bacho Kiro Initial Upper Paleolithic (IUP) associated lineage to the Ust’-Ishim, Tianyuan, and GoyetQ116-1 lineages;  (2) the ~35,000-years-old Bacho Kiro Cave individual BK1653 belonged to a population that was related, but not identical, to that of the GoyetQ116-1 individual. |
| Lipson et al. 2020 | 12/ 11 | N/A | (1) A lineage maximized in present-day West African groups (Lemande, Mende, and Yoruba) also contributed some ancestry to the ancient Shum Laka individual and to present-day Biaka and Mbuti;  (2) another ancestry component in Shum Laka is a deep-branching lineage maximized in the rainforest hunter-gatherers Biaka and Mbuti;  (3) “super-archaic” ancestry (i.e., diverging at the modern human/Neanderthal split point or deeper) contributed to Biaka, Mbuti, Shum Laka, Lemande, Mende, and Yoruba;  (4) a ghost modern human lineage (or lineages) contributed to Agaw, Mota, Biaka, Mbuti, Shum Laka, Lemande, Mende, and Yoruba. |
| Wang et al. 2021 | 12 / 8 | N/A | Admixture from a source related to Andamanese hunter-gatherers is almost universal in East Asians, occurring in the Jomon, Tibetan, Upper Yellow River Late Neolithic, West Liao River Late Neolithic, Taiwan Iron Age, and China Island Early Neolithic (Liangdao) groups. |
| Sikora et al. 2019 “West” | 13 / 6 | N/A | The Mal’ta (MA1_ANE) lineage received a gene flow from the Caucasus hunter-gatherer (CaucasusHG_LP or CHG) lineage. |
| Sikora et al. 2019 “East” | 14 / 6 | (2) European-related ancestry in the Kolyma, USR1, and Clovis lineages is closer to Mal’ta than to Yana. | (1) the Mal’ta (MA1_ANE) and Yana (Yana_UP) lineages received gene flow from a common East Asian-associated source diverging before the ones contributing to the Devil’s Cave (DevilsCave_N), Kolyma (Kolyma_M), USR1 (Alaska_LP), and Clovis (Clovis_LP) lineages;  (3) the Devil’s Cave lineage received no European-related gene flows, and Kolyma has less European-related ancestry than ancient Americans (USR1 and Clovis). |

*f-statistics*

All *ADMIXTOOLS* programs are based on the statistics $f_{2}$, $f_{3}$, and $f_{4}$, for population pairs, triplets, and quadruples, respectively.

$f_{2}$ quantifies the genetic drift separating two populations $A$ and $B$. For a single SNP, it is given by $f_{2}\left( A,B \right)=\frac{1}{M}\sum_{j} (a_{j}-b_{j})^{2}$, where $a_{j}$ and $b_{j}$ are the allele frequencies for SNP $j$ in populations $A$ and $B$. When allele frequencies are estimated using a small number of samples, this estimator of $f_{2}$ will be biased upwards. An unbiased estimator of $f_{2}$ is given by
 $f_{2}=\frac{1}{M}\sum_{j} (a_{j}-b_{j})^{2}-\frac{a_{j}(1-a_{j})}{n_{A,j}-1}-\frac{b_{j}(1-b_{j})}{n_{B,j}-1}$, where $n_{A,j}$ and $n_{B,j}$ are the observed allele counts in populations $A$ and $B$.

$f_{3}(A;B,C)=\frac{1}{M}\sum_{j} (a_{j}-b_{j}{)(a_{j}-c_{j})}$is the covariance of the allele frequency differences between populations $A$ and $B$, and the allele frequency differences between populations $A$ and $C$ (assuming that alleles are coded randomly, so that $a-b$ and $a-c$ are both 0 in expectation). Significantly negative values of $f_{3}(A;B,C)$ suggest that $A$ is a mixture of sources related to $B$ and $C$ (although the converse does not hold: $A$ might be admixed between $B$ and $C$ even if $f_{3}$ is positive).

$f_{4}(A,B;C,D)=\frac{1}{M}\sum_{j} (a_{j}-b_{j}{)(c_{j}-d_{j})}$is the covariance of the allele frequency differences between $A$ and $B$, and the allele frequency differences between $C$ and $D$. Significantly positive values of $f_{4}(A,B;C,D)$ (or equivalently significantly negative values of $f_{4}(A,B;D,C)$) reveal that $A$ and $B$ do not form a clade with respect to $C$ and $D$, and that some of the drift separating $A$ from $C$ is shared with the drift separating $B$ from $D$.

$f_{3}$ and $f_{4}$ can be written as linear combinations of $f_{2}$ statistics:

$f_{3}(A;B,C)=\frac{1}{2}(f_{2}(A,B)+f_{2}(A,C)-f_{2}(B,C))$	(Eq. 1)

$f_{4}(A,B;C,D)=\frac{1}{2}(f_{2}(A,D)+f_{2}(B,C)-f_{2}(A,C)-f_{2}(B,D))$	(Eq. 2)

This implies that all $f_{3}$- and $f_{4}$-statistics can be computed from $f_{2}$-statistics as long as they are defined on the same SNPs.

$L(g) = -\frac{1}{2} (f_{3, obs}-f_{3, fit})' Q^{-1}(f_{3, obs}-f_{3, fit})$	(Eq. 3)

Here, $f_{3, obs}$ are the observed $f_{3}$-statistics and $f_{3, fit}$ are the fitted $f_{3}$-statistics. Both are vectors of length $q=\frac{k(k+1)}{2}$ for $k$ populations excluding the outgroup. $Q$ is the $q\times q$ covariance matrix of $f_{3}$-statistics, where the diagonal entries are the $f_{3}$-statistic variances, and the off-diagonal entries are the covariances for all pairs of $f_{3}$-statistics. Just like the variances (the squared standard errors), the covariances are estimated from the jackknife leave-one-block-out $f_{3}$-statistics.

$f_{2, fit} = \sum_{p \in P} \prod_{a \in p} w_{a} \sum_{e \in p} w_{e}$	(Eq. 4)

The fitted $f_{2}$-statistics are then used to obtain fitted $f_{3}$-statistics using Eq. 1.

Step 3 uses the fitted and observed $f_{3}$-statistics to estimate the likelihood score using Eq. 3.

$L(g) = -\frac{1}{2} (f_{3, obs}-f_{3, fit})' Q^{-1}(f_{3, obs}-f_{3, fit})$, with $f_{3, obs}$ and $f_{3, fit}$ defined on the same set of SNPs. The out-of-sample likelihood score is defined in the same way, except that $f_{3, obs}$ and $f_{3, fit}$ are defined on mutually exclusive sets of SNP blocks, thereby preventing any overfitting. The covariance matrix $Q$ is defined on the same set of SNP blocks as $f_{3, fit}$. As described earlier, we use block-bootstrap to fit both graphs multiple times on different SNP blocks. In each bootstrap iteration, we use all SNP blocks which are not used in fitting the graph for estimating $f_{3, obs}$.

**Supplementary Tables**

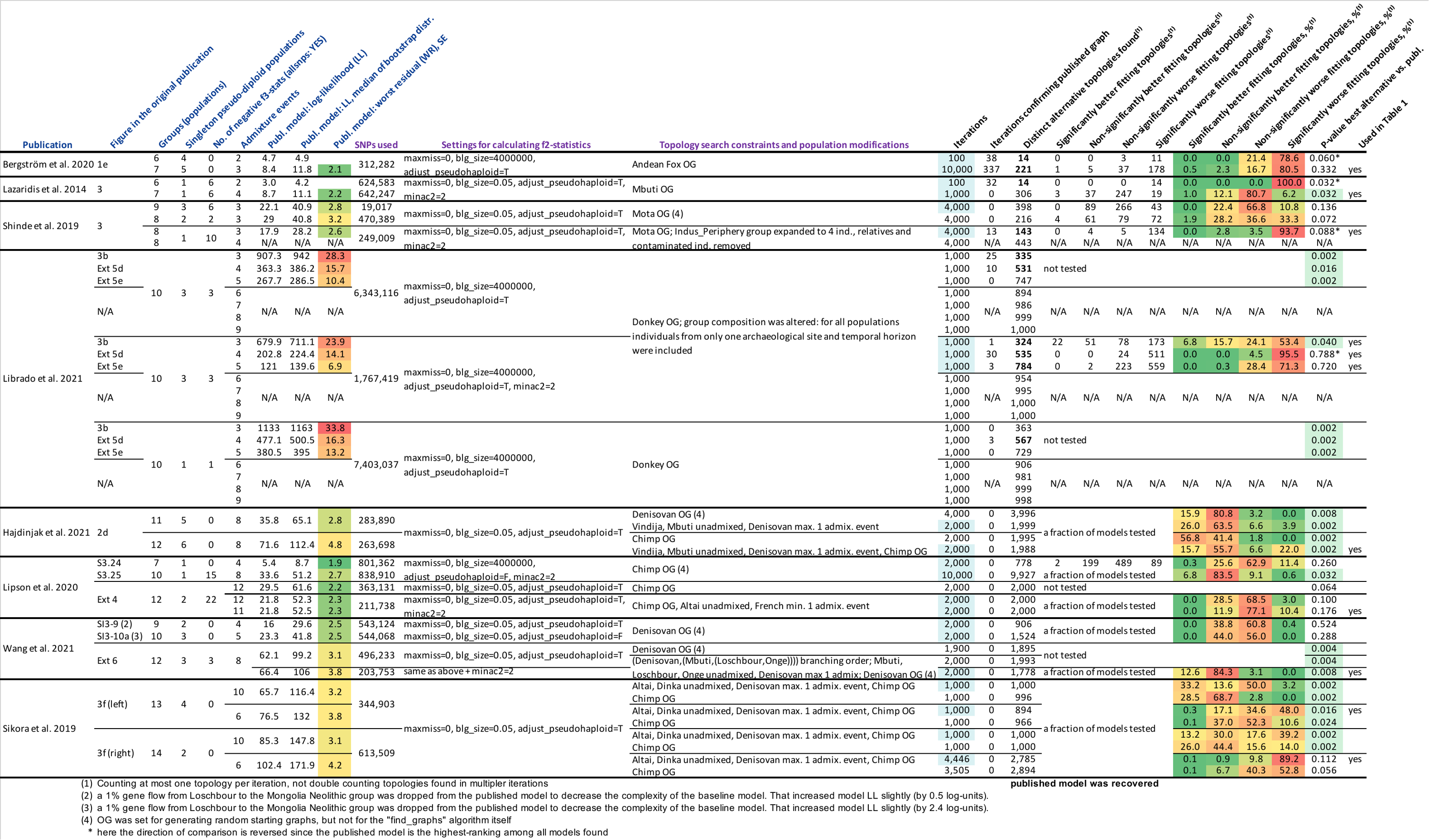

**Table S1: Published graphs in the context of automatically found graphs.** We compared 22 different graphs from 8 publications to alternative graphs inferred on the same or very similar data; these *findGraphs* runs are highlighted in blue in the “Iterations” column. In total, 51 *findGraphs* runs are summarized here since in some cases models more complex or less complex than the published one were explored and/or different population compositions were tested (see the “Topology search constraints and population modifications” column and footnotes for details). The columns with names in blue show various information on the published graphs or their modified versions and some properties of the published population sets. The columns with names in magenta show settings used for calculating *f*-statistics and for exploring the admixture graph space, and the number of SNPs used that depends on them. The columns with names in black summarize the outcomes of *findGraphs* runs, i.e., the properties of alternative model sets found.

**Used in Table 1**: Here the *findGraphs* runs featured in **Table 1** are marked.

**Table S2: Statistics for shotgun sequencing of individual I8726**

| Individual ID | I8726 |
| --- | --- |
| Archaeological IDs | SHAR_201 (Grave 201) |
| Skeletal element | Petrous bone |
| Location | Seistan, Shahr-i-Sokhta, Iran |
| Archaeological context date | 3100-3000 BCE |
| Latitude | 30.649857 |
| Longitude | 61.400311 |
| First publication of library | Narasimhan, Patterson et al. Science 2019 |
| Library ID | S8726.E1.L1 |
| HiSeqX10 lanes for shotgun sequencing | 3 |
| Molecular sex | Male |
| Mean coverage measured on 1240k autosomal targets | 2.60306 |

**Supplementary Items Available as Separate Files**

**5 supplementary tables are available as separate files (Tables S3-S7)**

**13 supplementary figures are available as separate files (Figures S1-S13)**
